# The Regulatory Logic of Planarian Stem Cell Differentiation

**DOI:** 10.1101/2024.08.23.608747

**Authors:** Alberto Pérez-Posada, Helena García-Castro, Elena Emili, Virginia Vanni, Cirenia Arias-Baldrich, Siebren Frölich, Simon J. van Heeringen, Nathan Kenny, Jordi Solana

## Abstract

Cell type identity is determined by gene regulatory networks (GRNs), comprising the expression of specific transcription factors (TFs) regulating target genes (TGs) via binding to open chromatin regions (OCRs). The regulatory logic of differentiation includes factors specific to one or multiple cell types, functioning in a combinatorial fashion. Classic approaches of GRN discovery used perturbational data to elucidate TF-TG links, but are laborious and not scalable across the tree of life. Single cell transcriptomics has emerged as a revolutionary approach to study gene expression with cell type resolution, but incorporating perturbational data is challenging. Planarians, with their pluripotent neoblast stem cells continuously giving rise to all cell types, offer an ideal model to attempt this integration. Despite extensive single cell transcriptomic studies, the transcriptional and chromatin regulation at the cell type level remains unexplored. Here, we investigate the regulatory logic of planarian stem cell differentiation by obtaining an organism-level integration of single cell transcriptomics and single cell accessibility data. We identify specific open chromatin profiles for major differentiated cell types and analyse their transcriptomic landscape, revealing distinct gene modules expressed in individual types and combinations of them. Integrated analysis unveils gene networks reflecting known TF interactions in each type and identifies TFs potentially driving differentiation across multiple cell types. To validate our predictions, we combined TF knockdown RNAi experiments with single cell transcriptomics. We focus on *hnf4*, a TF known to be expressed in gut phagocytes, and confirm its influence on other types, including parenchymal cells. Our results demonstrate high overlap between predicted targets and experimentally-validated differentially-regulated genes. Overall, our study integrates TFs, TGs and OCRs to reveal the regulatory logic of planarian stem cell differentiation, showcasing that the combination of single cell methods and perturbational studies will be key for characterising GRNs widely.

## Introduction

Gene regulation underlies many cellular decisions including cell fate and identity. Pluripotent stem cells undergo distinct molecular changes as they differentiate into mature cell types, including changes in gene expression and chromatin dynamics ^1^. These changes involve different kinds of genetic regulators, such as chromatin remodelers and transcription factors (TFs). The former play a critical role in the functioning of the latter, as chromatin marks and accessibility facilitate the binding and interaction of TFs with chromatin. Transcription factors are often pleiotropic, functioning in multiple cell types, stages or conditions in context-specific ways depending on the co-expression of other factors ^2-4^. Ultimately, these factors orchestrate the transcription of specific targets, thereby determining cell type identity. Thus, cell differentiation comprises the combined expression of TFs and the combined accessibility of open chromatin regions (OCRs) acting as cis-regulatory elements (CREs). This combination creates a ‘regulatory logic’ forming gene regulatory networks (GRNs) ^5-7^. While the general dynamics of this process have been studied in a number of model species ^8-11^, the mechanisms governing cell differentiation into various lineages remain largely unexplored in most multicellular organisms.

Single-cell methods have democratised the study of differentiation trajectories in a variety of animal species ^12-14^. The initial step in characterising the potential differentiation pathways of pluripotent stem cells consists in identifying their distinct differentiation products, a task accomplished through single-cell transcriptomics (scRNA-seq) ^15, 16^. This technique enables the identification of expressed transcripts within individual cells, allowing for the grouping of cells into specific cell types. Computational algorithms are then employed to reconstruct the transitional states between stem cells and each differentiated cell type ^17-19^. However, despite the ability to characterise the expression of transcripts, uncovering the GRNs governing their activity remains challenging.

Recently, novel single-cell methods have emerged to characterise the chromatin state to reveal OCRs and CREs. These methods leverage assay for transposase-accessible chromatin with sequencing (ATAC-seq) ^20, 21^, which identifies OCRs, including the enhancers and promoters that play a pivotal role in transcriptional regulation. Single-cell ATAC-seq (scATAC-seq) has been successfully employed in various models and paradigms ^22-28^. One major challenge lies in integrating scATAC-seq data with scRNA-seq data and extracting regulatory information from the combination of chromatin accessibility and expression data ^29, 30^. Recent single-cell technologies predict TF/target gene interactions across various contexts ^31, 32^ but often lack experimental validation, and it is unclear if these methods can scale to the whole organism level.

Planarians are an ideal model organism to address this challenge as they have a pluripotent stem cell type in their adults that constantly differentiates to replace aged cells of all cell types ^33, 34^. A single planarian stem cell can differentiate into all cell types of the adult worm ^35^. These cells also enable planarians’ amazing regenerative capacities ^36-38^. Transcription factors and epigenetic regulation have been already studied in planarians ^39-43^. Using scRNA-seq, the major differentiated cell types that mature from planarian stem cells have been described ^44, 45^. Planarians are also very amenable to gene knockdown by RNAi ^46^. Single-cell analysis techniques hold significant potential for investigating RNAi knockdown experiments, but there are still several challenges that need to be addressed ^47-50^. Cell dissociation techniques can trigger stress responses and introduce biases, resulting in cell death and variations in cell survival rates ^51-53^. Additionally, including different samples with current methods can introduce batch effects ^54^. However, fixative cell dissociation approaches like ACME can mitigate the first concern by minimising stress-induced effects ^55^. Moreover, combinatorial single-cell transcriptomic approaches like SPLiT-seq enable sample multiplexing and facilitate convenient multi-sample experiments ^56^. By combining ACME and SPLiT-seq, it becomes possible to analyse multi-sample experiments, such as RNAi knockdown studies, with greater efficiency and accuracy ^57, 58^.

Here we report the first integration of scRNA-seq and scATAC-seq in *Schmidtea mediterranea*, in a whole adult organism. We combined 98,363 single cell transcriptomes with 3,659 single cell ATAC profiles. Using the graph-based correlational tool WGCNA we predicted gene sets and OCRs active in one or more broad types. We identified key transcription factors involved in the differentiation of all major cell lineages of planarian stem cell differentiation. We predicted TFs influential in each broad cell type, and their targets, using ANANSE, a graph analysis computational approach ^59^. Finally, to validate our findings, we performed RNAi of *hnf4* coupled with single-cell analysis. Altogether, our experiments reveal the regulatory logic of planarian stem cell differentiation. Our results underscore that the characterisation of all differentiation trajectories and the GRNs that underlie them is possible combining single cell methods and perturbation experiments with single cell resolution.

## Results

### An integrated atlas of planarian stem and differentiated cells

To understand the regulation of differentiation from pluripotent stem cells to all adult cell types in planarians we generated an integrated multimodal single cell atlas with scRNA-seq and scATAC-seq data. We compiled previously generated datasets ^55, 57^ as well as newly generated experiments using ACME and SPLiT-seq (Figure 1A, Supplementary File 1). On the other hand, we used Trypsin dissociation and the 10X Genomics commercial approach to obtain a scATAC-seq dataset (Figure 1A). We mapped these datasets to the recently released version of the *Schmidtea mediterranea* genome ^33^, (Supplementary File 2). To analyse scATAC-seq data and obtain cell clusters we used CellRanger ^60^ and Seurat ^61^. This allowed us to obtain a dataset with 3,659 cells distributed in 11 clusters (Figure 1B). We processed SPLiT-seq data with our analysis pipeline ^55, 62^ and Seurat, to obtain a total of 98,363 cells in 59 cell clusters. We elucidated their identity with a previously published dataset (Figure 1B, Supplementary Figure 1, Supplementary File 3) ^57^. The average numbers of UMIs and genes quantified per cell remain low, as characteristic of SPLiT-seq (Supplementary Figure 2AB). However, the integrated approach increases the total number of reads in each cluster, and therefore, the total number of genes quantified (Supplementary Figure 2C). We then integrated the scRNA-seq and scATAC-seq datasets to obtain identities for the scATAC-cluster (Figure 1B-C, Supplementary Figure 3, Supplementary File 3) using canonical-correlation analysis (CCA) ^63, 64^. This approach leverages the correlation between the expression data and the scATAC signal detected within gene bodies. While the scATAC-seq data was shallower and less resolved than the scRNA-seq, the assay managed to detect open chromatin profiles for all major planarian broad cell types ^44, 45^, including neoblasts, three stages of epidermal differentiation, phagocytes, basal/goblet cells, muscle, neurons, parenchymal cells, protonephridia and secretory cells. We grouped scRNA-seq clusters in 11 corresponding broad groups using this integration data (Supplementary File 3, Supplementary Figure 3). To identify genes with both open chromatin and gene expression specific to each cluster we cross-referenced the markers of both datasets (Figure 1D, Supplementary Figure 4). We inspected the genomic regions identified, with their associated gene annotations and open chromatin peaks (Figure 1E). We then obtained genomic coverage tracks of the scATAC-seq (Figure 1F) and scRNA-seq (Figure 1G) signal of these regions, for each broad group. This analysis included a bulk ATAC-seq sample that showed good agreement with the scATAC-seq (Supplementary Figure 5). Altogether, these analyses revealed genes with both open chromatin features and RNA expression and validated the quality of the scATAC-seq data. This shows our integrated multimodal dataset captures the transcriptomic and epigenomic landscape of each planarian differentiated broad type.

**Figure 1.**
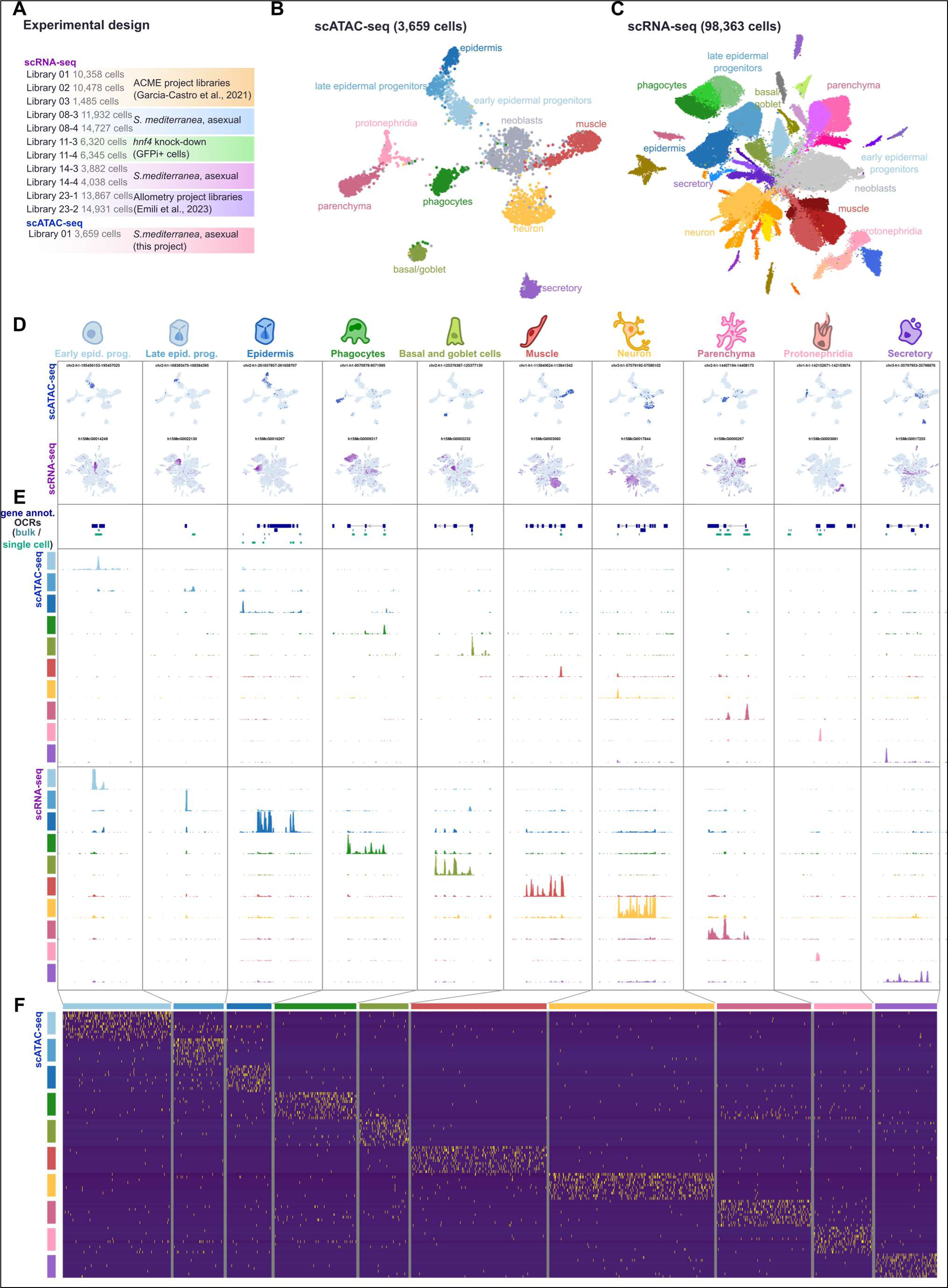
An integrated atlas of planarian differentiated cells. A: Summary of the experimental design of the different libraries. B: UMAP projection of 3,659 individual cellular profiles of chromatin accessibility in *S. mediterranea*. D: Feature plots of ten different open chromatin regions (up) and genes (down) specific to non-pluripotent broad cell types in *S. mediterranea*. E: Overview of specific gene expression and chromatin accessibility of each broad cell type, for the same features as in D. Top: Gene annotation and open chromatin regions detected in bulk and single cell ATAC-seq. Middle: genomic tracks of scATAC-seq of each cell type. Bottom: genomic tracks of scRNA-seq of each cell type. F: Heatmap of markers of scATAC-seq cells.

### The transcriptomic landscape of planarian stem and differentiated cells

Gene expression is highly pleiotropic. Some genes are expressed broadly in all cell types and tissues. Others are very highly specific to one cell type. Often, genes are expressed specifically in multiple cell types ^65, 66^, and it is the combination of genes expressed in each cell type that defines their identity.

In single-cell analysis, marker finding algorithms usually perform one-against-all comparisons. This approach can reveal genes very specific to any one cell type, but potentially fail to reveal genes with expression specific to multiple cell types. One approach that can overcome this limitation is Weighted Gene Coexpression Network Analysis (WGCNA) ^67, 68^ as it detects modules of correlated genes regardless of their correlation being in one or more cell types (Supplementary Note 1). As a pseudobulk approach, using WGCNA in our integrated scRNA-seq data leads to high sensitivity, despite the relative shallowness of individual single-cell data points. We obtained a total of 24 modules with average expression peaking in one single cell type (s modules), and interestingly, 53 modules with expression peaking in multiple cell types (m modules) (Figure 2A, Supplementary Figure 6, Supplementary File 4) (See Methods). These included modules made of genes with expression in similar cell types (e.g. in two or more types of neurons) but also modules expressed in distinct broad cell types (Supplementary Files 4, 5, 6, 7). As examples of modules with genes expressed in similar types, modules m08 and m23 contained genes expressed broadly in phagocytes and their progenitors, module m24 contained genes expressed broadly in all muscle types, module 51 contained genes expressed broadly in secretory types, and modules 32 and 33 contained genes expressed in several parenchymal cell types. Interestingly, many modules had peak expression in several distinct cell types. For instance, we identified modules with expression in neuronal and muscle types (m26, m27), modules with expression in the pharynx cell type and *psd+* cells (m19), which are both pharyngeal, and modules containing cilia genes (m50 and m53) with expression in epidermal, neuronal and protonephridial types. The most prominent feature was the presence of several modules with genes expressed in both gut cell types and parenchymal cells, including m37, m44, m45, m46 and m49. The latter module contained genes enriched for GO terms related to amino acid transmembrane transport metabolism, highlighting possible shared functions of these cell types. Our observations indicated that gut and parenchymal broad types are strongly associated in their gene expression patterns. Taken together, our WGCNA analysis revealed modules of genes expressed in multiple distinct cell types, revealing their similarities in gene expression.

**Figure 2.**
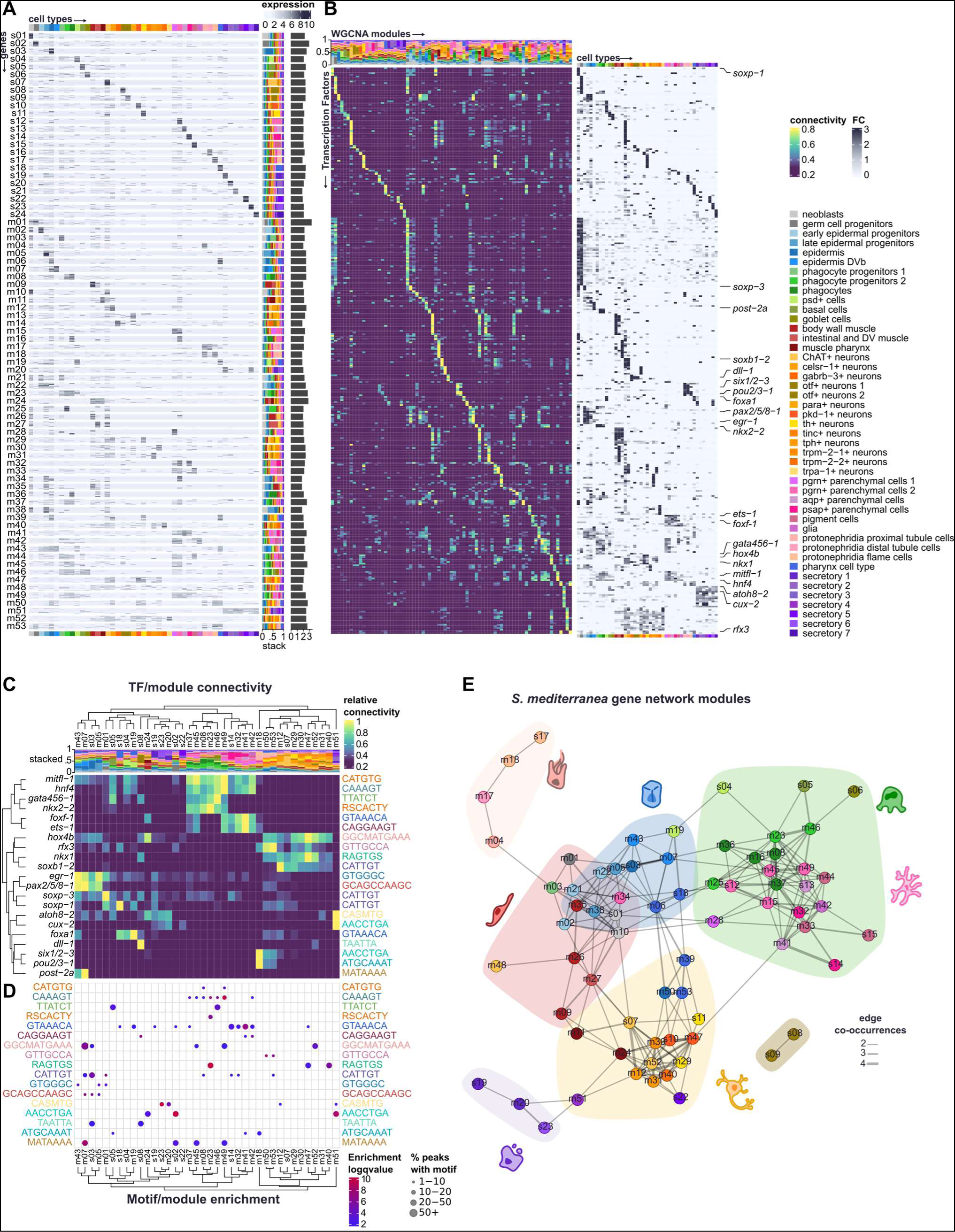
The transcriptomic landscape of planarian stem and differentiated cells. A: Heatmap of gene expression of genes from different WGCNA co-expression modules across cell types in *S. mediterranea*. Genes in rows, cell types in columns. A random subset of thirty genes per co-expression module is shown. Module identity labels on the left side. On the right side, stacked barplot of average gene expression of each co-expression module in each cell type (see Methods; Supplementary Note 1). On the right-most side, bar plot showing log number of genes per co-expression module. B: (left) Heatmap of Transcription Factor (TF) connectivity to each co-expression module in *S. mediterranea*. TFs in rows, modules in columns. TFs have been sorted by highest connectivity value. On the top, stacked barplot of average gene expression as explained in A. (right) Heatmap of gene expression fold change of the same TFs on each cell type. TFs in rows, cell types in columns. TFs follow the same order as in the heatmap to the left. Several TFs known in the literature are highlighted. C: Subset of left heatmap in B showing connectivity of well-known TFs in the literature to different co-expression modules in *S. mediterranea*. Right side, annotation of core sequence boxes of associated motifs (JASPAR database) D: Motif enrichment analysis on the promoters of the same co-expression modules as shown in C. Motifs (core boxes) in rows, modules in columns. Modules have been arranged in the same order as the heatmap in C. Motifs have been arranged following the same order of the core boxes associated to the TFs (rows) in the heatmap from C. E: Module-wise network of modules in *S. mediterranea*. Nodes represent co-expression modules, edges represent connections between them. Edge width indicates the number of occurrences of a given module-module connection in different analyses. Shaded areas represent module communities.

To understand the regulatory logic of these modules of gene coexpression, we then annotated transcription factors. Using an approach combining sequence homology, identification of DNA binding protein domains and literature curation (expanding on Neiro *et al*. ^39^ and similar to King *et al*. ^69^) we annotated 665 TFs (Supplementary File 2) and identified a set of 517 TFs with cell type specific expression in our scRNA-seq dataset. These TFs showed high connectivity (measured as the correlation to the average expression of all genes in each module) to one or more of the WGCNA modules we detected (Figure 2B); some of them were more highly connected to mixed modules than to specific modules, as we corroborated by investigating their expression profile (Figure 2B). For example, we detected high connectivity of well-known TFs such as *foxa* ^70^, *gata456-1* ^71^, and *foxf-1* ^72^, in modules m19, m46 and m41, respectively (Figure 2B; labels on the right side). Interestingly, the highest connectivity of transcription factor *hnf4* was to module m49, a mixed module of genes expressed in gut and parenchymal cells (Figure 2B). To further explore these agreements between TFs and gene modules, we performed motif enrichment analyses in the promoters of the genes of each respective module (Figure 2C, Supplementary File 8). We observed that promoters of the genes in these modules had enrichment for DNA motifs that have been previously associated with TFs and/or TF classes with high expression in the same cell types of the modules with high connectivity. For example, we found enrichment of CTGGGGC-box motifs associated with *egr* transcription factors ^73, 74^ in epidermal modules, to which *egr1* is highly connected. Similarly, the pou-associated box ATGCAAAT ^75, 76^ is enriched in the m18 protonephridia module to which *pou2/3-1* is connected. Finally, the CAAAGT box, associated with nuclear receptor TFs such as *hnf4* ^35, 43, 77^, appeared enriched in modules of gut and parenchymal cells (Figure 2C).

To further explore the dynamics of these modules we analysed their cross-connections (i.e the number of connections between genes of different modules), similarity in motif enrichment, similarity in functional category enrichment between modules, and the overall profile of TF connectivity of each module (Supplementary Figure 7 Supplementary Note 1). We retrieved similar connections between modules across all these analyses (Figure 2D). For example, and most prominently, parenchymal and gut modules were highly connected within themselves but they also shared many cross-connections. We also observed connections between muscle, secretory and neuronal modules, and also between neuronal, pharynx, and cilia modules.

Overall, our analyses suggest the existence of several major programmes of gene expression controlled by similar groups of TFs. As predicted, many of these were specific to one cell type but others were specific to multiple cell types. Our methods showcase the capability of single cell transcriptomics to elucidate the transcriptomic landscape of planarian cell types.

### Single cell ATAC-seq analysis reveals the chromatin accessibility landscape of planarian cells

Based on the idea that genes are expressed in multiple cell types and likely controlled by multiple TFs, we wondered if similar patterns were also observable at the chromatin level. To investigate this, we examined chromatin accessibility dynamics across cell types using single-cell data to generate pseudoreplicates ^78^. We split our sc-ATACseq data in two pseudoreplicates to perform differential chromatin accessibility analysis for each cell type (Figure 3A, Supplementary Figure 8A, Supplementary Files 9, 10). We first compared every cell type against the rest of cell types in order to identify cell type-specific open chromatin regions (OCRs). The resulting sets of differentially accessible OCRs showed similar motif enrichments to those detected in our WGCNA analysis (Figure 3B), supporting our previous observations. Our method reliably detected differentially accessible OCRs for every broad cell type except for neoblasts (Figure 3C). Moreover, marker analysis also revealed regions and motifs specific to each broad type, except for neoblast, which raised regions generally open in all cell types, as if they were constitutive regions (Supplementary Figure 8BC). To further corroborate these points, we selected markers of neoblasts using the scRNA-seq dataset, and observed their activity in the scATAC dataset. This analysis revealed that their signal was not neoblast-specific either (Supplementary Figure 8D). The OCRs derived by our differential chromatin accessibility analysis are distributed in chromatin accessibility peaks (Supplementary Figure 8E). These findings align with previous observations that neoblasts lack a specific chromatin signature ^40^.

**Figure 3.**
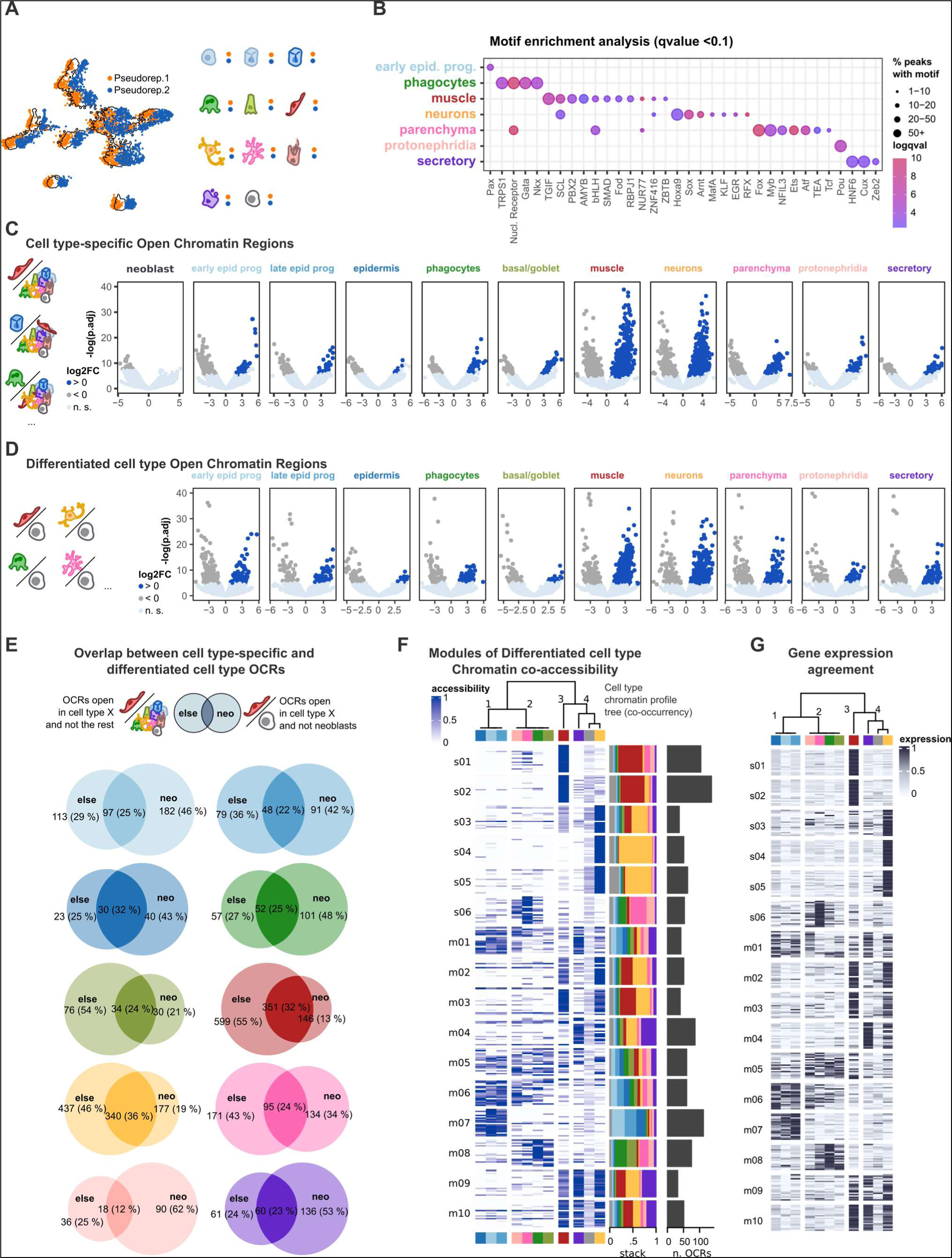
Chromatin dynamics of pluripotent and differentiated planarian cells. A: Graphical overview of the splitting of the scATAC-seq into pseudo-replicates. B: Motif enrichment analysis in cell type-specific (one-vs-all) sets of differentially accessible open chromatin regions (OCRs). C: Volcano plots showing log fold change (x-axis) and -log p. adjusted (y axis) of each cell type-specific (one-vs-all) differential chromatin accessibility analysis, one for each broad cell type. D: Same as C, for each of the differentiated cell type (one-vs-neoblasts) differential chromatin accessibility analysis. E: Venn diagrams of the overlap between cell type-specific and differentiated cell type OCRs. F: Heatmap of chromatin accessibility of differentiated cell type OCRs from different WGCNA co-accessibility modules across broad cell types in *S. mediterranea*. OCRs in rows, cell types in columns. A random subset of twenty OCRs per module is shown. On the right side, stacked barplot of average chromatin accessibility of each co-accessibility module in each broad cell type (see Methods; Supplementary Note 1). On the right-most side, bar plot showing number of OCRs per co-accessibility module. G: Heatmap of gene expression of genes associated to OCRs from different co-accessibility modules as in F. Genes in rows, cell types in columns. A subset of top-correlating genes is shown (see Supplementary Note 1 for details).

In addition to this one-versus-all approach, we also compared chromatin accessibility of every cell type against neoblasts in order to retrieve differentiated cell type OCRs (Figure 3D). This allowed us to retrieve a second set of differentially accessible OCRs for every major cell type which partially agreed with our cell type-specific chromatin regions (Figure 3E, Supplementary Figure 9A, Supplementary Files 9, 10), suggesting our second approach was detecting a different set of OCRs that may be accessible in other cell types and which we also observed by visual inspection of some of these OCRs (Supplementary Figure 9B). We performed a co-occurrence analysis that confirmed previously observed patterns, with strong associations between epidermal types, gut and parenchymal types, as well as muscle, neuron and secretory types (Supplementary Figure 9C). We further explored this by performing a second WGCNA analysis using these differentiated cell type OCRs as input, which successfully clustered the OCRs into modules of co-accessible regions in different cell types (Figure 3F, Supplementary Figure 9D, Supplementary File 11). These modules contained OCRs predominantly open in one cell type, such as muscle (modules s01 and s02) or neuronal cells (modules s04 and s05), but also modules of accessibility specific to multiple cell types. For example, we detected three modules of OCRs accessible in parenchymal and gut cells (s06, m05, m08), and also modules of OCRs open in muscle, neuron, and secretory cells (m02, m03, m09, m10). These combinations of cell types match the profiles of genes with expression specific to multiple cell types we observed in previous analyses (Figure 2A). To ascertain if these patterns of accessibility specific to multiple cell types translated into expression in these multiple cell types we associated each OCR to their closest gene model and analysed their gene expression patterns (Supplementary File 12). We visualised as a heatmap the genes with highest correlation between accessibility and expression (Supplementary Figure 9E-F, Figure 3G), which showed high agreement. This analysis shows that many genes associated with these OCRs were also expressed in the same cell types. We also used our OCR-closest gene association to investigate whether genes from our co-expression WGCNA modules had associated OCRs that are accessible in the same cell types as said WGCNA modules. For this we did a stacked barplot of the composition of chromatin co-accessibility WGCNA modules of associated OCRs to the genes of each WGCNA module (Supplementary Figure 9G). We observed a general trend of genes and OCRs expressed and accessible, respectively, in the same cell types. For example, co-expression modules s15-16 are composed of genes expressed in parenchyma and protonephridia cells, and these are associated to OCRs from co-accessibility module s01 which is accessible in muscle but also parenchyma and protonephridia cells (Supplementary Figure 9H). Likewise, co-expression modules m30-m33 are composed of genes expressed in neurons, and these are associated to OCRs from co-accessibility modules s03, s04, s05, m02, and m03 which are accessible in neurons. Another example are co-expression modules m41, m42, m44, m45 and m49 which are composed of genes expressed in gut and parenchymal cells and are associated to OCRs from parenchyma and gut co-accessibility modules s06 and m08. Together, these agreements suggest these OCRs could be part of the regulatory logic driving the expression of these genes in each cell type. Overall our approach retrieved complementary sets of OCRs for every differentiated cell type in planarians, which potentially account for the regulatory programmes of gene expression in the major planarian cell types.

### ANANSE network analysis predicts influential TFs for planarian cell fate specification

Recent efforts in the community have sought to integrate multimodal data such as gene expression and chromatin dynamics to establish associations between TFs and target genes (TGs) via binding of TFs to regions of open chromatin, and to study these regulatory programmes at the network level ^7, 79, 80^. To combine gene expression and chromatin accessibility data, we used ANANSE ^59^, a tool that leverages gene expression, chromatin accessibility, distance from the Transcription Start Site (TSS), and motif enrichment analysis, in an additive model to create networks of TFs and TGs, assigning a score to the interaction between each TF-TG (Figure 4A). ANANSE uses motif databases and orthology assignment to associate motifs to TFs (Supplementary File 13, Supplementary Figure 10A, Supplementary Note 2). We aggregated our scRNA-seq and scATAC-seq in pseudobulk data to isolate the gene expression and chromatin signal of every independent broad cell type, to generate a network for each, for a total of eleven networks of TF-target genes (Supplementary Figure 10B). We pruned these networks for lowly-scored interactions and constructed TF-TG graphs which we used to calculate a centrality score (Supplementary Note 2) for each TF in each network. Correlating these profiles of TF centrality for each network revealed that epidermal cell types clustered together, as expected, and that the most similar cell type to gut phagocytes were the parenchymal and basal/goblet cells (Figure 4B, Supplementary Figure 10C). This suggests that not only their gene expression and chromatin accessibility patterns are similar, but also their transcription factor-based regulation.

**Figure 4.**
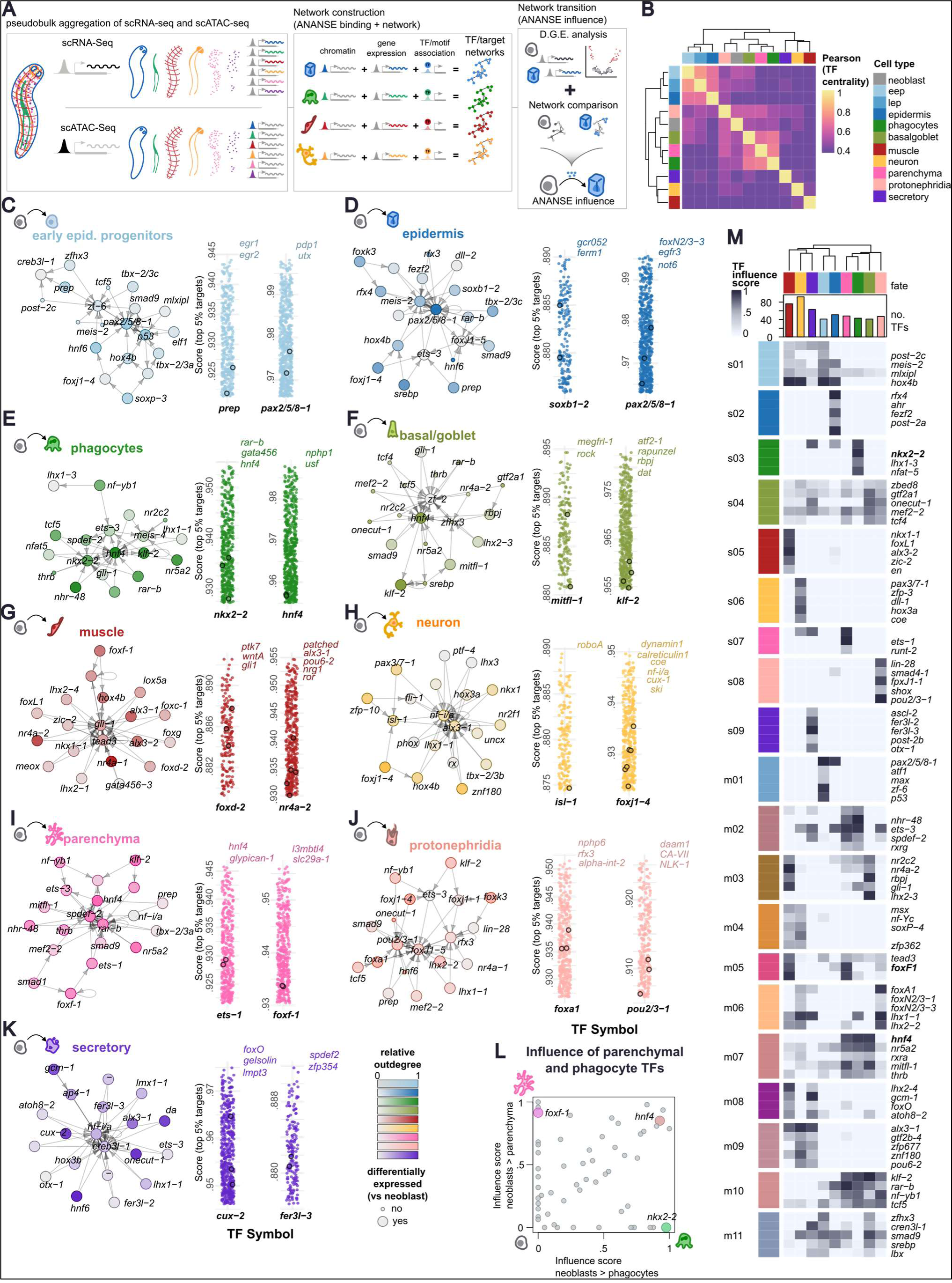
TF/target gene regulatory networks of planarian cell differentiation. A: Graphic summary of ANANSE. B: Pearson correlation of cell types based on profile of TF centrality across networks. C-K: (left) differential network and (right) top targets of two example TFs of the differential network when comparing neoblasts to (C) early epidermal progenitors, (D) epidermis, (E) phagocytes, (F) basal/goblet cells, (G) muscle cells, (H) neurons, (I) parenchymal cells, (J) protonephridia cells, and (K) secretory cells. Colour intensity of TFs in networks indicates relative outdegree (fraction of emitting connections). Outlined dots in stripcharts indicate the position of the labelled genes. L: Scatter plot showing influence scores of different TFs for the transitions from neoblast to phagocytes and from neoblasts to parenchyma, showing some TFs have high influence scores in more than one cell type. M: Heatmap of top co-influential TFs for different modules of co-influential TFs. TFs in rows, fates in columns. Top tree: clustering of fates based on the co-influence profile of these TFs.

To investigate if these similarities go beyond the molecular signature of differentiated cell types, we used *ANANSE influence* to compare the cell fate networks from neoblasts to every differentiated cell type (Figure 4A). *ANANSE influence* uses differential gene expression data to compare the two networks and identify the so-called influential factors –TFs whose expression changes the most and show highest binding to differentially expressed genes between two networks. Comparing a differentiated cell type network against the neoblasts network can predict key TFs driving the differentiation of pluripotent stem cells to that differentiated cell type. Thus, we performed differential gene expression analysis on our RNA dataset comparing every broad cell type against neoblasts independently (Supplementary Figure 11A, Supplementary File 14). We used this data to generate the cell fate networks from neoblast to every major cell type, and retrieved dozens of influential TFs for each cell fate alongside their target genes, a summarised excerpt of which can be seen in Figures 4C-K (Supplementary Figure 11B-J, Supplementary Files 15, 16, 17). Our predicted influence networks recapitulate well-known TFs from the scientific literature, such as nkx2−2 in phagocytes ^73, 81^, or *pou2/3-*1 in protonephridia ^76^. Some of the predicted targets also align with expectations extracted from the literature, such as *gata456*, predicted target of *nkx2-2* in phagocytes, or *ca-VII*, predicted target of *pou2/3-1* in protonephridia. These observations lend support to our ANANSE analysis and suggest it can predict interactions between TFs and target genes.

We noticed several transcription factors appear as influential for more than one fate, such as *hnf4* in phagocytes, basal/goblet cells, and parenchymal cells (Figures 4E,F,I). Several of these were consistent with previous knowledge. For instance, *foxf-1* was most influential in parenchyma and muscle, as recently described ^72^ but not in phagocytes. Conversely *nkx2-2* was among the most influential TFs in phagocytes as expected ^73, 81^, but not in parenchyma, with *hnf4* among the top ranked by influence in both (Figure 4L). To further investigate co-influence we clustered these TFs in groups of co-influence, detecting sets of TFs that share a similar profile of influence over one or multiple fates (Figure 4M, Supplementary Figure 11K, Supplementary File 18, Supplementary Note 2; See Methods). For instance, module m05 contains TFs co-influential in muscle and parenchyma (Supplementary Figure 11L), including *foxf-1* ^72^. Our groups of co-influentiality include TFs influential in protonephridia/neurons (m06), muscle/ secretory cells (m08), and neurons/muscle (m04). A further group had high scores in parenchymal and gut cells (m07), which included *hnf4* among the top five. Together with our previous analyses, this suggests that *hnf4* is an important TF to regulate the differentiation of neoblasts into cell types other than gut phagocytes.

Overall, our results show that our scRNA-seq/scATAC-seq integration model is capable of elucidating the regulatory logic of planarian stem cell differentiation. Our results suggest that this similarity of networks underlies the differentiation process, achieved by a combinatory logic of broad, co-influential, and cell type-specific TFs.

### *hnf4* RNAi induces depigmentation, head loss and lethality

Our previous findings revealed that many transcription factors are expressed in multiple cell types, including some previously thought to be exclusive to a single type. For instance, *hnf4* was initially reported to be expressed in the gut and their progenitors, the gamma neoblasts ^73^, consistent with a role in phagocyte differentiation. Recent single-cell transcriptomic studies have also revealed a prominent expression in parenchymal cells ^44, 45^. Our results indicated that *hnf4-*related motifs are highly enriched in genes expressed in both phagocytes and parenchymal cell clusters, as well as in the OCRs of those cell types. Our ANANSE analysis predicted *hnf4* as a top influential factor for both gut cell types and parenchyma.

To further explore the role of *hnf4*, we performed *hnf4* RNAi experiments in two independent biological replicates. We injected 25 animals per replicate with *hnf4* dsRNA and 25 animals with *gfp* dsRNA, as negative control, over 3 consecutive days. We monitored them from days 9-15 post-injection and characterised the phenotype of the *hnf4* knockdown. The first observed effects were small depigmentation and necrotic patches. These lesions frequently appeared in the pre-pharyngeal region, but also affected other body parts. Gradually, the pre-pharyngeal region accumulates depigmentation and tissue damage, which can result in the head being cleaved from the body or disintegrating (Supplementary Figure 12). The resulting headless tails did not regenerate and eventually died. Nine days post-injection, all animals treated with control *gfp* dsRNA presented healthy phenotypes. Although most worms treated with *hnf4* dsRNA were healthy, 9 out of 50 (18%) already showed an altered phenotype at this timepoint. From them, 3 out of 50 (6%) had no head. Five healthy-phenotype worms were selected from each replicate and condition and further monitored from day 12 to 15. During this time, all control-treated worms remained healthy. On day 12, 60% of hnf4-treated animals showed head damage or headless phenotypes. By day 15, every animal treated with *hnf4* had at least some degree of head damage. Overall, the *hnf4* knockdown has a strong phenotype which mostly affects the pre-pharyngeal body and ultimately leads to head loss and death. We were able to characterise this phenotype in both biological replicates.

### *hnf4* RNAi results in aberrant phagocyte progenitors that fail to activate phagocyte differentiation genes

To validate our ANANSE observations, we performed SPLiT-seq on our *hnf4* RNAi experiment samples (Figure 5A). We reasoned that this approach can measure systematically all genes in computationally dissected broad type. Therefore, contrary to classic approaches such as *in situ* hybridisation and qPCR, the results can be compared with each of the ANANSE generated networks, interactions and influence scores.

**Figure 5.**
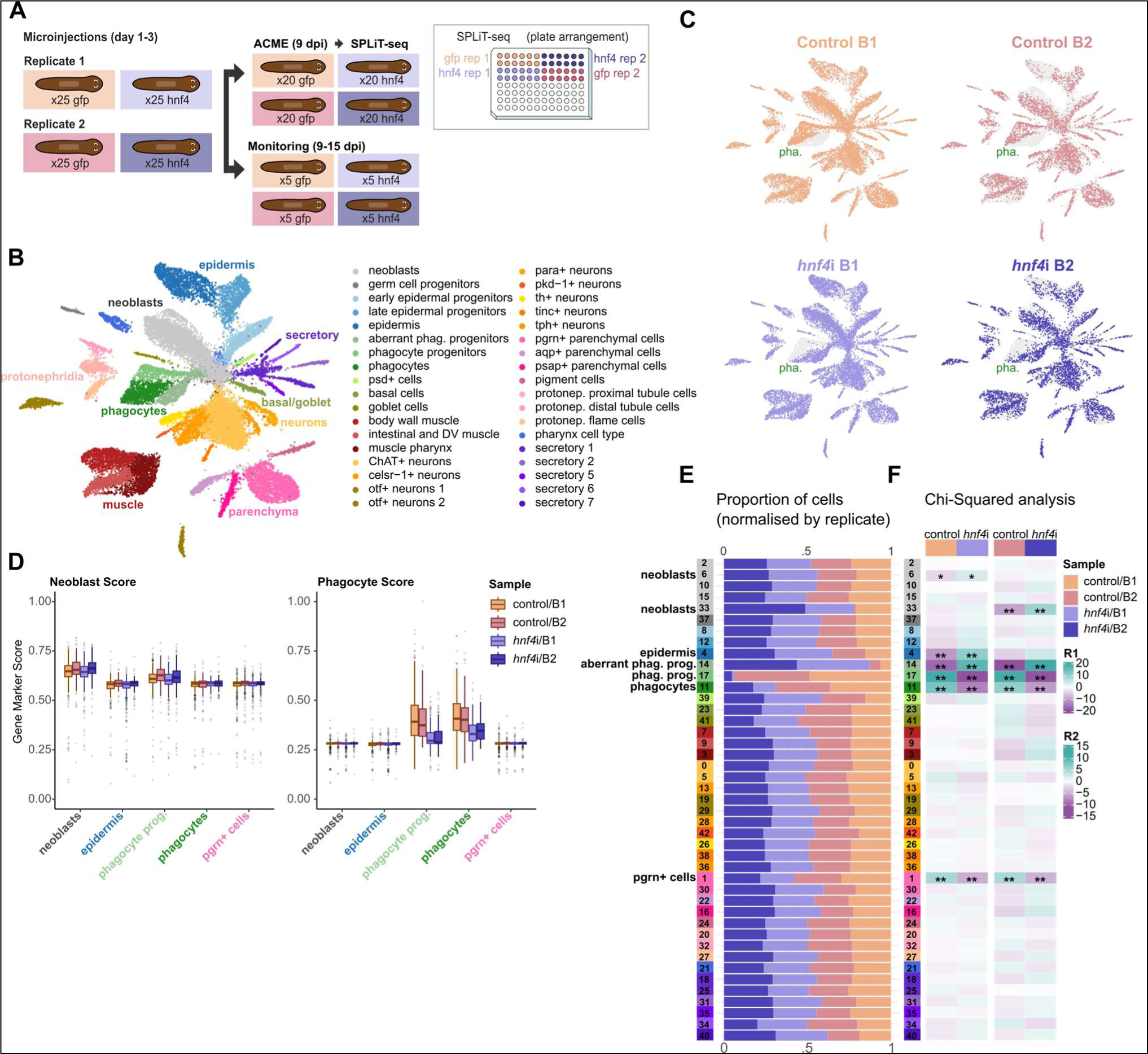
Knock-down of *hnf4* in planarian stem cells from a single cell analysis perspective. A: Overview of the experimental design of the knock-down. B: UMAP projection of 26,596 individual cell transcriptomes in the *hnf4*i dataset. C: Distribution of cells from the different conditions and replicates across the UMAP projection. D: Gene score of neoblasts (left) and phagocytes (right) markers in neoblasts, epidermis, phagocyte progenitors, phagocytes, and *pgrn*+ parenchymal cells of each condition and replicate. Center line, median; box limits, upper and lower quartiles; whiskers, 1.5x interquartile range; points, outliers. E: Fraction of cells from each condition and replicate on each Seurat cluster. Row colours indicate transferred cell identities. Name labels for cell clusters with significant differences in abundance (see F). F: Heatmap of residuals from Chi-squared post-hoc analysis of cell abundances in each Seurat cluster. Asterisks represent level of significance; * : p < .05, **: p < .01.

We analysed this independent dataset with our single-cell transcriptomic analysis pipeline, transferring labels using the scRNA-seq (Figure 1) dataset. This experiment yielded 41,016 single-cell transcriptomes, including all known major planarian cell types (Figure 5B, Supplementary Figure 13A, Supplementary File 19). We observed that phagocyte progenitor cluster 17 consisted almost entirely of *control*(*RNAi*) cells, while another cluster of phagocyte progenitors, cluster 14, was predominantly composed of *hnf4*(*RNAi*) cells (Figure 5C). We termed cluster 14 cells as “aberrant phagocyte progenitors” as they resemble phagocyte progenitors but are mostly observed in the knockdown condition. In the transfer labels analysis, cluster 14 was the only cluster that received labels from 2 distinct broad types, neoblasts and phagocyte progenitors (Supplementary Figure 13B).

We then investigated the nature of the aberrant phagocyte progenitor cluster. One possibility is that these cells fail to activate genes related to phagocyte biology. Alternatively, they might fail to deactivate genes related to stem cell biology. To differentiate between these scenarios, we scored the aggregated expression of neoblast and phagocyte markers in neoblasts, phagocyte progenitors and phagocytes from each experimental group, including epidermis and *progranulin*+ parenchymal cells as controls (Supplementary File 20). Remarkably, we found no significant differences between control and RNAi samples for the neoblast marker score, but the phagocyte score was significantly reduced in phagocytes and phagocyte progenitors (Figure 5E). This analysis indicated that *hnf4*(*RNAi*) phagocyte progenitors fail to activate phagocyte genes.

### *hnf4* knockdown leads to a decreased proportion of gut phagocytes and parenchymal cells

We visualised cell proportions in each cluster and replicate (Figure 5F) and statistically analysed these proportions in each biological replicate to identify significant and reproducible changes (Figure 5G, Chi-squared test) (Supplementary File 19). These analyses showed that phagocyte progenitors are significantly and reproducibly reduced in *hnf4* RNAi samples of both replicates. Similarly, the number of fully differentiated phagocytes significantly decreased in *hnf4* RNAi samples of both replicates. Importantly, the only other cluster significantly reduced in *hnf4* RNAi samples of both replicates was cluster 1, containing *progranulin*+ parenchymal cells. All other clusters were not significantly affected except for clusters 4, 6, and 33. Cluster 4, the major epidermal cell cluster, was significantly increased in *hnf4* RNAi in one biological replicate, likely due to the proportion-based test and the decrease in other clusters. Clusters 6 and 33, containing neoblasts, were significantly increased in the knockdown condition of one biological replicate, and likely contain neoblasts early in the differentiation process towards aberrant phagocyte progenitors. Taken together, these analyses showed that *hnf4* RNAi led to decreased proportions of both gut phagocytes and parenchymal cells.

### *hnf4* regulates independent but overlapping transcriptional gene expression programmes in phagocytes and parenchymal cells

Our multiplexed approach includes biological replicates, which are essential in bulk RNA-seq for identifying significantly regulated genes. Several researchers have highlighted that replicates are equally important for single-cell pseudobulk methods ^47^. However, the high cost of such methods presents a challenge. Single-cell combinatorial barcoding techniques, like SPLiT-seq, offer a solution by allowing the multiplexing of multiple samples within a single experiment, thereby reducing costs and batch effects.

To analyse gene expression changes in each cell type, we aggregated gene expression counts based on cluster identity and sample origin. This allowed us to create pseudo-bulk count tables for each cell type and broad group separately, enabling computational dissection of each cell type within each sample. We then employed DESeq2 ^82^ for differential gene expression (DGE) analysis, incorporating biological replicates for each cell type (Figure 6A, Supplementary File 21). To evaluate the response rate to the knockdown in each cell type, we analysed the relationship between cluster size and the number of Differentially Expressed Genes (DEGs) detected in each broad cell group (Figure 6B) and individual cell cluster (Supplementary Figure 14A,B). This analysis showed that most DEGs are in phagocytes and parenchymal cells.

**Figure 6.**
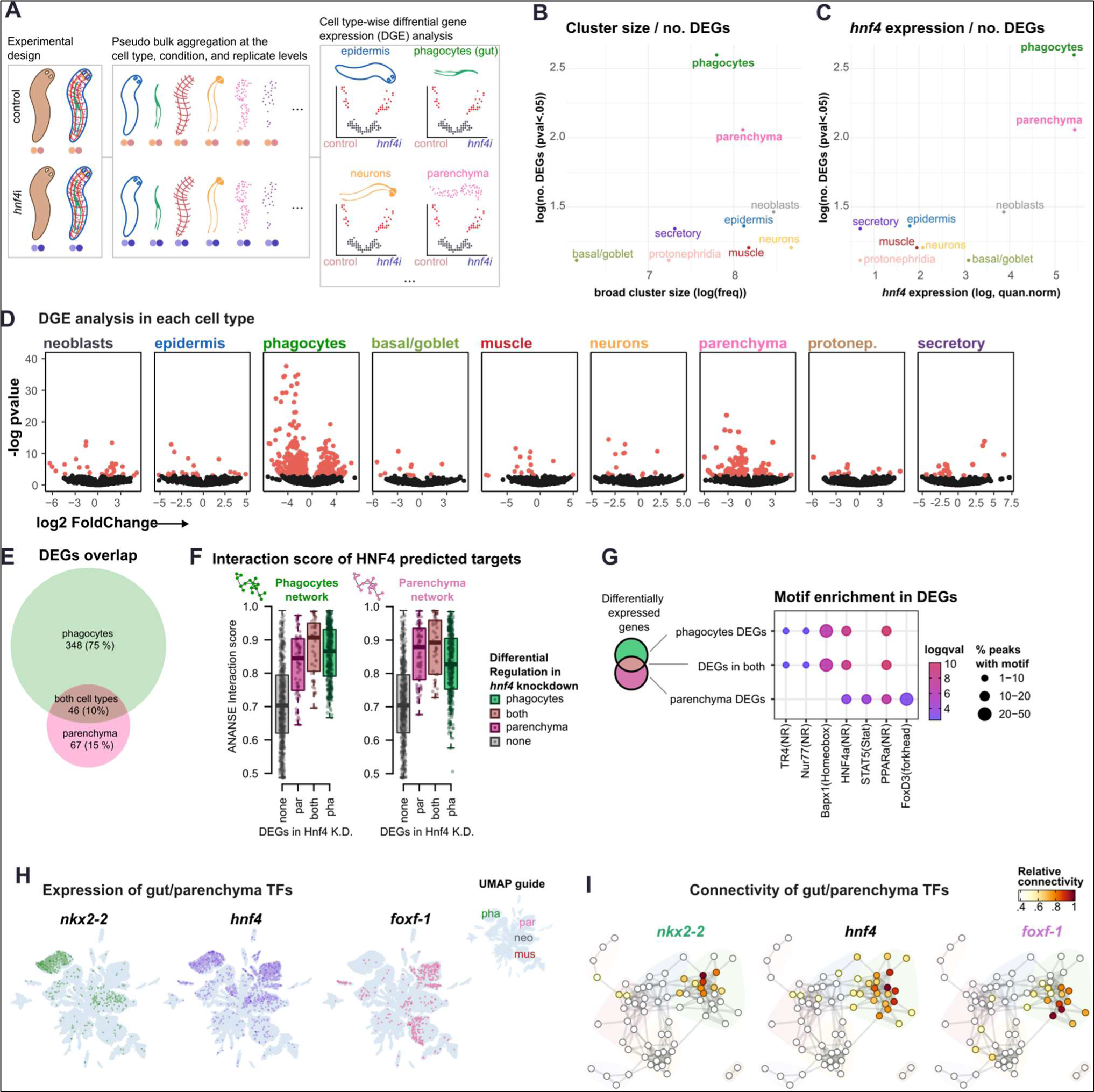
Differential gene expression of *hnf4*i knock-down in different cell types. A: Schematic of computational dissection of the scRNA-seq dataset to perform differential gene expression (DGE) analysis between control and *hnf4*i cells. B: Scatter plot showing the number of differentially expressed genes on each broad cell type (DEGs) in relation to the broad cell type cluster size (number of cells). C: Scatter plot showing the number of DEGs on each broad cell type in relation to the level of expression of *hnf4* on each broad cell type. D: Volcano plots of DEGs on each cell type. X axis: log fold change of expression in *hnf4*i cells relative to control cells. Y axis: negative log p-value. E: Venn diagram showing the overlap of DEGs in phagocyte and parenchymal cells. F: Box plots showing the predicted ANANSE interaction score between *hnf4* and target genes. Center line, median; box limits, upper and lower quartiles; whiskers, 1.5x interquartile range; points, data points. G: Motif enrichment in DEGs based on the overlap between phagocytes and parenchymal cells. H: Feature plots of the expression of *nkx2-2*, *hnf4*, and *foxf-1* in our whole scRNA dataset. I: Connectivity of *nkx2-2*, *hnf4*, and *foxf-1* to the different WGCNA co-expression modules (see Figure 2).

We visualised DEGs in each broad cell type as volcano plots. These confirmed that most DEGs occurred in phagocytes and parenchymal cells (Figure 6C,D). The most highly significant DEGs in both corresponded to downregulated genes, consistent with a role of *hnf4* as a transcriptional activator. Many genes were significantly up- or down-regulated in both phagocytes and parenchymal cells (46, 10%), but many others were differentially regulated only in phagocytes (348, 75%) or parenchymal cells (67, 15%) (Figure 6E). These gene sets are enriched in broadly different gene ontology terms (Supplementary Figure 14C-F), suggesting they are functionally independent.

We then aimed to determine if DEGs detected in our *in vivo* knockdown data overlapped with *in silico* predicted *hnf4* target genes from our ANANSE analysis. Importantly, these two analyses are entirely independent. To analyse this overlap, we examined the interaction score of DEGs in phagocytes, parenchymal cells, and those regulated in both, comparing these to the rest of the genes (Figure 6F). On average, *in vivo* DEGs of all three groups had increased *in silico* predicted ANANSE interaction scores in both the phagocyte and parenchymal cell ANANSE networks. Altogether, these analyses showed that *hnf4* regulates independent but overlapping gene expression programs in both gut phagocytes and parenchymal cells. These results validate our *in silico* ANANSE analysis and lend support to our ANANSE predictions for all other TFs and interactions.

Finally, we questioned what other factors could be responsible for the differences between phagocytes and parenchymal cells. We analysed motifs enriched in all three DEG groups (Figure 6G). Consistently, we found nuclear receptor HNF4 motifs in all three groups, but we also found motifs specific to phagocyte DEGs and parenchymal cell DEGs. Interestingly, a homeobox factor motif was highly enriched in phagocytes, and a Fox factor was highly enriched in parenchymal cells (Figure 6G). We hypothesised that these could correspond to *nkx2-2* and *foxF-1* (Figure 6H), which have been shown to be important for phagocyte and parenchymal cell differentiation respectively ^72, 73, 81^. We examined the connectivity of these factors with WGCNA modules (Figure 6I). Indeed, *nkx2-2* had higher connectivity to phagocyte-only modules, and *foxF-1* to parenchymal cell modules. Altogether, these analyses corroborate that DEGs detected *in vivo* have *hnf4*-related motifs as well as motifs of other factors expressed in each of the two cell types.

## Discussion

Gene regulation involves the combinatorial expression of transcription factors (TFs) and the accessibility of chromatin regions (OCRs) acting as cis-regulatory elements (CREs), forming a ‘regulatory logic’ that creates Gene Regulatory Networks (GRNs). GRNs are typically represented as graphs with genes as nodes and CRE-mediated interactions as edges. Traditional GRN inference through perturbational experiments reveals direct gene interactions but may miss indirect ones and is limited by experimental constraints. For instance, in planarians they require laborious, gene-specific studies often limited to one cell type or tissue, highlighting the need for comprehensive techniques to assess regulatory networks in regenerative animals. Single-cell technologies predict TF/target gene interactions but they often lack experimental validation and their scalability to the whole organism level is unclear.

In this study we profiled the transcriptional landscape and chromatin dynamics of planarian cell types using single cell technologies. Using correlational and pseudobulk computational dissection approaches, we detected modules of combinatorial gene expression and chromatin accessibility, uncovering potential TF/target gene interactions. We observed similar trends of combinatorial expression, chromatin accessibility, and TF fate influence in two independent sources of data from different modalities (sc-RNA-Seq and scATAC-Seq).

Our data suggest multiple TFs are influential in multiple cell types, and OCRs follow similar combinatorial logic. The main example are phagocytes and parenchymal cell types, with numerous genes co-expressed as observed in our correlational WGCNA, but also a large number of shared OCRs specific to differentiated cell types compared to neoblasts, as well as a number of TFs influential in both. This may be caused by functional similarities as both have phagocytic activity ^72^, or a lineage relationship. We observed similar trends in neurons and secretory cells, which might be related to their shared secretory nature. Of note, goblet cells are also thought to be secretory, but share fewer connections with neurons and/or secretory cells than with gut phagocytes and parenchyma, suggesting that this relationship is lineage-driven. Future studies will address whether these associations reflect functional or lineage relationships, or a mixture.

Our analyses identified *hnf4* as a regulator of both phagocyte and parenchymal cell identity, which we validated with a knockdown experiment. We detected previously undescribed changes both in cell abundance and gene expression in both types. Interestingly, gene expression and motif enrichment analysis suggest TFs *nkx2* and *foxf1* are differentially specific to each of these types, consistent with previous studies ^72, 73, 81^. Therefore, this supports the notion that the discrimination between phagocyte identity and parenchymal cell identity depends on the expression of *hnf4*+*nkk2* and *hnf4*+*foxf1* respectively. We envision that future studies will decode similar relationships in GRNs with functional links akin to logic gates ^83^.

Of note, the accessible regions of neoblast specific genes are open in most other cell types, resembling constitutive promoters. This confirms previous results obtained by tissue fractionation ^40^, revealing that planarian neoblasts follow chromatin regulation rules distinct from those of their differentiated counterparts. Many regenerating animals have stem cells that are similar to planarian neoblasts, at least at the gene expression level ^84-86^. Recently, we described that in the annelid *Pristina leidyi* most cell types can be connected by computational differentiation trajectories to a population of *piwi*+ cells ^62^. Interestingly, these express in high levels most chromatin regulators, something that was also described in planarian neoblasts ^87^. However, their chromatin properties remain largely unexplored, and this is the case for a vast majority of regenerating animals. Combining single cell transcriptomic and accessibility techniques will allow us to elucidate if this is a universal feature of animal stem cells.

Our results were limited by the resolution of the scATAC-seq, which resolved the major broad cell types, including epidermis, muscle, neurons, phagocytes, parenchymal cells, protonephridia, secretory cells and basal/goblet cells. These can be understood as the major “fates” in other GRNs, and discriminating major fates is also challenging using classic techniques such as *in situ* hybridisation. However, scRNA-seq has a higher clustering resolution, discriminating individual cell types. Future studies will also see increased resolution of scATAC-seq.

Overall, our data demonstrates that GRN computational inference can co-exist with classic functional approaches, as the former is able to formulate data-driven hypotheses of TF influence in cell fates that can be functionally validated by perturbation assays coupled with single cell analysis. We already explored this possibility by studying the relationship between the ANANSE predictive score of *hnf4*-gene interactions and the differential gene expression on hnf4i cells and found it to be positively correlated. Future studies will incorporate a larger number of knockdowns, exploiting the scalability of combinatorial barcoding single cell transcriptomic methods. Our study pioneers this avenue, which will lead to decoding the GRNs underlying regeneration and development broadly in the animal kingdom.

## Materials and Methods

### Experimental batches

This study comprises data from 6 independent experiments: 1 scATAC-seq and 5 scRNA-seq (batches 1, 8, 11, 14 and 23). In addition, each scRNA-seq batch comprises multiple libraries (batch 1: libraries 1-3, batch 8: libraries 8.3 and 8.4, batch 11: libraries 11.3 and 11.4, batch 14: libraries 14.3 and 14.4, and batch 23: libraries 23.1 and 23.2) (Figure 1A).

### Animal culture and collection

All libraries used in this study were generated from asexual *Schmidtea mediterranea* worms derived from the clonal line *Berlin-1* ^88^. The animals were kept at 18-20°C in 1x Montjuic water (1.6 mM NaCl, 1.0 mM CaCl2, 1.0 mM MgSO4, 0.1 mM MgCl, 0.1 mM KCl, and 1.2 mM NaHCO3, dissolved in deionised water) at pH 7.0. Planarians were fed cow liver once or twice per week and starved 7 days minimum before any experimental procedure. Animal collection consisted of a random selection of mixed-size healthy individuals (1-10 mm), except for batches 11 and 23. For batch 11, we selected 6-8 mm animals, as is the standard size for dsRNA injection. Animal collection for batch 23 was performed as described in Emili et al. 2023 ^57^.

### Knock-down by RNAi

Batch 11 was generated using knockdown samples treated with *gfp* (control) or *hnf4* dsRNA. These samples were obtained according to the following protocol:

**Primary PCR:** To amplify *hnf4,* we used cDNA from wild type *S. mediterranea* worms. To amplify *gfp*, we used a DNA miniprep (13 ng/uL) of enhanced GFP in a pAGW vector provided by the Drosophila Genomics Resource Center. Primary PCR was performed using 2 µL of cDNA/DNA miniprep, 2 µL of 10x Standard Taq Reaction Buffer (NEB), 0.4 µL of dNTPs (2.5 µM), 0.2 µL of Hot Start Taq DNA Polymerase (NEB), 4 µL of Primer Forward (2.5 μM), 4 µL of Primer Reverse (2.5 μM) and 7.4 µL of water. The primer sequences were as follows: ggccgcggCGCTGAAATAGCCAGTCACA (hnf4-F), gccccggccGCCGCTTCAGGTGATATGTT (hnf4-R), ggccgcggGTCTATATCATGGCCGACAAG (gfp-F) and gccccggccACTGGGTGCTCAGGTAGTGGT (gfp-R). *Hnf4* primers were designed from the GenBank sequence JF802199.1. Both primer pairs included linkers for Universal T7 primers: ggccgcgg (linker-F) and gccccggcc (linker-R). The thermocycler programme used was: 94°C (30 s); 35 cycles at 94°C (20 s), 55 °C (20 s) and 68°C (30 s); and 68°C (5 min). We assessed PCR products in a 1% agarose gel, cut the bands under UV light, and froze them in 50 µL of nuclease-free water at -20°C.

**Secondary PCR:** Samples were thawed and centrifuged at maximum speed for 1 min to extract cDNA from the gel bands. The supernatants were collected and used as cDNA input for the secondary PCR. We prepared 100 µL reactions using 3 µL of cDNA, 2 µL of dNTPs (2.5 µM), 10 µL of 10x Standard Taq Reaction Buffer, 1 µL of Hot Start Taq DNA Polymerase, 82 µL of water, 1 µL of Universal T7-F5’ primer (25 µM, gagaattctaatacgactcactatagggccgcgg) and 1 µL of Universal T7-R3’ primer (25 µM, agggatcctaatacgactcactataggccccggc). Samples ran in a thermocycler as follows: 94 °C (30 s); 5 cycles at 94 °C (20 s), 50 °C (20 s) and 68°C (30 s); then 35 cycles at 94°C (20 s), 65°C (20 s) and 68°C (30 s); and 68°C (5 min). The size of the bands was assessed by running 10 µL/sample in a 1% agarose gel. The remaining volume was purified by 0.75x (*hnf4*) or 1.6x (*gfp*) SPRI size selection (KAPA Pure Beads, Roche) according to the manufacturer’s protocol. Purified samples were eluted in 20 µL of nuclease-free water.

**dsRNA synthesis:** For each sample, we mixed 1 µg of purified cDNA, 12.5 µL of 2x Express Buffer (T7 RiboMAX, Promega), 2.5 µL of Express Mix (T7 RiboMAX, Promega), and up to 25 µL of nuclease-free water, and incubated for 4 hours at 37°C. Then, we added 2.5 µL of DNase (1 U/µL, T7 RiboMAX, Promega) and incubated for another 30 min at 37°C. After incubation, reactions were stopped with 375 µL of Stop Solution (1M NH4OAc, 10 mM EDTA, 0.2% SDS). The resulting dsRNA was purified using phenol:chloroform. We added 1 µL of GlycoBlue and 400 µL of acid phenol:chloroform (pH 4.5, Thermo Fisher) per reaction and vortexed thoroughly. We centrifuged for 5 min and collected the aqueous top phase into a new tube. We added 400 µL of chloroform, centrifuged for 5 min, and collected the top phase again. To precipitate pellets, we added 1 mL of cold ethanol, vortexed and centrifuged for 15 min. Pellets were washed in 1 mL of 70% ethanol and centrifuged for 10 min. We discarded supernatants and let the pellets dry for 5 min at 37°C. Pellets were resuspended in 10-20 µL of nuclease-free water. All centrifugations were performed at 4°C and maximum speed. As a quality check, we ran 0.5 µL of purified dsRNA in a 1% agarose gel. Finally, we measured the concentration in a Nanodrop.

**Injections and harvest:** For injections, we used worms that were 6-8 mm in length. Each animal was injected with 0.1 μg of dsRNA for 3 consecutive days (0.3 μg in total). We generated two replicates per condition (*gfp* and *hnf4*) that were both biological and technical, as they were processed by different researchers. We injected 25 animals per replicate with *hnf4* dsRNA and 25 animals per replicate with *gfp* dsRNA, using a Nanoject II Auto-Nanoliter Injector (Drummond Scientific Company). At 9 days post-injection, counted from the last day of injections, we harvested 20 animals per replicate and condition and dissociated them using ACME (as described below). The remaining 5 animals were kept uncut and monitored from day 9 to 15 post-injection.

### ACME dissociation

Tissue dissociation and fixation were performed using ACME as described in ^55^ with the following modifications: Incubation time was 35 min for batch 11, and 45 min for batches 14 and 23. After incubation, all samples were pipetted up and down as in the original protocol. Then, batches 11, 14 and 23 were kept on ACME solution (on ice) for 3 consecutive rounds of filtration to help remove cell aggregates and undissociated tissue fragments. Samples were first filtered through 50 μL and 30-40 μL strainers (Celltrics). Then, they were centrifuged at 1000 g for 5 min (4°C) to reduce the volume of the solution to 1-2 mL; by discarding part of the supernatant and resuspending the pellet in the remaining volume. Samples were then filtered into 15 mL tubes using 1 mL filter tips (Flowmi). To remove ACME solution and wash cells, we added 7-8 mL of buffer (1x PBS 1% BSA) to the filtered samples and centrifuged at 1000 g for 5 min (4°C). The supernatant was discarded, samples were resuspended in 900 μL of buffer and filtered one last time, using a 40 μm strainer, into 1.5 mL tubes. We added 100 μL of DMSO per sample and froze them at -80°C.

### SPLiT-Seq

Batch 1 was entirely processed using the SPLiT-seq protocol described in García-Castro *et al*. ^55^. Batches 8, 11, 14 and 23 were processed using the modifications introduced in Leite *et al*. ^89^ with the following variations:

**Sample preparation:** Frozen dissociated cells (unsorted) were thawed, centrifuged twice at 1000g for 5 min (4 °C), resuspending in ∼450 μL of buffer (1× PBS 1% BSA), and filtered through a 50 μL strainer (CellTrics). For each sample, we stained a 1:3 aliquot (50 μL sample + 100 μL buffer) for 20 min at RT (in the dark) using 1.5 μL of DRAQ5 (0.5 mM stock) and 0.6 μL of Concanavalin-A conjugated with AlexaFluor 488 (1 mg/mL stock, Invitrogen). The remaining cells were kept at 4°C. To estimate cell concentration (singlets/μL), three measurements of 10 μL were taken for each aliquot by flow cytometry. To count singlet events we used a FSC-H vs FSC-A gate to select singlets, a Concanavalin-A positive gate to select cells with cytoplasm and a DRAQ5 positive gate to select cells vs cellular debris. The remaining unstained cells were diluted according to this singlet cell count, in 0.5x PBS, to a final working concentration of ∼625k cells/mL (5000 singlets per well).

**Plate loading:** Batch 11 comprises four different samples: *gfp* RNAi replicate 1, *hnf4* RNAi replicate 1, *gfp* RNAi replicate 2 and *hnf4* RNAi replicate 2. Each of these samples were loaded separately into specific wells (12 wells per sample) during round 1 of barcoding, so they could be deconvoluted during the bioinformatic analysis.

**FACS:** For each SPLiT-seq experiment, we sorted two separated libraries obtaining the following numbers: 19k cells (library 8.3), 24k cells (library 8.4), 17k cells (library 11.3), 18.5k cells (library 11.4), 6.1k cells (library 14.3), 6.6k cells (library 14.4), 19k cells (library 23.1) and 19k cells (library 23.2).

**PCR amplification:** Samples were amplified for 10-11 qPCR cycles.

### scATAC-Seq library preparation

Nuclei suspensions for scATAC-seq were obtained from trypsin dissociated cells. Essentially, we chopped planarians into small pieces on ice, and incubated the pieces in 2-4 ml of PBS containing 1% BSA and 1% Trypsin for 25-30 minutes at room temperature, gently pipetting up and down until fragments were completely dissociated. Cells were then pelleted at 1000xg for 5 minutes at 4°C, and resuspended in n 4-5 ml of PBS containing 1% BSA. We filtered the cells through a 40 µm cell strainer (Becton-Dickinson) and through a 20 µm nylon net filter (Millipore). We pelleted cells at 1000xg for 5 minutes at 4°C, and added 100 µl of chilled lysis buffer (10 mM Tris-HCl (pH 7.4), 10 mM NaCl, 3 mM MgCl2, 0.1% Tween-20, 0.1% Nonidet P40 Substitute, 0.01% Digitonin, 1% BSA in Nuclease-free Water) to the cell pellet and mixed it by pipetting up and down 10 times. We incubated the mixture on ice for 3-5 minutes, and added 1 ml of chilled Wash Buffer (10mM Tris-HCl pH 7.4, 10mM NaCl, 3 mM MgCl2, 1% BSA, 0.1% Tween-20 in Nuclease-free Water) to the lysed cells and mixed it by pipetting up and down 5 times. We centrifuged the mixture at 500 rcf for 5 minutes at 4°C and removed the supernatant carefully without disturbing the nuclei pellet. Nuclei were then resuspended in Diluted Nuclei Buffer (10X Genomics) and counted before injection in the 10X Genomics Chromium, following the manufacturer’s protocol. The libraries were then amplified using Nextera library prep and sequenced in a NextSeq Illumina sequencer to obtain 75PE reads.

### Bulk ATAC-Seq library preparation

To generate nuclei suspension for bulk ATAC-seq, we flash frozen planarians for 1 min in liquid nitrogen and resuspended in cold lysis buffer (10 mM Tris-HCl (pH 7.4), 10 mM NaCl, 3 mM MgCl2, 0.1% Nonidet P40 Substitute, in Nuclease-free Water). While in lysis buffer on ice, planarians were dissociated by smashing them against a 100um cell strainer with the aid of a syringe plunger. The resulting nuclei suspension was centrifuged (500 rcf for 5 min at 4°C) and resuspended in PBS 0.04% BSA. A fraction (50ul) of the nuclei suspension was labelled with DRAQ5 (1.5ul of 0.5mM stock), counted with a cytometer (as in the Sample preparation section of the SPLiT-Seq above), and the volume to have 20k nuclei was calculated. The resulting volume was centrifuged and resuspended in 50ul of tagmentation buffer (5 mM MgCl2, 10 mM Tris HCl, 9.4% Dimethylformamide in Nuclease-free water), 2.5 ul of custom loaded Tn5 (5′ TCGTCGGCAGCGTCAGATGTGTATAAGAGACAG and 5′ GTCTCGTGGGCTCGGAGATGTGTATAAGAGACAG) was added to the nuclei suspension and incubated at 37°C for 30 minutes. After tagmentation, we resuspended nuclei in stop reaction mix (20 mM EDTA, 0.5 mM Spermidine, in nuclease-free water) and incubated at 37°C for 15 minutes. We proceeded with the library preparation by purifying the tagmentation product using the Monarch PCR DNA Cleanup Kit (New England Biolabs) and following manufacturer’s instructions. The tagmentation product was eluted in 10ul of Nuclease-free Water, and the DNA concentration was measured using the NanoDrop. To evaluate the optimal number of amplification cycles, we first ran a qPCR using 1ul of tagmented DNA, mixed with 5ul of 2× Kapa HiFi HotStart ReadyMix (Roche), 0.5μl of PCR_PF (25 μM, 5’-AATGATACGGCGACCACCGAGATCTACACAATCCGCGTCGTCGGCAGCGTCAGATGTGTAT), 0.5μl of PCR_PR (25 μM, 5’-CAAGCAGAAGACGGCATACGAGATTCATTAGGGTCTCGTGGGCTCGGAGATGTG), 3 μL of nuclease-free water and 0.5 ul of EvaGreen® Dye (20X in Water, Biotium). We then amplified 1ul of tagmented DNA using the same mix of the qPCR without EvaGreen, with the following conditions: 30 s at 98 °C; 10s at 98 °C, 30s at 65 °C, and 60s at 72 °C repeated for 11 cycles; final elongation for 5 minutes at 72 °C. Finally, we purified the PCR products, quantified it with NanoDrop, and check the tagmentation profile using an Agilent 2100 bioanalyzer. The library was sequenced in a NovaSeq X Plus PE150 Illumina sequencer.

### Gene functional annotation

#### Prediction of protein sequences

We first extracted the coding sequences from the latest version of the *S. mediterranea* ^33^ using AGAT ‘agat_convert_sp_gff2gtf.pl’ (https://www.doi.org/10.5281/zenodo.3552717) and standard parameters. The resulting set of coding sequences was then transformed to protein sequence using TransDecoder v5.5.0 (https://github.com/TransDecoder/TransDecoder). first, we ran ‘TransDecoder.LongOrfs’ with standard parameters; second, we ran hmmscan vs Pfam database and BLAST vs Swissprot database, with parameters:‘-max_target_seqs 1 -evalue 1e-5’ and default parameters respectively, to gather supporting evidence for coding transcripts; third, we ran ‘TransDecoder.Predict’ with parameters ‘--retain_pfam_hits pfam.domtblout --retain_blastp_hits blastp.outfmt6 --single_best_only‘. We manually curated and removed gene and transcript features with few or no hits against any known protein in our annotation databases (see below) that were overlapping with other features that did have hits against such databases.

#### Querying against previous annotations

The resulting set of predicted protein sequences (hereafter referred to as proteome) was queried against three previous genome annotations of *S. mediterranea* –Dresden v4, Dresden v6 ^90, 91^, and the extended annotation used in Garcia Castro et al. ^55^, to retrieve reciprocal best hits. Briefly, we ran BLASTp against each set of predicted protein sequences of each genome annotation version, using standard parameters. Secondly, genes without a clear one-to-one reciprocal match were queried more leniently against the previous versions; we retrieved all hits with e-value <.001.

#### eggNOG functional annotation

The resulting proteome of *S. mediterranea* was queried using EggNOG mapper41 with the parameters: ‘-m diamond --sensmode sensitive --target_orthologs all --go_evidence non-electronic’ against the EggNOG metazoa database. From the EggNOG output, GO term, functional category COG, and gene name association files, were generated using custom bash code.

#### Transcription Factor annotation

The resulting proteome of *S. mediterranea* was queried for evidence of Transcription Factor (TF) homology using (i) InterProScan ^92^ against the Pfam ^93^, PANTHER ^94^, and (ii) SUPERFAMILY ^95, 96^ domain databases with standard parameters, (iii) using BLAST reciprocal best hits ^97^ against swissprot transcription factors ^98^, and (iv) using OrthoFinder ^99^ with standard parameters against a set of model organisms (Human, Zebrafish, Mouse, *Drosophila*) with well annotated transcription factor databases (following AnimalTFDB v4.0 ^100^). For the latter, a given *S. mediterranea* gene was counted as TF if at least another TF gene from any of the species belonged to the same orthogroup as the *S. mediterrane*a gene. The different sources of evidence were pooled together and we kept those *S. mediterranea* genes with at least two independent sources of TF evidence. In addition to this, we also added to our list all the *S. mediterranea* genes with a match of high query coverage and e-value <0.001 against any of the TFs reported in Neiro *et al*. ^39^. The resulting list of 665 genes was manually curated to assign a TF class to each gene based on their sources of evidence.

#### Transcription Factor motif annotation

The resulting proteome of *S. mediterranea* was queried for TF motif annotation using gimmemotifs motif2factors, using the proteomes and the JASPAR 2020 TF annotation of *H. sapiens* and *M. musculus* as reference, as well as the proteomes of several protostome metazoan species (Supplementary File 13, Supplementary Figure 10A) to provide additional phylogenetic signal for the automated transfer. Secondly, we ran the JASPAR similarity prediction tool ^101^ on those TFs retrieved with our sequence homology annotation that did not get any transferred motif, using the JASPAR 2024 motif database which overlaps with the JASPAR 2020 database. We manually curated this motif annotation by adding any TF with an associated motif by Neiro *et al*. ^39^. The resulting list of 401 (out of 665) TFs was subsequently used for running ANANSE.

#### Definition of promoters

We defined promoters as gene regions ranging 200bp upstream the transcription start site (TSS) of genes and 200bp of the TSS using ‘bedtools flank’ ^102^ with standard parameters.

### Single Cell Transcriptomic Analysis

#### Gene annotation pre-processing

We parsed the genome annotation GFF3 file of *S. mediterranea* and converted it to GTF format using AGAT ‘agat_convert_sp_gff2gtf.pl’ (https://www.doi.org/10.5281/zenodo.3552717), after which we used a custom python script to add the gene_id, transcript_id, gene_name and transcript_name fields. This was done in order to comply with the requirements of dropseqtools and the SPLiT-Seq pipeline workflow (https://github.com/RebekkaWegmann/splitseq_toolbox) which envelops algorithms from Drop-seq_tools-2.3.0 (https://github.com/broadinstitute/Drop-seq) (hereafter dropseqtools; see below).

#### SPLiT-seq read processing

Single cell RNA-seq libraries were pre-processed as previously described ^55, 57, 62^. A total of 170173041 reads were sequenced. These were assayed for QC purposes using FastQC (https://www.bioinformatics.babraham.ac.uk/projects/fastqc). We concatenated shallow and deep sequencing using cat. We used CutAdapt v2.8 ^103^ to trim read 1 (transcripts) and read 2 (UMI and barcodes) sequences, using standard parameters. Reads were checked to be in phase using grep, and the resulting phased reads were paired using pairfq makepairs (https://github.com/sestaton/Pairfq). The resulting reads were transformed into sam format using picard FastqToSam and mapped against the *S. mediterranea* genome (with a tailored annotation for dropseqtools; see above) using the SPLiT-Seq pipeline workflow wrapper that uses Picard (http://broadinstitute.github.io/picard/), STAR ^104^, and dropseqtools.

#### Generation of scRNA Matrices

For each library, the resulting outputs of running the SPLiT-Seq pipeline (tag_bam_with_gene_function) were used to generate a gene x cell sparse matrix keeping cells with a minimum of 100 genes detected. When required, such as libraries 1-3 (from an experimental design containing *D. japonica* cells) and 11.3 and 11.4 (with *hnf4*i cells), any cells not coming from unperturbed *S. mediterranea* organisms were discarded by re-running the command ‘DigitalExpression’ from dropseqtools using a set of white-listed barcodes corresponding to *gfp*i control cells.

#### Matrix concatenations

The resulting matrices were concatenated using Seurat v4.3.0 ^61^ in R v4.0.3 ^105^. Briefly, we created independent Seurat Objects for each separate library and performed Proportional Fitting-log1p-Proportional Fitting normalisation ^106^ on each separate matrix. We labelled cells from each library with their respective library ID (Supplementary File 1) and merged all the Seurat objects into a single concatenated sparse matrix.

#### Seurat Analysis

The concatenated sparse matrix was queried for highly variable genes using ‘SelectIntegrationFeatures()’ from the Seurat package. We scaled the data and performed PCA using the function ‘runPCA()’ with the following parameters: ‘npcs = 120’ . We integrated the data using the function ‘runHarmony()’ from Harmony ^107^ and the following parameters: ‘dims.use = 1:120, theta = 3, lambda = 3, nclust = 40, max.iter.harmony = 20’. We identified neighbours for the k-nn graph using the function ‘FindNeighbors()’ and the following parameters = ‘dims = 1:120, k.param = 35’. We identified clusters using the function ‘FindClusters()’ with the Louvain algorithm ^108^ and the following parameters: ‘resolution = 2, algorithm = 1, random.seed = 75’. We computed an UMAP projection using the function ‘RunUMAP()’ and the following parameters: ‘dims = 1:120, reduction = “harmony”, n.neighbors = 35, min.dist = 0.5, spread = 1, metric = “euclidean”, seed.use = 1’ . To find cell markers, we used the function ‘FindAllMarkers()’ with the following parameters: ‘only.pos = TRUE, return.thresh = 1, logfc.threshold = 0’, and subsequently sorted these markers based on average logFC to keep the top 30 markers per gene cluster.

#### Alignment to reference dataset

We first ported the AnnData object from our previous study on allometry of cell types Emili et al. ^57^ (hereafter the Sizes object) to Seurat using custom python and R scripts. After transferring and transforming into a Seurat object, we ran the function ‘FindTransferAnchors()’ from Seurat using our Seurat analysis object (see section above) as query and the Sizes object as the reference, and the following parameters: ‘dims = 120’. The predicted labels of each cell in the query dataset were added as a metadata column. We assigned the most frequent predicted label to each Seurat cluster we obtained, and manually curated ambiguous assignments checking diagnostic markers from the reference dataset. Label transferring was visualised using igraph ^109^.

#### Pseudobulk computational dissection and normalisation (without conditions or replicates)

We aggregated the counts of every gene in each cluster using a custom R function, which yielded a gene x cluster matrix of expression. In parallel to this, we created a gene x cluster matrix that quantifies how many cells from each cluster are expressing a gene, using a custom R function with the parameter ‘min_counts = 1’. To adjust for expression dynamics across clusters from very different sizes (in terms of number of cells), we calculated a “cell weight” matrix that leverages the expression matrix and the cell number matrix to calculate, for every gene in every cluster, a “score” of expression constraint using the formula: *w_ij_ = 1 - exp(- C_ij_) = 1 - exp(- a_ij_ / b_ij_)*; where *w_ij_* is the cell weight of gene i in cluster j, *a_ij_* is the fraction of cells from cluster j expressing gene i, and *b_ij_* is the fraction of cells NOT from cluster j expressing gene i. This weighs down the expression of genes with counts scattered among a low fraction of cells in large clusters as opposed to smaller clusters that comparatively capture less reads but more frequently inside that cluster. We ran this code inside a custom R function with the parameters ‘min_counts = 30’ for genes with at least 30 counts through the dataset, and the parameter ‘min_cells = 3’ to retrieve expression information for genes expressed in at least three cells of a given cluster of interest’.

We normalised the matrix of expression by “library size” using the ‘DESeqDataSetFromMatrix()’ function with parameter ‘design = ∼ condition’ and the ‘counts()’ function with parameter ‘normalised = TRUÈ from the R package DESeq2 ^82^. This normalised expression matrix was weighed using the cell weights described above. The resulting cell-weighted normalised expression matrix was used for downstream analyses.

#### Co-occurrency analysis

Co-occurrency analysis was performed as previously described ^110, 111^. Briefly, the pseudobulk normalised matrix was subjected to Pearson correlation at the cell type level using bootstrapping and subsampling, using the function ‘treeFromEnsembleClustering()’ with the following parameters: ‘h = c(0.75,0.9), p = 0.05, n = 1000, bootstrap = FALSE, clustering_algorithm = “hclust”, clustering_method = “average”, cor_method = “pearson”’, and using all the genes in the pseudobulk matrix as ‘vargenes’.

#### WGCNA analysis

We subsetted our normalised expression matrix to keep genes with CV > 1.25 and scaled it across rows, and subjected this to the WGCNA algorithm ^68^. We picked soft power beta = 8 as it resulted in the highest fit to a scale-free topology model ^67^. The resulting adjacency matrix was weighted using topology overlap and the resulting TOM matrix was clustered using the function ‘hclust()’ with the following parameters: ‘method = “average”’. The resulting gene tree was cut into different modules of co-expression using the ‘cutreeDynamic()’ function from the WGCNA package ^68^ with the following parameters: ‘deepSplit = 3, pamRespectsDendro = FALSE, minClusterSize = 50’. We reclassified the resulting modules in specific or mixed modules based, for each module, on the number of outliers (value > (1.5 x standard deviation) + mean) on the distribution of upper quartiles of expression in cell clusters of the genes in that module (“s” if only one cluster clearly showcased a higher expression of the genes of that module, “m” if two or more clusters showcased higher expression of the genes of that module). In addition to this, we sorted these modules based on the identity of their clusters of peak expression. To create profiles of the relative amount of gene expression of each module, we calculated the average expression profile per module, and did an internal normalisation by dividing every count on each cluster by the sum of counts on all clusters. The resulting frequencies of expression in each cluster represent the average expression signal in each cluster, and these were visualised as stacked bar plots using the ComplexHeatmap R package ^112^. We subsetted each module to randomly retrieve thirty genes and visualised their expression profiles using the ComplexHeatmap R package ^112^.

#### Transcription factor expression analysis

We subsetted the pseudobulk normalised expression matrix to keep only the 665 genes classified as TFs by our previous analysis (see above). We correlated the expression of these TFs against the average expression profile of every WGCNA module (so-called connectivity score in WGCNA). The resulting expression and connectivity matrices were visualised using the ComplexHeatmap R package.

#### WGCNA motif enrichment analysis

We retrieved the promoters of the gene from each WGCNA module and performed motif enrichment analysis using the findMotifsGenome.pl wrapper from the HOMER suite ^113^ with the following parameters: ‘-p 12 -mis 3 -mset vertebrates’, and using as background a subsample of the promoters of every *S. mediterranea* gene that did not belong to the queried module. We concatenated the motif enrichment results into a single matrix which we filtered to keep only motifs with qvalue < 0.1 . Next, we manually curated the association of significant motifs and high-connectivity TFs by inspecting, for a given module X, if the sequence logos of the motifs enriched in promoters of module X resembled those of any group of TFs highly connected to module X. For this, we contrasted the HOMER sequence logos to the JASPAR 2024 database ^114^. These results were visualised using the ggplot2 (https://link.springer.com/book/10.1007/978-3-319-24277-4) and ComplexHeatmap R packages ^112^.

#### WGCNA graph analysis

We generated a large graph taking the TOM matrix as an input adjacency matrix using the R package igraph ^109^. We pruned lowly-scored interactions and used a threshold that maximised the number of evenly-sized connected components. We calculated cross-connections between modules by constructing a gene x module matrix used to count how many genes from each module are direct neighbours to a given gene x. We normalised this matrix by dividing the number of connections of gene x to each module by the size of the module that gene x is part of. These numbers were later aggregated at the module level to retrieve the number of normalised cross-connections between modules. The resulting matrix was transformed into a graph using ‘graph_from_adjacency_matrix()’ from igraph with parameters ‘mode = “upper”, weighted = TRUE, diag = FALSÈ, and the number of cross-connections was used for edge size to highlight the largest amounts of cross-connections.

In addition to this, we created three more module-wise graphs. We first correlated the motif enrichment profile of each module (specifically, the percentage of regions -in this case, gene promoters-with enrichment of each significant detected motif) and used this as an adjacency matrix for a module-wise motif enrichment graph. We then performed functional category enrichment of the genes of every module to obtain a module x functional category enrichment matrix using a custom wrapper of Fisher’s test. We correlated the functional category enrichment profile of each module (specifically, the percentages of enrichment) and used this as an adjacency matrix for a module-wise functional category graph. Lastly, we correlated the TF connectivity profile of each module and used this as an adjacency matrix for a module-wise TF connectivity similarity graph.

We merged these four graphs using igraph and retained pairwise module connections detected in at least two of our four analyses to generate a module-wise similarity graph. We detected communities of modules using the function ‘cluster_label_prop()’ and the list of instances of co-occurring module-module edges as weights for the parameter ‘weights’.

#### Gene Ontology Enrichment analysis

Gene Ontology Enrichment analysis was run using a custom wrapper of the R package ‘topGÒ ^115^, using all the genes with detected expression in our pseudobulk expression matrix (see above) and the classicFisher test with elim algorithm. Gene Ontology terms with less than five annotated genes in the whole genome of *S. mediterranea* (see eggNOG functional annotation above) were discarded. These results were visualised using the ggplot2 ^116^.

#### Pseudobulk computational dissection of scRNA (with conditions and/or replicates)

We created a pseudo-bulk supermatrix leveraging the cell annotation data derived from our scRNA-Seq analysis (e.g. clustering and broad cell types) along with additional cell information like sample characteristics (e.g. experiment, condition, replicates). This was achieved by aggregating, for a given gene X, cell type Y, experiment I, and replicate J, all the counts of gene X from cells belonging to the same cluster Y and under identical conditions (experiment I and replicate J). This process effectively generated a supermatrix with genes in rows and ‘pseudo-samples’ (combinations of cell type, experiment, and replicate) in columns. These pseudo-samples encompassed various combinations of cell types and conditions, such as RNA interference treatment and replicates (e.g. biological, technical, library etc.).

#### Differential Gene Expression Analysis (one-versus-neoblasts)

We performed DGE analysis to compare differentiated cell types against neoblasts as follows: For a given contrast (i.e. comparison of cell type X vs neoblast), we first extracted the relevant pseudo-samples from the pseudo-bulk supermatrix. Secondly, because we were comparing cell types, we used “cell type” as conditions and we used the different batches of experiments (libraries 1-3, libraries 8.3 and 8.4, libraries 11.3 and 11.4, libraries 14.3 and 14.4, and libraries 23.1 and 23.2) as replicates. Third, we ran DESeq2 ^82^ inside a custom wrapper with a contrast of “condition 1” (cell type X) relative to “condition 2” (neoblasts). Genes were identified as differentially expressed if having a p-adjusted below 0.05 (negative binomial test).

### Single Cell ATAC-seq Analysis

#### Generation of scATAC Matrices

The sc-ATACseq library was mapped using CellRanger ^60^. We created a reference index for CellRanger using cellranger mkref and providing the genome fasta and the GTF (see above). We then mapped the scATAC-seq reads using ‘cellranger-atac’ with the reference index and standard parameters to create a region x cell sparse matrix for downstream analysis.

#### Seurat/Signac analysis

The resulting region x cell sparse matrix was loaded onto Seurat and Signac alongside the genome annotation used for mapping, to create a chromatin assay using the function ‘CreateChromatinAssay()’ and parameters ‘min.features = 45’, after which we turned into a Seurat object. We ran the function ‘NucleosomeSignal()’ with standard parameters to map signal from nucleosomes, and subset the Seurat object with the following parameters: ‘peak_region_fragments < 1000 & pct_reads_in_peaks > 15 & nucleosome_signal < 1’. We identified neighbours for the k-nn graph using the function ‘FindNeighbors()’ and the following parameters: ‘reduction = “lsi”, dims = 2:30, k.param = 10’. We identified clusters using the function ‘FindClusters()’ with the Louvain algorithm ^108^ and the following parameters: ‘resolution = 1, algorithm = 3’. We computed an UMAP projection using the function ‘RunUMAP()’ and the following parameters: ‘dims = 2:30, reduction = “lsi”, seed.use = 1’ . To find cell markers in the scATAC data (either in the “peaks” or “RNA” assays), we used the function ‘FindAllMarkers()’ with the following parameters: ‘only.pos = TRUE, min.pct = 0.1, logfc.threshold = 0.1’.

#### Alignment between datasets

To identify the cell type of the scATAC cells, we aligned the two datasets to transfer the scATAC seurat cluster identity to cells in the scRNA object. We did it this way because the scATAC-seq data had lower resolution. This in turn allows us to know what broad cell type of the scRNA data corresponds to scATAC clusters. We first calculated the Gene Activity function using the ‘GeneActivity()’ function of signac with standard parameters. We normalised the resulting Gene Activity data using the function ‘NormalizeData()’ and the parameters: ‘normalization.method = “LogNormalize”, scale.factor = median(nCount_RNA)’, where ‘nCount_RNÀ was the number of counts detected per cell in the scATAC Seurat object. We intersected the scRNA object variable features and the genes detected with gene activity in the scATAC object, and used these as common features for aligning the datasets using the function ‘FindTransferAnchors()’ with the following parameters: ‘reduction = “cca”, k.anchor = 5, k.filter = NA, k.score = 10, max.features = 1000’, and using the scRNA object as query and the scATAC as reference. We transferred the labels using the ‘TransferData()’ function with the following parameters: ‘weight.reduction = scrna_pca, dims = 2:30’, where ‘scrna_pcà was the PCA calculated for the scRNA object. The predicted labels of each cell in the query dataset were added as a metadata column. We assigned the most frequent predicted label (the scATAC cluster) to each cell type cluster we annotated in the scRNA dataset. We manually annotated the scATAC clusters after inspecting the results of the label transferring. Label transferring was visualised using igraph ^109^.

#### Pseudobulk computational dissection of scATAC-Seq (for differential chromatin accessibility analysis)

We created a pseudo-bulk supermatrix with OCRs in rows and “pseudo-samples” in columns in the same fashion as described above. For replicates, we randomly split the cells into two pseudoreplicates. For one-versus-all analyses, we ran independent pseudo-bulk count aggregations labelling every non-cell of interest as “else”, aggregating counts from cells different from the cell type of interest (e.g. muscle cells vs non-muscle cells).

#### Differential chromatin accessibility analysis (one-versus-all)

We performed differential chromatin accessibility analysis (DCA) to compare each differentiated cell type against the rest as follows: For a given contrast (e.g. muscle vs everything else), we first ran the pseudobulk count aggregation as described above. Secondly, we ran DESeq2 ^82^ inside a custom wrapper with a contrast of “condition 1” (cell type X) relative to “condition 2” (everything else). Genes were identified as differentially expressed if having a p-adjusted below 0.05 (negative binomial test).

#### Differential chromatin accessibility analysis (one-versus-neoblasts)

We performed differential chromatin accessibility analysis to compare differentiated cell types against neoblasts as follows: For a given contrast (i.e. comparison of cell type X vs neoblast), we first extracted the relevant pseudo-samples from the pseudo-bulk supermatrix of scATAC-seq. Secondly, because we were comparing cell types, we used “cell type” as conditions and the pseudoreplicates as replicates. Third, we ran DESeq2 ^82^ inside a custom wrapper with a contrast of “condition 1” (cell type X) relative to “condition 2” (neoblasts). Genes were identified as differentially expressed if having a p-adjusted below 0.05 (negative binomial test).

#### One-versus-all OCR Motif Enrichment Analysis

We performed motif enrichment analysis using the ‘findMotifsGenome.pl’ wrapper from the HOMER suite ^113^ with standard parameters, using the OCRs of interest as foreground and the rest of OCRs detected by cellranger as background. We concatenated the motif enrichment results into a single matrix which we filtered to keep only motifs with qvalue < 0.1 . These were visualised using the ggplot2^116^ and ComplexHeatmap R packages ^112^.

#### Association of OCRs to genes

We associated every OCR to a gene using ‘bedtools closestbed’ ^102^ with the following parameters:‘-k 1 -D ref -a all_peaks_sorted.bed -b TSS.bed’, where ‘all_peaks_sorted.bed’ is the BED file of all OCR coordinates and ‘TSS.bed’ is the BED file of the TSS coordinates of all genes.

#### Differentially accessible OCR WGCNA analysis

We first ran a pseudobulk aggregation without replicates or conditions, only the cell types, as described above. The resulting matrix of OCR counts was normalised by “library size” using the ‘DESeqDataSetFromMatrix()’ function with parameter ‘design = ∼ condition’ and the ‘counts()’ function with parameter ‘normalised = TRUÈ from the R package DESeq2 ^82^. We subsetted this matrix to keep all the OCRs detected as significantly open (log2FC > 0) in any of the differentiated cell types of our DCA analysis. We performed cell type co-occurrency as described above using this normalised matrix to retrieve a tree of cell type similarity based on their profile of chromatin accessibility in these differentially accessible OCRs. We scaled this matrix across rows and ran WGCNA using soft power 8 after visual inspection of the scale-free topology fit and median connectivity dynamics. The resulting TOM matrix was turned into a dissimilarity matrix and clustered using the R function ‘hclust’ with parameters ‘method = “ward.D2”’ . We cut the clustering in different modules of co-accessibility using the function ‘cutreeDynamic()’ from the WGCNA package ^68^ and the following parameters: ‘deepSplit = 4, pamRespectsDendro = FALSE, minClusterSize = 30’. We reclassified the modules as described above for the WGCNA co-expression modules. To assess the likeness in expression of the genes associated to these OCRs, we ran the pseudobulk aggregation with broad cell type labels on the scRNA-seq dataset, and normalised the counts as described above. We then subsetted and reordered this matrix to keep the genes associated to the differentially accessible OCRs used in the ATAC WGCNA analysis. We correlated the expression profile of every gene with the average accessibility profile of the ATAC module of their associated OCRs and retrieved the top 20 highly-correlating genes with each module for visualisation using the ComplexHeatmap package.

#### Bulk ATAC-seq analysis

We mapped the bulk ATAC-seq reads using a custom perl wrapper of bowtie2 as described in Pérez-Posada *et al*. ^117^. We then called for peaks using a custom bash wrapper of MACS2 as described in Pérez-Posada *et al*. ^117^, with the following parameters: ‘-f BED --nomodel --extsize 100 --shift 45 -- buffer-size 50000 -g 840213658 -p 0.001’.

### Gene Track and Chromatin Visualisation

#### Generation of gene annotation tracks

We parsed the GTF gene annotation track using GenomicRanges and rtracklayer ^118^ in R, and transformed it to a suitable format using the R package GenomicRanges ^119^.

#### Generation of scRNA-seq alignment tracks

For every gene_function_tagged BAM file of our scRNA-seq libraries, we ran ‘sintò (https://github.com/timoast/sinto) to split them into different bam files, one per broad cell type, by filtering reads labelled with barcodes from cells assigned to that cell type, using the following parameters: ‘-barcodetag XC’. After splitting, we merged all the bam files from different libraries but from the same cell type to create one unique alignment file per cell type. We used bamCoverage to convert them to BigWig files with standard parameters parameters. BigWig files were visualised using the ‘GenomicRanges’ and ‘gviz’ R packages ^119, 120^.

#### Generation of scATAC-seq-seq alignment tracks

We split the scATAC-seq BAM file generated by ‘cellranger-atac’ using ‘sintò and the list of cell barcodes labelled for each cell type, to generate independent BAM files for each cell type. We used bamCoverage to convert them to BigWig files with the following parameters: ‘--normalizeUsing BPM’. BigWig files were visualised using the ‘GenomicRanges’ and ‘gviz’ R packages ^119, 120^.

#### Genomic Coordinates visualisation

We retrieved the genomic coordinates of scATAC-seq markers and plotted them using a custom gviz^120^ wrapper in R. These were manually inspected.

#### Chromatin enrichment profiles

Chromatin plots were generated and visualised using the GenomicRanges ^119^and EnrichedHeatmap package ^121^.

### ANANSE analysis

#### Chromatin pre-processing

We used the same BAM files generated using sinto (https://github.com/timoast/sinto) as input for ANANSE binding. For a peak catalogue file, we re-centered the coordinates of the peaks called by cellranger-atac around the summit of the signal using a custom python wrapper of ‘bedtools slop’ ^102^ and ‘samtools mpileup’ ^122^. We kept peaks with a minimum of 5 counts detected.

#### Gene expression pre-processing

We generated independent tables of normalised counts for each broad cell type using the pseudobulk approach without replicates or conditions, only broad cell type labels, as described above.

#### ANANSE binding

For each broad cell type, we ran ANANSE binding ^59^ with parameters ‘--jaccard-cutoff 0.2’, providing the genome fasta, the respective broad cell type BAM file, the summit-centered peak file, and the motif2factors lookup database we generated using gimme motif2factors (see above).

#### ANANSE network

For each broad cell type, we ran ANANSE network ^59^ with parameters: ‘--full-output’, providing the genome fasta, the respective broad cell type counts table, the GTF annotation of S. mediterranea, and the resulting .h5 ANANSE binding file for the respective cell type.

#### ANANSE influence

We ran ANANSE influence ^59^ for every pair of contrasts (from neoblast network to a differentiated cell type X), using the following parameters: ‘-n 12 -i 250000’. The resulting output (influence data frame and differential network) were analysed in R using ‘igraph’ ^109^.

#### ANANSE network pre-processing, graph analysis and visualisation

The resulting ANANSE networks were imported into R and analysed as follows. For every network, we subsetted the network to keep interactions above score value 0.8. We created graph objects using igraph and removed genes without neighbours.

We calculated centrality, out-centrality, in-degree, and out-degree for every gene using igraph. Relative out-degree was calculated as in-degree divided by the sum of in-degree and out-degree. We calculated the number of active TFs as the number of genes with outdegree above 0. For every network, we extracted the centrality values for all the TF genes in the network. We collapsed these values together in a TF x cell type network matrix of centrality values. We correlated the columns of this matrix using the following parameters: ‘method = “ward.D2”’. For graph visualisation, we subsetted the graphs to keep the influential TFs from the ANANSE influence analysis, and we kept the top 2 interactions (based on ANANSE’s prob score) of each TF. The resulting graphs were plotted using the graphopt algorithm from igraph ^109^.

For target visualisation of each network, we first retrieved the target genes of each of the top five TFs of each fate. For each TF in each fate, we kept interactions with target genes above the .95 quantile (top 5% interactions). We visualised them using the ‘geom_jitter()’ function from the R package ggplot2 ^116^. Names of target genes detected in the literature from PlanMine ^90, 91^ were manually curated and shown next to the stripchart.

To detect groups of co-influential TFs, we collapsed the influence score values of each TF in the network of each fate into a single TF x cell fate influence score matrix. We clustered these TFs using Pearson correlation of their influence profile and the method ‘ward.D2’. The resulting tree was cut above 0.7 to retrieve small clusters of TFs with similar influence profiles across fates. These clusters were automatedly sorted as described above for WGCNA modules. Within each cluster, we correlated every TF with the average co-influence profile of their parent cluster, and sorted them in descending order. We chose the top five TFs for each cluster for visualisation using the R package ‘ComplexHeatmap’ ^112^.

### HNF4a Knock-down scRNA-seq analysis

#### scRNA-seq pre-processing and generation of scRNA matrices

We pre-processed the reads of libraries 11.3 and 11.4, from the *hnf4*i experiment, as discussed above. To generate scRNA matrices for each library, the resulting outputs of running the SPLiT-Seq pipeline (tag_bam_with_gene_function) were used to generate a gene x cell sparse matrix keeping cells with a minimum of 100 genes detected using the command ‘DigitalExpression’ from dropseqtools.

#### Matrix concatenations

The resulting matrices were concatenated using Seurat v4.3.0 ^61^ in R v4.0.3. Briefly, we created independent Seurat Objects for each separate library and performed Proportional Fitting-log1p-Proportional Fitting normalisation ^106^ on each separate matrix. We ran ‘FindVariableFeatures’ using the parameters: ‘selection.method = “vst”, features = 20000’. We labelled cells from each library with their respective library ID (Supplementary File 19) and merged all the Seurat objects into a single object with a concatenated sparse matrix.

#### Seurat Analysis

The concatenated sparse matrix was queried for highly variable genes using ‘SelectIntegrationFeatures()’ from the Seurat package. We scaled the data and performed PCA using the function ‘runPCA()’ with the following parameters: ‘npcs = 50’ . We integrated the data using the function ‘runHarmony()’ from Harmony ^107^ and the following parameters: ‘dims.use = 1:50, theta = 3, lambda = 3, nclust = 40, max.iter.harmony = 20’. We identified neighbours for the k-nn graph using the function ‘FindNeighbors()’ and the following parameters = ‘dims = 1:50, k.param = 65’. We identified clusters using the function ‘FindClusters()’ with the Louvain algorithm ^108^ and the following parameters: ‘resolution = 2.5, algorithm = 1, random.seed = 75’. We computed an UMAP projection using the function ‘RunUMAP()’ and the following parameters: ‘dims = 1:50, reduction = “harmony”, n.neighbors = 65, min.dist = 0.5, spread = 1, metric = “euclidean”, seed.use = 1’.

#### Alignment to reference dataset

To align the *hnf4*i dataset to our whole atlas analysed in this study, we ran the function ‘FindTransferAnchors()’ from Seurat using the *hnf4*i Seurat object as query and the Seurat object of our scRNA atlas of thirteen libraries as the reference, and the following parameters: ‘dims = 70’. The predicted labels of each cell in the query dataset were added as a metadata column. We assigned the most frequent predicted label to each Seurat cluster we obtained. Upon inspection and downstream analyses, we manually curated the assignment of phagocyte progenitors from the *hnf4*i libraries (as opposed to the control libraries) as aberrant phagocyte progenitors. Label transferring was visualised using igraph ^109^.

#### Cell abundance analysis

To calculate the over- or under-representation of cells from experimental condition in our dataset, we performed pre- and post-hoc Chi-Squared test as described previously ^57^using a custom R wrapper. These resulting residuals were visualised using ComplexHeatmap ^112^.

#### Gene score analysis

We first retrieved markers for each cell type of interest (neoblasts, phagocyte progenitors, phagocytes, epidermis) using the function ‘FindMarkers’ from Seurat and the following parameters: ‘only.pos = TRUE, return.thresh = 1, logfc.threshold = 0’, and from this retrieved the top 50 markers based on log fold change. For each set of markers of interest, we calculated a gene score as follows: for each cell, we first calculated the average expression of the genes of interest; we then calculated the average expression of a random subsample of five percent of all genes; then we subtract the second value from the first value. These values, one per cell, were pooled based on cell cluster and experimental condition, and plotted and visualised using ggplot2 ^116^.

#### Differential Gene Expression Analysis

To perform differential gene expression (DGE) analysis on each cell type separately, we first computationally dissected the *hnf4*i dataset to retrieve a super matrix of genes in rows and “pseudo-samples” in columns, each “pseudo-sample” being a combination of cell type, experimental condition, and replicate, as described above. For each type, we first subsetted this supermatrix to extract the columns relevant to the cell type of interest. We then filtered out genes with less than one count in at least two replicates. We then ran DGE analysis using a custom R wrapper of DESeq2 ^82^, with the following contrast: ‘c(“condition”,”HNF4i”,”control”)’. Genes were identified as differentially expressed genes (DEGs) if having a p-value below 0.05 (negative binomial test). For each cell type, the results of DESeq2 (log2FC and -log p value) were plotted using ggplot2 ^116^. This analysis was performed first using broad cell types, but also using the individual, narrow cell types.

#### Downstream analysis of Differentially Expressed Genes

We retrieved the lists of DEGs from the cell types with the highest fraction of DEGs in their analysis. We calculated the overlap between these using the eulerr R package (For each cell type, the results of DESeq2 (log2FC and -log p value) were plotted using ggplot2 ^116^. For each set (exclusive to phagocytes, exclusive to parenchyma, common to both, or all of them together) performed Gene ontology enrichment and motif enrichment analyuses as described above.

To visualise the *hnf4* interaction predicted by ANANSE on the DEGs, we imported the ANANSE networks of phagocytes and parenchyma into R, and subsetted them to keep only interactions stemming from *hnf4*. We labelled these genes as DEG in each of the sets (exclusive to phagocytes, exclusive to parenchyma, or common to both) and visualised their predicted score.

## Acknowledgements

Research at the Solana lab at Oxford Brookes University and at the Living Systems Institute is supported by MRC grants (MR/S007849/1 and MR/W017539/1), a BBSRC Grant (BB/V014447/1) and a Leverhulme Trust grant (RPG-2019-332 and RPG-2023-330) to JS. HG-C and EE were supported by Nigel Groome studentships from Oxford Brookes University. S.F. and S.J.v.H. were supported by the Netherlands Organization for Scientific Research (NWO grant 016.Vidi.189.081) to S.J.v.H. We thank Isabel Liao for advice and discussions with the weight normalisation, María Rosselló for advice and discussions with the differential gene expression analysis, and all members of the Solana lab for useful discussion, input and assistance. We thank the Wellcome Centre for Human Genetics for their expertise in generating the scATAC-seq dataset, especially Rory Bowden and Hubert Slawinski. Flow cytometry was performed at the Sir William Dunn School of Pathology Flow Cytometry Facility, University of Oxford with the assistance of Dr Robert Hedley.

## Data availability

The datasets supporting the conclusions of this article are available in:

GEO (whole project, including scRNA-seq reads, scATAC-seq reads, ATAC-seq reads, as well as processed files such as .RDS Seurat objects): GSE274286

Code: https://github.com/scbe-lab/regulatory_logic

## Author contributions

J.S. conceived the study, designed the experiments, and provided supervision. H.G.C. and E.E. generated cell dissociations and performed single-cell transcriptomic experiments using *Schmidtea mediterranea*, both in unperturbed and knock-down animals. J.S. generated the library of scATAC-Seq. V.V. generated the bulk ATAC-Seq library. C.A.B. and N.J.K. performed preliminary bioinformatic analyses. S.F. performed preliminary ANANSE bioinformatic analyses. J.S. and A.P.P. designed the final bioinformatic analyses. A.P.P. performed all the final bioinformatic analyses. S.F. and S.J.V.H. contributed to the interpretation of the ANANSE network data. A.P.P. generated the figures. A.P.P. and J.S. wrote the manuscript, with contributions from all other authors. All authors read and approved the final version of the manuscript.

