## Supplementary material for "The Regulatory Logic of Planarian Stem Cell Differentiation": supp_file_06.pdf

### Module s01

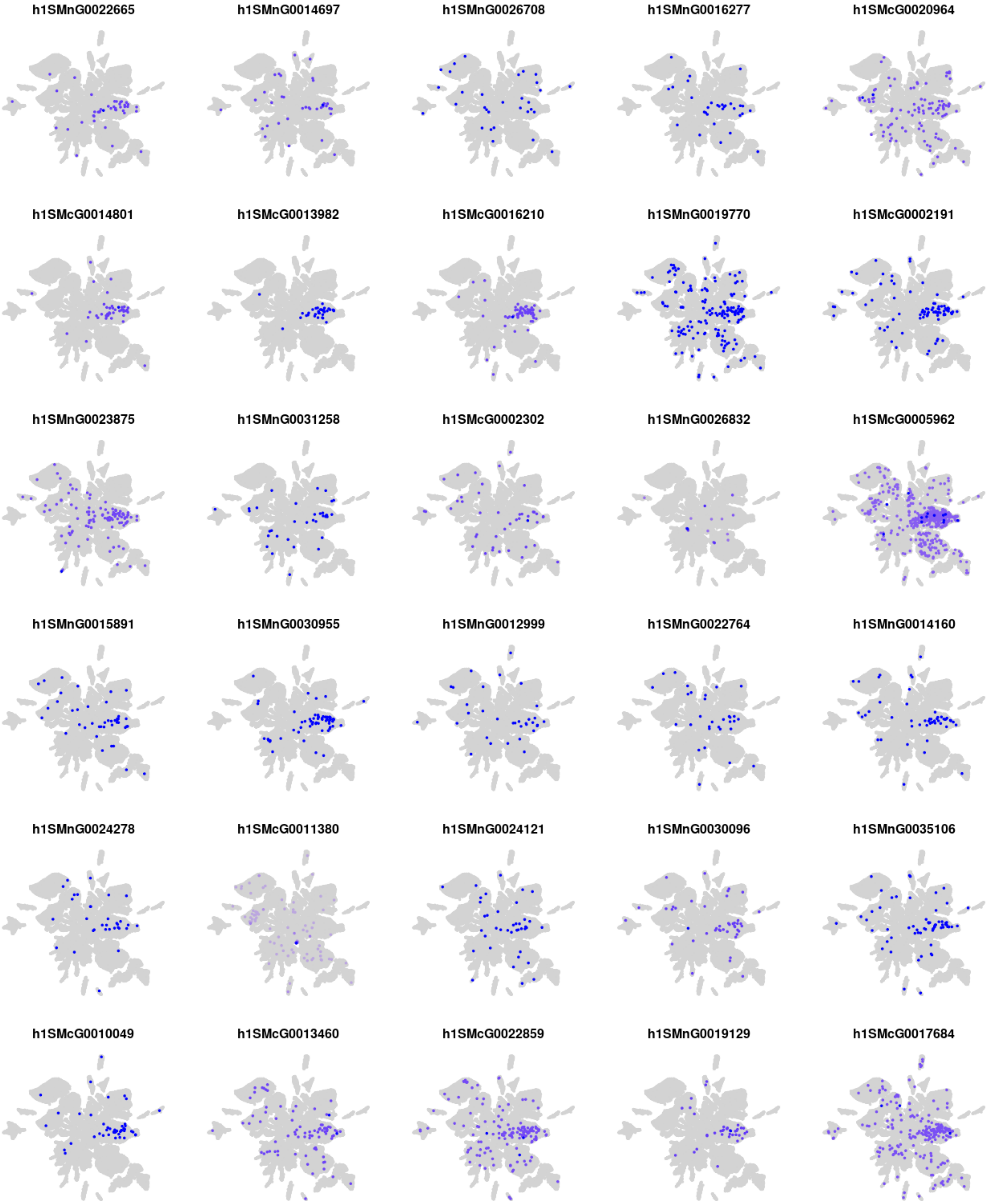
h1SMnG0022665
h1SMnG0014697
h1SMnG0026708
h1SMnG0016277
h1SMcG0020964
h1SMcG0014801
h1SMcG0013982
h1SMcG0016210
h1SMnG0019770
h1SMcG0002191
h1SMnG0023875
h1SMnG0031258
h1SMcG0002302
h1SMnG0026832
h1SMcG0005962
h1SMnG0015891
h1SMnG0030955
h1SMnG0012999
h1SMnG0022764
h1SMnG0014160
h1SMnG0024278
h1SMcG0011380
h1SMnG0024121
h1SMnG0030096
h1SMnG0035106
h1SMcG0010049
h1SMcG0013460
h1SMcG0022859
h1SMnG0019129
h1SMcG0017684

### Module s02

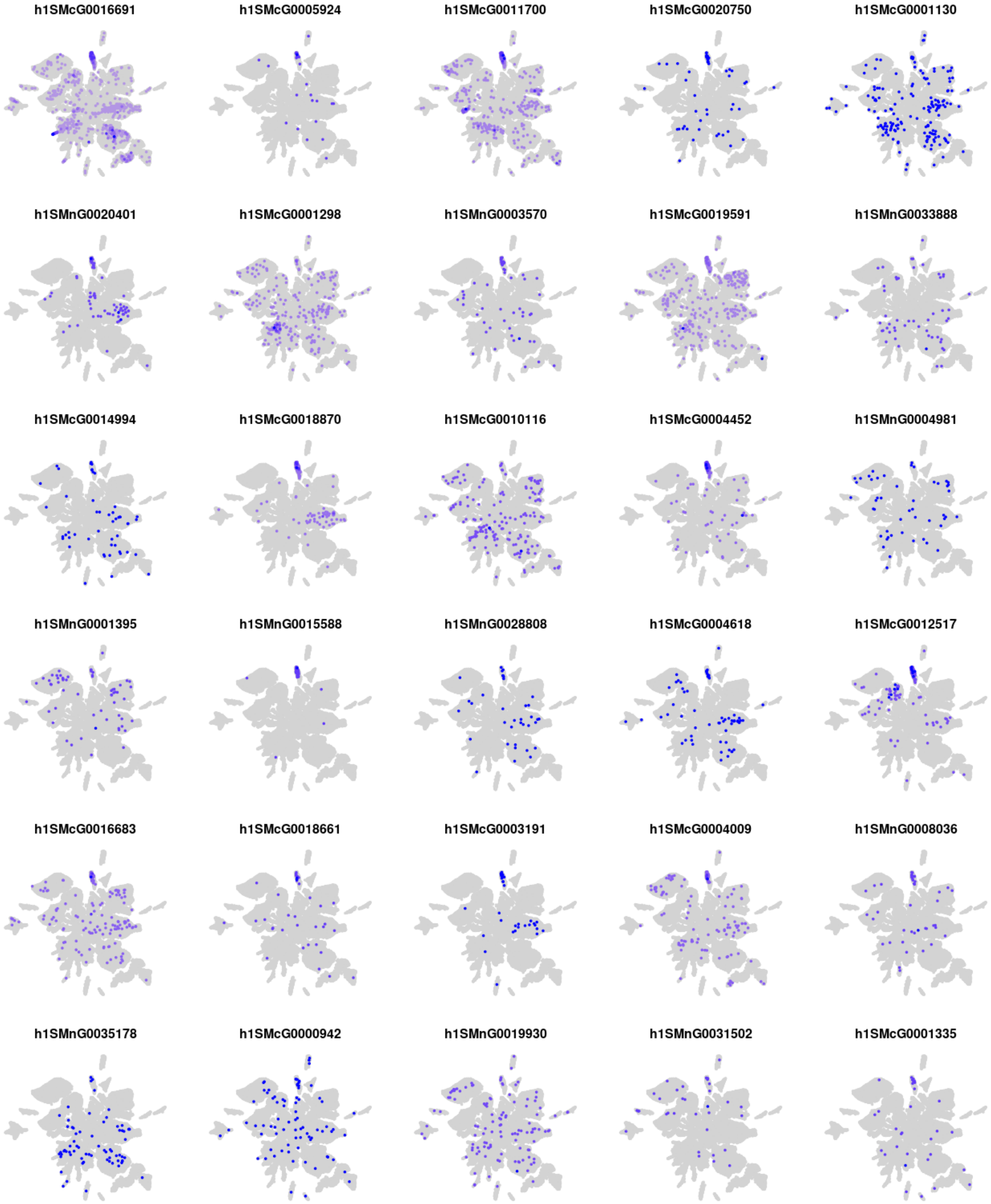
h1SMcG0016691
h1SMcG0005924
h1SMcG0011700
h1SMcG0020750
h1SMcG0001130
h1SMnG0020401
h1SMcG0001298
h1SMnG0003570
h1SMcG0019591
h1SMnG0033888
h1SMcG0014994
h1SMcG0018870
h1SMcG0010116
h1SMcG0004452
h1SMnG0004981
h1SMnG0001395
h1SMnG0015588
h1SMnG0028808
h1SMcG0004618
h1SMcG0012517
h1SMcG0016683
h1SMcG0018661
h1SMcG0003191
h1SMcG0004009
h1SMnG0008036
h1SMnG0035178
h1SMcG0000942
h1SMnG0019930
h1SMnG0031502
h1SMcG0001335

### Module s03

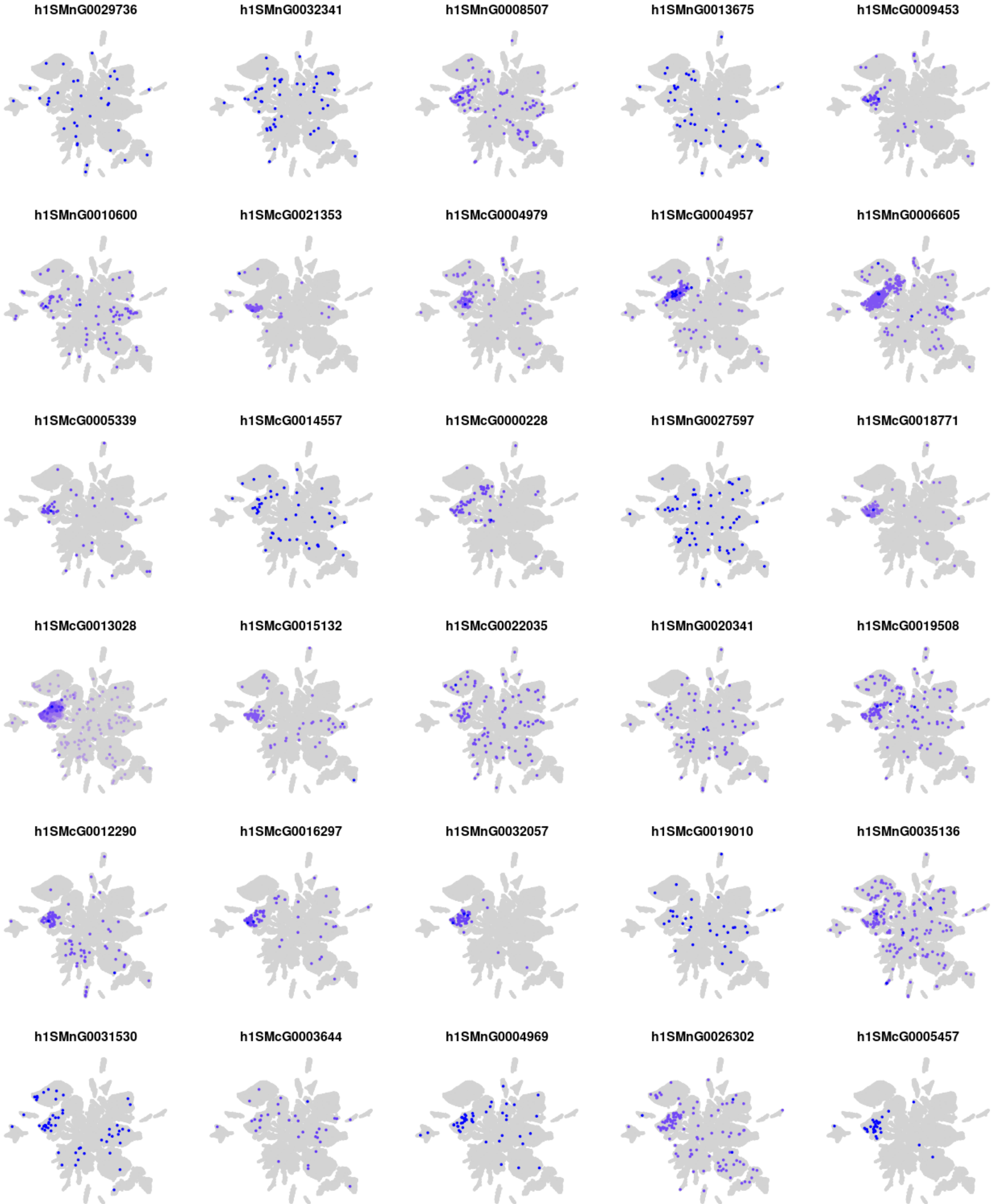
h1SMnG0029736
h1SMnG0032341
h1SMnG0008507
h1SMnG0013675
h1SMcG0009453
h1SMnG0010600
h1SMcG0021353
h1SMcG0004979
h1SMcG0004957
h1SMnG0006605
h1SMcG0005339
h1SMcG0014557
h1SMcG0000228
h1SMnG0027597
h1SMcG0018771
h1SMcG0013028
h1SMcG0015132
h1SMcG0022035
h1SMnG0020341
h1SMcG0019508
h1SMcG0012290
h1SMcG0016297
h1SMnG0032057
h1SMcG0019010
h1SMnG0035136
h1SMnG0031530
h1SMcG0003644
h1SMnG0004969
h1SMnG0026302
h1SMcG0005457

### Module s04

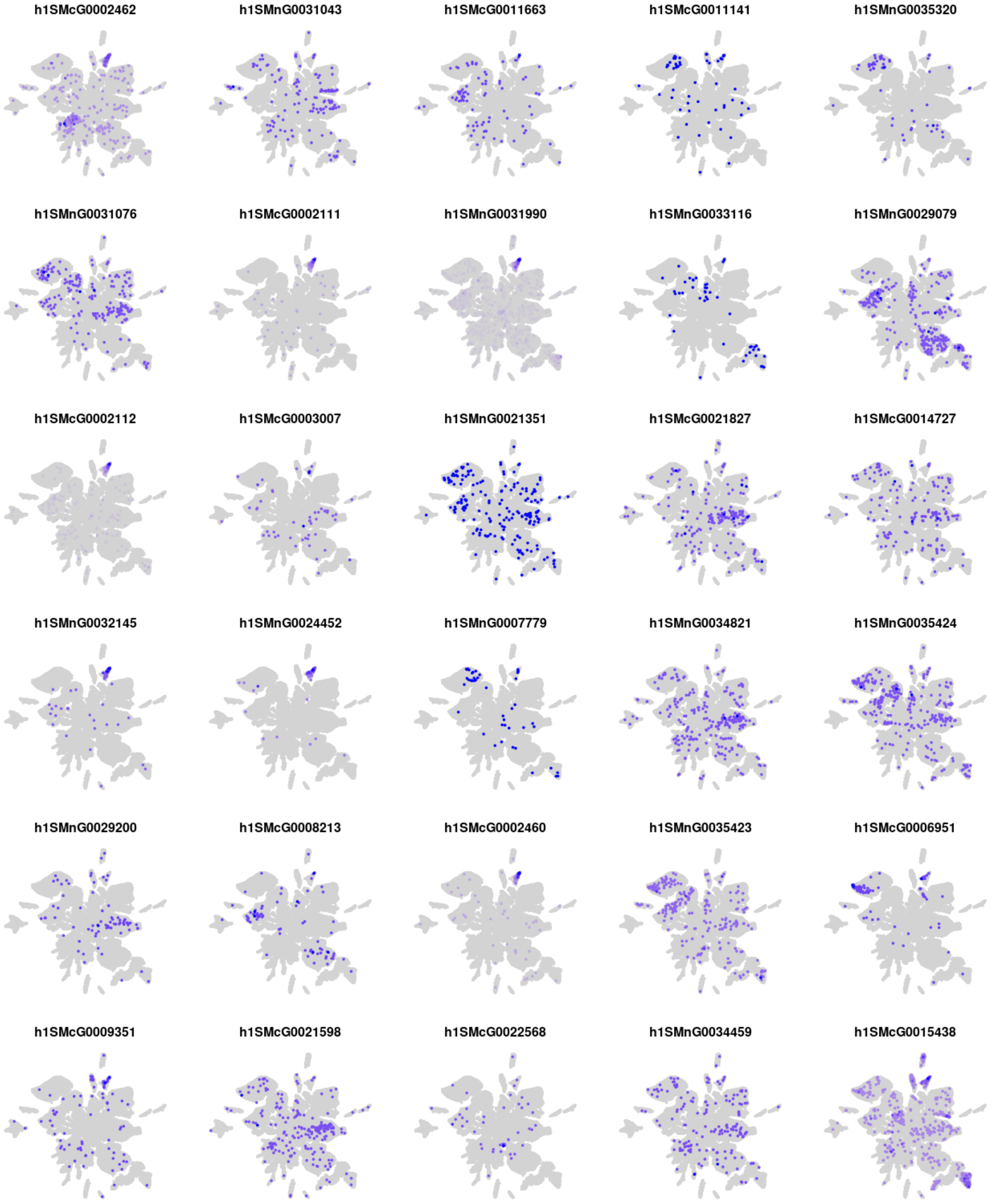
h1SMcG0002462
h1SMnG0031043
h1SMcG0011663
h1SMcG0011141
h1SMnG0035320
h1SMnG0031076
h1SMcG0002111
h1SMnG0031990
h1SMnG0033116
h1SMnG0029079
h1SMcG0002112
h1SMcG0003007
h1SMnG0021351
h1SMcG0021827
h1SMcG0014727
h1SMnG0032145
h1SMnG0024452
h1SMnG0007779
h1SMnG0034821
h1SMnG0035424
h1SMnG0029200
h1SMcG0008213
h1SMcG0002460
h1SMnG0035423
h1SMcG0006951
h1SMcG0009351
h1SMcG0021598
h1SMcG0022568
h1SMnG0034459
h1SMcG0015438

### Module s05

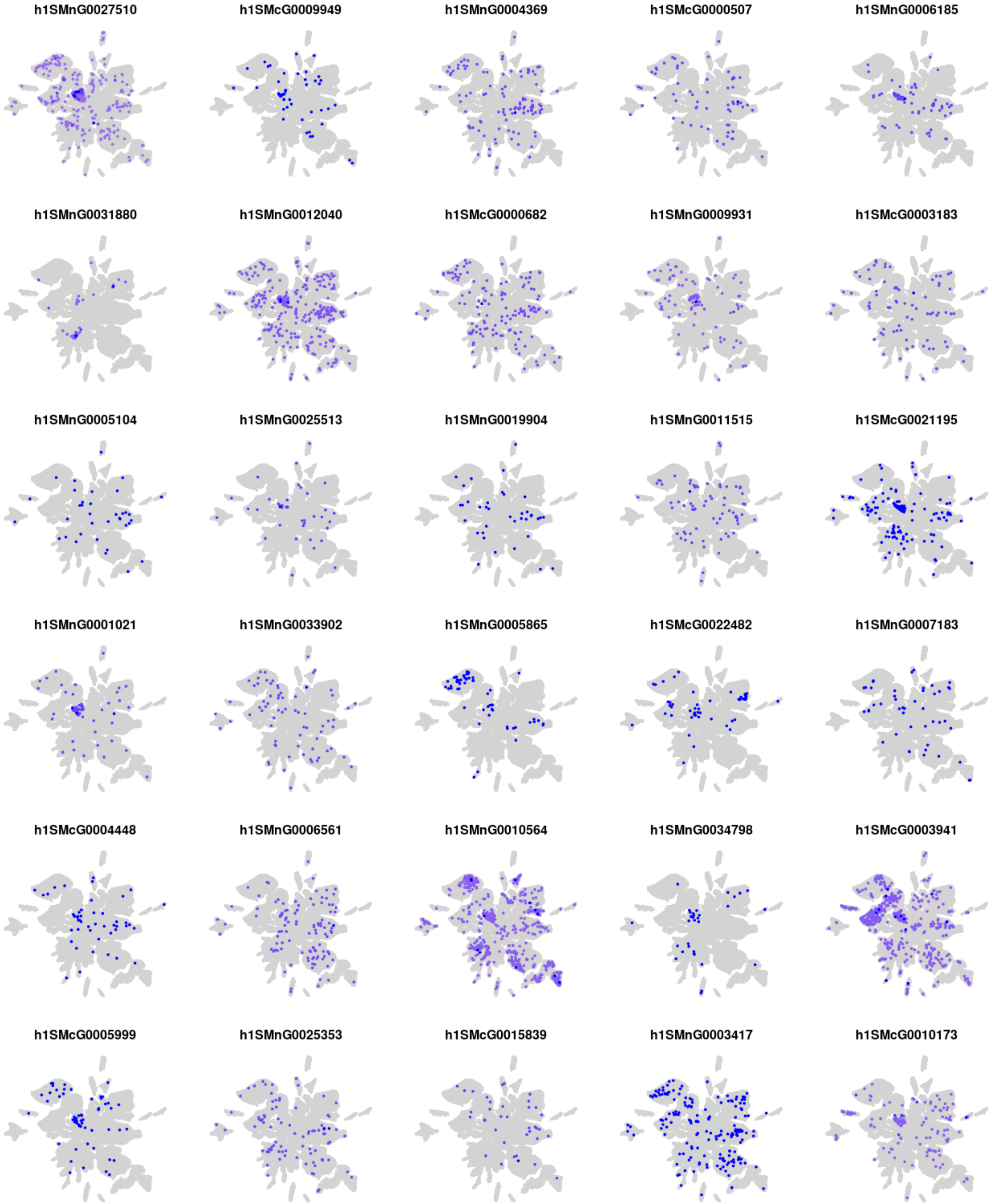
h1SMnG0027510
h1SMcG0009949
h1SMnG0004369
h1SMcG0000507
h1SMnG0006185
h1SMnG0031880
h1SMnG0012040
h1SMcG0000682
h1SMnG0009931
h1SMcG0003183
h1SMnG0005104
h1SMnG0025513
h1SMnG0019904
h1SMnG0011515
h1SMcG0021195
h1SMnG0001021
h1SMnG0033902
h1SMnG0005865
h1SMcG0022482
h1SMnG0007183
h1SMcG0004448
h1SMnG0006561
h1SMnG0010564
h1SMnG0034798
h1SMcG0003941
h1SMcG0005999
h1SMnG0025353
h1SMcG0015839
h1SMnG0003417
h1SMcG0010173

### Module s06

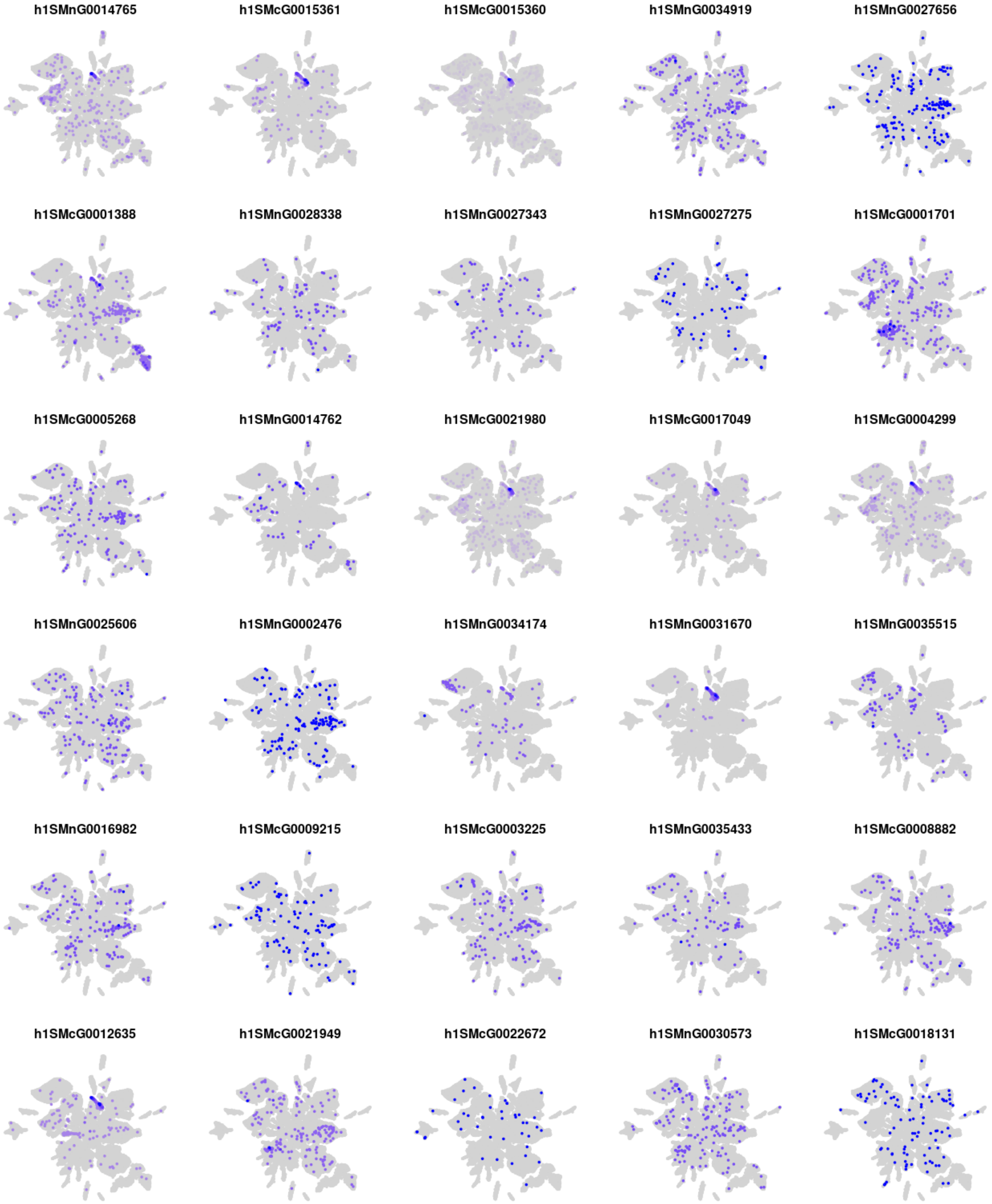
h1SMnG0014765
h1SMcG0015361
h1SMcG0015360
h1SMnG0034919
h1SMnG0027656
h1SMcG0001388
h1SMnG0028338
h1SMnG0027343
h1SMnG0027275
h1SMcG0001701
h1SMcG0005268
h1SMnG0014762
h1SMcG0021980
h1SMcG0017049
h1SMcG0004299
h1SMnG0025606
h1SMnG0002476
h1SMnG0034174
h1SMnG0031670
h1SMnG0035515
h1SMnG0016982
h1SMcG0009215
h1SMcG0003225
h1SMnG0035433
h1SMcG0008882
h1SMcG0012635
h1SMcG0021949
h1SMcG0022672
h1SMnG0030573
h1SMcG0018131

### Module s07

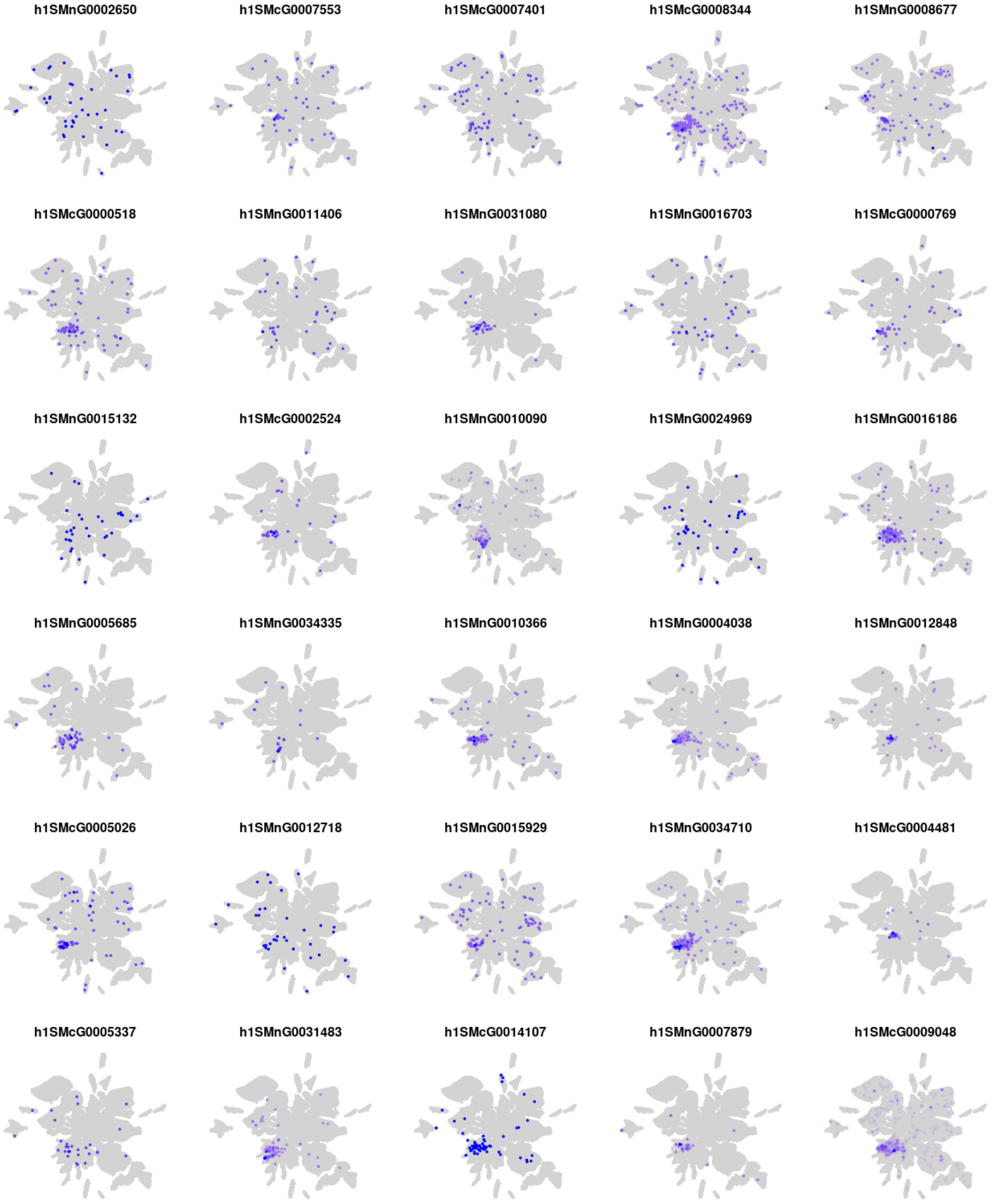
h1SMnG0002650
h1SMcG0007553
h1SMcG0007401
h1SMcG0008344
h1SMnG0008677
h1SMcG0000518
h1SMnG0011406
h1SMnG0031080
h1SMnG0016703
h1SMcG0000769
h1SMnG0015132
h1SMcG0002524
h1SMnG0010090
h1SMnG0024969
h1SMnG0016186
h1SMnG0005685
h1SMnG0034335
h1SMnG0010366
h1SMnG0004038
h1SMnG0012848
h1SMcG0005026
h1SMnG0012718
h1SMnG0015929
h1SMnG0034710
h1SMcG0004481
h1SMcG0005337
h1SMnG0031483
h1SMcG0014107
h1SMnG0007879
h1SMcG0009048

### Module s08

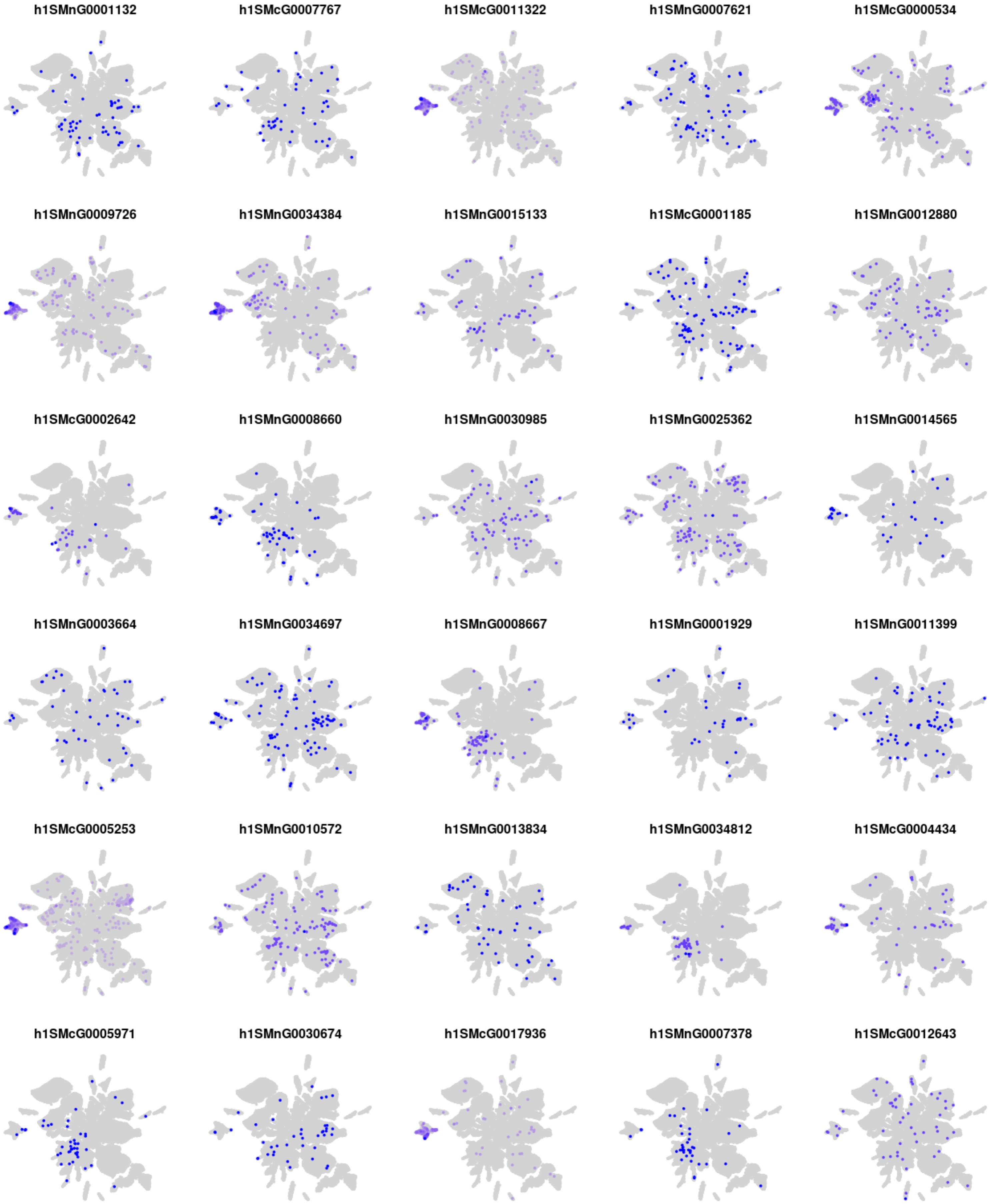
h1SMnG0001132
h1SMcG0007767
h1SMcG0011322
h1SMnG0007621
h1SMcG0000534
h1SMnG0009726
h1SMnG0034384
h1SMnG0015133
h1SMcG0001185
h1SMnG0012880
h1SMcG0002642
h1SMnG0008660
h1SMnG0030985
h1SMnG0025362
h1SMnG0014565
h1SMnG0003664
h1SMnG0034697
h1SMnG0008667
h1SMnG0001929
h1SMnG0011399
h1SMcG0005253
h1SMnG0010572
h1SMnG0013834
h1SMnG0034812
h1SMcG0004434
h1SMcG0005971
h1SMnG0030674
h1SMcG0017936
h1SMnG0007378
h1SMcG0012643

### Module s09

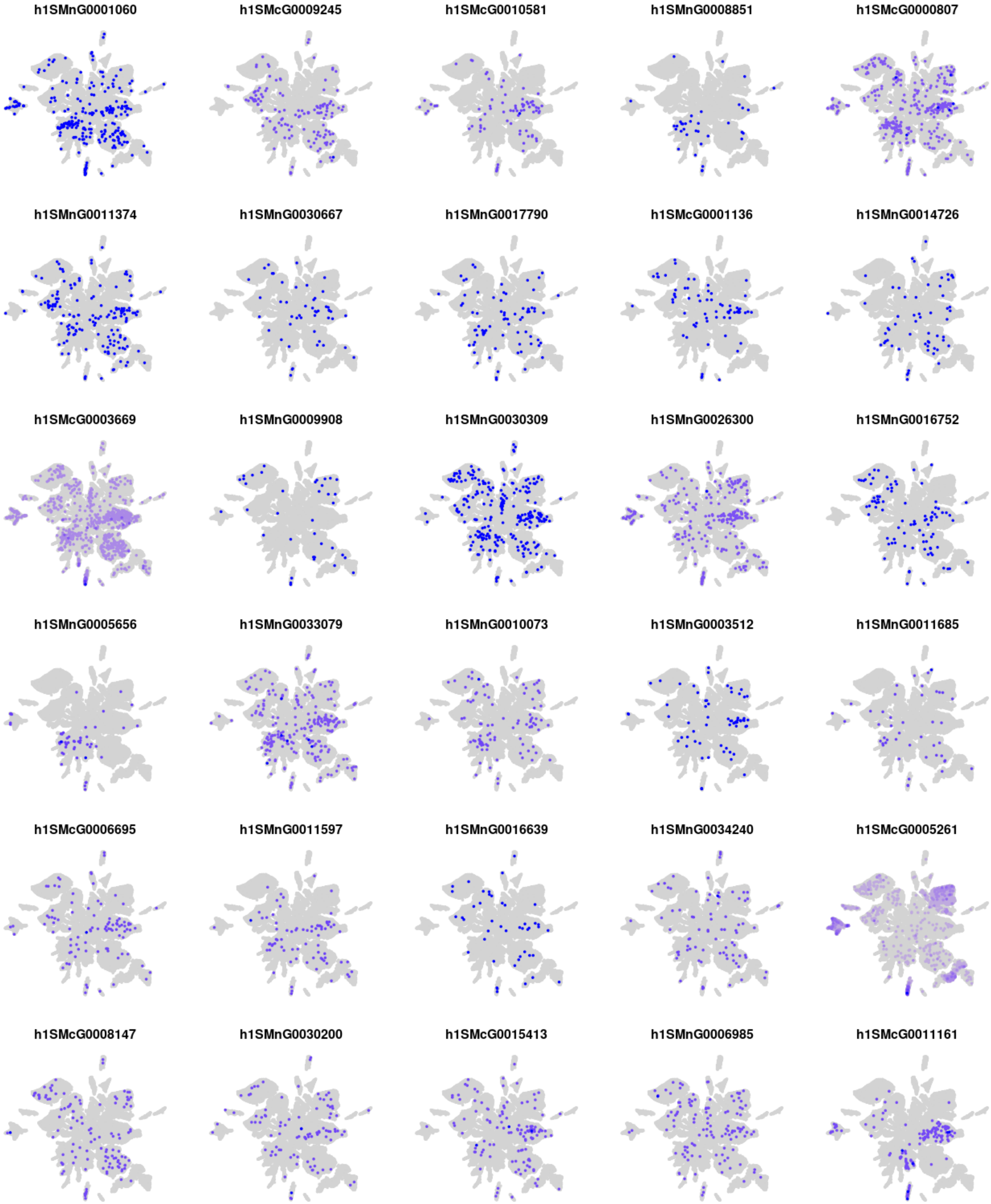
h1SMnG0001060
h1SMcG0009245
h1SMcG0010581
h1SMnG0008851
h1SMcG0000807
h1SMnG0011374
h1SMnG0030667
h1SMnG0017790
h1SMcG0001136
h1SMnG0014726
h1SMcG0003669
h1SMnG0009908
h1SMnG0030309
h1SMnG0026300
h1SMnG0016752
h1SMnG0005656
h1SMnG0033079
h1SMnG0010073
h1SMnG0003512
h1SMnG0011685
h1SMcG0006695
h1SMnG0011597
h1SMnG0016639
h1SMnG0034240
h1SMcG0005261
h1SMcG0008147
h1SMnG0030200
h1SMcG0015413
h1SMnG0006985
h1SMcG0011161

### Module s10

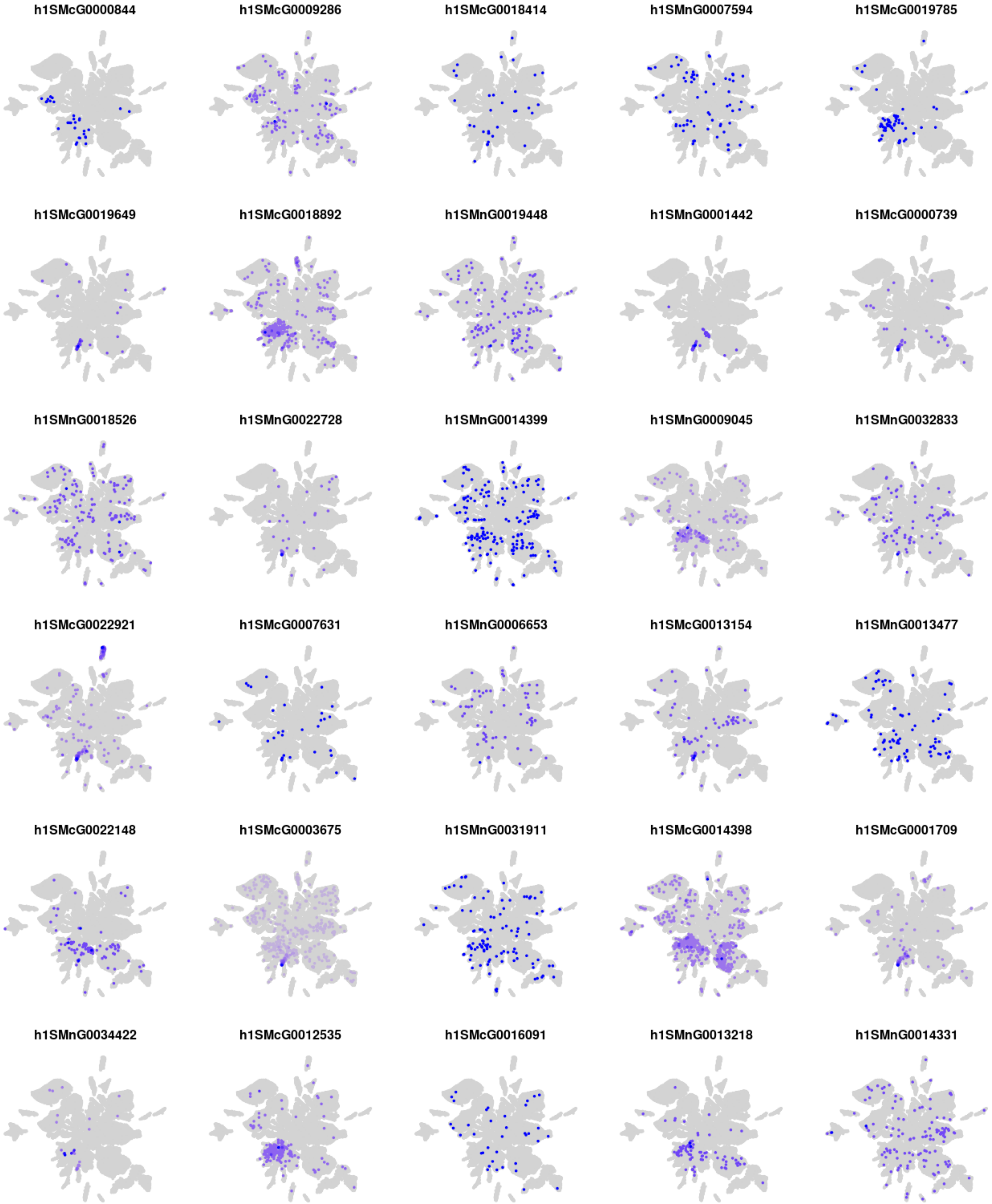

h1SMcG0000844
h1SMcG0009286
h1SMcG0018414
h1SMnG0007594
h1SMcG0019785
h1SMcG0019649
h1SMcG0018892
h1SMnG0019448
h1SMnG0001442
h1SMcG0000739
h1SMnG0018526
h1SMnG0022728
h1SMnG0014399
h1SMnG0009045
h1SMnG0032833
h1SMcG0022921
h1SMcG0007631
h1SMnG0006653
h1SMcG0013154
h1SMnG0013477
h1SMcG0022148
h1SMcG0003675
h1SMnG0031911
h1SMcG0014398
h1SMcG0001709
h1SMnG0034422
h1SMcG0012535
h1SMcG0016091
h1SMnG0013218
h1SMnG0014331

### Module s11

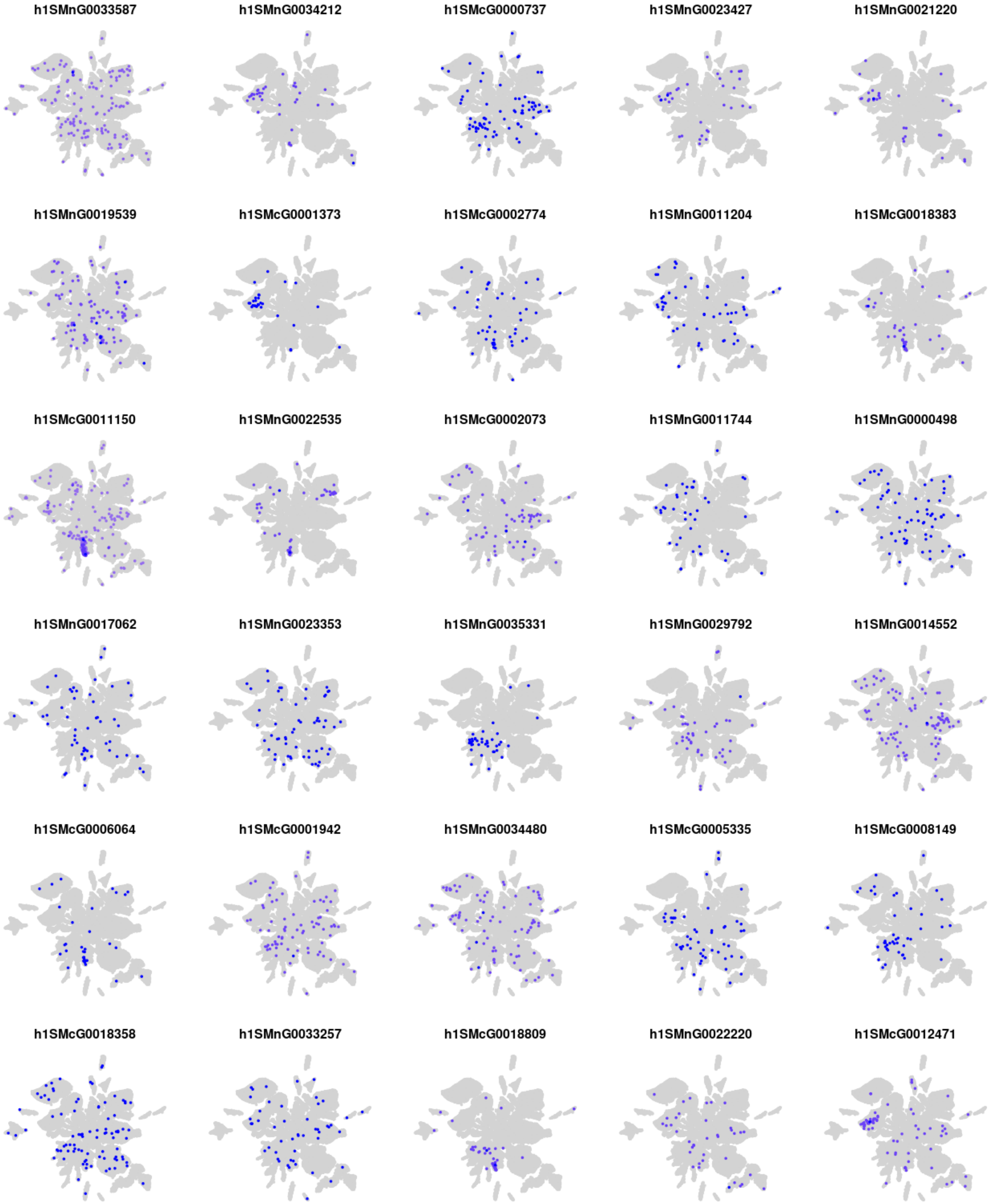
h1SMnG0033587
h1SMnG0034212
h1SMcG0000737
h1SMnG0023427
h1SMnG0021220
h1SMnG0019539
h1SMcG0001373
h1SMcG0002774
h1SMnG0011204
h1SMcG0018383
h1SMcG0011150
h1SMnG0022535
h1SMcG0002073
h1SMnG0011744
h1SMnG0000498
h1SMnG0017062
h1SMnG0023353
h1SMnG0035331
h1SMnG0029792
h1SMnG0014552
h1SMcG0006064
h1SMcG0001942
h1SMnG0034480
h1SMcG0005335
h1SMcG0008149
h1SMcG0018358
h1SMnG0033257
h1SMcG0018809
h1SMnG0022220
h1SMcG0012471

### Module s12

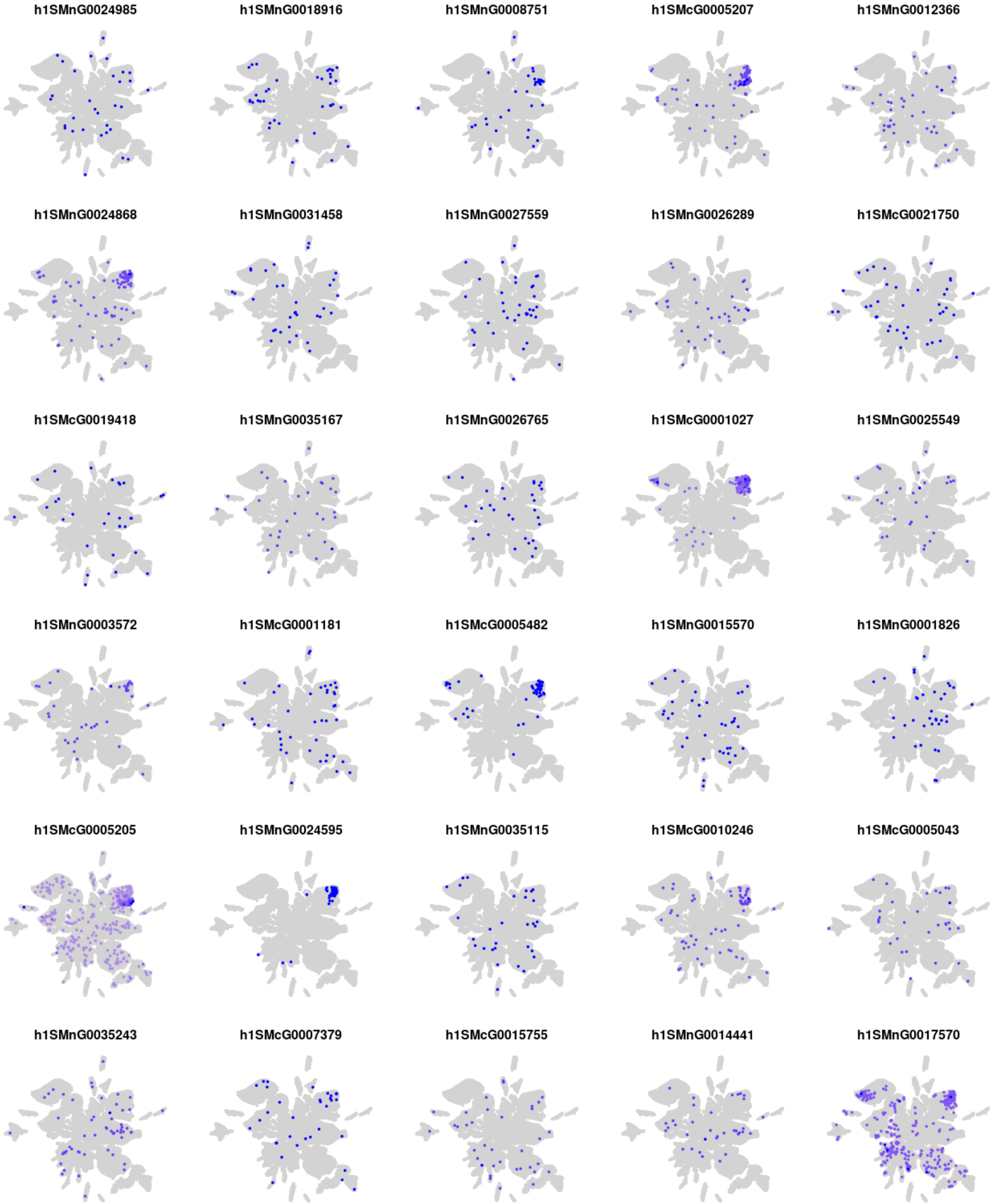
h1SMnG0024985
h1SMnG0018916
h1SMnG0008751
h1SMcG0005207
h1SMnG0012366
h1SMnG0024868
h1SMnG0031458
h1SMnG0027559
h1SMnG0026289
h1SMcG0021750
h1SMcG0019418
h1SMnG0035167
h1SMnG0026765
h1SMcG0001027
h1SMnG0025549
h1SMnG0003572
h1SMcG0001181
h1SMcG0005482
h1SMnG0015570
h1SMnG0001826
h1SMcG0005205
h1SMnG0024595
h1SMnG0035115
h1SMcG0010246
h1SMcG0005043
h1SMnG0035243
h1SMcG0007379
h1SMcG0015755
h1SMnG0014441
h1SMnG0017570

### Module s13

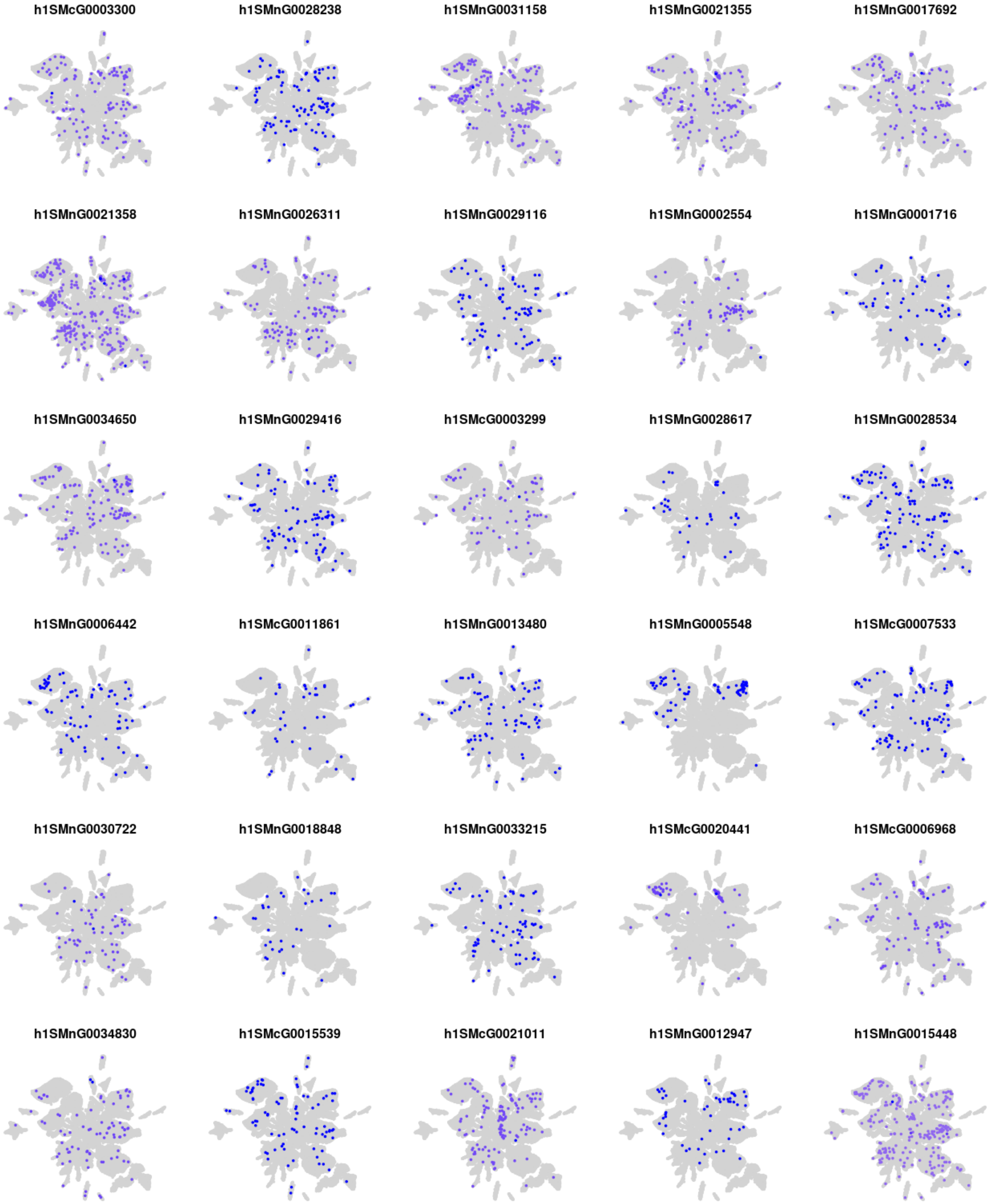
h1SMcG0003300
h1SMnG0028238
h1SMnG0031158
h1SMnG0021355
h1SMnG0017692
h1SMnG0021358
h1SMnG0026311
h1SMnG0029116
h1SMnG0002554
h1SMnG0001716
h1SMnG0034650
h1SMnG0029416
h1SMcG0003299
h1SMnG0028617
h1SMnG0028534
h1SMnG0006442
h1SMcG0011861
h1SMnG0013480
h1SMnG0005548
h1SMcG0007533
h1SMnG0030722
h1SMnG0018848
h1SMnG0033215
h1SMcG0020441
h1SMcG0006968
h1SMnG0034830
h1SMcG0015539
h1SMcG0021011
h1SMnG0012947
h1SMnG0015448

### Module s14

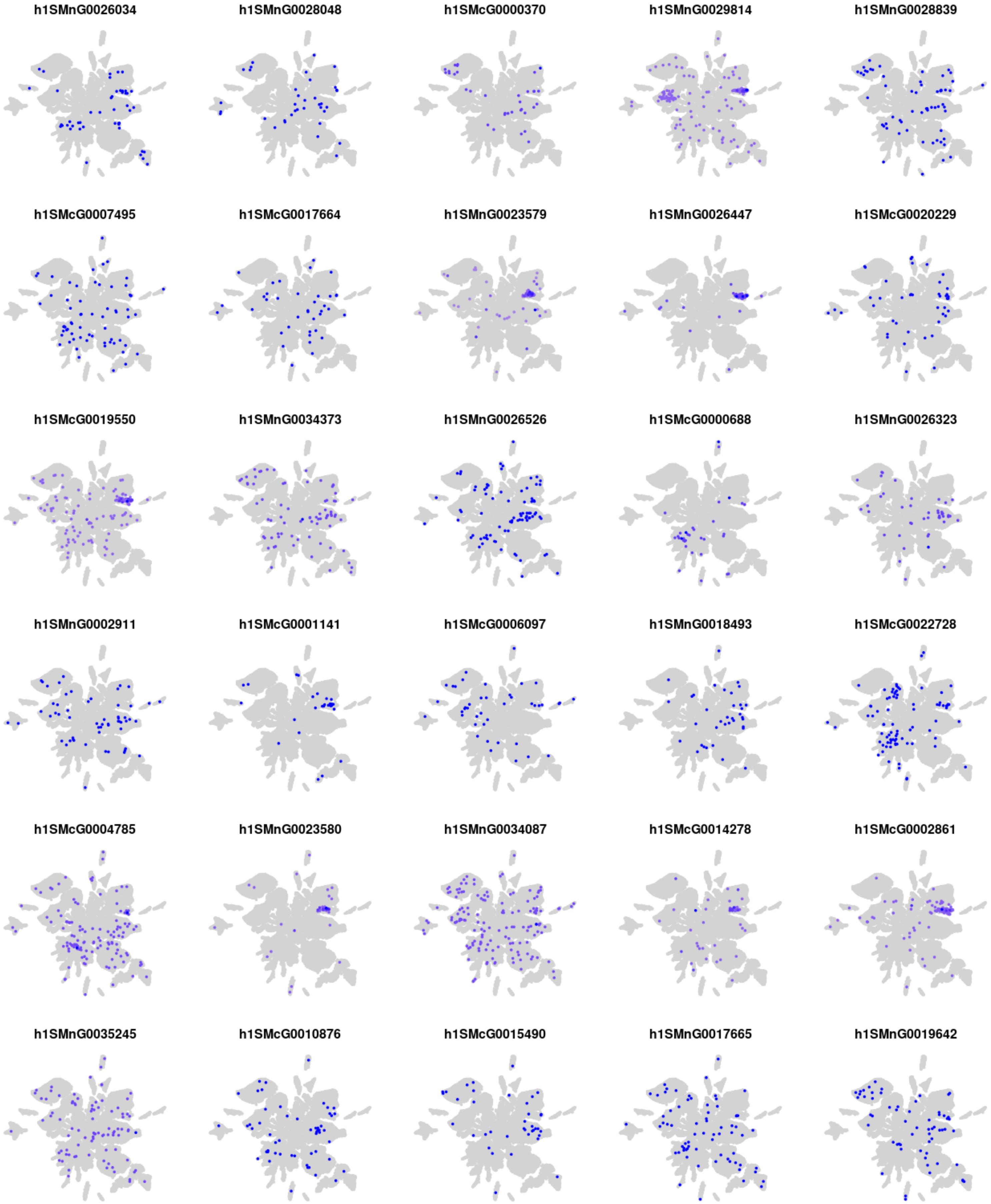
h1SMnG0026034
h1SMnG0028048
h1SMcG0000370
h1SMnG0029814
h1SMnG0028839
h1SMcG0007495
h1SMcG0017664
h1SMnG0023579
h1SMnG0026447
h1SMcG0020229
h1SMcG0019550
h1SMnG0034373
h1SMnG0026526
h1SMcG0000688
h1SMnG0026323
h1SMnG0002911
h1SMcG0001141
h1SMcG0006097
h1SMnG0018493
h1SMcG0022728
h1SMcG0004785
h1SMnG0023580
h1SMnG0034087
h1SMcG0014278
h1SMcG0002861
h1SMnG0035245
h1SMcG0010876
h1SMcG0015490
h1SMnG0017665
h1SMnG0019642

### Module s15

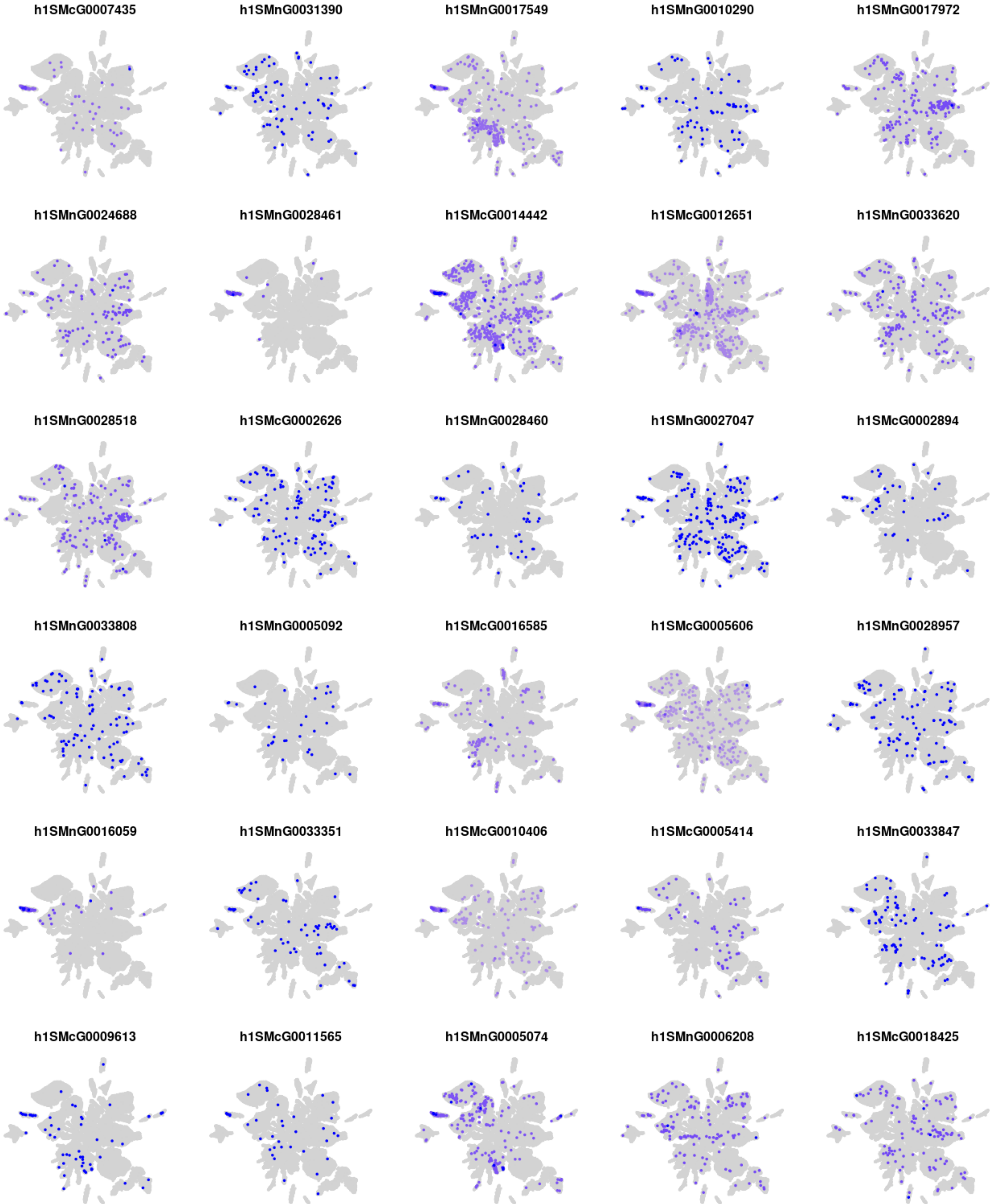
h1SMcG0007435
h1SMnG0031390
h1SMnG0017549
h1SMnG0010290
h1SMnG0017972
h1SMnG0024688
h1SMnG0028461
h1SMcG0014442
h1SMcG0012651
h1SMnG0033620
h1SMnG0028518
h1SMcG0002626
h1SMnG0028460
h1SMnG0027047
h1SMcG0002894
h1SMnG0033808
h1SMnG0005092
h1SMcG0016585
h1SMcG0005606
h1SMnG0028957
h1SMnG0016059
h1SMnG0033351
h1SMcG0010406
h1SMcG0005414
h1SMnG0033847
h1SMcG0009613
h1SMcG0011565
h1SMnG0005074
h1SMnG0006208
h1SMcG0018425

### Module s16

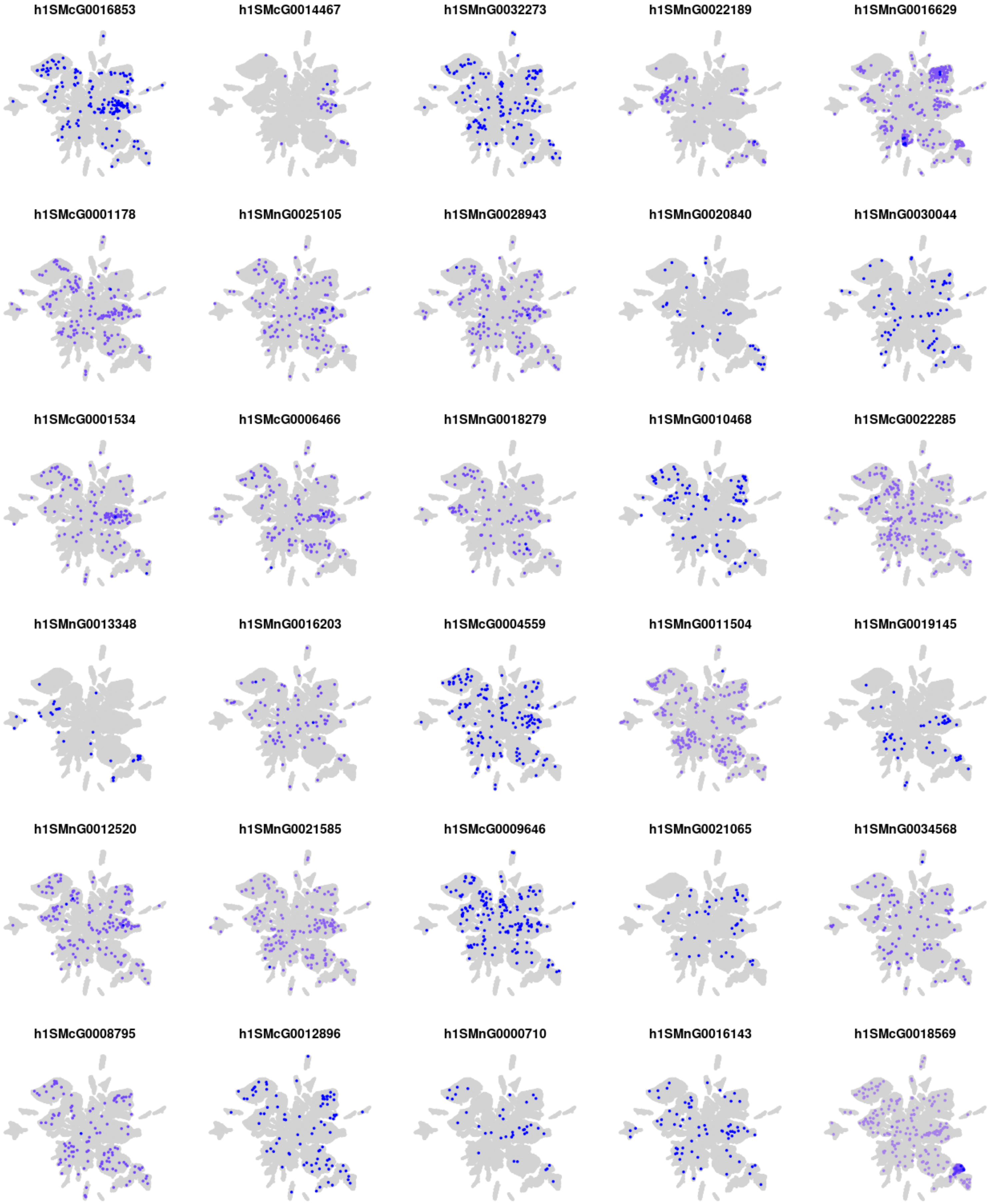
h1SMcG0016853
h1SMcG0014467
h1SMnG0032273
h1SMnG0022189
h1SMnG0016629
h1SMcG0001178
h1SMnG0025105
h1SMnG0028943
h1SMnG0020840
h1SMnG0030044
h1SMcG0001534
h1SMcG0006466
h1SMnG0018279
h1SMnG0010468
h1SMcG0022285
h1SMnG0013348
h1SMnG0016203
h1SMcG0004559
h1SMnG0011504
h1SMnG0019145
h1SMnG0012520
h1SMnG0021585
h1SMcG0009646
h1SMnG0021065
h1SMnG0034568
h1SMcG0008795
h1SMcG0012896
h1SMnG0000710
h1SMnG0016143
h1SMcG0018569

### Module s17

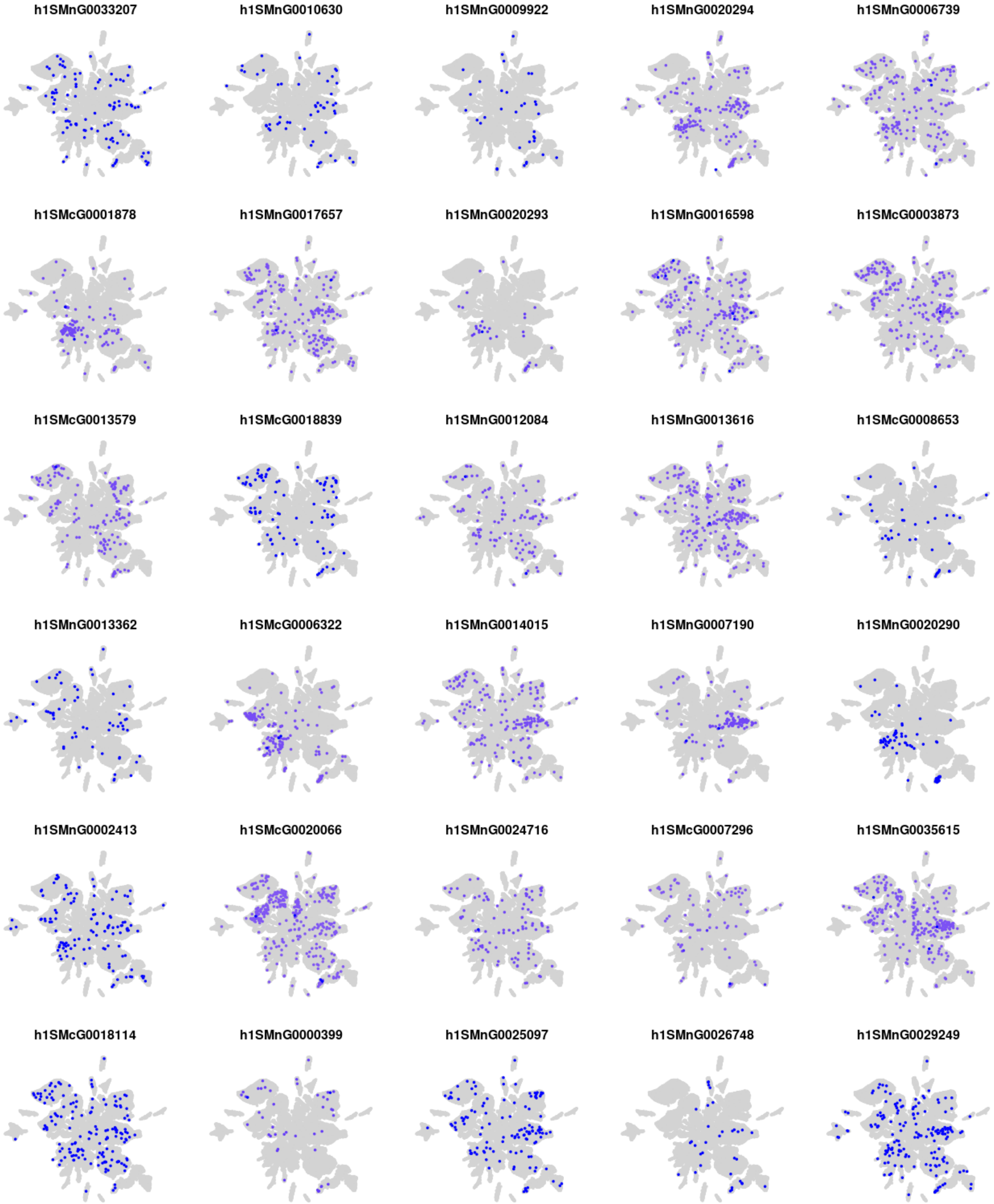
h1SMnG0033207
h1SMnG0010630
h1SMnG0009922
h1SMnG0020294
h1SMnG0006739
h1SMcG0001878
h1SMnG0017657
h1SMnG0020293
h1SMnG0016598
h1SMcG0003873
h1SMcG0013579
h1SMcG0018839
h1SMnG0012084
h1SMnG0013616
h1SMcG0008653
h1SMnG0013362
h1SMcG0006322
h1SMnG0014015
h1SMnG0007190
h1SMnG0020290
h1SMnG0002413
h1SMcG0020066
h1SMnG0024716
h1SMcG0007296
h1SMnG0035615
h1SMcG0018114
h1SMnG0000399
h1SMnG0025097
h1SMnG0026748
h1SMnG0029249

### Module s18

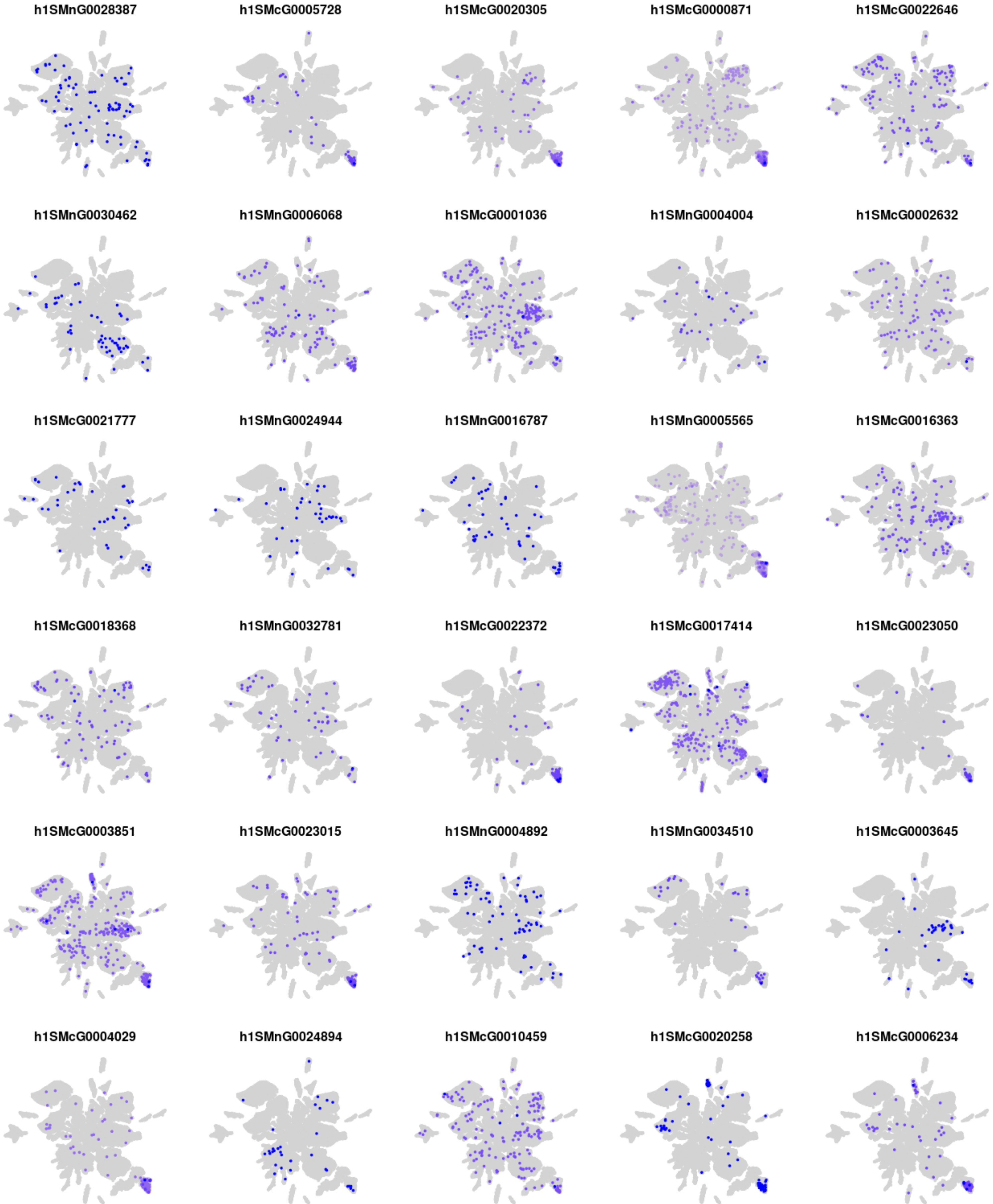
h1SMnG0028387
h1SMcG0005728
h1SMcG0020305
h1SMcG0000871
h1SMcG0022646
h1SMnG0030462
h1SMnG0006068
h1SMcG0001036
h1SMnG0004004
h1SMcG0002632
h1SMcG0021777
h1SMnG0024944
h1SMnG0016787
h1SMnG0005565
h1SMcG0016363
h1SMcG0018368
h1SMnG0032781
h1SMcG0022372
h1SMcG0017414
h1SMcG0023050
h1SMcG0003851
h1SMcG0023015
h1SMnG0004892
h1SMnG0034510
h1SMcG0003645
h1SMcG0004029
h1SMnG0024894
h1SMcG0010459
h1SMcG0020258
h1SMcG0006234

### Module s19

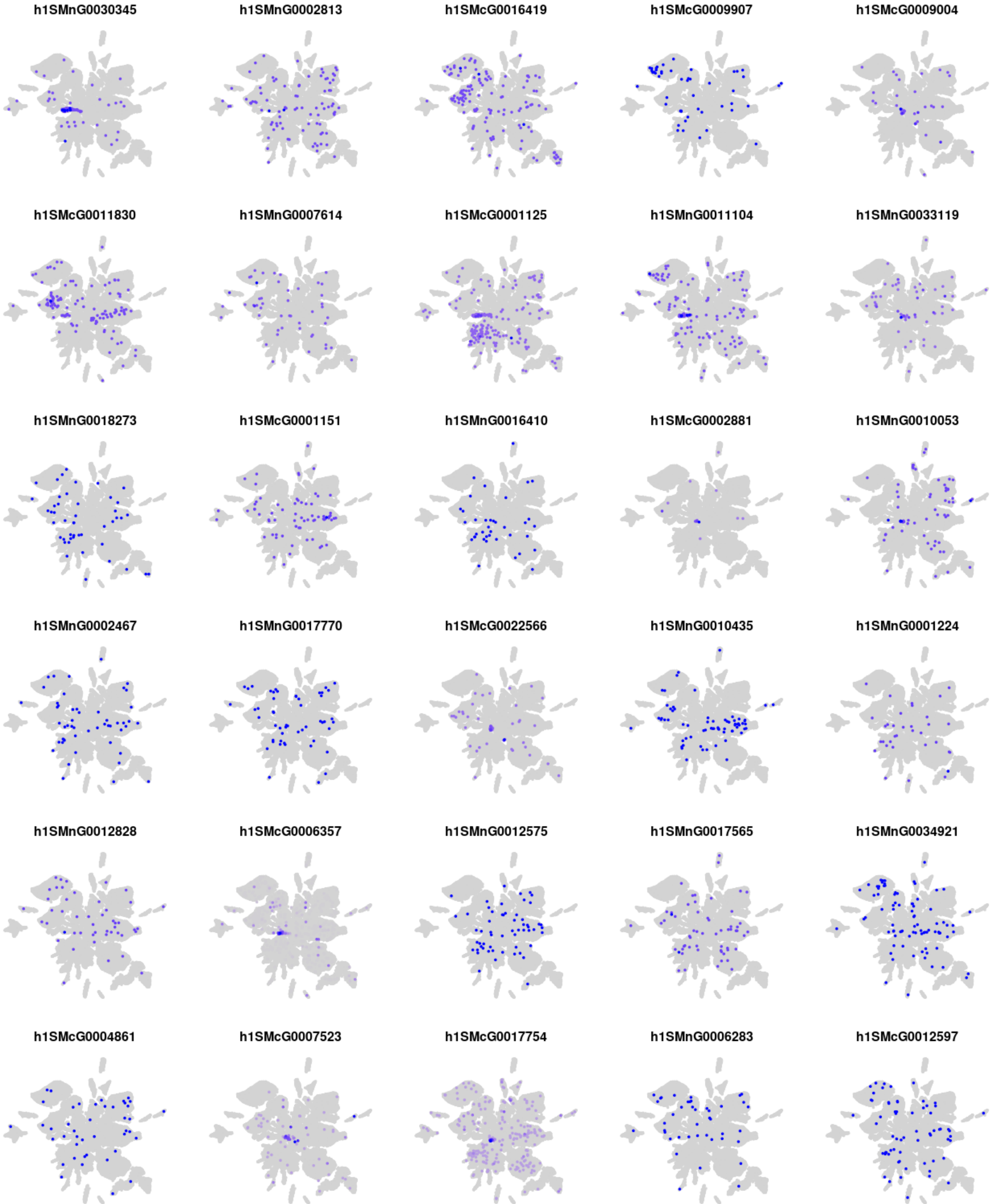
h1SMnG0030345
h1SMnG0002813
h1SMcG0016419
h1SMcG0009907
h1SMcG0009004
h1SMcG0011830
h1SMnG0007614
h1SMcG0001125
h1SMnG0011104
h1SMnG0033119
h1SMnG0018273
h1SMcG0001151
h1SMnG0016410
h1SMcG0002881
h1SMnG0010053
h1SMnG0002467
h1SMnG0017770
h1SMcG0022566
h1SMnG0010435
h1SMnG0001224
h1SMnG0012828
h1SMcG0006357
h1SMnG0012575
h1SMnG0017565
h1SMnG0034921
h1SMcG0004861
h1SMcG0007523
h1SMcG0017754
h1SMnG0006283
h1SMcG0012597

### Module s20

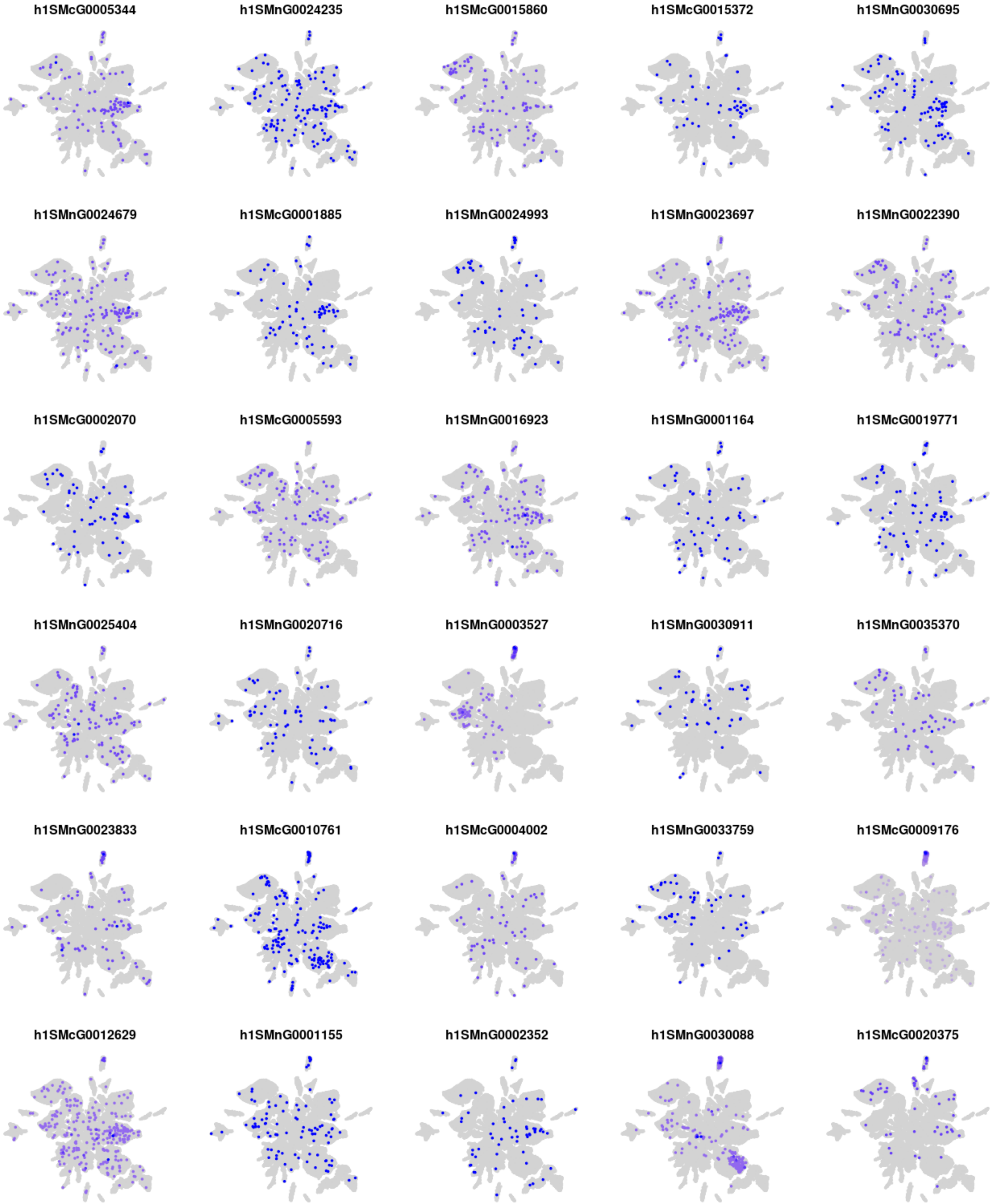
h1SMcG0005344
h1SMnG0024235
h1SMcG0015860
h1SMcG0015372
h1SMnG0030695
h1SMnG0024679
h1SMcG0001885
h1SMnG0024993
h1SMnG0023697
h1SMnG0022390
h1SMcG0002070
h1SMcG0005593
h1SMnG0016923
h1SMnG0001164
h1SMcG0019771
h1SMnG0025404
h1SMnG0020716
h1SMnG0003527
h1SMnG0030911
h1SMnG0035370
h1SMnG0023833
h1SMcG0010761
h1SMcG0004002
h1SMnG0033759
h1SMcG0009176
h1SMcG0012629
h1SMnG0001155
h1SMnG0002352
h1SMnG0030088
h1SMcG0020375

### Module s21

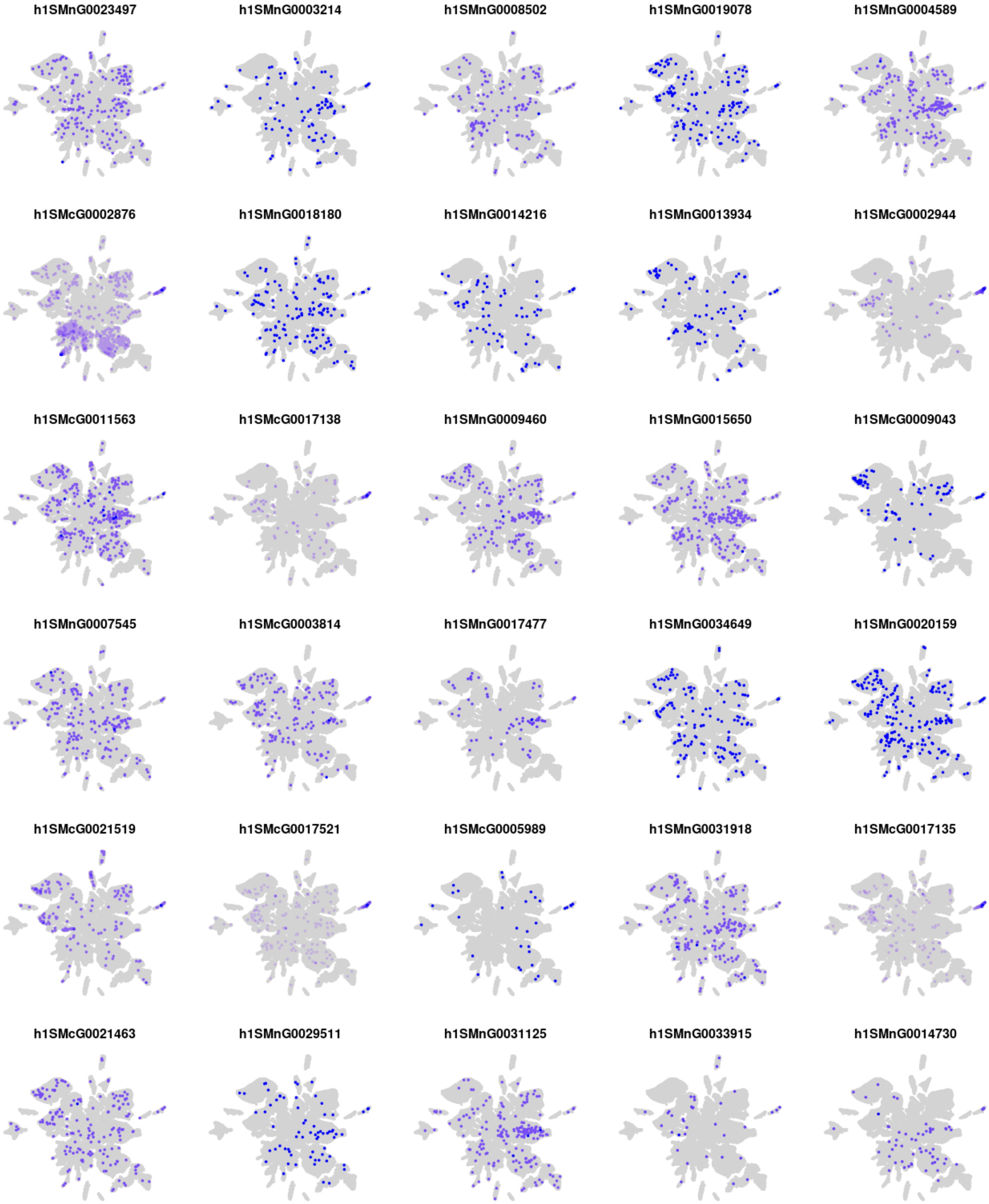
h1SMnG0023497
h1SMnG0003214
h1SMnG0008502
h1SMnG0019078
h1SMnG0004589
h1SMcG0002876
h1SMnG0018180
h1SMnG0014216
h1SMnG0013934
h1SMcG0002944
h1SMcG0011563
h1SMcG0017138
h1SMnG0009460
h1SMnG0015650
h1SMcG0009043
h1SMnG0007545
h1SMcG0003814
h1SMnG0017477
h1SMnG0034649
h1SMnG0020159
h1SMcG0021519
h1SMcG0017521
h1SMcG0005989
h1SMnG0031918
h1SMcG0017135
h1SMcG0021463
h1SMnG0029511
h1SMnG0031125
h1SMnG0033915
h1SMnG0014730

### Module s22

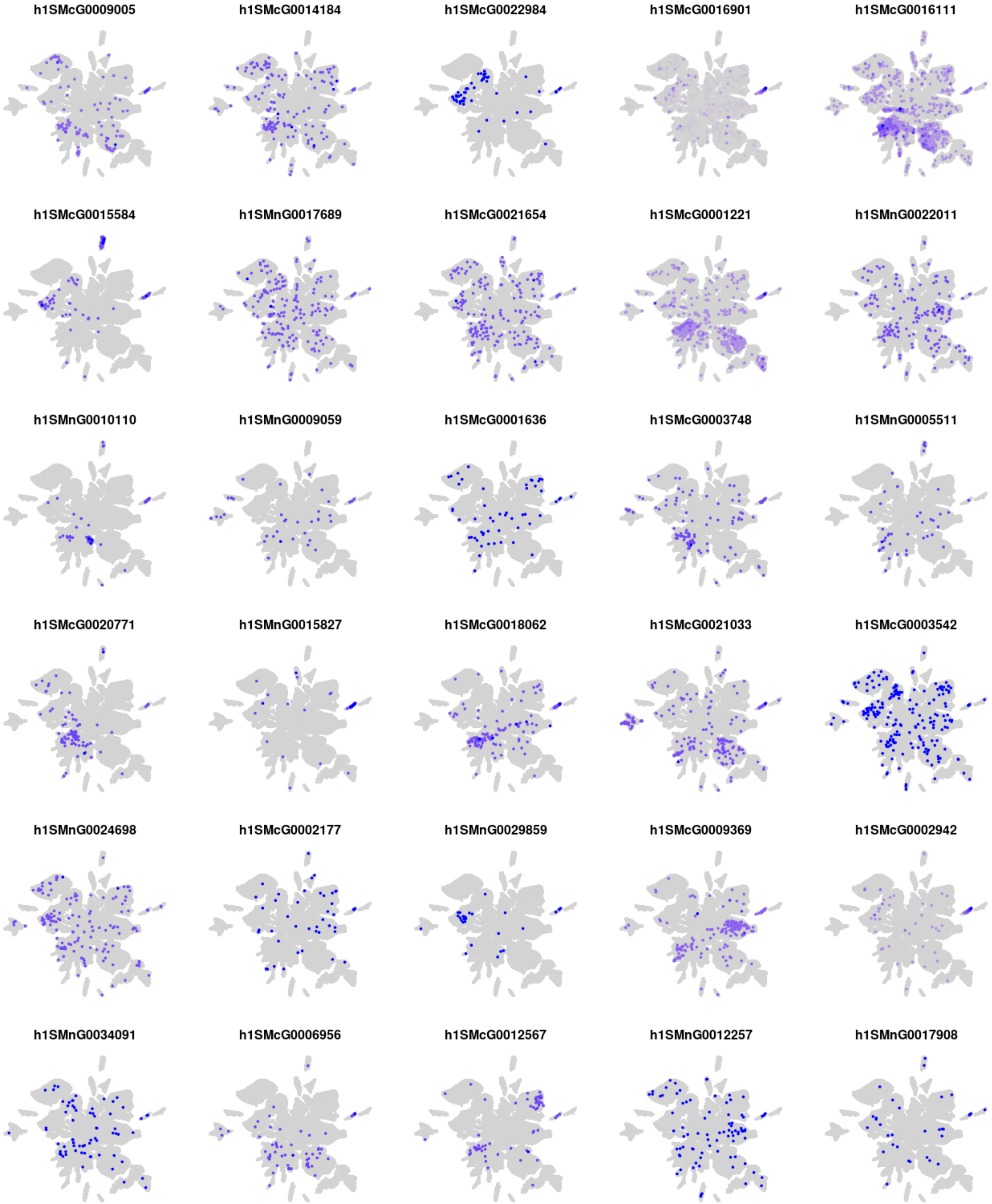
h1SMcG0009005
h1SMcG0014184
h1SMcG0022984
h1SMcG0016901
h1SMcG0016111
h1SMcG0015584
h1SMnG0017689
h1SMcG0021654
h1SMcG0001221
h1SMnG0022011
h1SMnG0010110
h1SMnG0009059
h1SMcG0001636
h1SMcG0003748
h1SMnG0005511
h1SMcG0020771
h1SMnG0015827
h1SMcG0018062
h1SMcG0021033
h1SMcG0003542
h1SMnG0024698
h1SMcG0002177
h1SMnG0029859
h1SMcG0009369
h1SMcG0002942
h1SMnG0034091
h1SMcG0006956
h1SMcG0012567
h1SMnG0012257
h1SMnG0017908

### Module s23

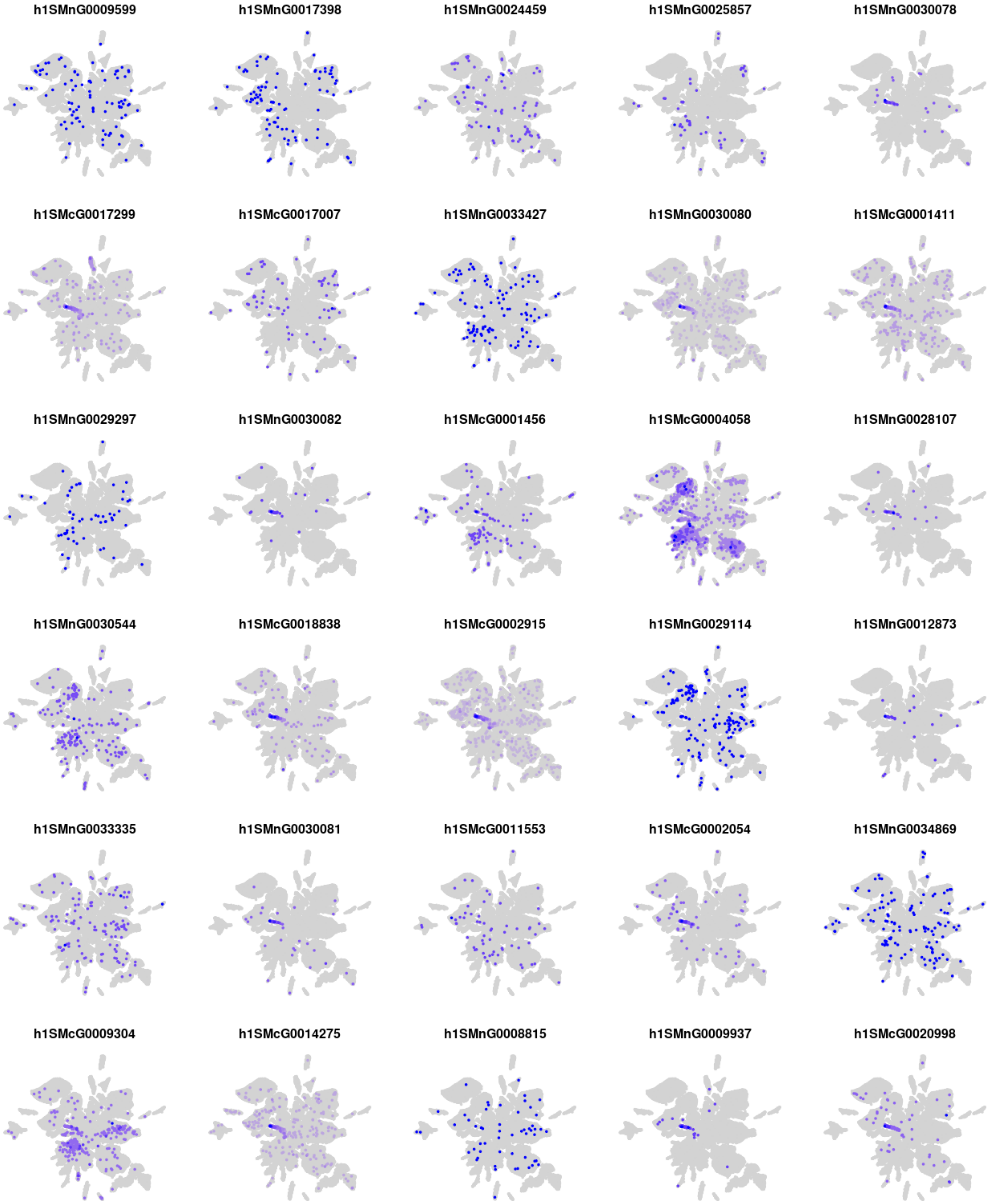

h1SMnG0009599
h1SMnG0017398
h1SMnG0024459
h1SMnG0025857
h1SMnG0030078
h1SMcG0017299
h1SMcG0017007
h1SMnG0033427
h1SMnG0030080
h1SMcG0001411
h1SMnG0029297
h1SMnG0030082
h1SMcG0001456
h1SMcG0004058
h1SMnG0028107
h1SMnG0030544
h1SMcG0018838
h1SMcG0002915
h1SMnG0029114
h1SMnG0012873
h1SMnG0033335
h1SMnG0030081
h1SMcG0011553
h1SMcG0002054
h1SMnG0034869
h1SMcG0009304
h1SMcG0014275
h1SMnG0008815
h1SMnG0009937
h1SMcG0020998

### Module s24

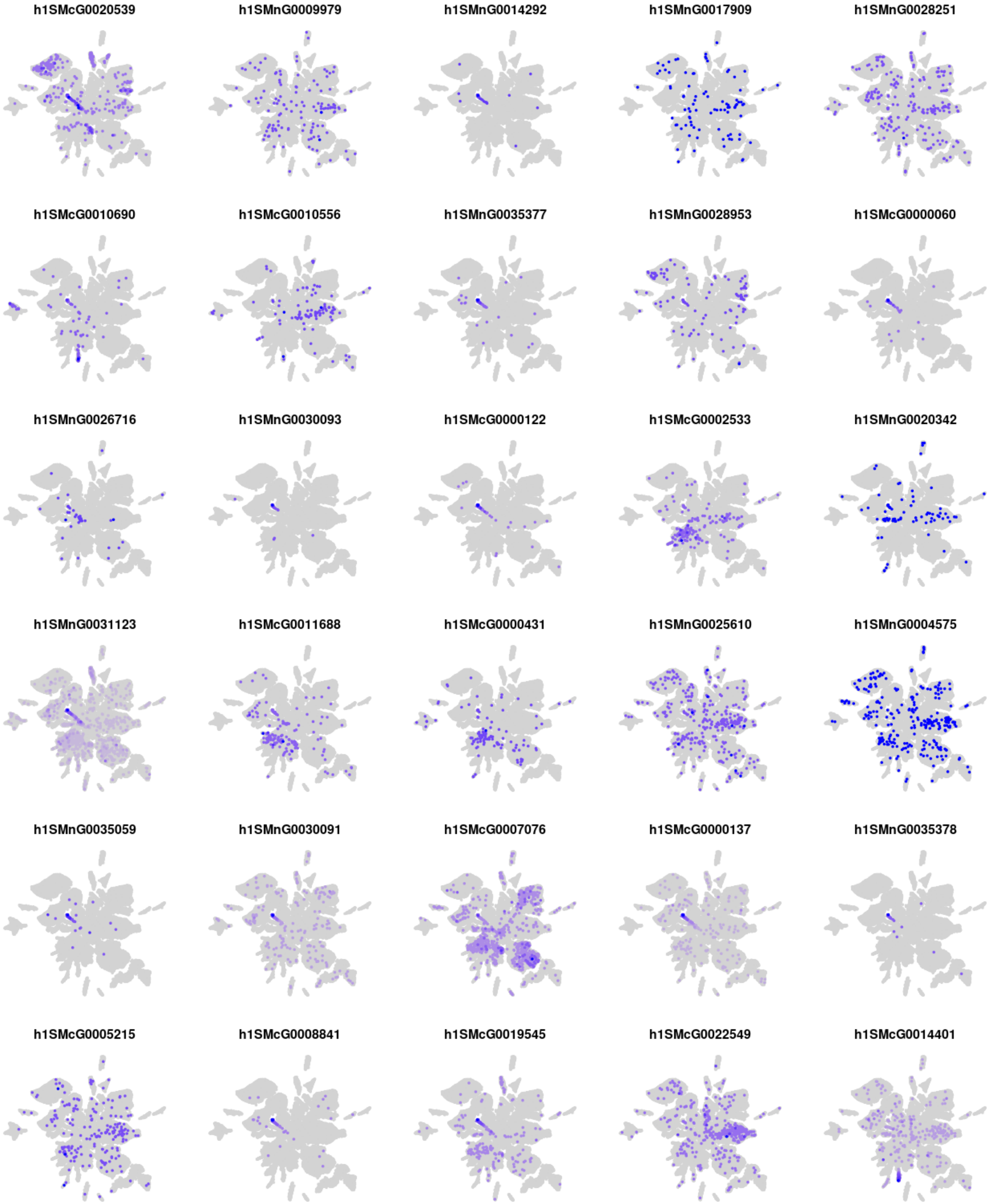
h1SMcG0020539
h1SMnG0009979
h1SMnG0014292
h1SMnG0017909
h1SMnG0028251
h1SMcG0010690
h1SMcG0010556
h1SMnG0035377
h1SMnG0028953
h1SMcG0000060
h1SMnG0026716
h1SMnG0030093
h1SMcG0000122
h1SMcG0002533
h1SMnG0020342
h1SMnG0031123
h1SMcG0011688
h1SMcG0000431
h1SMnG0025610
h1SMnG0004575
h1SMnG0035059
h1SMnG0030091
h1SMcG0007076
h1SMcG0000137
h1SMnG0035378
h1SMcG0005215
h1SMcG0008841
h1SMcG0019545
h1SMcG0022549
h1SMcG0014401

#### Module m01

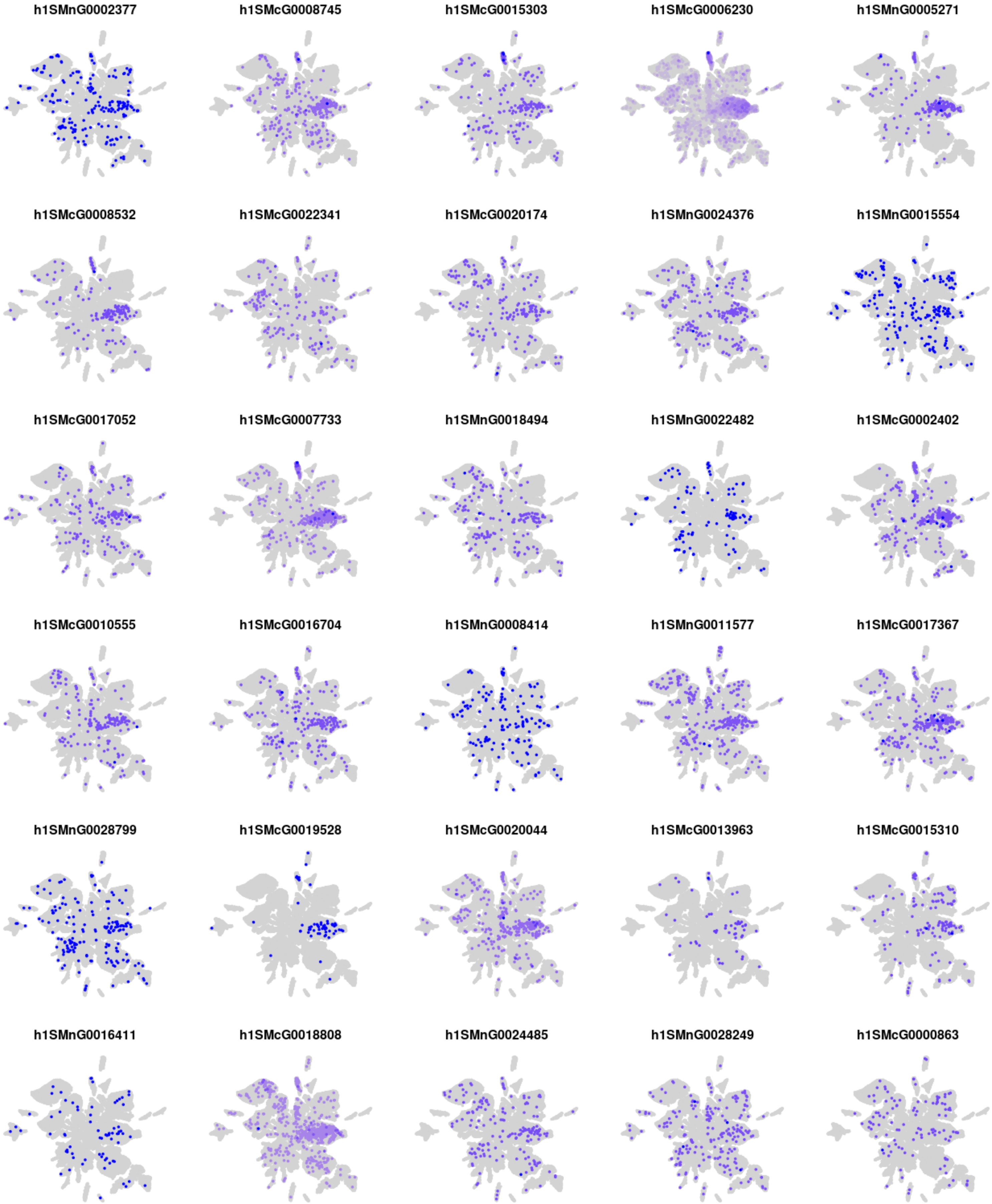
h1SMnG0002377
h1SMcG0008745
h1SMcG0015303
h1SMcG0006230
h1SMnG0005271
h1SMcG0008532
h1SMcG0022341
h1SMcG0020174
h1SMnG0024376
h1SMnG0015554
h1SMcG0017052
h1SMcG0007733
h1SMnG0018494
h1SMnG0022482
h1SMcG0002402
h1SMcG0010555
h1SMcG0016704
h1SMnG0008414
h1SMnG0011577
h1SMcG0017367
h1SMnG0028799
h1SMcG0019528
h1SMcG0020044
h1SMcG0013963
h1SMcG0015310
h1SMnG0016411
h1SMcG0018808
h1SMnG0024485
h1SMnG0028249
h1SMcG0000863

### Module m02

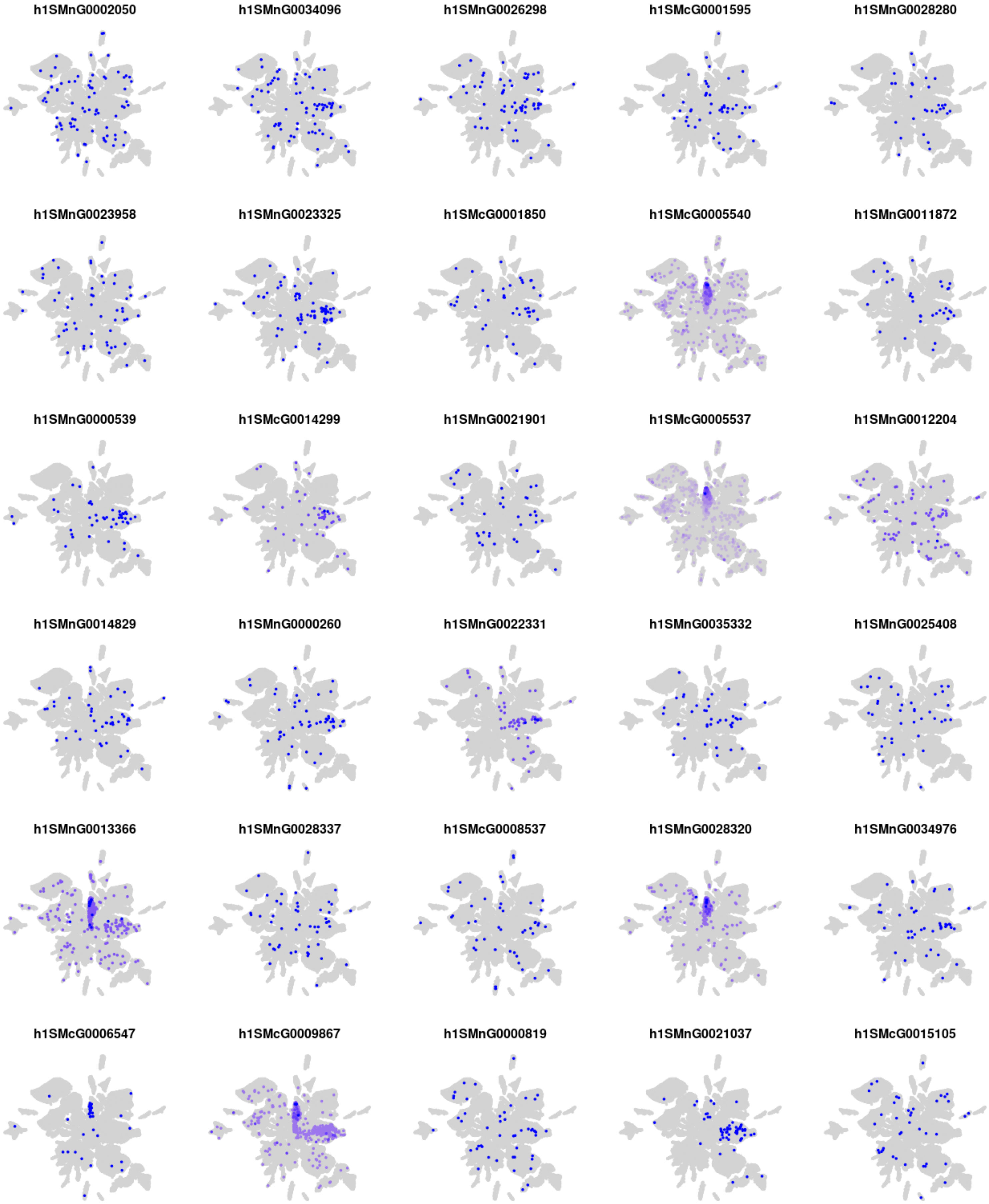
h1SMnG0002050
h1SMnG0034096
h1SMnG0026298
h1SMcG0001595
h1SMnG0028280
h1SMnG0023958
h1SMnG0023325
h1SMcG0001850
h1SMcG0005540
h1SMnG0011872
h1SMnG0000539
h1SMcG0014299
h1SMnG0021901
h1SMcG0005537
h1SMnG0012204
h1SMnG0014829
h1SMnG0000260
h1SMnG0022331
h1SMnG0035332
h1SMnG0025408
h1SMnG0013366
h1SMnG0028337
h1SMcG0008537
h1SMnG0028320
h1SMnG0034976
h1SMcG0006547
h1SMcG0009867
h1SMnG0000819
h1SMnG0021037
h1SMcG0015105

### Module m03

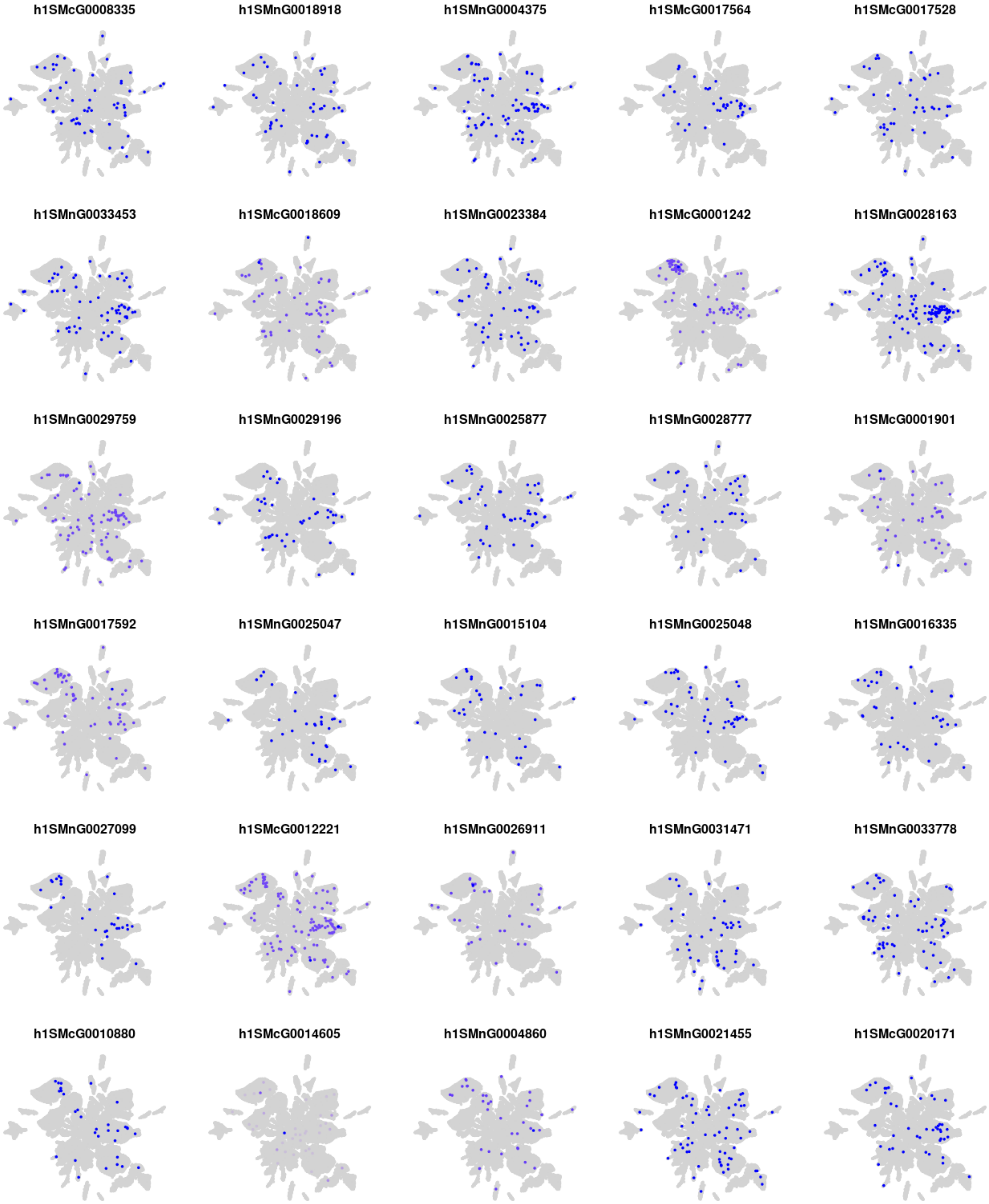
h1SMcG0008335
h1SMnG0018918
h1SMnG0004375
h1SMcG0017564
h1SMcG0017528
h1SMnG0033453
h1SMcG0018609
h1SMnG0023384
h1SMcG0001242
h1SMnG0028163
h1SMnG0029759
h1SMnG0029196
h1SMnG0025877
h1SMnG0028777
h1SMcG0001901
h1SMnG0017592
h1SMnG0025047
h1SMnG0015104
h1SMnG0025048
h1SMnG0016335
h1SMnG0027099
h1SMcG0012221
h1SMnG0026911
h1SMnG0031471
h1SMnG0033778
h1SMcG0010880
h1SMcG0014605
h1SMnG0004860
h1SMnG0021455
h1SMcG0020171

#### Module m04

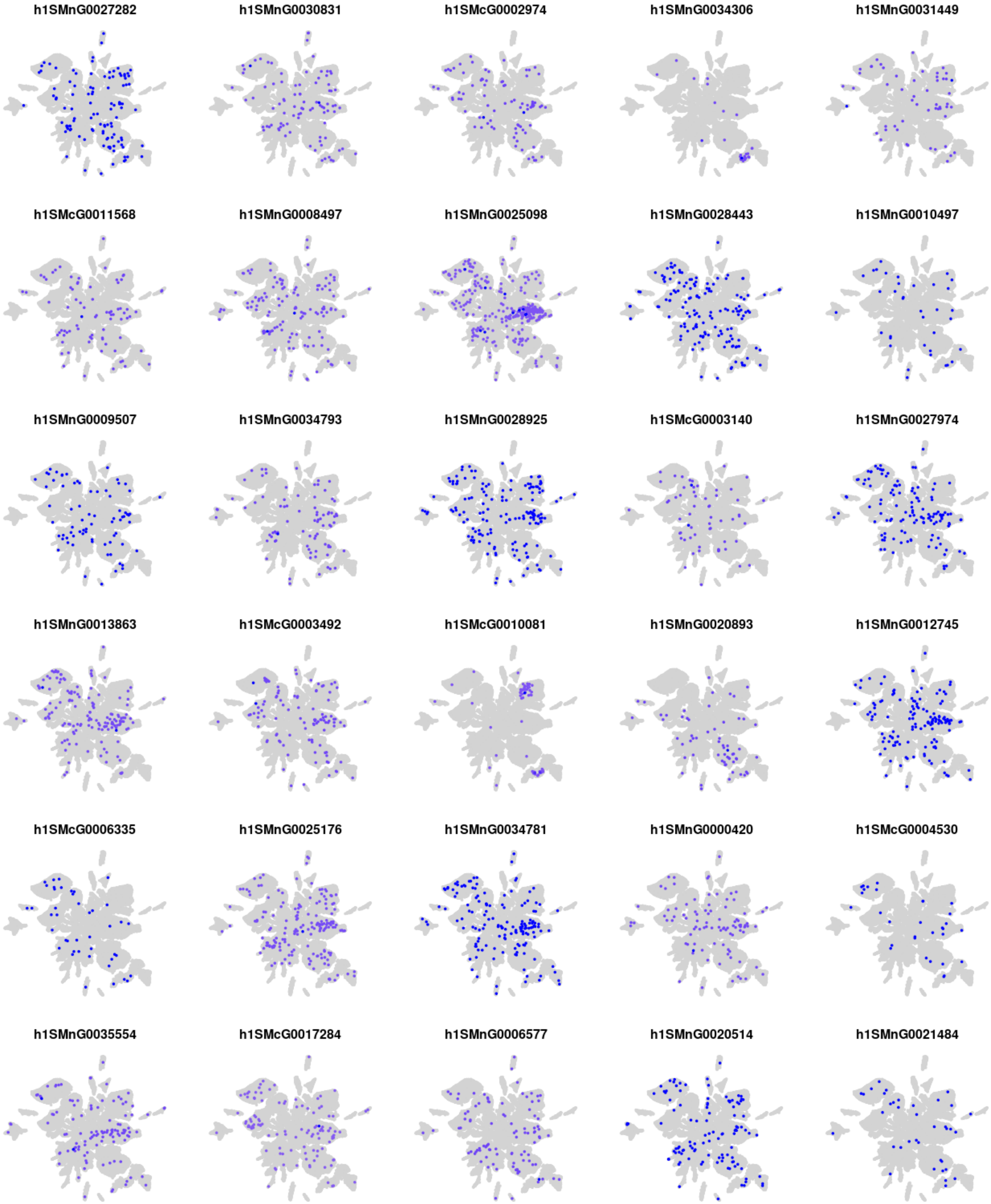
h1SMnG0027282
h1SMnG0030831
h1SMcG0002974
h1SMnG0034306
h1SMnG0031449
h1SMcG0011568
h1SMnG0008497
h1SMnG0025098
h1SMnG0028443
h1SMnG0010497
h1SMnG0009507
h1SMnG0034793
h1SMnG0028925
h1SMcG0003140
h1SMnG0027974
h1SMnG0013863
h1SMcG0003492
h1SMcG0010081
h1SMnG0020893
h1SMnG0012745
h1SMcG0006335
h1SMnG0025176
h1SMnG0034781
h1SMnG0000420
h1SMcG0004530
h1SMnG0035554
h1SMcG0017284
h1SMnG0006577
h1SMnG0020514
h1SMnG0021484

#### Module m05

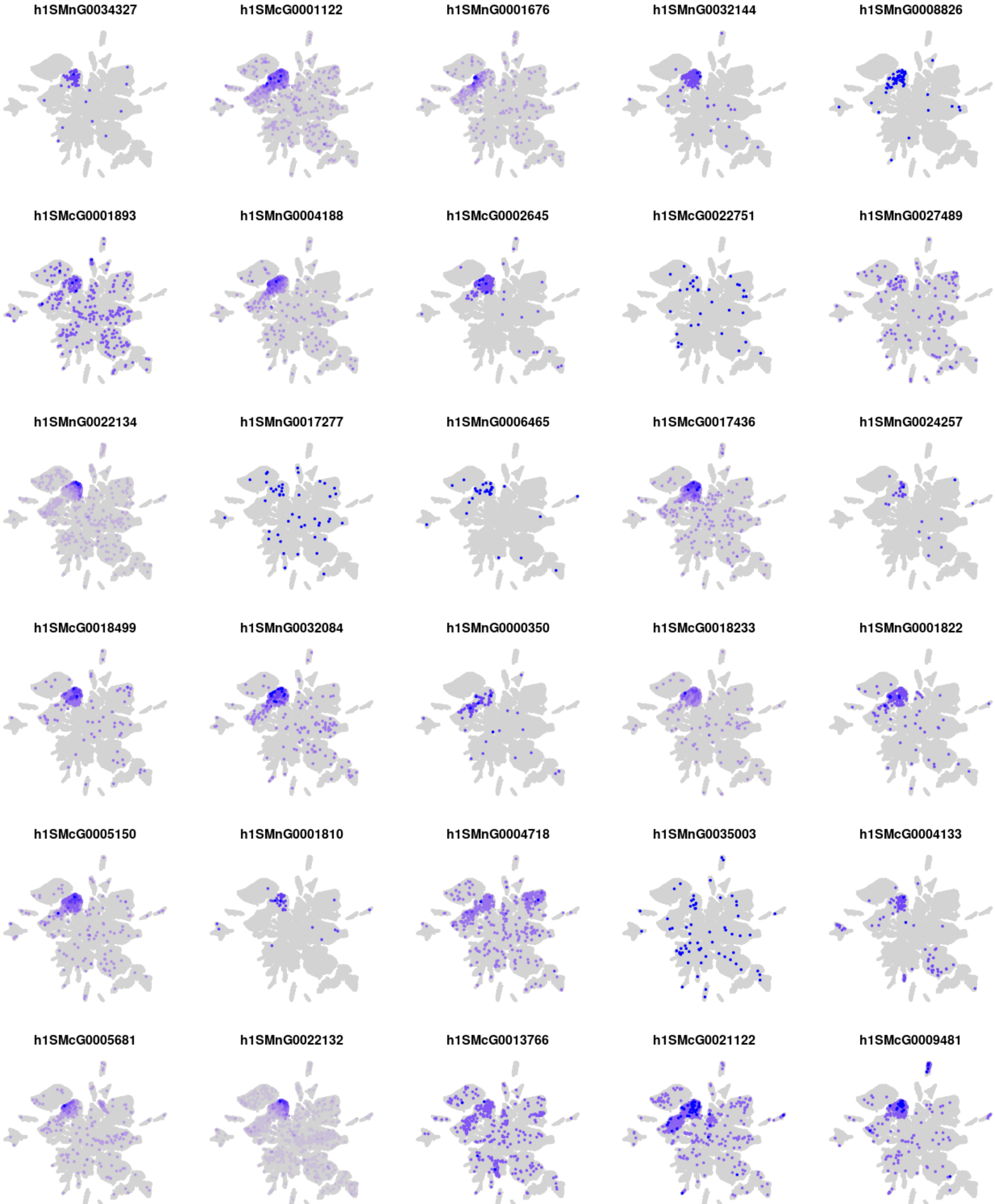
h1SMnG0034327
h1SMcG0001122
h1SMnG0001676
h1SMnG0032144
h1SMnG0008826
h1SMcG0001893
h1SMnG0004188
h1SMcG0002645
h1SMcG0022751
h1SMnG0027489
h1SMnG0022134
h1SMnG0017277
h1SMnG0006465
h1SMcG0017436
h1SMnG0024257
h1SMcG0018499
h1SMnG0032084
h1SMnG0000350
h1SMcG0018233
h1SMnG0001822
h1SMcG0005150
h1SMnG0001810
h1SMnG0004718
h1SMnG0035003
h1SMcG0004133
h1SMcG0005681
h1SMnG0022132
h1SMcG0013766
h1SMcG0021122
h1SMcG0009481

### Module m06

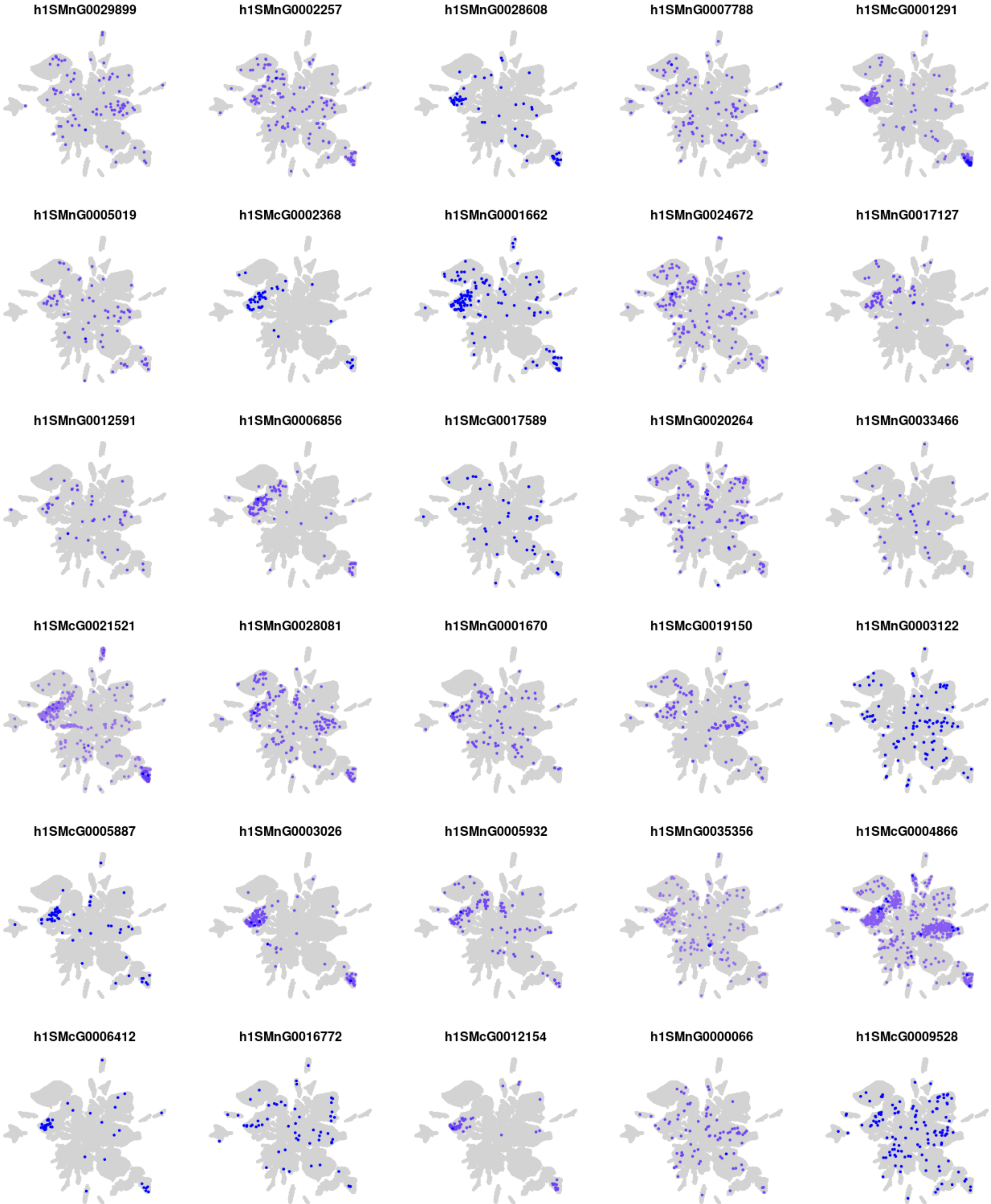
h1SMnG0029899
h1SMnG0002257
h1SMnG0028608
h1SMnG0007788
h1SMcG0001291
h1SMnG0005019
h1SMcG0002368
h1SMnG0001662
h1SMnG0024672
h1SMnG0017127
h1SMnG0012591
h1SMnG0006856
h1SMcG0017589
h1SMnG0020264
h1SMnG0033466
h1SMcG0021521
h1SMnG0028081
h1SMnG0001670
h1SMcG0019150
h1SMnG0003122
h1SMcG0005887
h1SMnG0003026
h1SMnG0005932
h1SMnG0035356
h1SMcG0004866
h1SMcG0006412
h1SMnG0016772
h1SMcG0012154
h1SMnG0000066
h1SMcG0009528

### Module m07

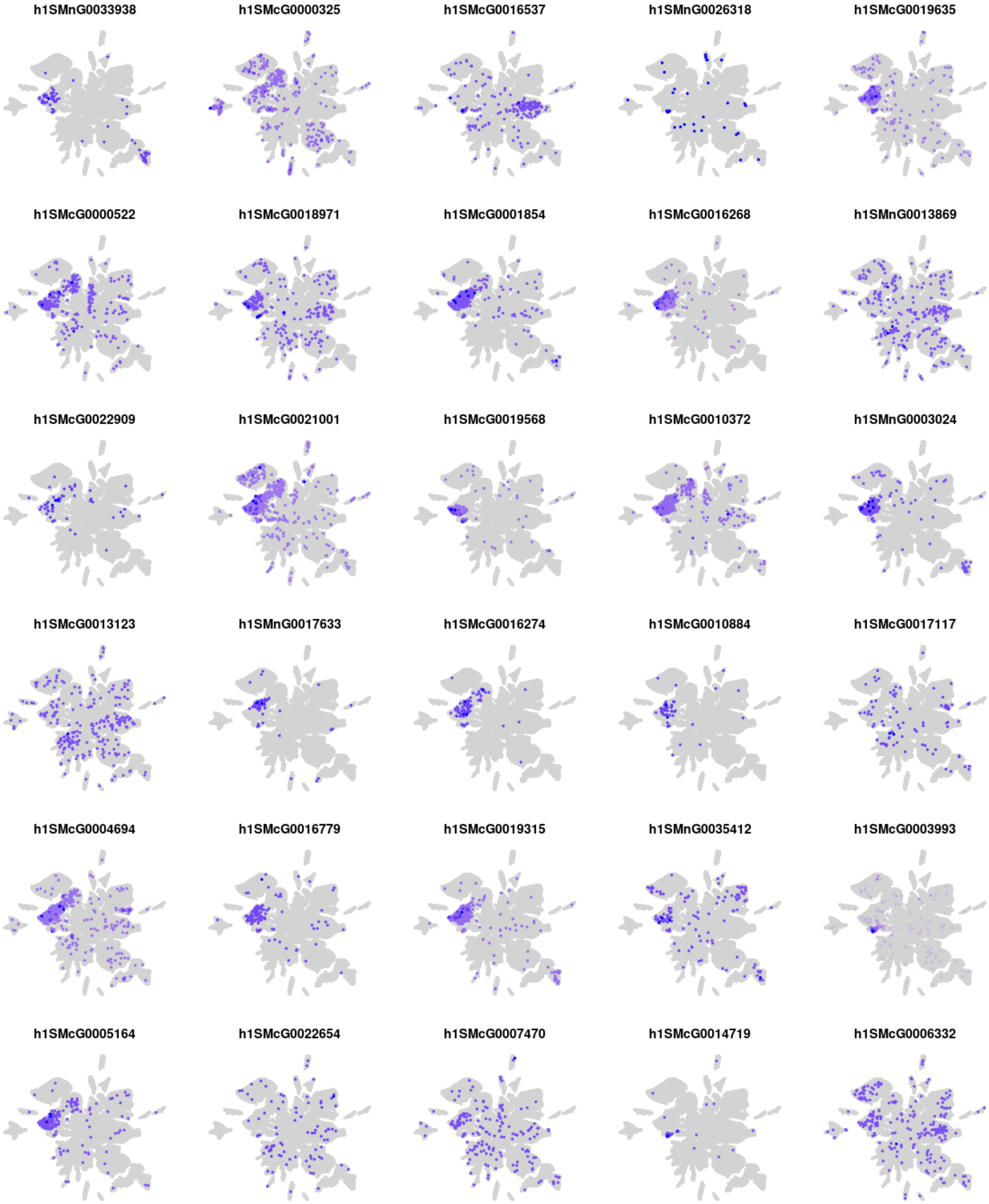
h1SMnG0033938
h1SMcG0000325
h1SMcG0016537
h1SMnG0026318
h1SMcG0019635
h1SMcG0000522
h1SMcG0018971
h1SMcG0001854
h1SMcG0016268
h1SMnG0013869
h1SMcG0022909
h1SMcG0021001
h1SMcG0019568
h1SMcG0010372
h1SMnG0003024
h1SMcG0013123
h1SMnG0017633
h1SMcG0016274
h1SMcG0010884
h1SMcG0017117
h1SMcG0004694
h1SMcG0016779
h1SMcG0019315
h1SMnG0035412
h1SMcG0003993
h1SMcG0005164
h1SMcG0022654
h1SMcG0007470
h1SMcG0014719
h1SMcG0006332

### Module m08

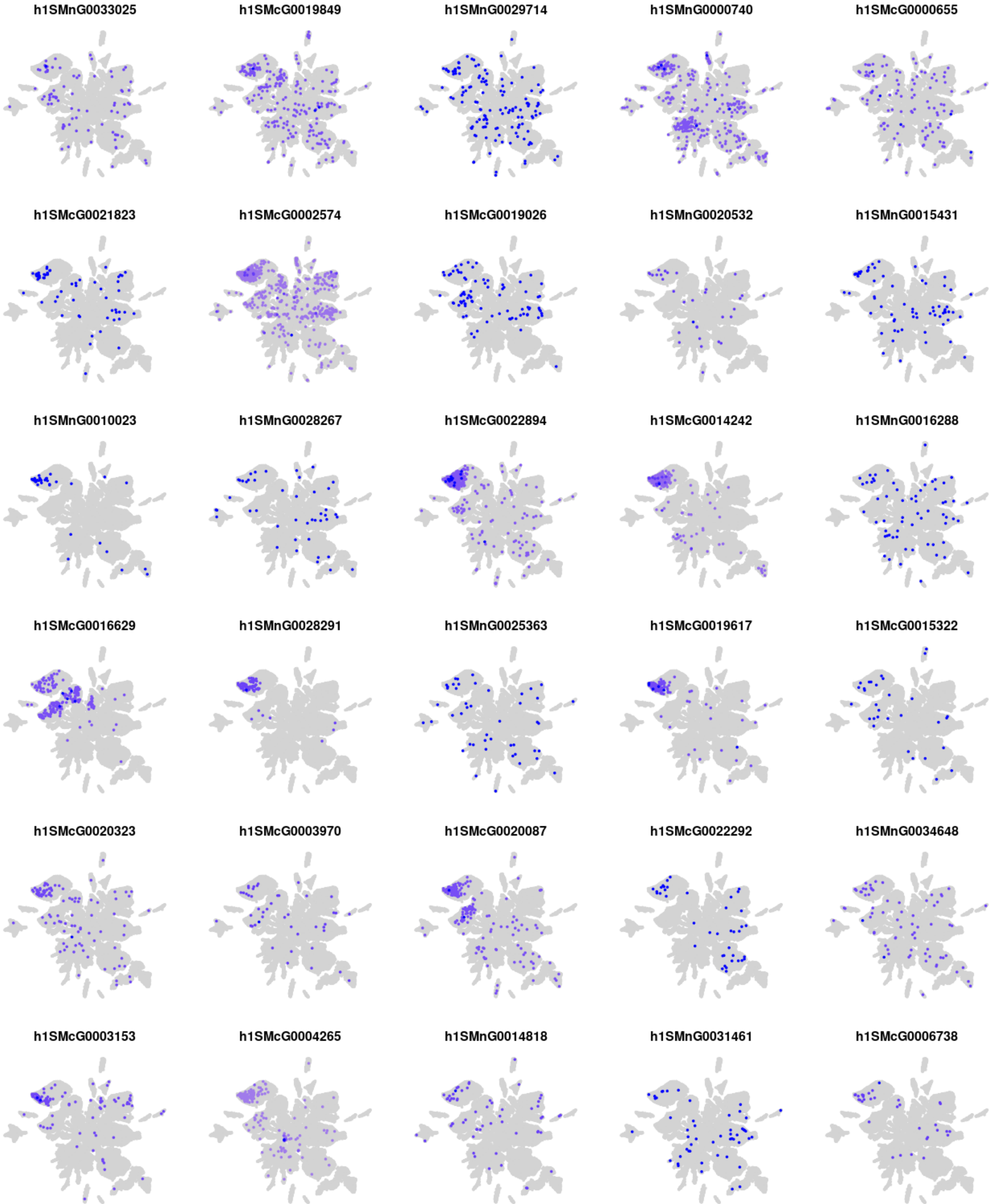
h1SMnG0033025
h1SMcG0019849
h1SMnG0029714
h1SMnG0000740
h1SMcG0000655
h1SMcG0021823
h1SMcG0002574
h1SMcG0019026
h1SMnG0020532
h1SMnG0015431
h1SMnG0010023
h1SMnG0028267
h1SMcG0022894
h1SMcG0014242
h1SMnG0016288
h1SMcG0016629
h1SMnG0028291
h1SMnG0025363
h1SMcG0019617
h1SMcG0015322
h1SMcG0020323
h1SMcG0003970
h1SMcG0020087
h1SMcG0022292
h1SMnG0034648
h1SMcG0003153
h1SMcG0004265
h1SMnG0014818
h1SMnG0031461
h1SMcG0006738

### Module m09

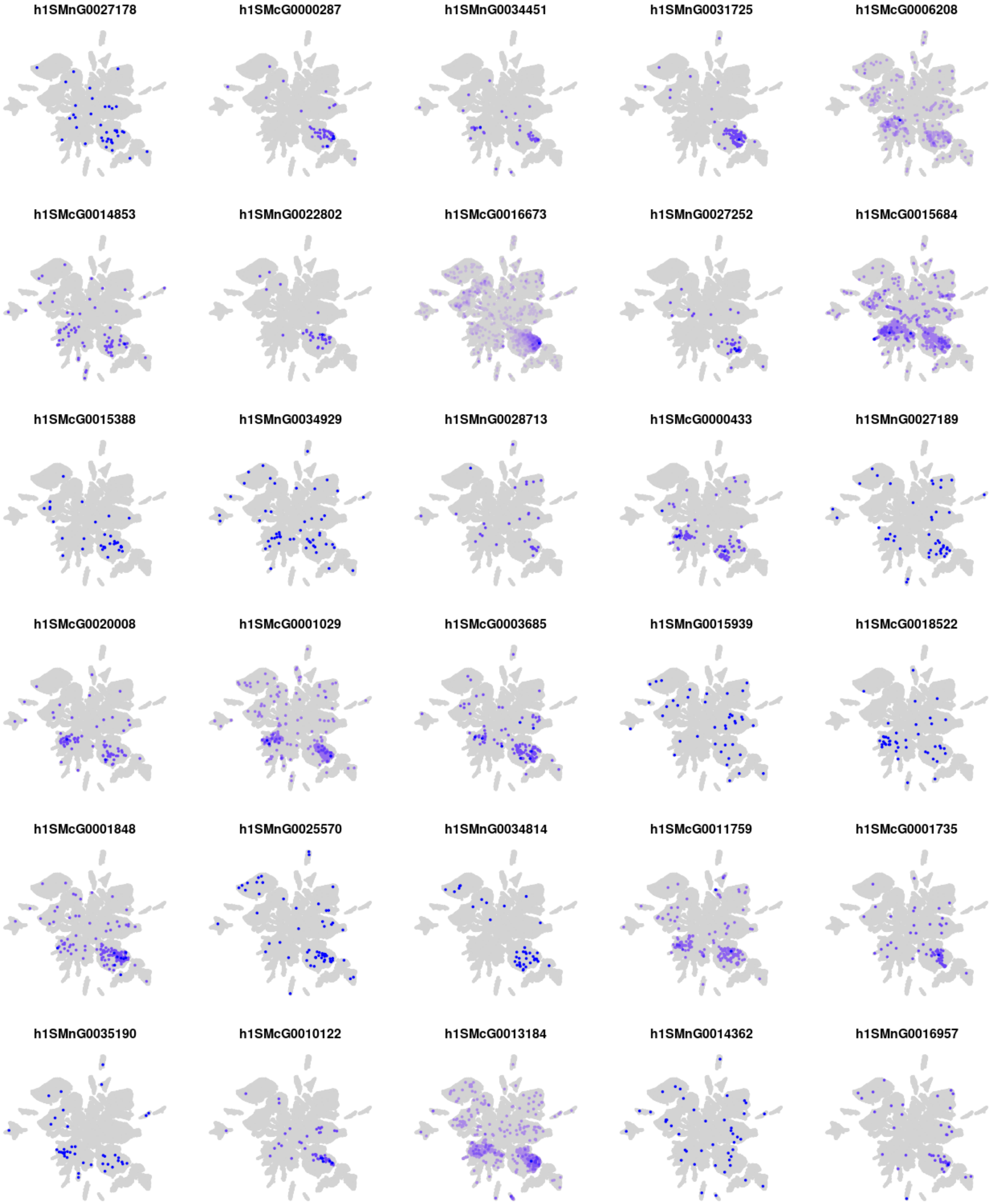
h1SMnG0027178
h1SMcG0000287
h1SMnG0034451
h1SMnG0031725
h1SMcG0006208
h1SMcG0014853
h1SMnG0022802
h1SMcG0016673
h1SMnG0027252
h1SMcG0015684
h1SMcG0015388
h1SMnG0034929
h1SMnG0028713
h1SMcG0000433
h1SMnG0027189
h1SMcG0020008
h1SMcG0001029
h1SMcG0003685
h1SMnG0015939
h1SMcG0018522
h1SMcG0001848
h1SMnG0025570
h1SMnG0034814
h1SMcG0011759
h1SMcG0001735
h1SMnG0035190
h1SMcG0010122
h1SMcG0013184
h1SMnG0014362
h1SMnG0016957

### Module m10

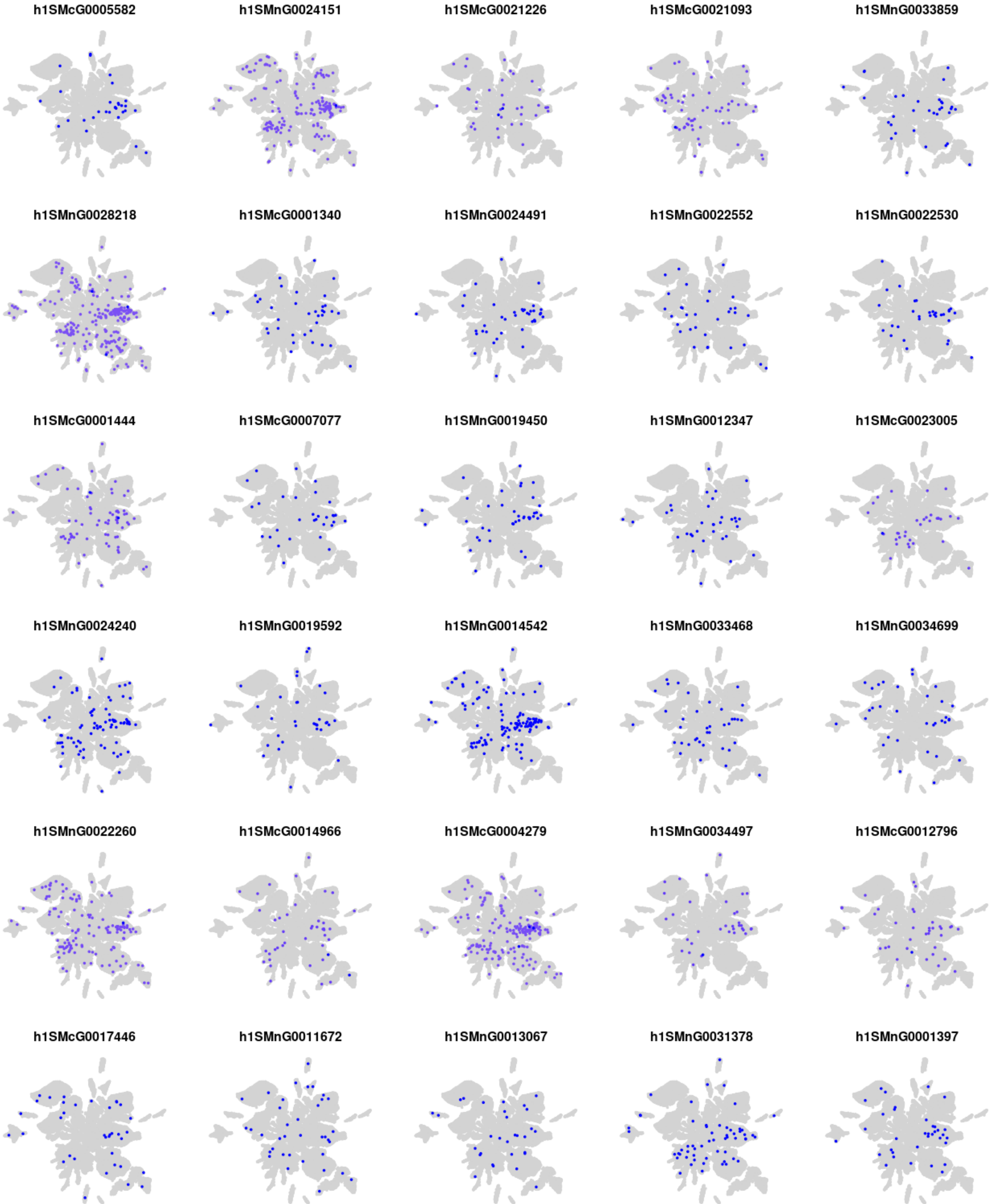

h1SMcG0005582
h1SMnG0024151
h1SMcG0021226
h1SMcG0021093
h1SMnG0033859
h1SMnG0028218
h1SMcG0001340
h1SMnG0024491
h1SMnG0022552
h1SMnG0022530
h1SMcG0001444
h1SMcG0007077
h1SMnG0019450
h1SMnG0012347
h1SMcG0023005
h1SMnG0024240
h1SMnG0019592
h1SMnG0014542
h1SMnG0033468
h1SMnG0034699
h1SMnG0022260
h1SMcG0014966
h1SMcG0004279
h1SMnG0034497
h1SMcG0012796
h1SMcG0017446
h1SMnG0011672
h1SMnG0013067
h1SMnG0031378
h1SMnG0001397

### Module m11

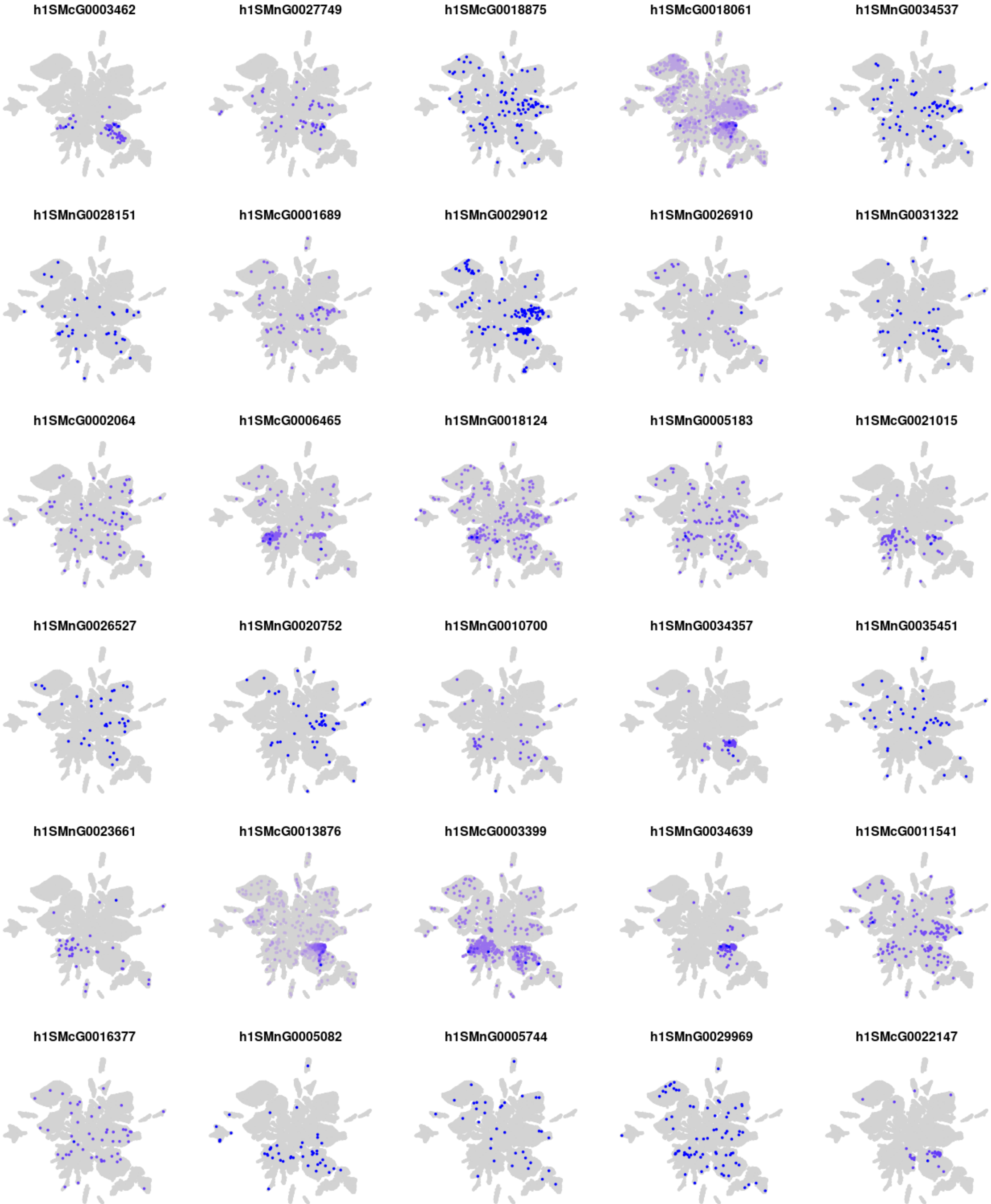
h1SMcG0003462
h1SMnG0027749
h1SMcG0018875
h1SMcG0018061
h1SMnG0034537
h1SMnG0028151
h1SMcG0001689
h1SMnG0029012
h1SMnG0026910
h1SMnG0031322
h1SMcG0002064
h1SMcG0006465
h1SMnG0018124
h1SMnG0005183
h1SMcG0021015
h1SMnG0026527
h1SMnG0020752
h1SMnG0010700
h1SMnG0034357
h1SMnG0035451
h1SMnG0023661
h1SMcG0013876
h1SMcG0003399
h1SMnG0034639
h1SMcG0011541
h1SMcG0016377
h1SMnG0005082
h1SMnG0005744
h1SMnG0029969
h1SMcG0022147

### Module m12

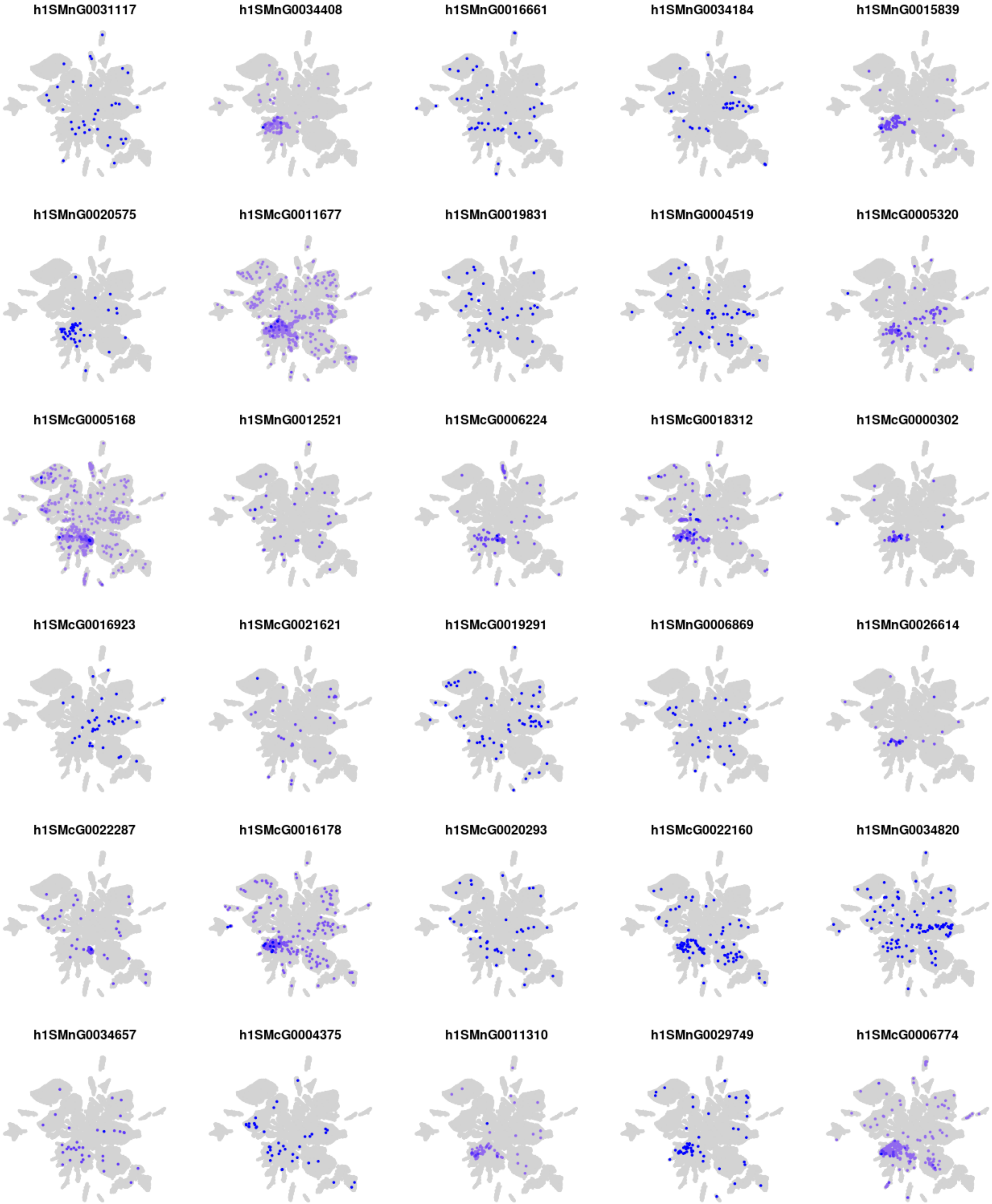
h1SMnG0031117
h1SMnG0034408
h1SMnG0016661
h1SMnG0034184
h1SMnG0015839
h1SMnG0020575
h1SMcG0011677
h1SMnG0019831
h1SMnG0004519
h1SMcG0005320
h1SMcG0005168
h1SMnG0012521
h1SMcG0006224
h1SMcG0018312
h1SMcG0000302
h1SMcG0016923
h1SMcG0021621
h1SMcG0019291
h1SMnG0006869
h1SMnG0026614
h1SMcG0022287
h1SMcG0016178
h1SMcG0020293
h1SMcG0022160
h1SMnG0034820
h1SMnG0034657
h1SMcG0004375
h1SMnG0011310
h1SMnG0029749
h1SMcG0006774

#### Module m13

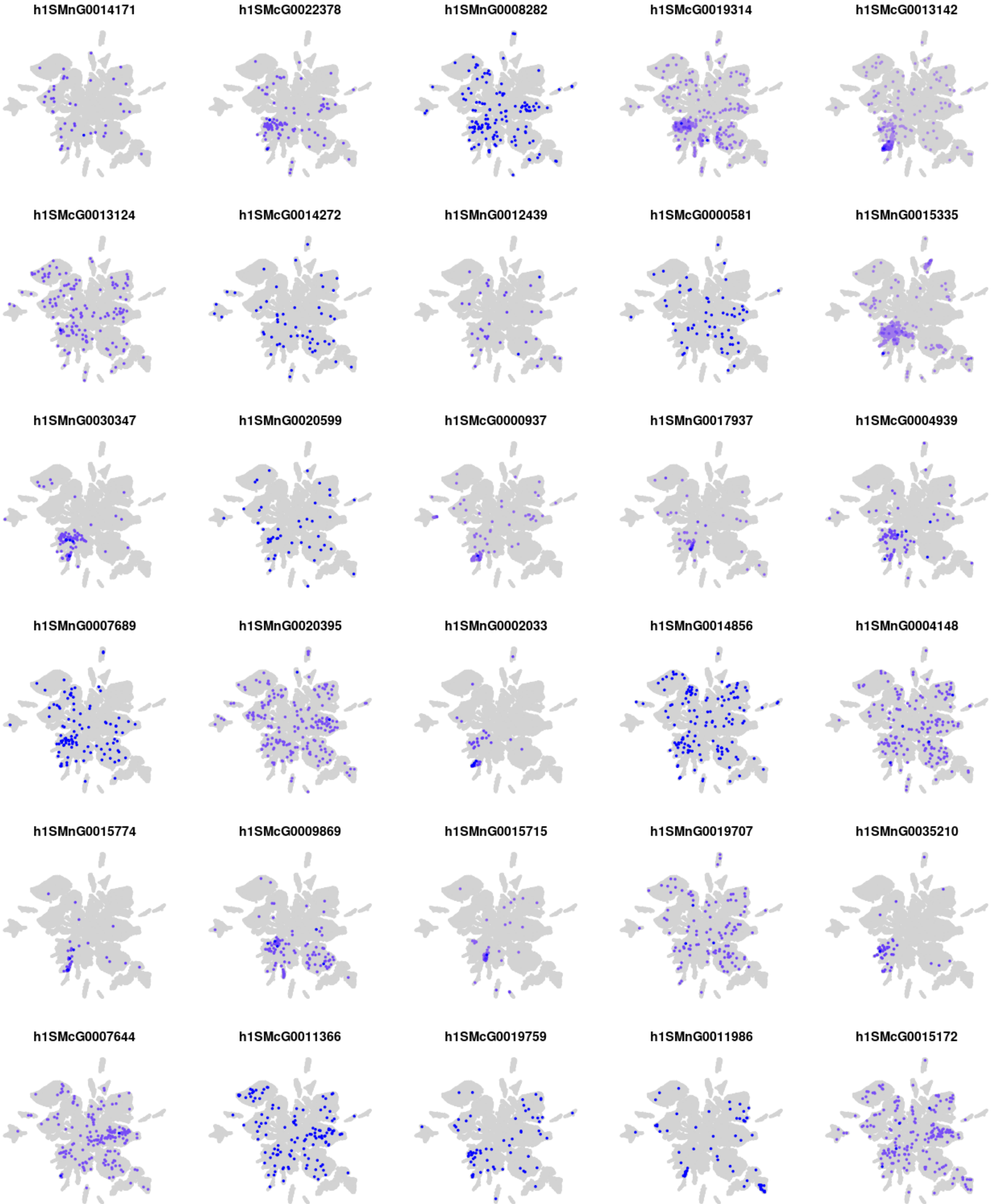

h1SMnG0014171
h1SMcG0022378
h1SMnG0008282
h1SMcG0019314
h1SMcG0013142
h1SMcG0013124
h1SMcG0014272
h1SMnG0012439
h1SMcG0000581
h1SMnG0015335
h1SMnG0030347
h1SMnG0020599
h1SMcG0000937
h1SMnG0017937
h1SMcG0004939
h1SMnG0007689
h1SMnG0020395
h1SMnG0002033
h1SMnG0014856
h1SMnG0004148
h1SMnG0015774
h1SMcG0009869
h1SMnG0015715
h1SMnG0019707
h1SMnG0035210
h1SMcG0007644
h1SMcG0011366
h1SMcG0019759
h1SMnG0011986
h1SMcG0015172

### Module m14

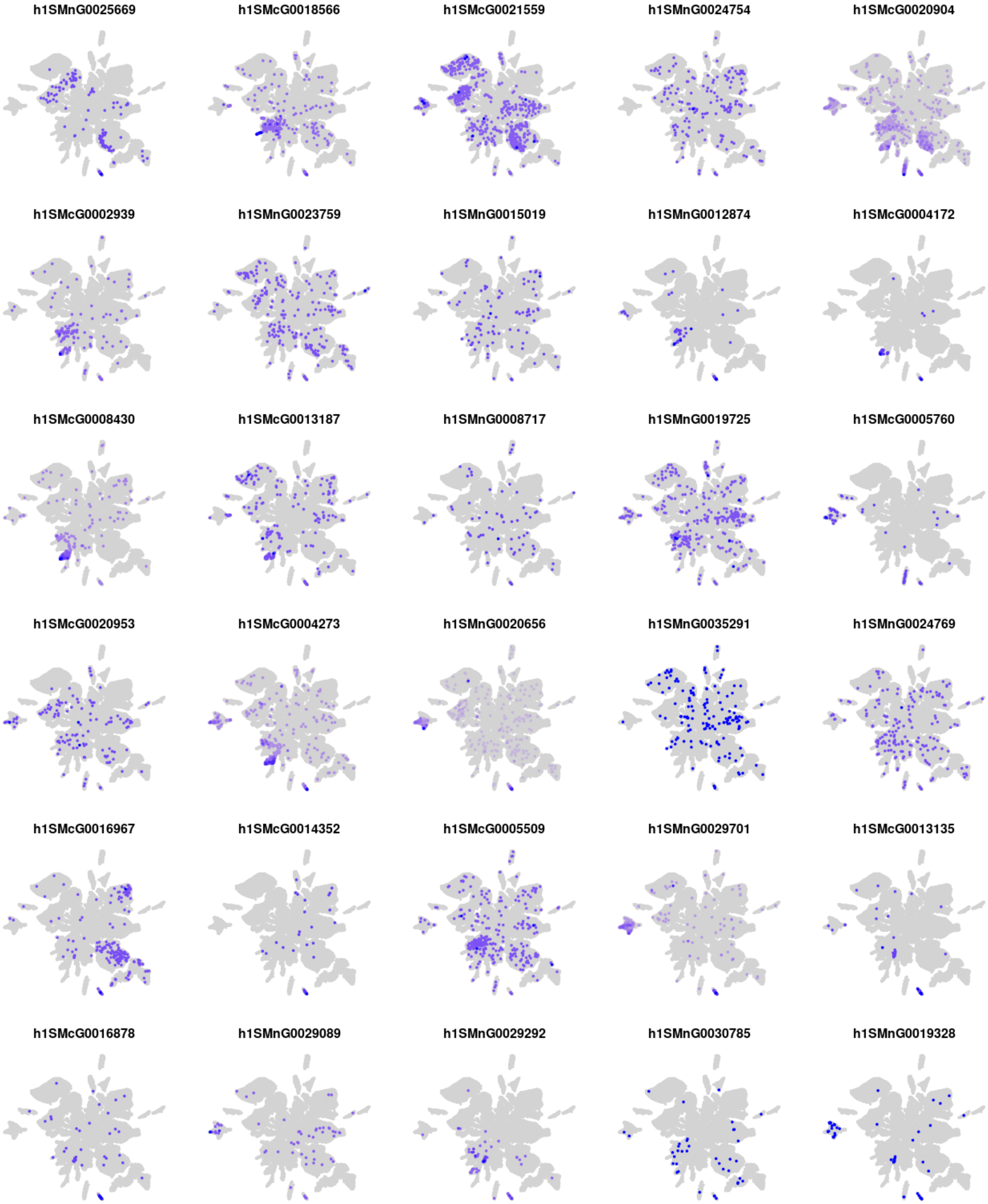
h1SMnG0025669
h1SMcG0018566
h1SMcG0021559
h1SMnG0024754
h1SMcG0020904
h1SMcG0002939
h1SMnG0023759
h1SMnG0015019
h1SMnG0012874
h1SMcG0004172
h1SMcG0008430
h1SMcG0013187
h1SMnG0008717
h1SMnG0019725
h1SMcG0005760
h1SMcG0020953
h1SMcG0004273
h1SMnG0020656
h1SMnG0035291
h1SMnG0024769
h1SMcG0016967
h1SMcG0014352
h1SMcG0005509
h1SMnG0029701
h1SMcG0013135
h1SMcG0016878
h1SMnG0029089
h1SMnG0029292
h1SMnG0030785
h1SMnG0019328

### Module m15

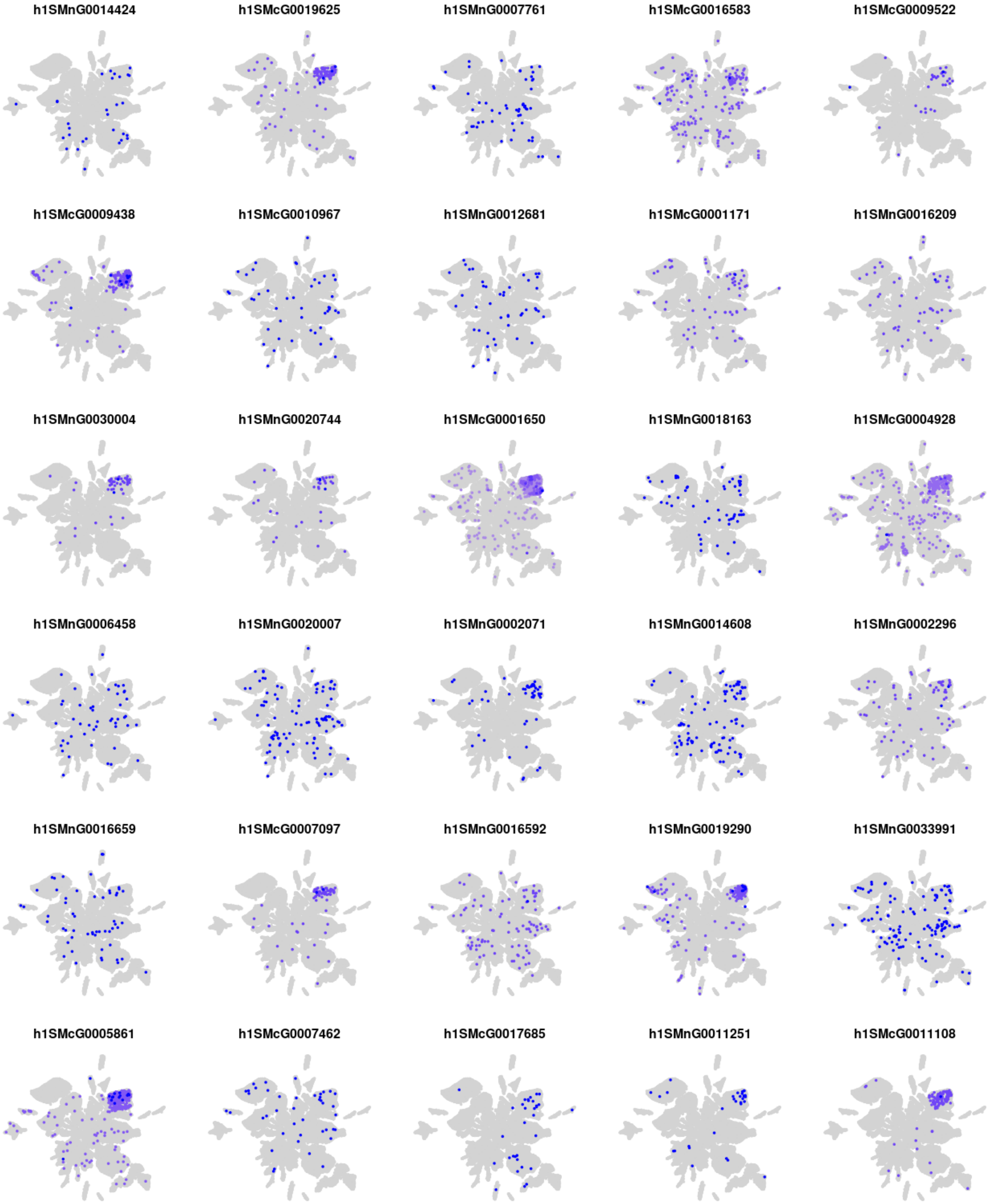
h1SMnG0014424
h1SMcG0019625
h1SMnG0007761
h1SMcG0016583
h1SMcG0009522
h1SMcG0009438
h1SMcG0010967
h1SMnG0012681
h1SMcG0001171
h1SMnG0016209
h1SMnG0030004
h1SMnG0020744
h1SMcG0001650
h1SMnG0018163
h1SMcG0004928
h1SMnG0006458
h1SMnG0020007
h1SMnG0002071
h1SMnG0014608
h1SMnG0002296
h1SMnG0016659
h1SMcG0007097
h1SMnG0016592
h1SMnG0019290
h1SMnG0033991
h1SMcG0005861
h1SMcG0007462
h1SMcG0017685
h1SMnG0011251
h1SMcG0011108

### Module m16

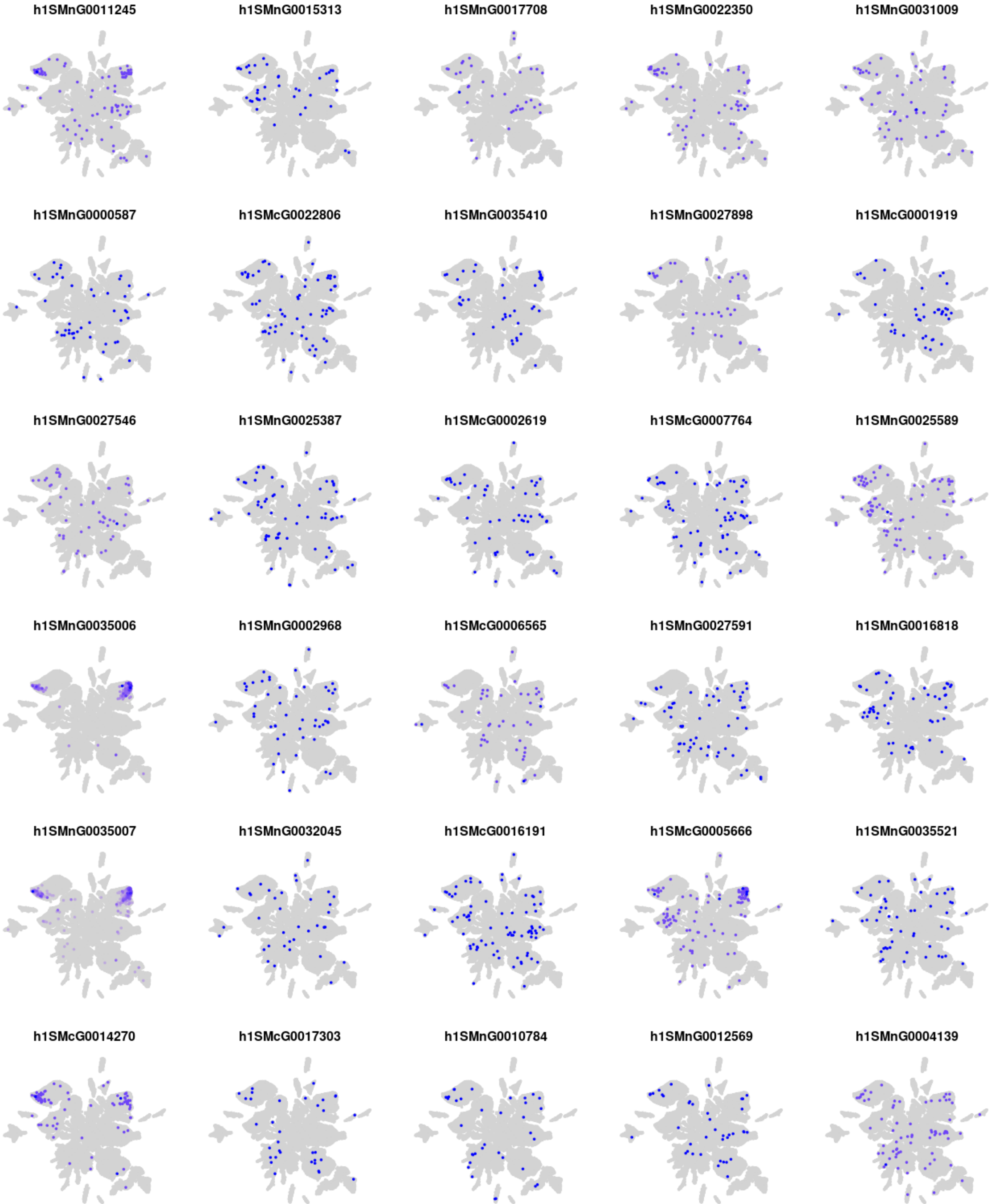
h1SMnG0011245
h1SMnG0015313
h1SMnG0017708
h1SMnG0022350
h1SMnG0031009
h1SMnG0000587
h1SMcG0022806
h1SMnG0035410
h1SMnG0027898
h1SMcG0001919
h1SMnG0027546
h1SMnG0025387
h1SMcG0002619
h1SMcG0007764
h1SMnG0025589
h1SMnG0035006
h1SMnG0002968
h1SMcG0006565
h1SMnG0027591
h1SMnG0016818
h1SMnG0035007
h1SMnG0032045
h1SMcG0016191
h1SMcG0005666
h1SMnG0035521
h1SMcG0014270
h1SMcG0017303
h1SMnG0010784
h1SMnG0012569
h1SMnG0004139

### Module m17

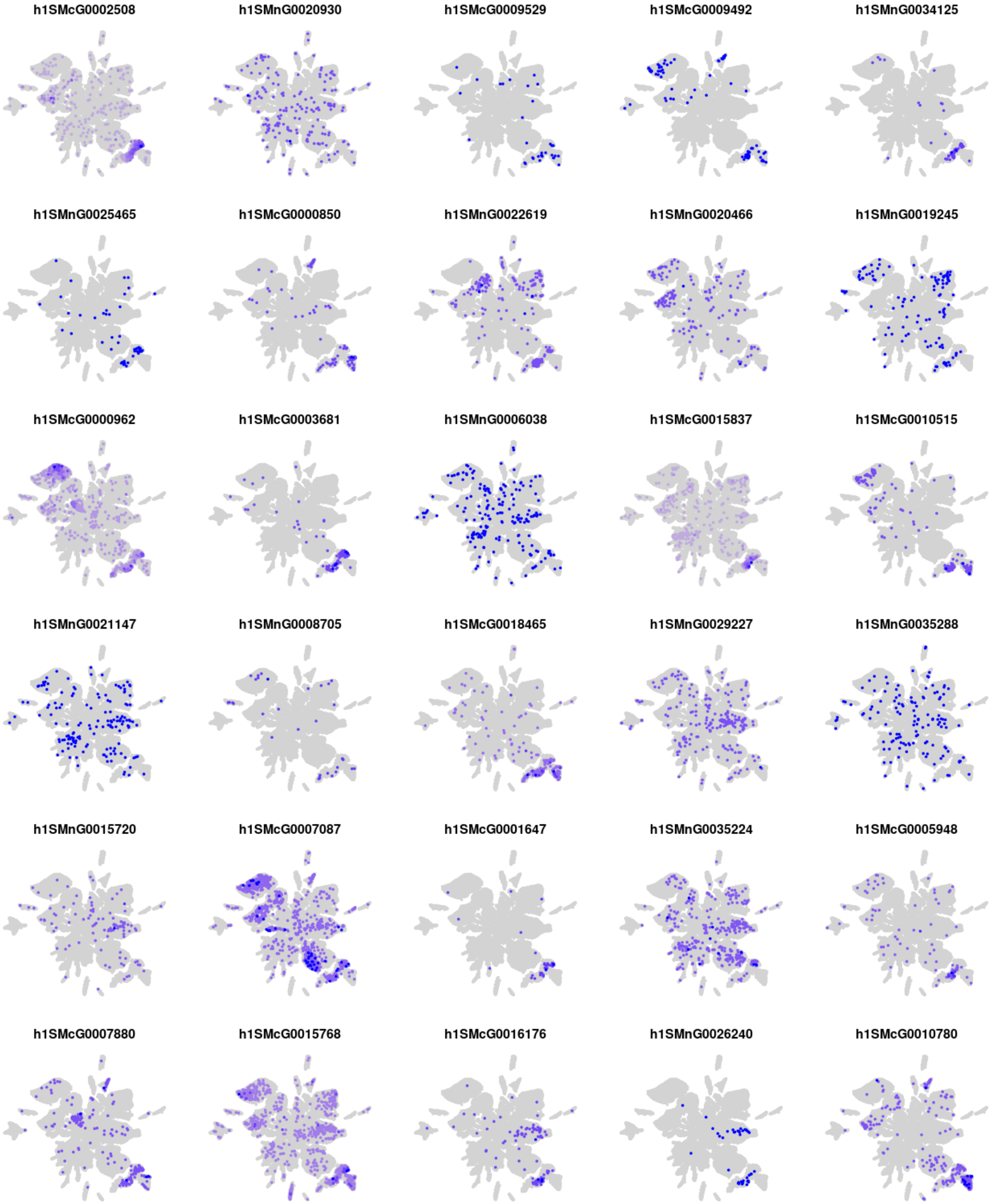
h1SMcG0002508
h1SMnG0020930
h1SMcG0009529
h1SMcG0009492
h1SMnG0034125
h1SMnG0025465
h1SMcG0000850
h1SMnG0022619
h1SMnG0020466
h1SMnG0019245
h1SMcG0000962
h1SMcG0003681
h1SMnG0006038
h1SMcG0015837
h1SMcG0010515
h1SMnG0021147
h1SMnG0008705
h1SMcG0018465
h1SMnG0029227
h1SMnG0035288
h1SMnG0015720
h1SMcG0007087
h1SMcG0001647
h1SMnG0035224
h1SMcG0005948
h1SMcG0007880
h1SMcG0015768
h1SMcG0016176
h1SMnG0026240
h1SMcG0010780

### Module m18

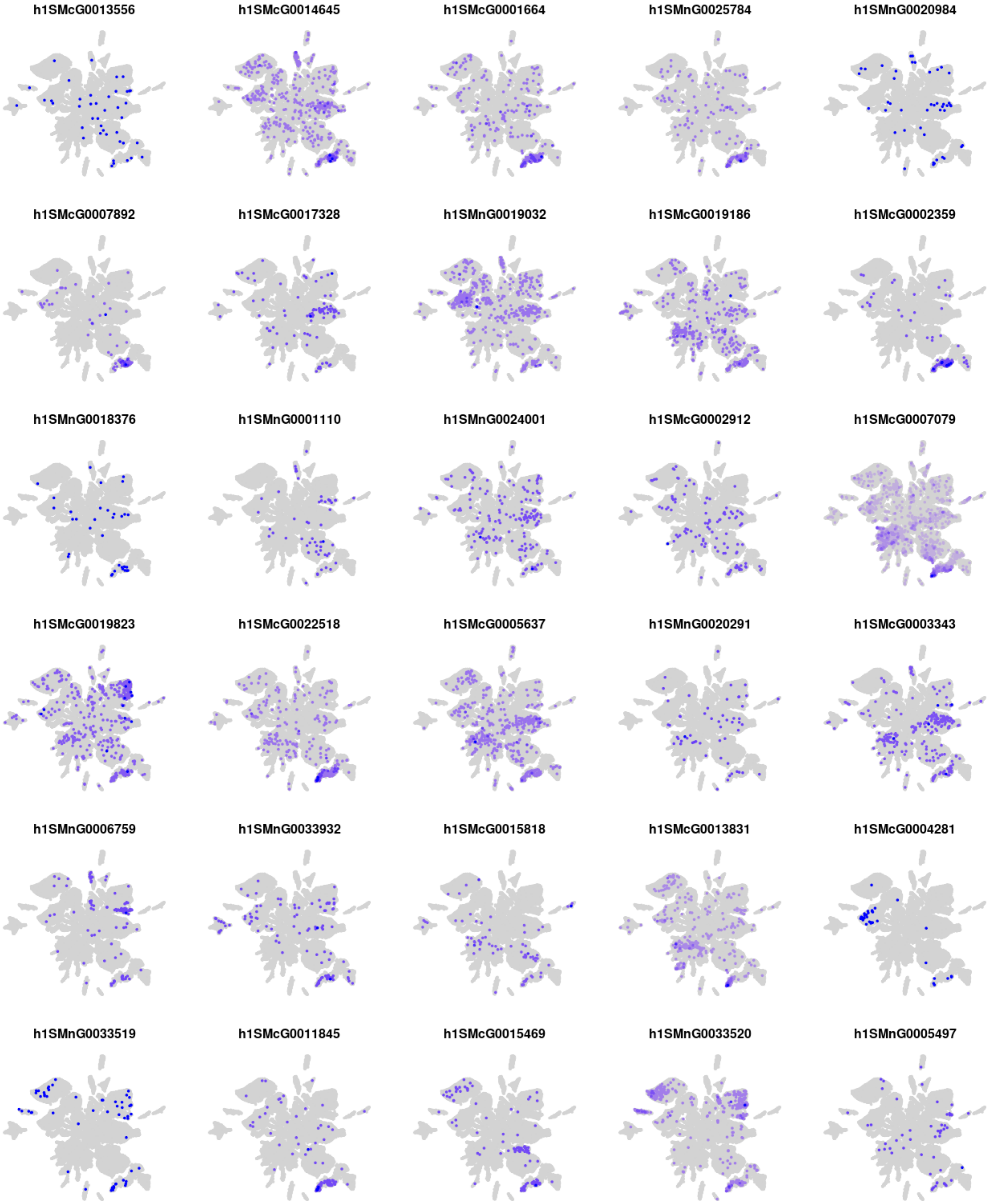
h1SMcG0013556
h1SMcG0014645
h1SMcG0001664
h1SMnG0025784
h1SMnG0020984
h1SMcG0007892
h1SMcG0017328
h1SMnG0019032
h1SMcG0019186
h1SMcG0002359
h1SMnG0018376
h1SMnG0001110
h1SMnG0024001
h1SMcG0002912
h1SMcG0007079
h1SMcG0019823
h1SMcG0022518
h1SMcG0005637
h1SMnG0020291
h1SMcG0003343
h1SMnG0006759
h1SMnG0033932
h1SMcG0015818
h1SMcG0013831
h1SMcG0004281
h1SMnG0033519
h1SMcG0011845
h1SMcG0015469
h1SMnG0033520
h1SMnG0005497

### Module m19

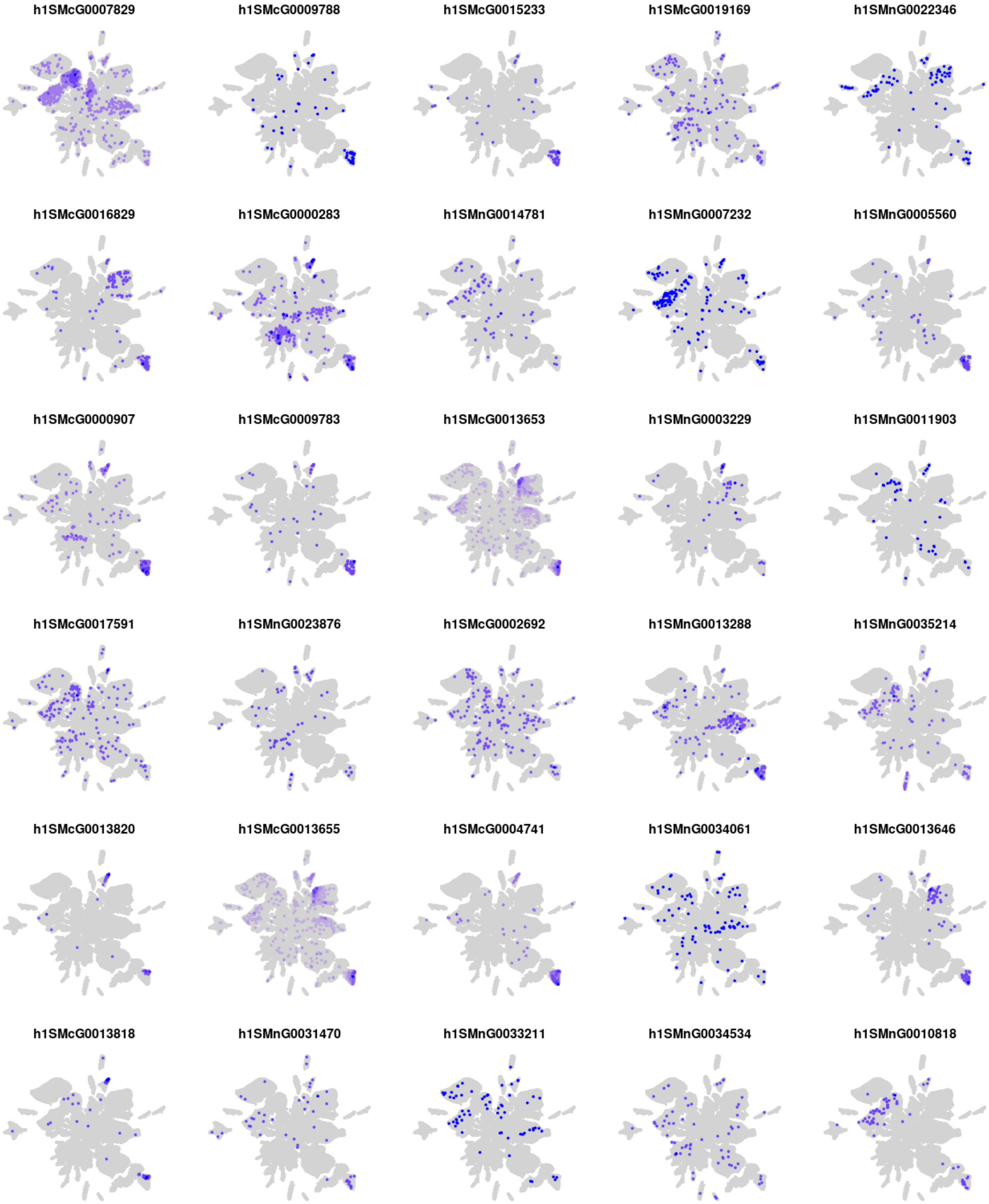
h1SMcG0007829
h1SMcG0009788
h1SMcG0015233
h1SMcG0019169
h1SMnG0022346
h1SMcG0016829
h1SMcG0000283
h1SMnG0014781
h1SMnG0007232
h1SMnG0005560
h1SMcG0000907
h1SMcG0009783
h1SMcG0013653
h1SMnG0003229
h1SMnG0011903
h1SMcG0017591
h1SMnG0023876
h1SMcG0002692
h1SMnG0013288
h1SMnG0035214
h1SMcG0013820
h1SMcG0013655
h1SMcG0004741
h1SMnG0034061
h1SMcG0013646
h1SMcG0013818
h1SMnG0031470
h1SMnG0033211
h1SMnG0034534
h1SMnG0010818

### Module m20

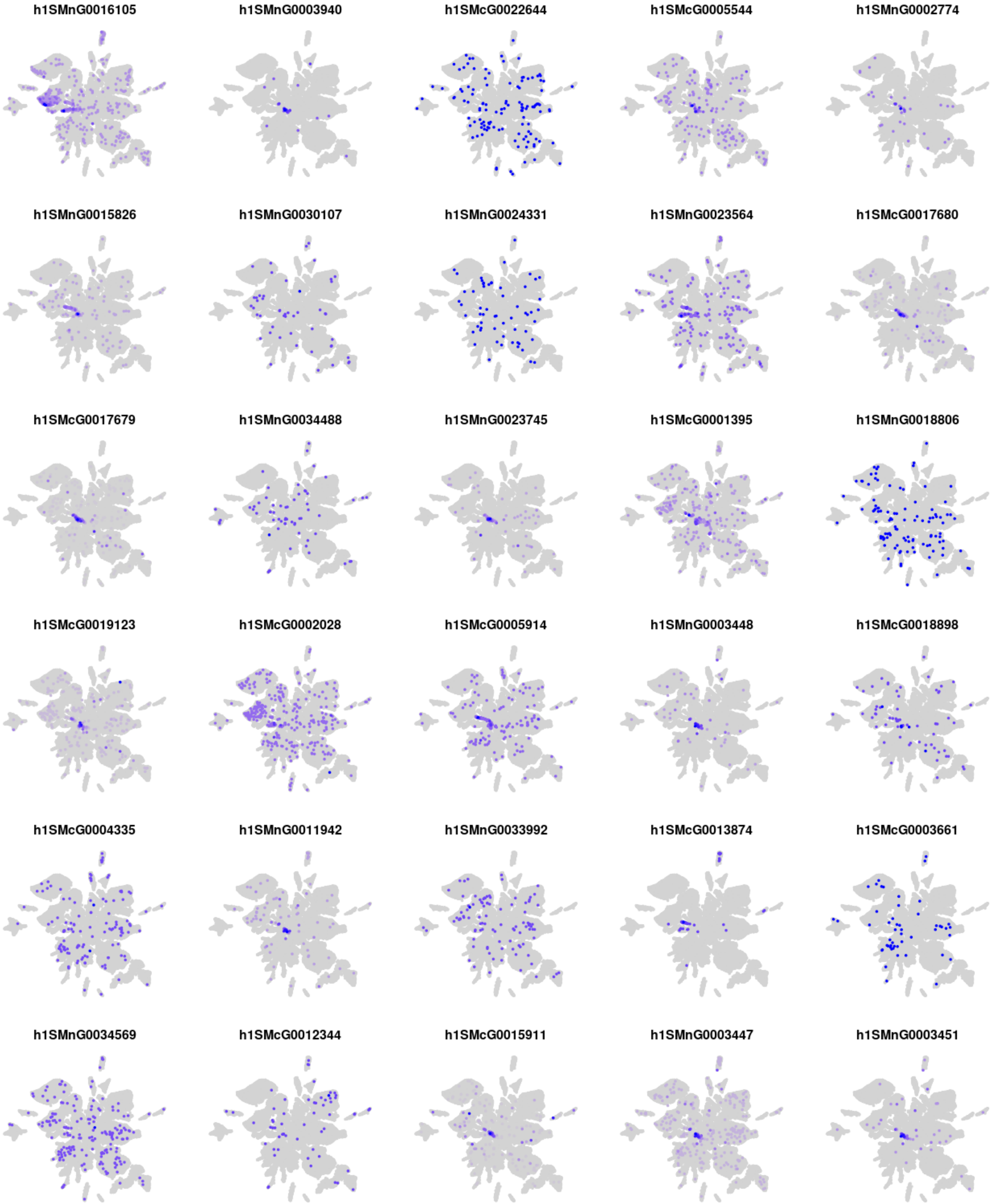

h1SMnG0016105
h1SMnG0003940
h1SMcG0022644
h1SMcG0005544
h1SMnG0002774
h1SMnG0015826
h1SMnG0030107
h1SMnG0024331
h1SMnG0023564
h1SMcG0017680
h1SMcG0017679
h1SMnG0034488
h1SMnG0023745
h1SMcG0001395
h1SMnG0018806
h1SMcG0019123
h1SMcG0002028
h1SMcG0005914
h1SMnG0003448
h1SMcG0018898
h1SMcG0004335
h1SMnG0011942
h1SMnG0033992
h1SMcG0013874
h1SMcG0003661
h1SMnG0034569
h1SMcG0012344
h1SMcG0015911
h1SMnG0003447
h1SMnG0003451

### Module m21

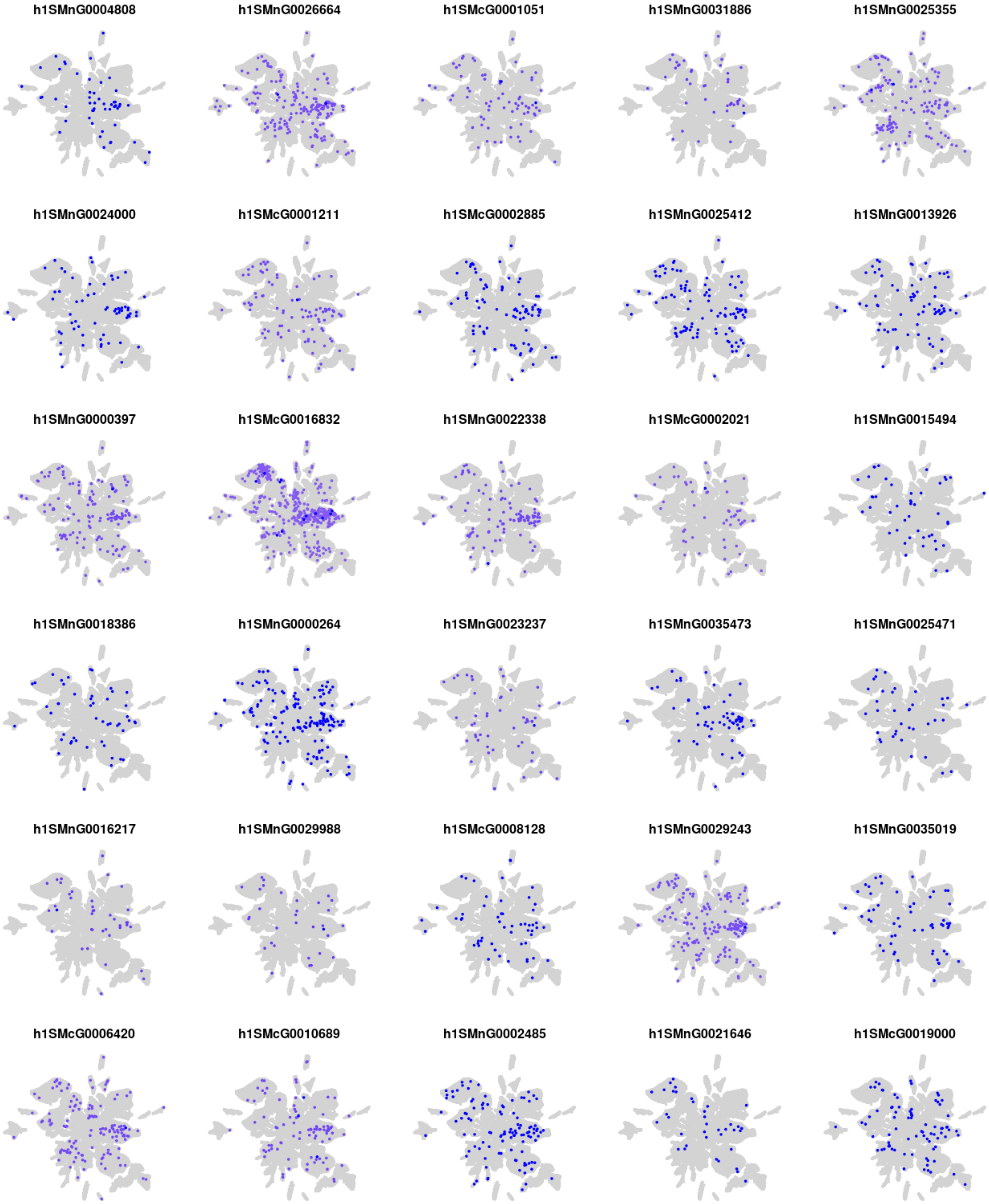

h1SMnG0004808
h1SMnG0026664
h1SMcG0001051
h1SMnG0031886
h1SMnG0025355
h1SMnG0024000
h1SMcG0001211
h1SMcG0002885
h1SMnG0025412
h1SMnG0013926
h1SMnG0000397
h1SMcG0016832
h1SMnG0022338
h1SMcG0002021
h1SMnG0015494
h1SMnG0018386
h1SMnG0000264
h1SMnG0023237
h1SMnG0035473
h1SMnG0025471
h1SMnG0016217
h1SMnG0029988
h1SMcG0008128
h1SMnG0029243
h1SMnG0035019
h1SMcG0006420
h1SMcG0010689
h1SMnG0002485
h1SMnG0021646
h1SMcG0019000

### Module m22

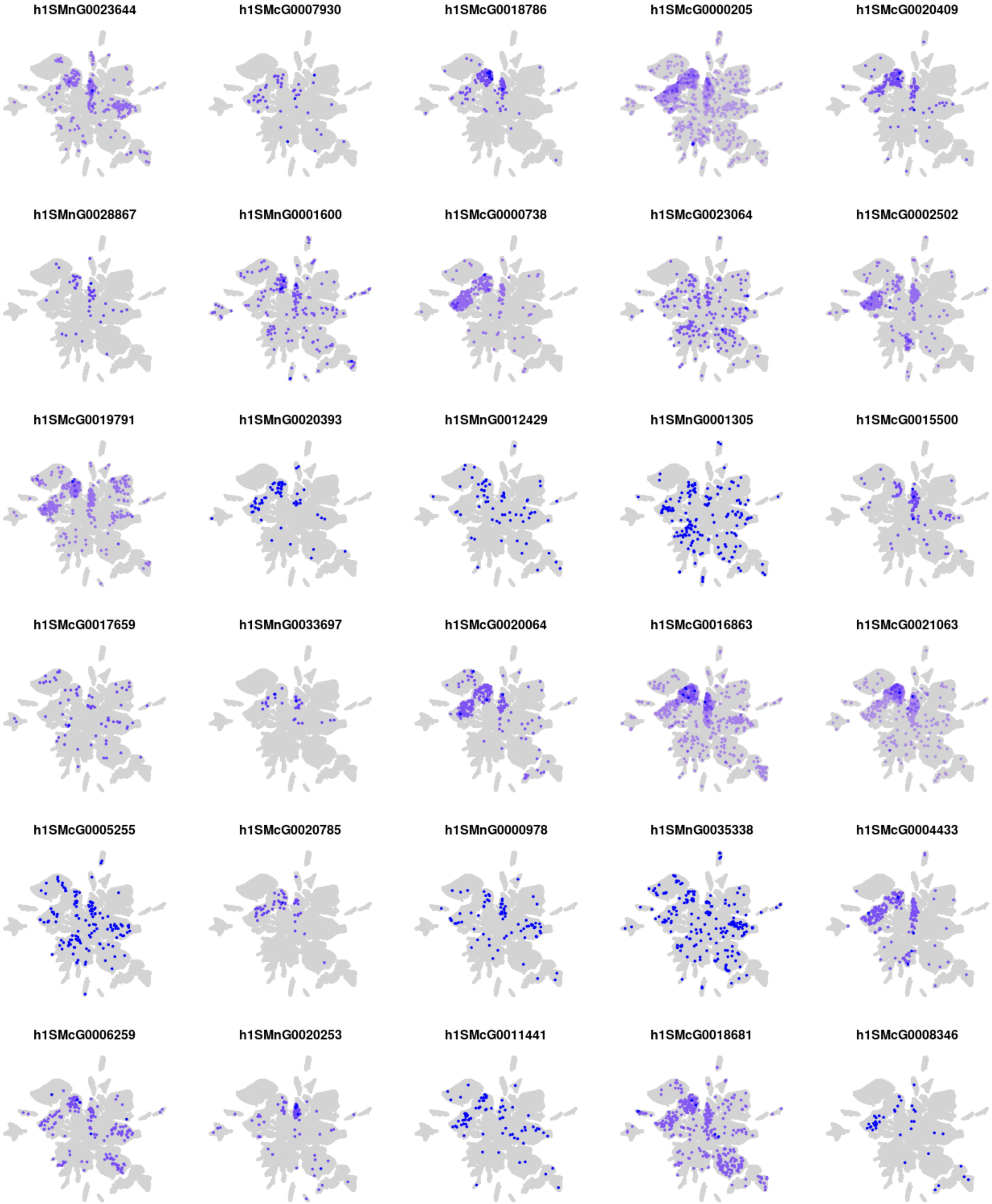
h1SMnG0023644
h1SMcG0007930
h1SMcG0018786
h1SMcG0000205
h1SMcG0020409
h1SMnG0028867
h1SMnG0001600
h1SMcG0000738
h1SMcG0023064
h1SMcG0002502
h1SMcG0019791
h1SMnG0020393
h1SMnG0012429
h1SMnG0001305
h1SMcG0015500
h1SMcG0017659
h1SMnG0033697
h1SMcG0020064
h1SMcG0016863
h1SMcG0021063
h1SMcG0005255
h1SMcG0020785
h1SMnG0000978
h1SMnG0035338
h1SMcG0004433
h1SMcG0006259
h1SMnG0020253
h1SMcG0011441
h1SMcG0018681
h1SMcG0008346

### Module m23

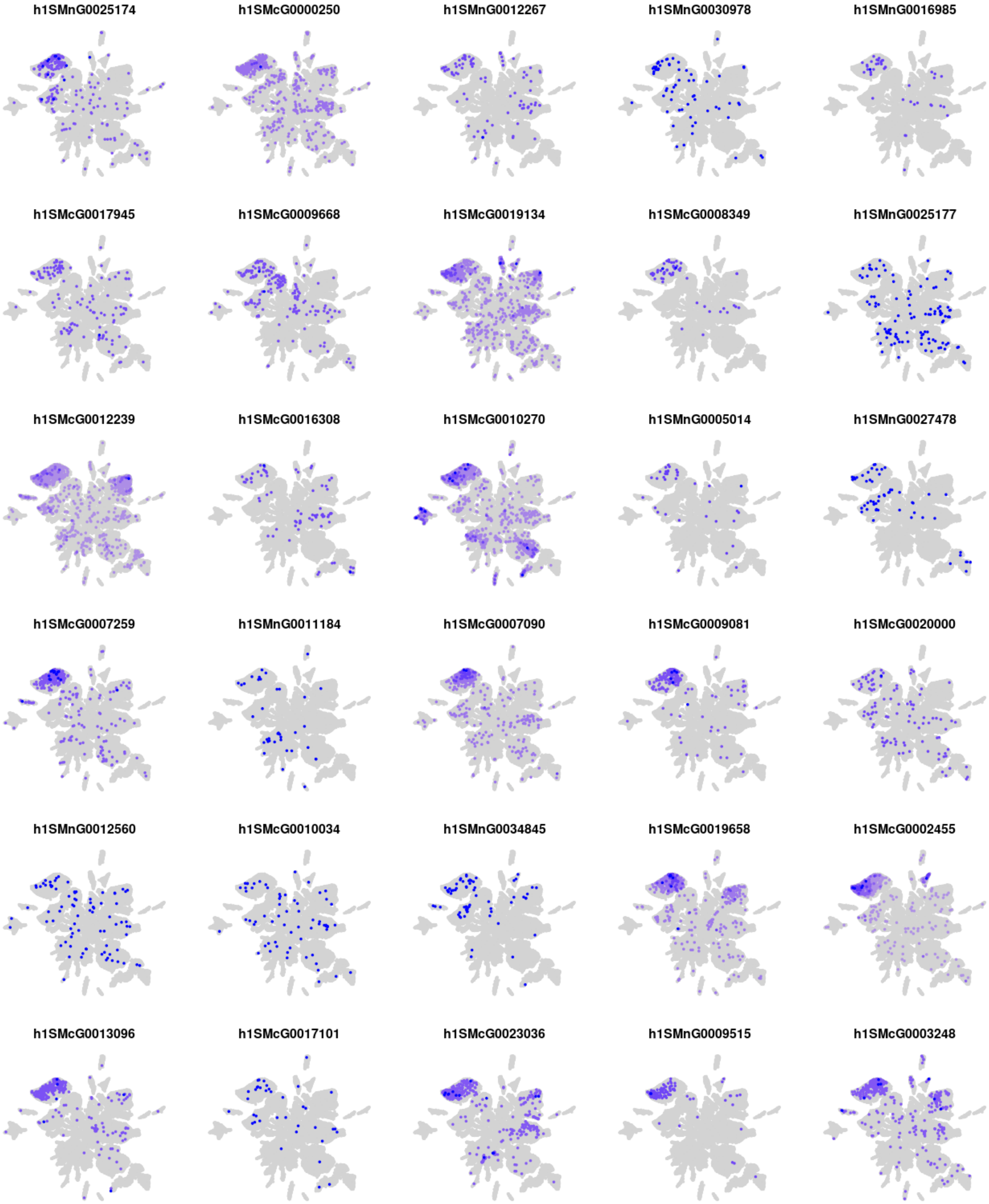
h1SMnG0025174
h1SMcG0000250
h1SMnG0012267
h1SMnG0030978
h1SMnG0016985
h1SMcG0017945
h1SMcG0009668
h1SMcG0019134
h1SMcG0008349
h1SMnG0025177
h1SMcG0012239
h1SMcG0016308
h1SMcG0010270
h1SMnG0005014
h1SMnG0027478
h1SMcG0007259
h1SMnG0011184
h1SMcG0007090
h1SMcG0009081
h1SMcG0020000
h1SMnG0012560
h1SMcG0010034
h1SMnG0034845
h1SMcG0019658
h1SMcG0002455
h1SMcG0013096
h1SMcG0017101
h1SMcG0023036
h1SMnG0009515
h1SMcG0003248

### Module m24

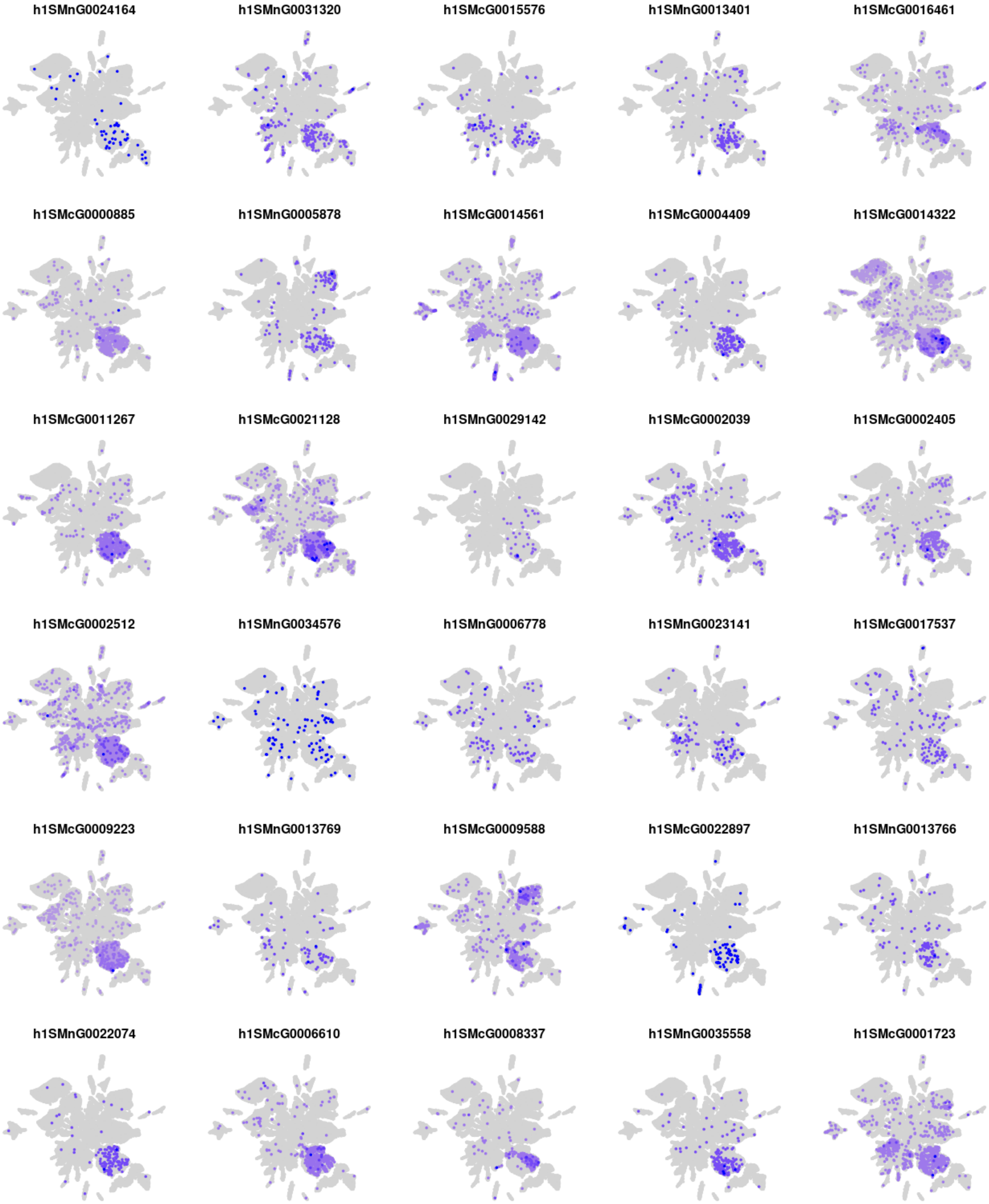
h1SMnG0024164
h1SMnG0031320
h1SMcG0015576
h1SMnG0013401
h1SMcG0016461
h1SMcG0000885
h1SMnG0005878
h1SMcG0014561
h1SMcG0004409
h1SMcG0014322
h1SMcG0011267
h1SMcG0021128
h1SMnG0029142
h1SMcG0002039
h1SMcG0002405
h1SMcG0002512
h1SMnG0034576
h1SMnG0006778
h1SMnG0023141
h1SMcG0017537
h1SMcG0009223
h1SMnG0013769
h1SMcG0009588
h1SMcG0022897
h1SMnG0013766
h1SMnG0022074
h1SMcG0006610
h1SMcG0008337
h1SMnG0035558
h1SMcG0001723

### Module m25

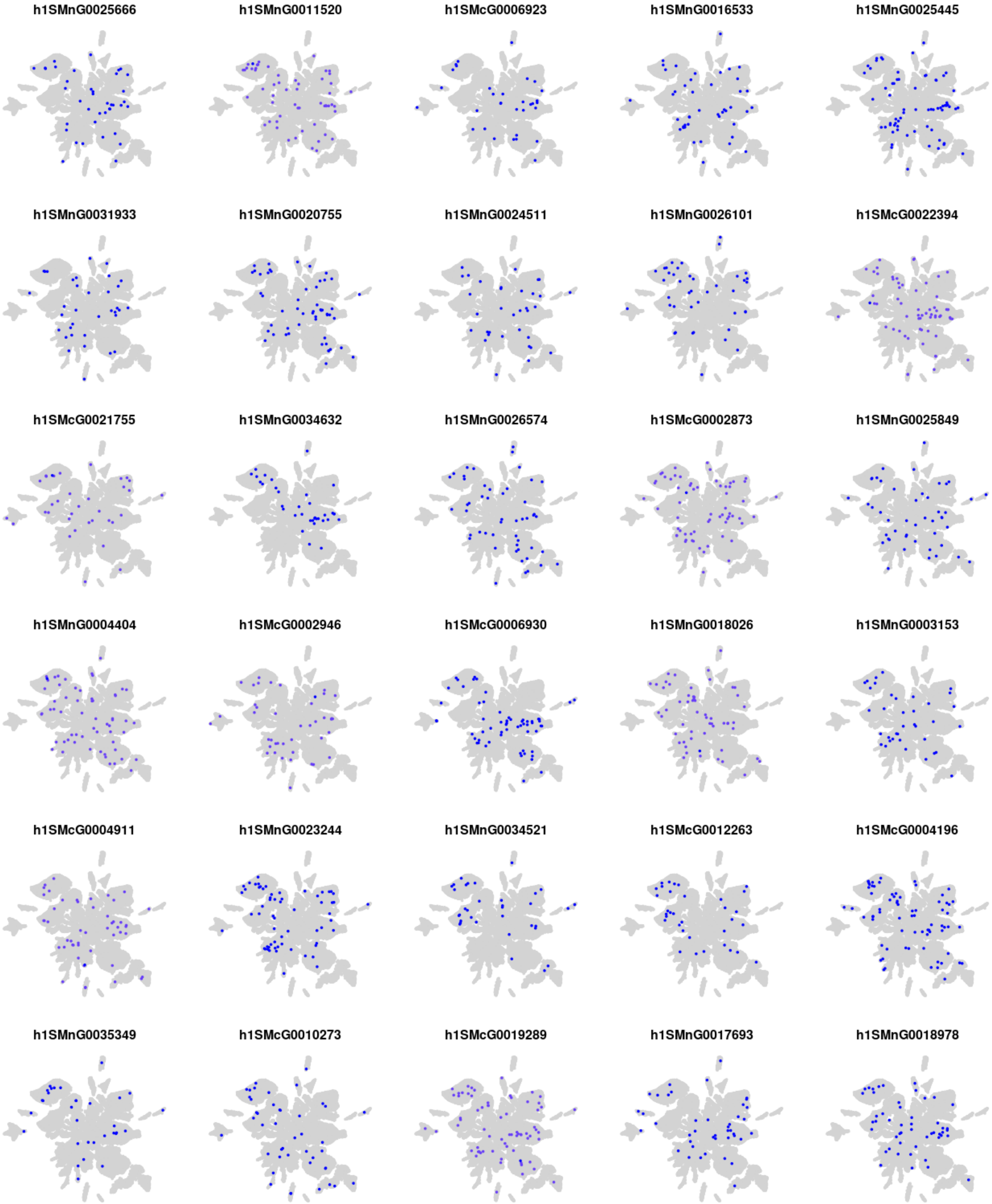
h1SMnG0025666
h1SMnG0011520
h1SMcG0006923
h1SMnG0016533
h1SMnG0025445
h1SMnG0031933
h1SMnG0020755
h1SMnG0024511
h1SMnG0026101
h1SMcG0022394
h1SMcG0021755
h1SMnG0034632
h1SMnG0026574
h1SMcG0002873
h1SMnG0025849
h1SMnG0004404
h1SMcG0002946
h1SMcG0006930
h1SMnG0018026
h1SMnG0003153
h1SMcG0004911
h1SMnG0023244
h1SMnG0034521
h1SMcG0012263
h1SMcG0004196
h1SMnG0035349
h1SMcG0010273
h1SMcG0019289
h1SMnG0017693
h1SMnG0018978

### Module m26

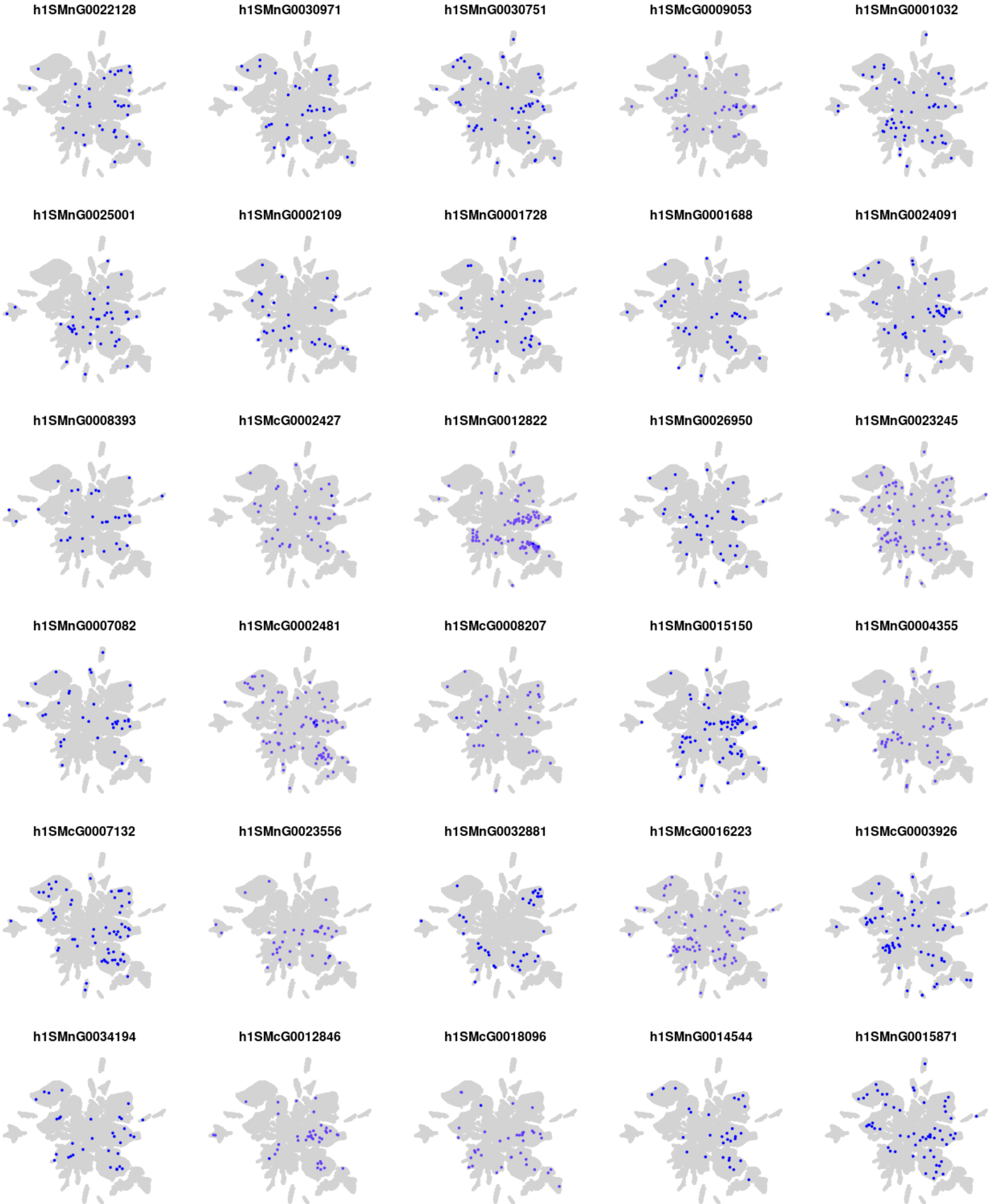
h1SMnG0022128
h1SMnG0030971
h1SMnG0030751
h1SMcG0009053
h1SMnG0001032
h1SMnG0025001
h1SMnG0002109
h1SMnG0001728
h1SMnG0001688
h1SMnG0024091
h1SMnG0008393
h1SMcG0002427
h1SMnG0012822
h1SMnG0026950
h1SMnG0023245
h1SMnG0007082
h1SMcG0002481
h1SMcG0008207
h1SMnG0015150
h1SMnG0004355
h1SMcG0007132
h1SMnG0023556
h1SMnG0032881
h1SMcG0016223
h1SMcG0003926
h1SMnG0034194
h1SMcG0012846
h1SMcG0018096
h1SMnG0014544
h1SMnG0015871

### Module m27

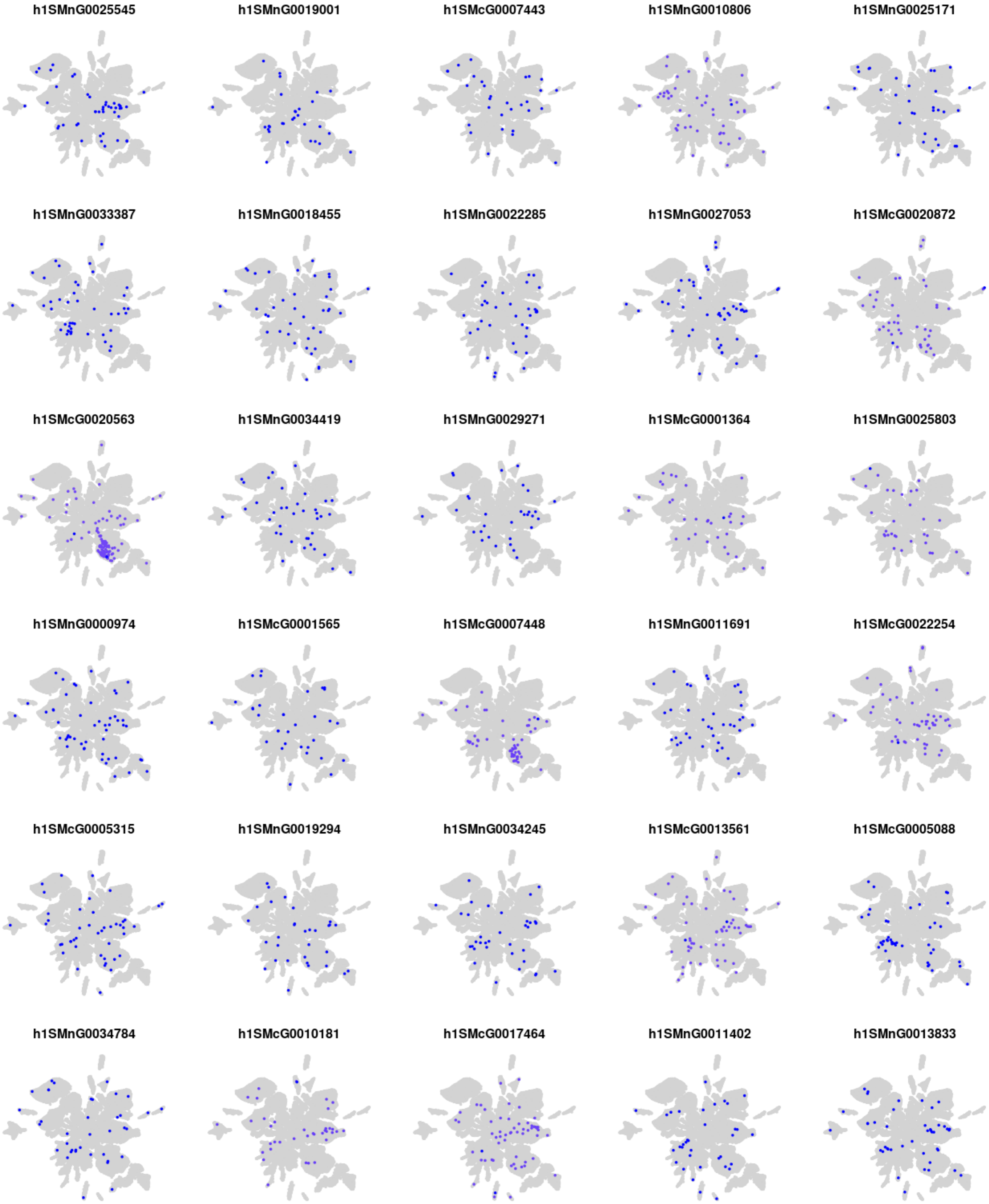

h1SMnG0025545
h1SMnG0019001
h1SMcG0007443
h1SMnG0010806
h1SMnG0025171
h1SMnG0033387
h1SMnG0018455
h1SMnG0022285
h1SMnG0027053
h1SMcG0020872
h1SMcG0020563
h1SMnG0034419
h1SMnG0029271
h1SMcG0001364
h1SMnG0025803
h1SMnG0000974
h1SMcG0001565
h1SMcG0007448
h1SMnG0011691
h1SMcG0022254
h1SMcG0005315
h1SMnG0019294
h1SMnG0034245
h1SMcG0013561
h1SMcG0005088
h1SMnG0034784
h1SMcG0010181
h1SMcG0017464
h1SMnG0011402
h1SMnG0013833

### Module m28

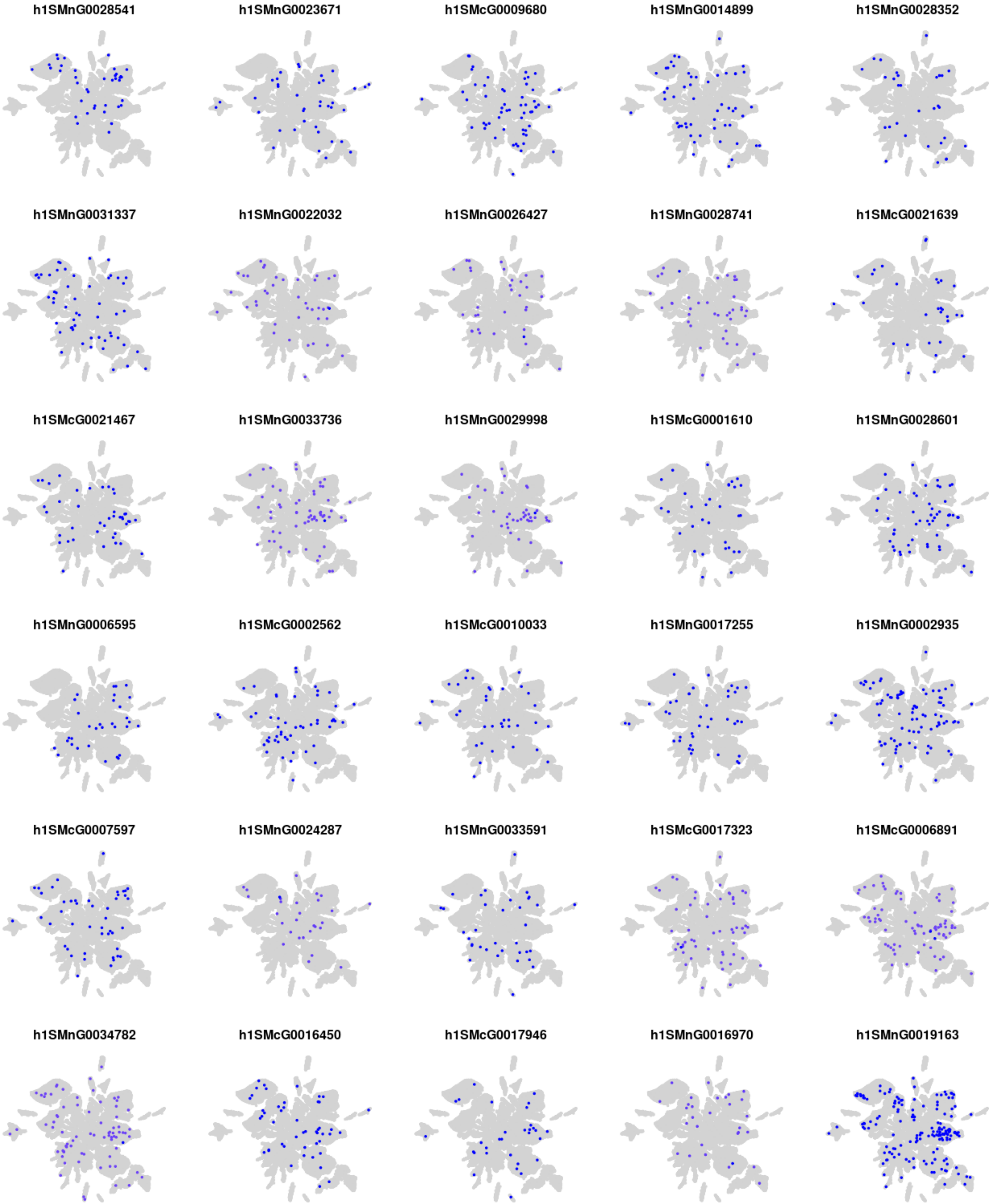
h1SMnG0028541
h1SMnG0023671
h1SMcG0009680
h1SMnG0014899
h1SMnG0028352
h1SMnG0031337
h1SMnG0022032
h1SMnG0026427
h1SMnG0028741
h1SMcG0021639
h1SMcG0021467
h1SMnG0033736
h1SMnG0029998
h1SMcG0001610
h1SMnG0028601
h1SMnG0006595
h1SMcG0002562
h1SMcG0010033
h1SMnG0017255
h1SMnG0002935
h1SMcG0007597
h1SMnG0024287
h1SMnG0033591
h1SMcG0017323
h1SMcG0006891
h1SMnG0034782
h1SMcG0016450
h1SMcG0017946
h1SMnG0016970
h1SMnG0019163

### Module m29

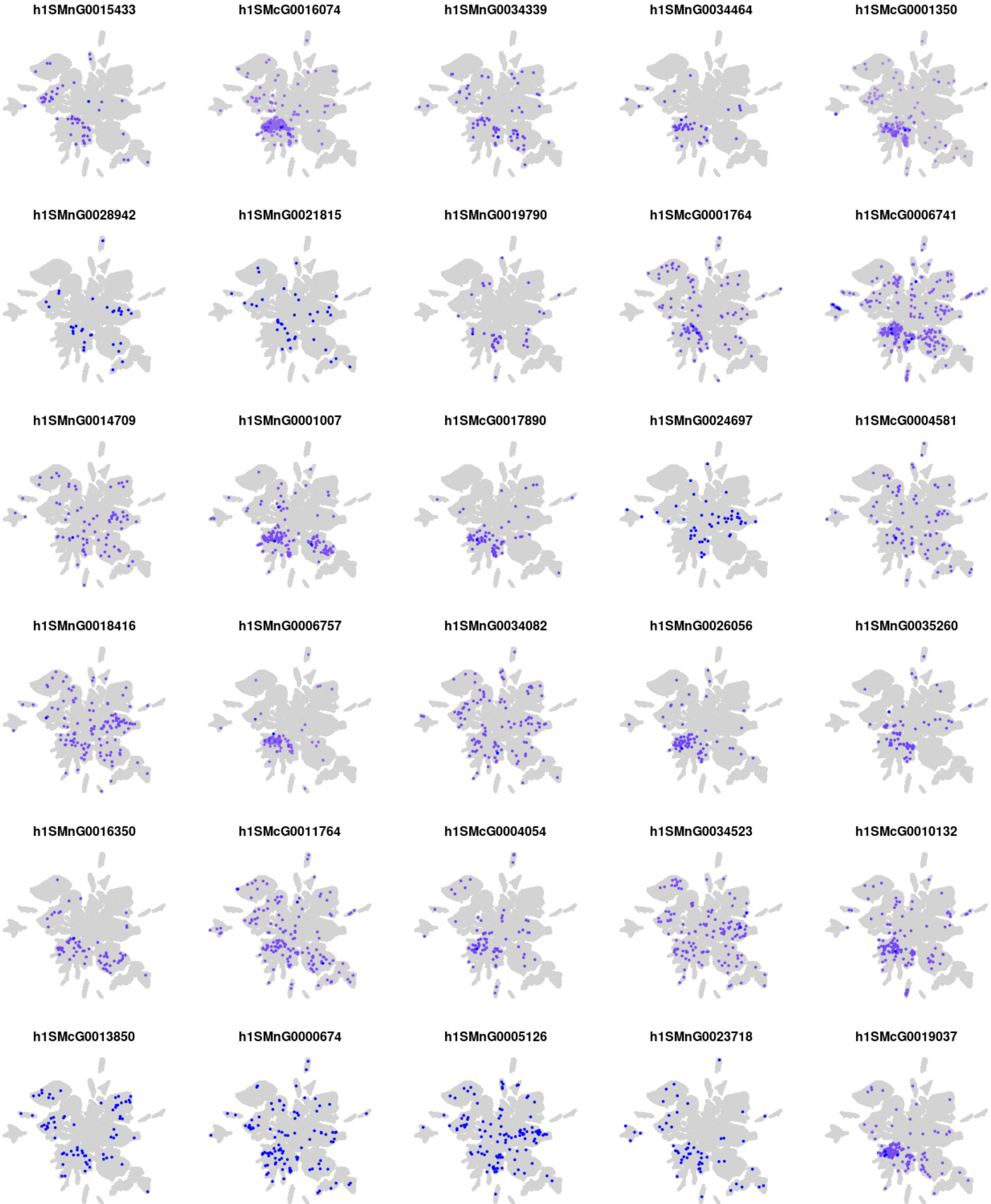
h1SMnG0015433
h1SMcG0016074
h1SMnG0034339
h1SMnG0034464
h1SMcG0001350
h1SMnG0028942
h1SMnG0021815
h1SMnG0019790
h1SMcG0001764
h1SMcG0006741
h1SMnG0014709
h1SMnG0001007
h1SMcG0017890
h1SMnG0024697
h1SMcG0004581
h1SMnG0018416
h1SMnG0006757
h1SMnG0034082
h1SMnG0026056
h1SMnG0035260
h1SMnG0016350
h1SMcG0011764
h1SMcG0004054
h1SMnG0034523
h1SMcG0010132
h1SMcG0013850
h1SMnG0000674
h1SMnG0005126
h1SMnG0023718
h1SMcG0019037

### Module m30

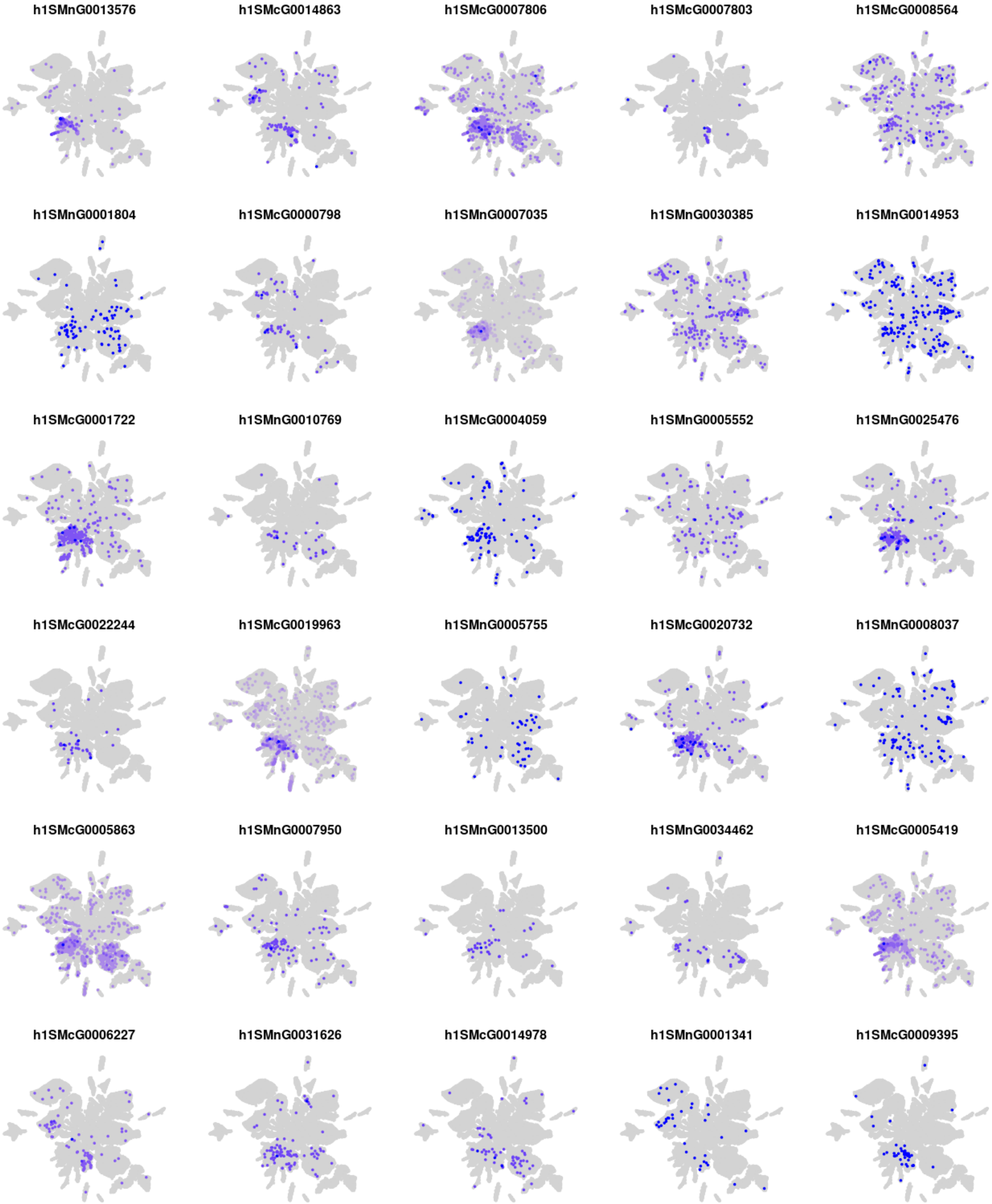

h1SMnG0013576
h1SMcG0014863
h1SMcG0007806
h1SMcG0007803
h1SMcG0008564
h1SMnG0001804
h1SMcG0000798
h1SMnG0007035
h1SMnG0030385
h1SMnG0014953
h1SMcG0001722
h1SMnG0010769
h1SMcG0004059
h1SMnG0005552
h1SMnG0025476
h1SMcG0022244
h1SMcG0019963
h1SMnG0005755
h1SMcG0020732
h1SMnG0008037
h1SMcG0005863
h1SMnG0007950
h1SMnG0013500
h1SMnG0034462
h1SMcG0005419
h1SMcG0006227
h1SMnG0031626
h1SMcG0014978
h1SMnG0001341
h1SMcG0009395

### Module m31

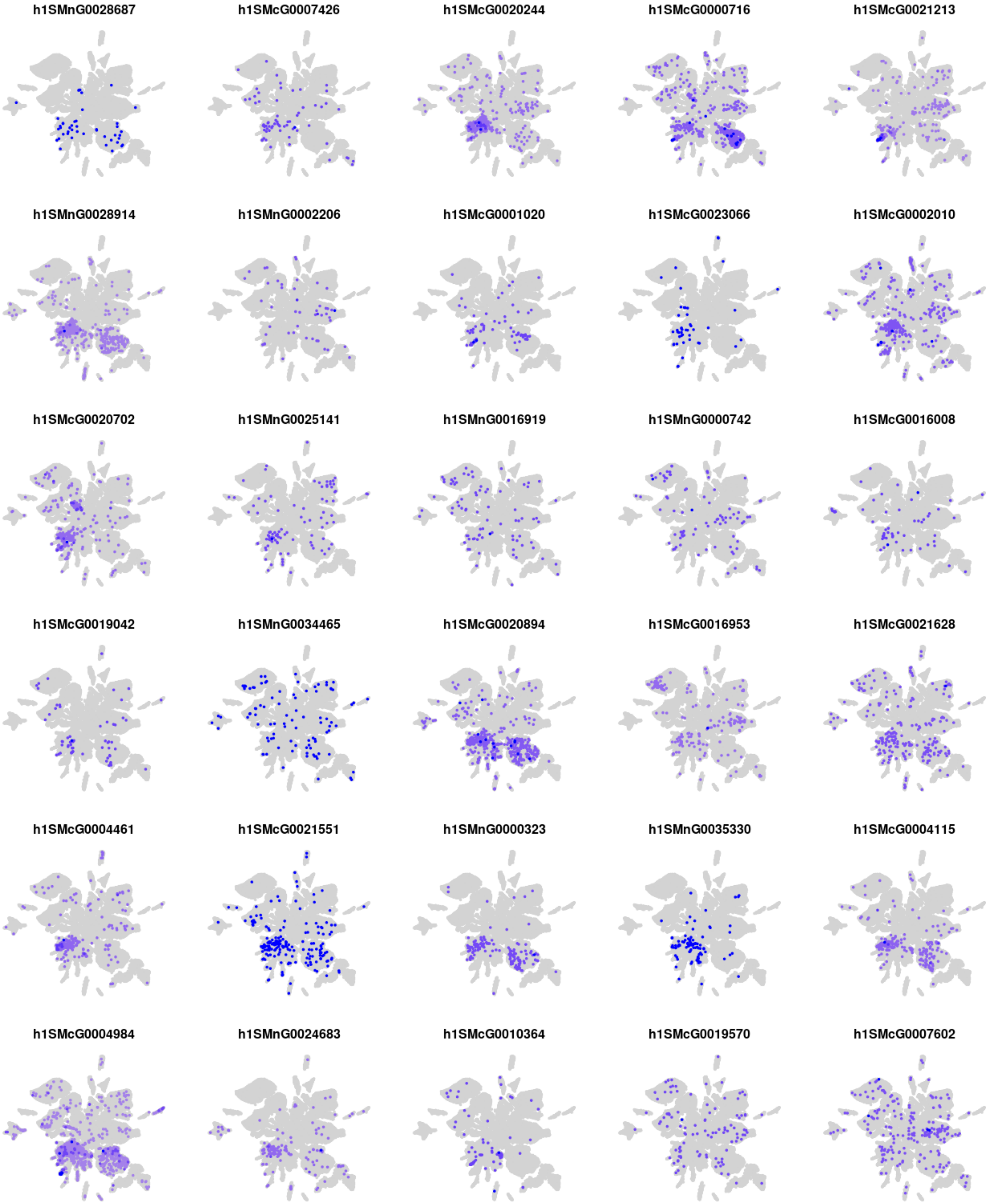
h1SMnG0028687
h1SMcG0007426
h1SMcG0020244
h1SMcG0000716
h1SMcG0021213
h1SMnG0028914
h1SMnG0002206
h1SMcG0001020
h1SMcG0023066
h1SMcG0002010
h1SMcG0020702
h1SMnG0025141
h1SMnG0016919
h1SMnG0000742
h1SMcG0016008
h1SMcG0019042
h1SMnG0034465
h1SMcG0020894
h1SMcG0016953
h1SMcG0021628
h1SMcG0004461
h1SMcG0021551
h1SMnG0000323
h1SMnG0035330
h1SMcG0004115
h1SMcG0004984
h1SMnG0024683
h1SMcG0010364
h1SMcG0019570
h1SMcG0007602

### Module m32

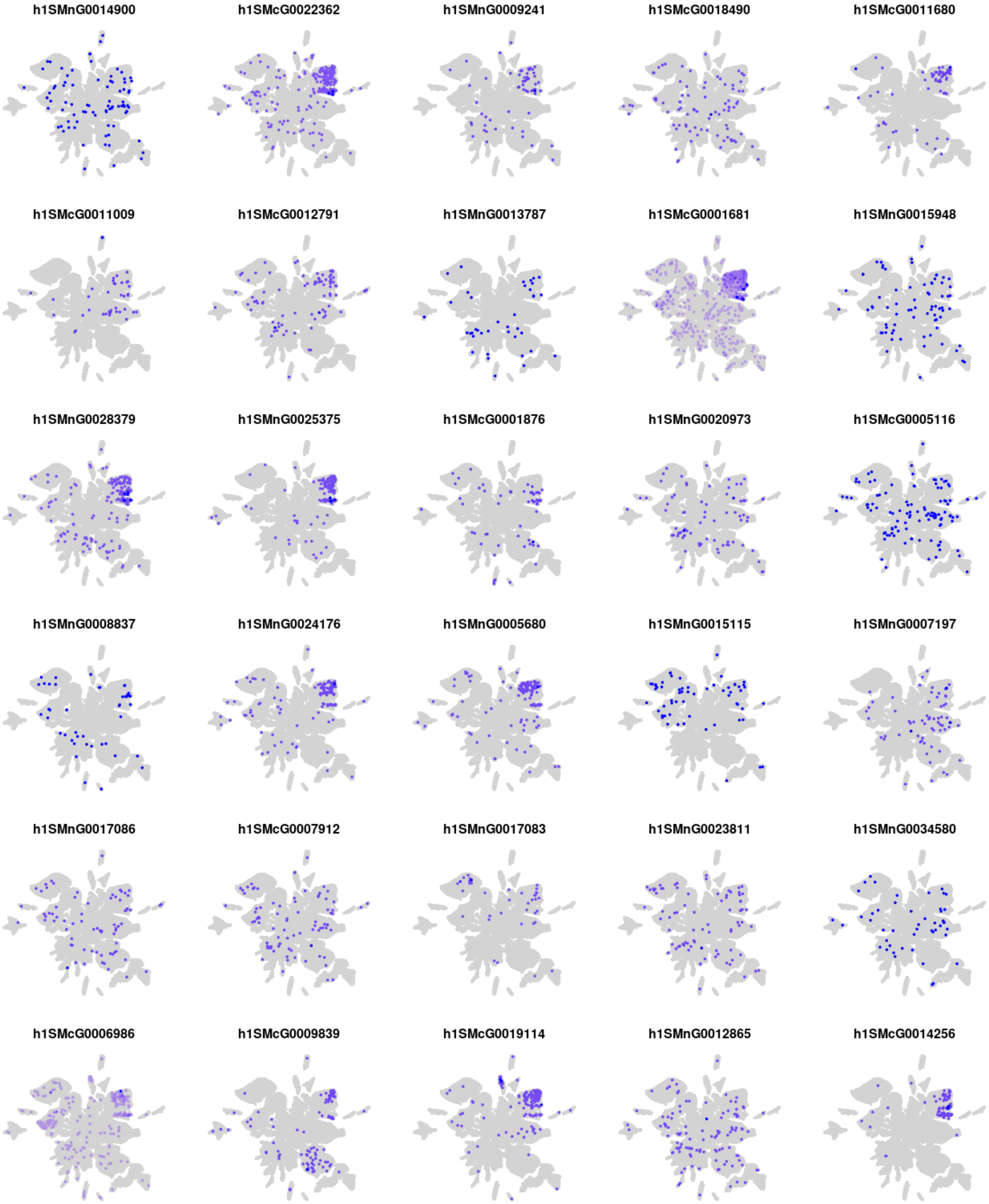

h1SMnG0014900
h1SMcG0022362
h1SMnG0009241
h1SMcG0018490
h1SMcG0011680
h1SMcG0011009
h1SMcG0012791
h1SMnG0013787
h1SMcG0001681
h1SMnG0015948
h1SMnG0028379
h1SMnG0025375
h1SMcG0001876
h1SMnG0020973
h1SMcG0005116
h1SMnG0008837
h1SMnG0024176
h1SMnG0005680
h1SMnG0015115
h1SMnG0007197
h1SMnG0017086
h1SMcG0007912
h1SMnG0017083
h1SMnG0023811
h1SMnG0034580
h1SMcG0006986
h1SMcG0009839
h1SMcG0019114
h1SMnG0012865
h1SMcG0014256

### Module m33

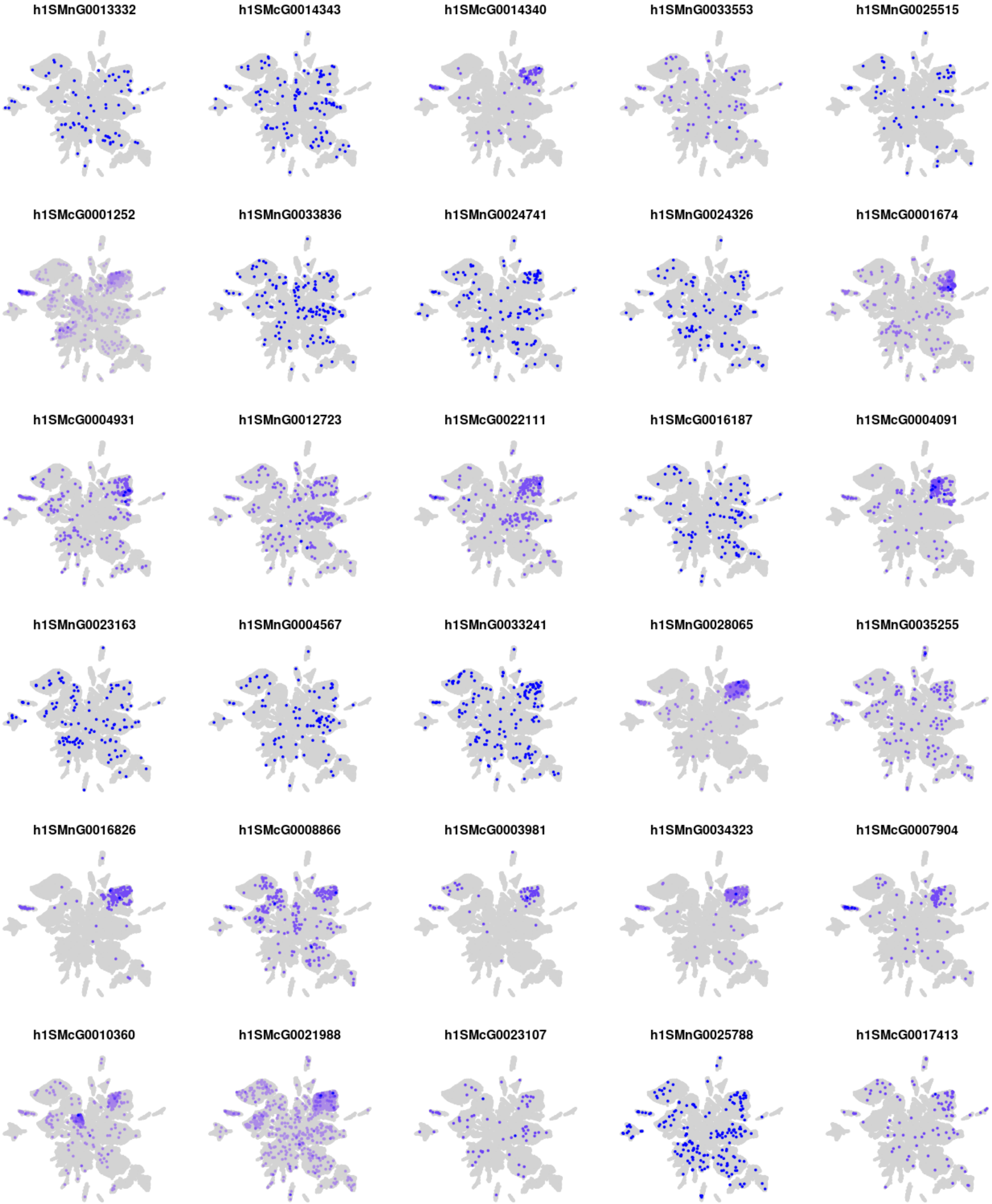
h1SMnG0013332
h1SMcG0014343
h1SMcG0014340
h1SMnG0033553
h1SMnG0025515
h1SMcG0001252
h1SMnG0033836
h1SMnG0024741
h1SMnG0024326
h1SMcG0001674
h1SMcG0004931
h1SMnG0012723
h1SMcG0022111
h1SMcG0016187
h1SMcG0004091
h1SMnG0023163
h1SMnG0004567
h1SMnG0033241
h1SMnG0028065
h1SMnG0035255
h1SMnG0016826
h1SMcG0008866
h1SMcG0003981
h1SMnG0034323
h1SMcG0007904
h1SMcG0010360
h1SMcG0021988
h1SMcG0023107
h1SMnG0025788
h1SMcG0017413

### Module m34

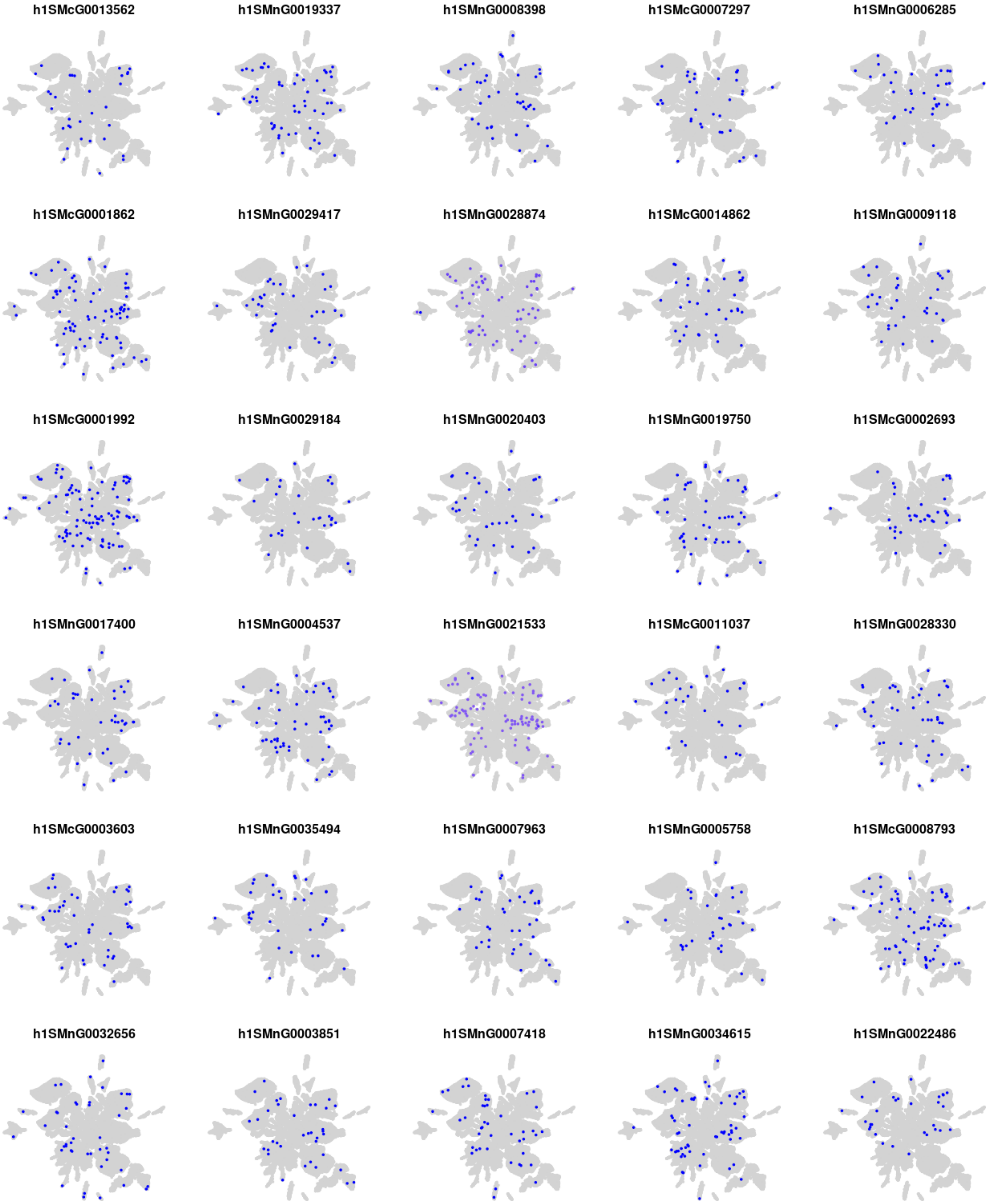
h1SMcG0013562
h1SMnG0019337
h1SMnG0008398
h1SMcG0007297
h1SMnG0006285
h1SMcG0001862
h1SMnG0029417
h1SMnG0028874
h1SMcG0014862
h1SMnG0009118
h1SMcG0001992
h1SMnG0029184
h1SMnG0020403
h1SMnG0019750
h1SMcG0002693
h1SMnG0017400
h1SMnG0004537
h1SMnG0021533
h1SMcG0011037
h1SMnG0028330
h1SMcG0003603
h1SMnG0035494
h1SMnG0007963
h1SMnG0005758
h1SMcG0008793
h1SMnG0032656
h1SMnG0003851
h1SMnG0007418
h1SMnG0034615
h1SMnG0022486

### Module m35

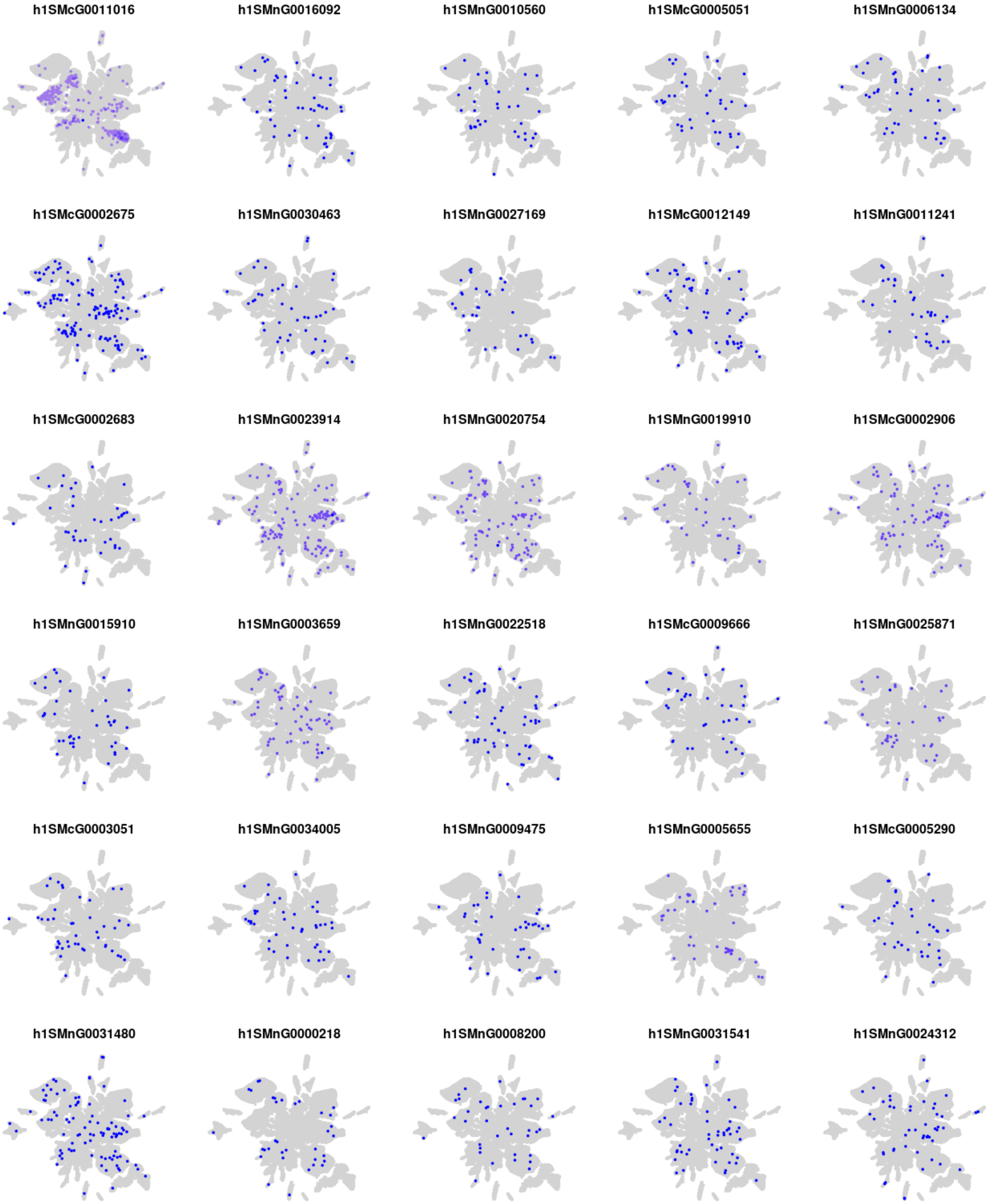

h1SMcG0011016
h1SMnG0016092
h1SMnG0010560
h1SMcG0005051
h1SMnG0006134
h1SMcG0002675
h1SMnG0030463
h1SMnG0027169
h1SMcG0012149
h1SMnG0011241
h1SMcG0002683
h1SMnG0023914
h1SMnG0020754
h1SMnG0019910
h1SMcG0002906
h1SMnG0015910
h1SMnG0003659
h1SMnG0022518
h1SMcG0009666
h1SMnG0025871
h1SMcG0003051
h1SMnG0034005
h1SMnG0009475
h1SMnG0005655
h1SMcG0005290
h1SMnG0031480
h1SMnG0000218
h1SMnG0008200
h1SMnG0031541
h1SMnG0024312

### Module m36

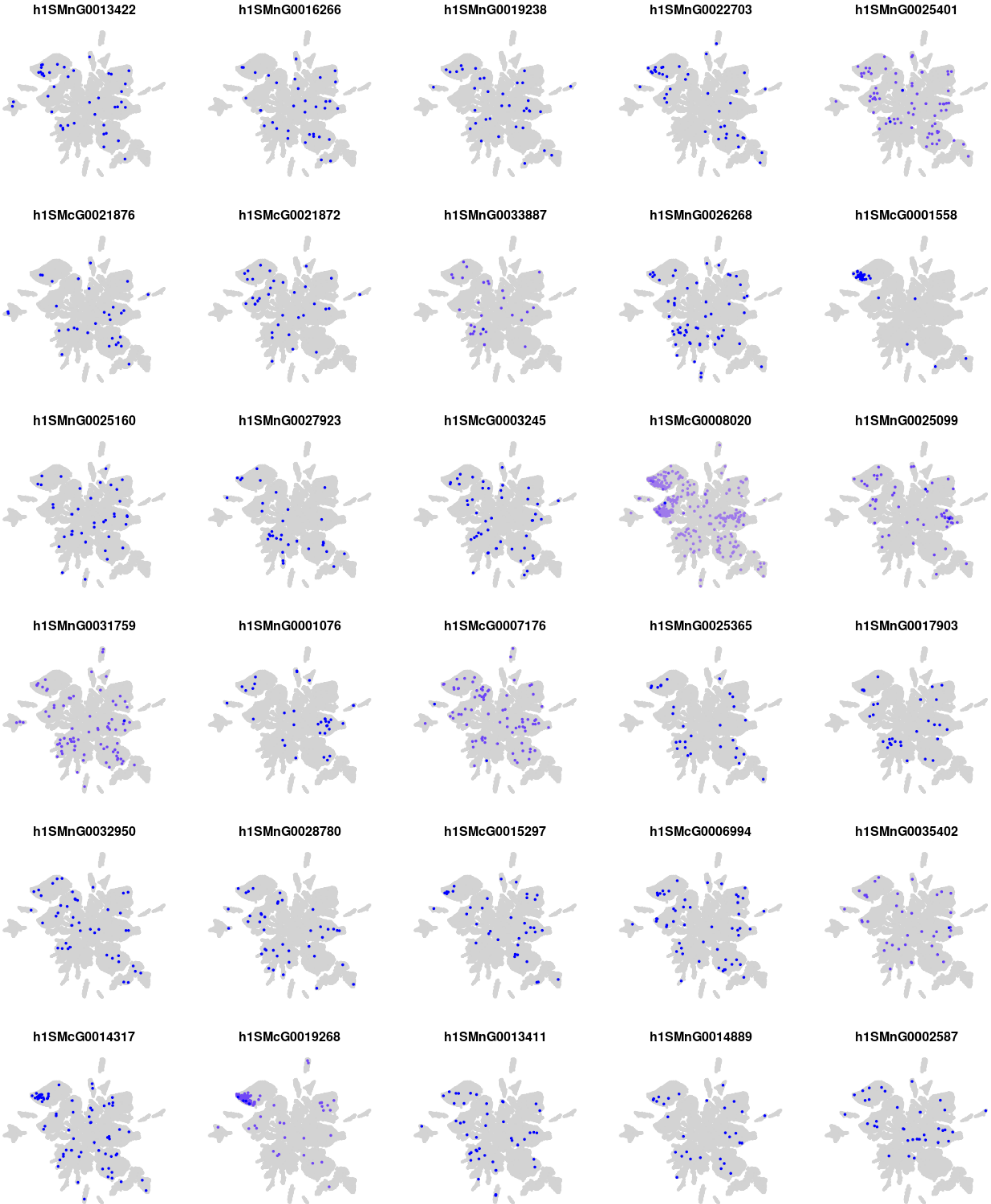

h1SMnG0013422
h1SMnG0016266
h1SMnG0019238
h1SMnG0022703
h1SMnG0025401
h1SMcG0021876
h1SMcG0021872
h1SMnG0033887
h1SMnG0026268
h1SMcG0001558
h1SMnG0025160
h1SMnG0027923
h1SMcG0003245
h1SMcG0008020
h1SMnG0025099
h1SMnG0031759
h1SMnG0001076
h1SMcG0007176
h1SMnG0025365
h1SMnG0017903
h1SMnG0032950
h1SMnG0028780
h1SMcG0015297
h1SMcG0006994
h1SMnG0035402
h1SMcG0014317
h1SMcG0019268
h1SMnG0013411
h1SMnG0014889
h1SMnG0002587

#### Module m37

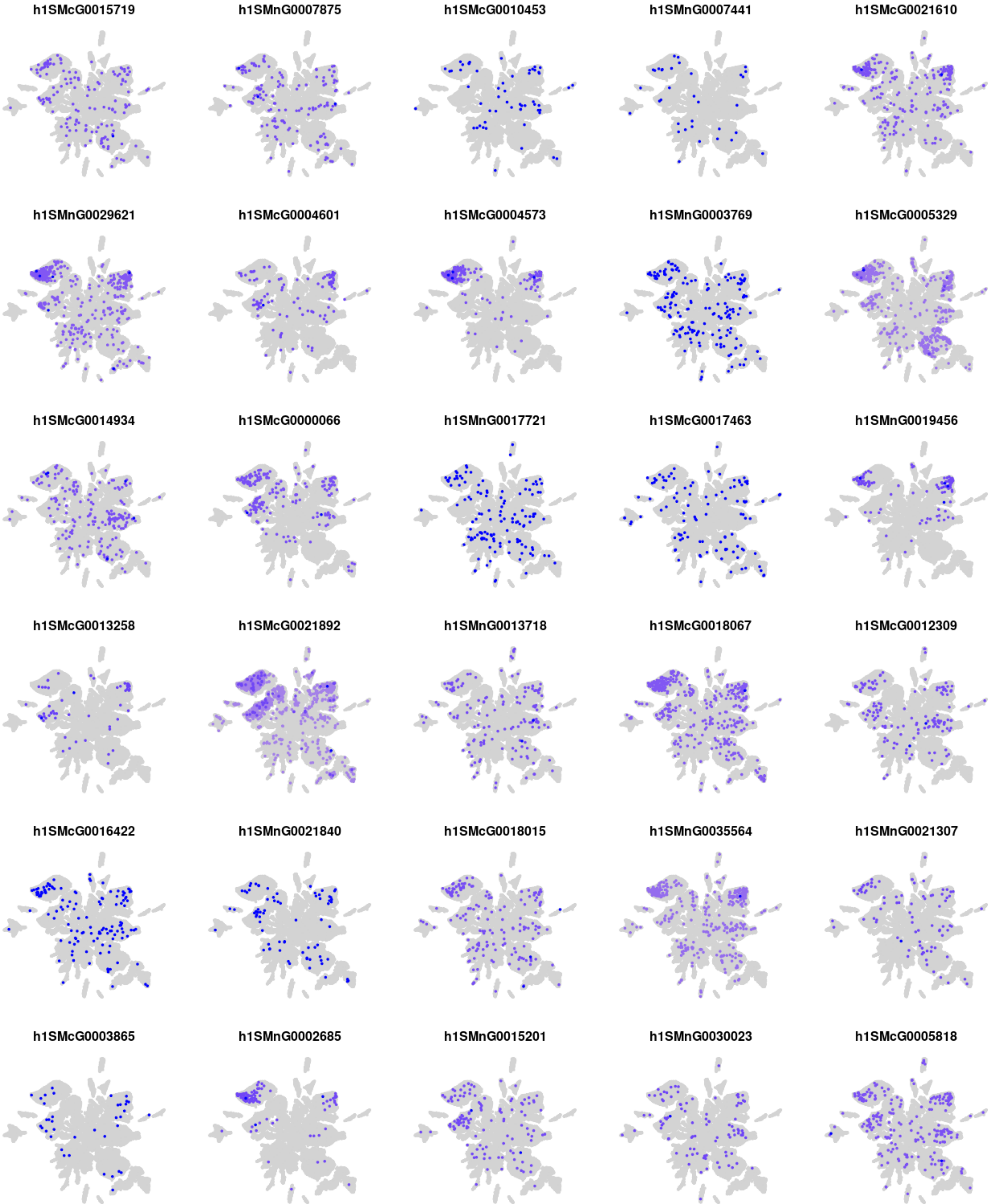
h1SMcG0015719
h1SMnG0007875
h1SMcG0010453
h1SMnG0007441
h1SMcG0021610
h1SMnG0029621
h1SMcG0004601
h1SMcG0004573
h1SMnG0003769
h1SMcG0005329
h1SMcG0014934
h1SMcG0000066
h1SMnG0017721
h1SMcG0017463
h1SMnG0019456
h1SMcG0013258
h1SMcG0021892
h1SMnG0013718
h1SMcG0018067
h1SMcG0012309
h1SMcG0016422
h1SMnG0021840
h1SMcG0018015
h1SMnG0035564
h1SMnG0021307
h1SMcG0003865
h1SMnG0002685
h1SMnG0015201
h1SMnG0030023
h1SMcG0005818

### Module m38

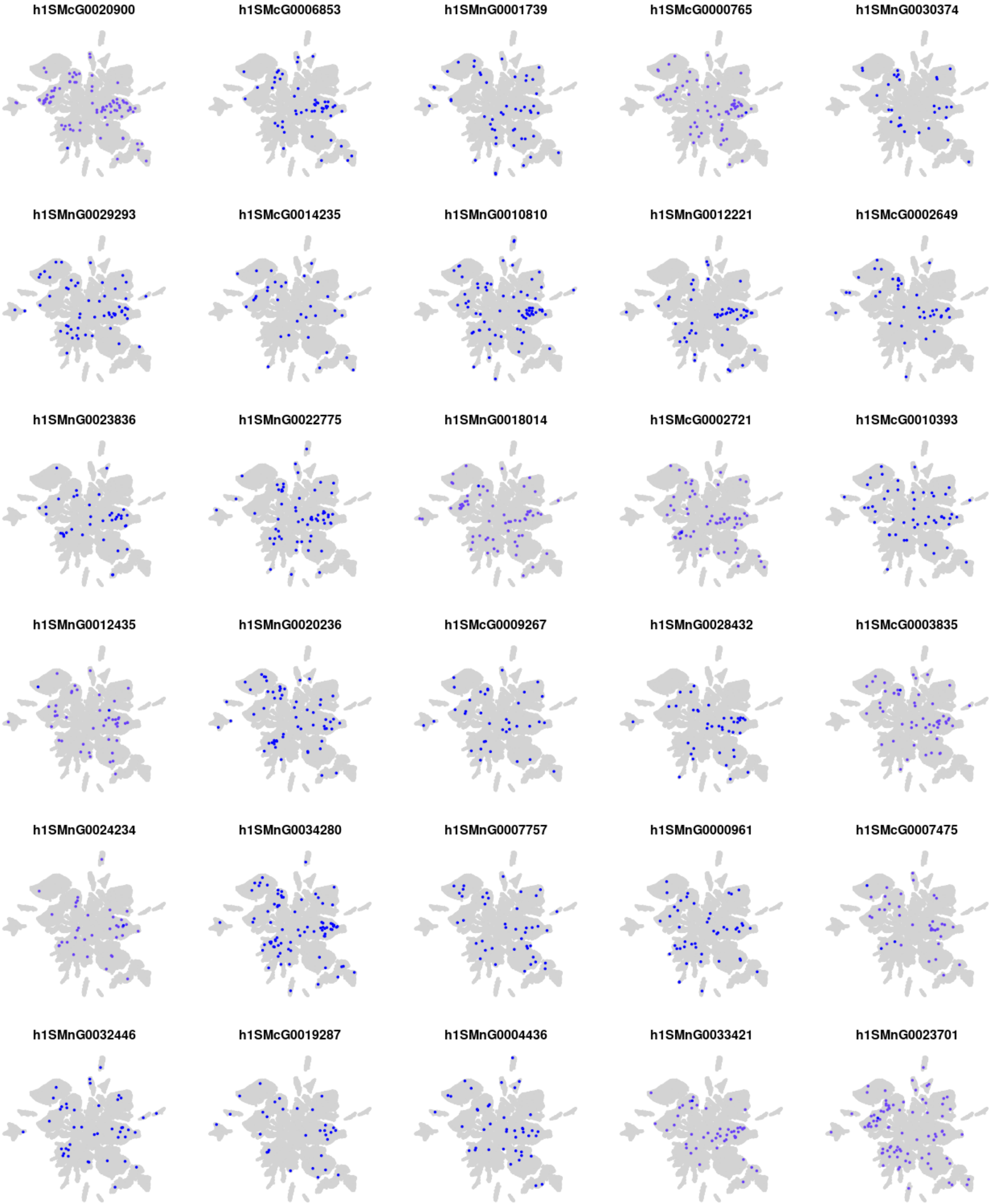
h1SMcG0020900
h1SMcG0006853
h1SMnG0001739
h1SMcG0000765
h1SMnG0030374
h1SMnG0029293
h1SMcG0014235
h1SMnG0010810
h1SMnG0012221
h1SMcG0002649
h1SMnG0023836
h1SMnG0022775
h1SMnG0018014
h1SMcG0002721
h1SMcG0010393
h1SMnG0012435
h1SMnG0020236
h1SMcG0009267
h1SMnG0028432
h1SMcG0003835
h1SMnG0024234
h1SMnG0034280
h1SMnG0007757
h1SMnG0000961
h1SMcG0007475
h1SMnG0032446
h1SMcG0019287
h1SMnG0004436
h1SMnG0033421
h1SMnG0023701

### Module m39

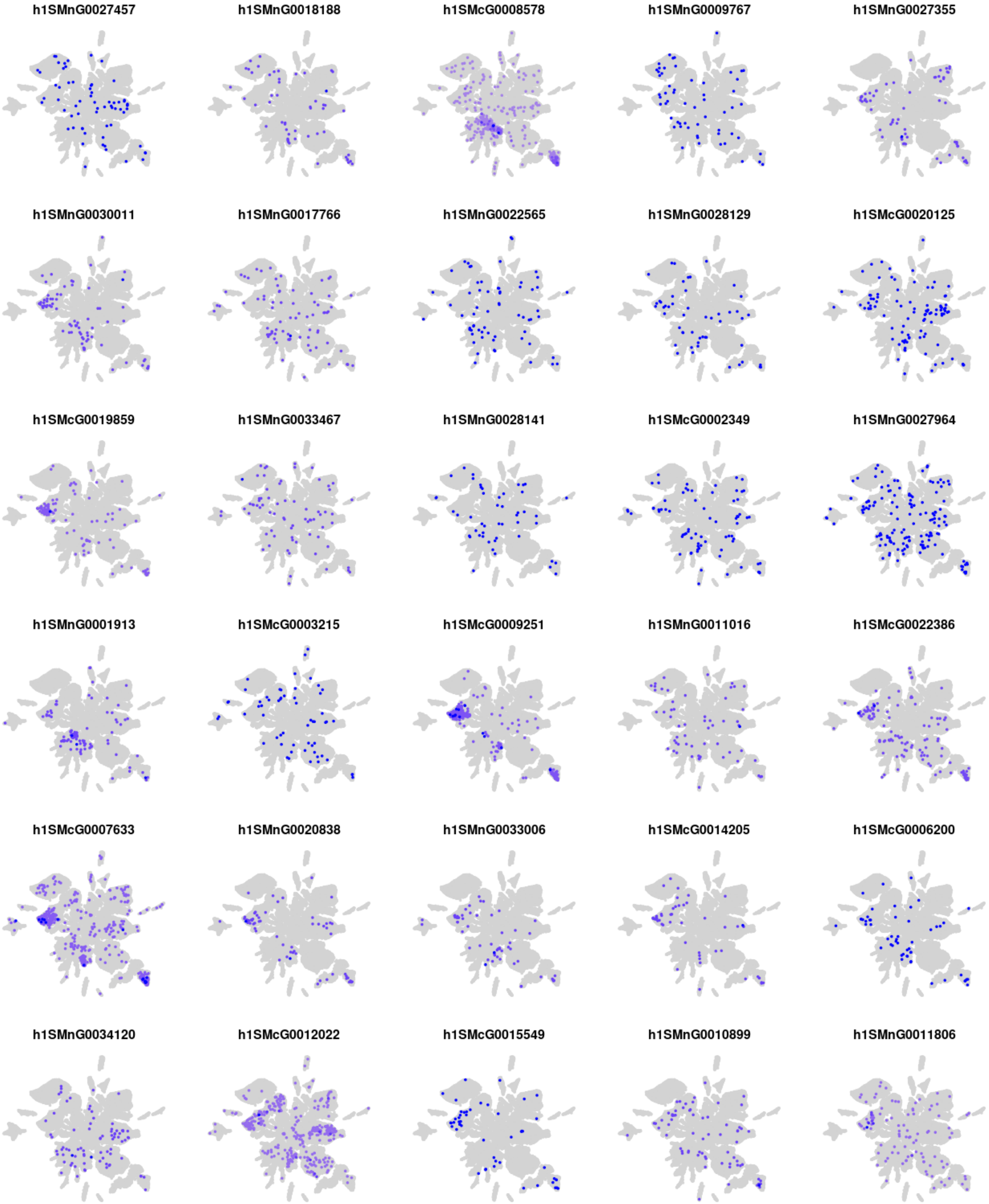
h1SMnG0027457
h1SMnG0018188
h1SMcG0008578
h1SMnG0009767
h1SMnG0027355
h1SMnG0030011
h1SMnG0017766
h1SMnG0022565
h1SMnG0028129
h1SMcG0020125
h1SMcG0019859
h1SMnG0033467
h1SMnG0028141
h1SMcG0002349
h1SMnG0027964
h1SMnG0001913
h1SMcG0003215
h1SMcG0009251
h1SMnG0011016
h1SMcG0022386
h1SMcG0007633
h1SMnG0020838
h1SMnG0033006
h1SMcG0014205
h1SMcG0006200
h1SMnG0034120
h1SMcG0012022
h1SMcG0015549
h1SMnG0010899
h1SMnG0011806

### Module m40

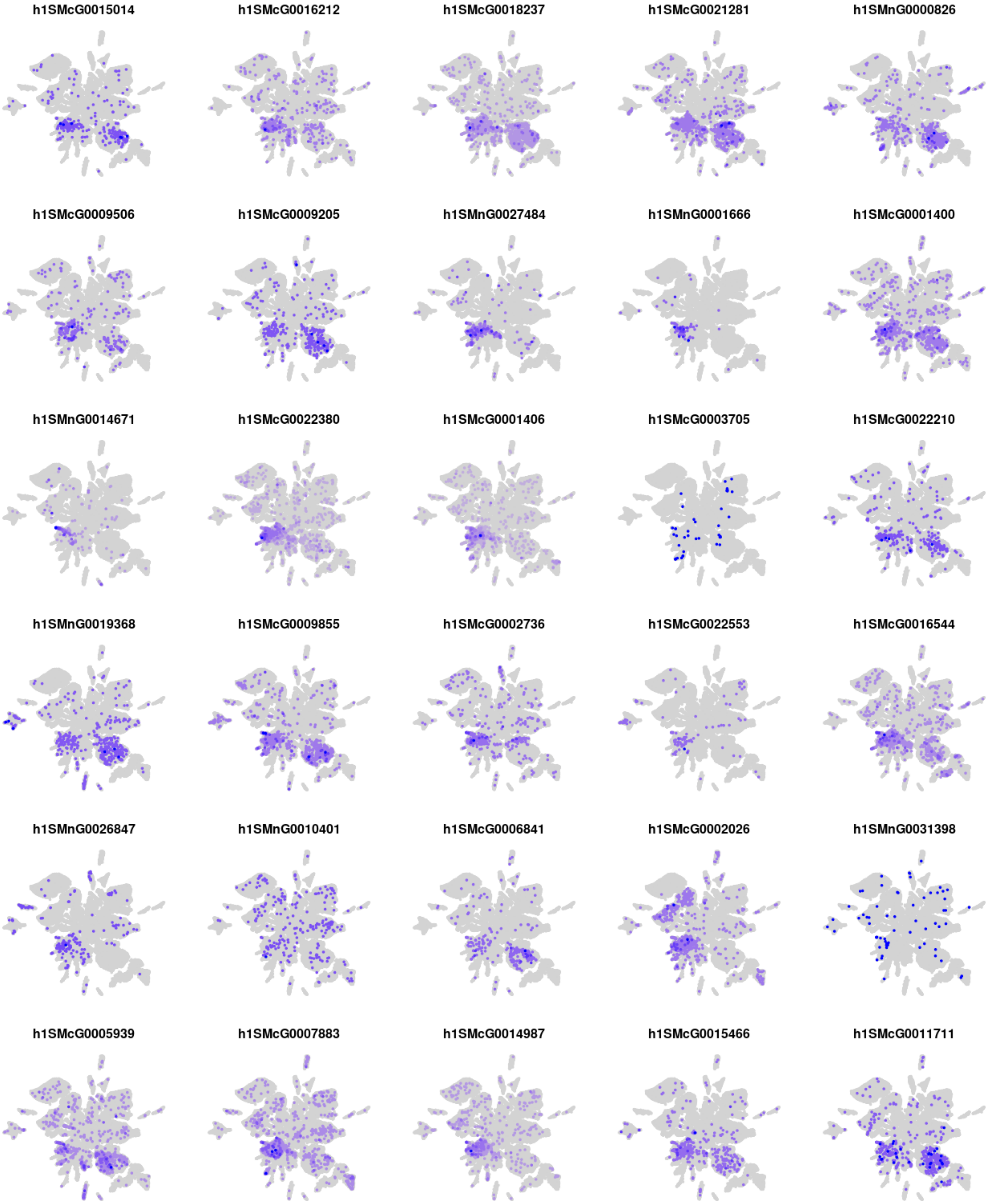
h1SMcG0015014
h1SMcG0016212
h1SMcG0018237
h1SMcG0021281
h1SMnG0000826
h1SMcG0009506
h1SMcG0009205
h1SMnG0027484
h1SMnG0001666
h1SMcG0001400
h1SMnG0014671
h1SMcG0022380
h1SMcG0001406
h1SMcG0003705
h1SMcG0022210
h1SMnG0019368
h1SMcG0009855
h1SMcG0002736
h1SMcG0022553
h1SMcG0016544
h1SMnG0026847
h1SMnG0010401
h1SMcG0006841
h1SMcG0002026
h1SMnG0031398
h1SMcG0005939
h1SMcG0007883
h1SMcG0014987
h1SMcG0015466
h1SMcG0011711

#### Module m41

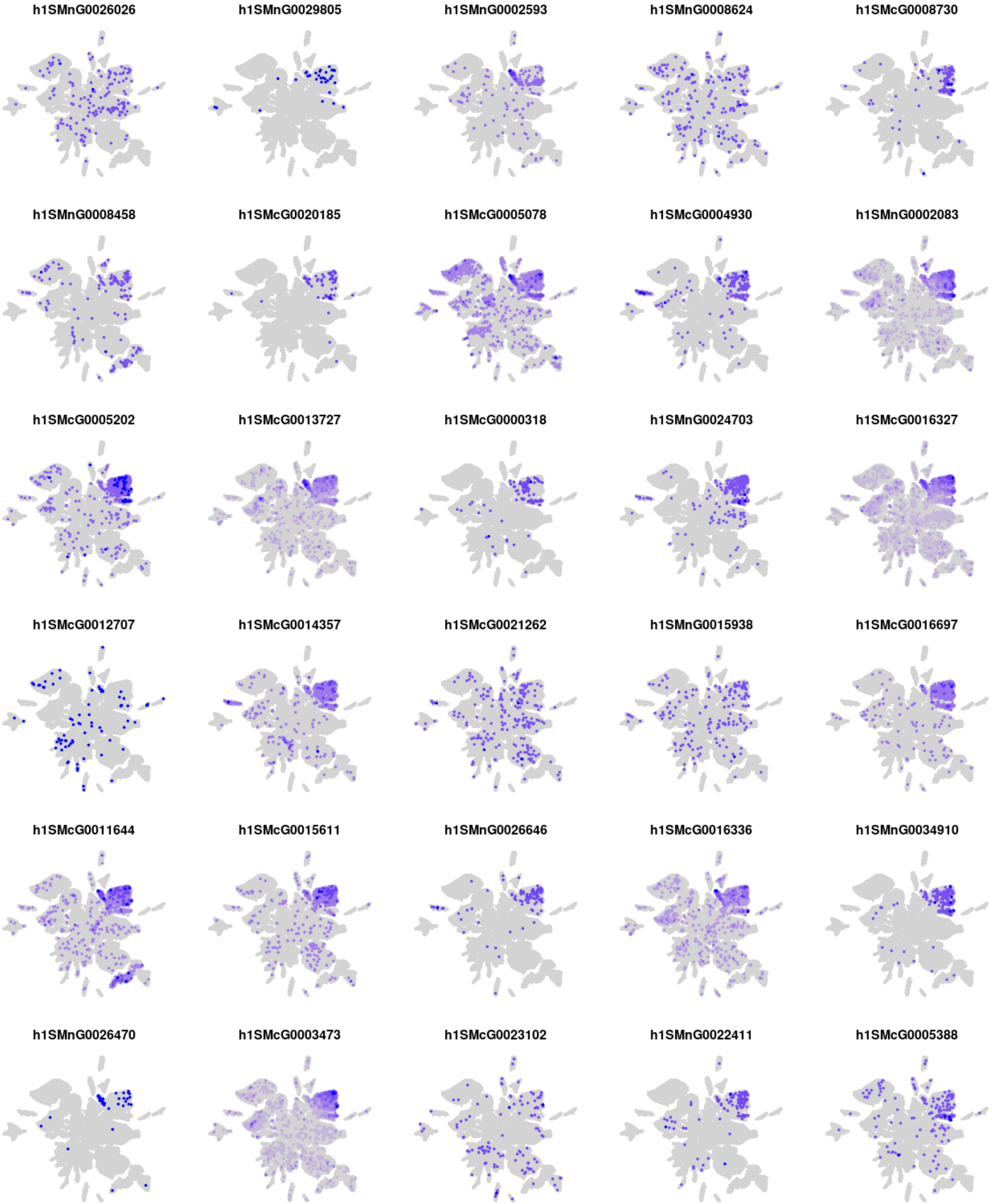
h1SMnG0026026
h1SMnG0029805
h1SMnG0002593
h1SMnG0008624
h1SMcG0008730
h1SMnG0008458
h1SMcG0020185
h1SMcG0005078
h1SMcG0004930
h1SMnG0002083
h1SMcG0005202
h1SMcG0013727
h1SMcG0000318
h1SMnG0024703
h1SMcG0016327
h1SMcG0012707
h1SMcG0014357
h1SMcG0021262
h1SMnG0015938
h1SMcG0016697
h1SMcG0011644
h1SMcG0015611
h1SMnG0026646
h1SMcG0016336
h1SMnG0034910
h1SMnG0026470
h1SMcG0003473
h1SMcG0023102
h1SMnG0022411
h1SMcG0005388

### Module m42

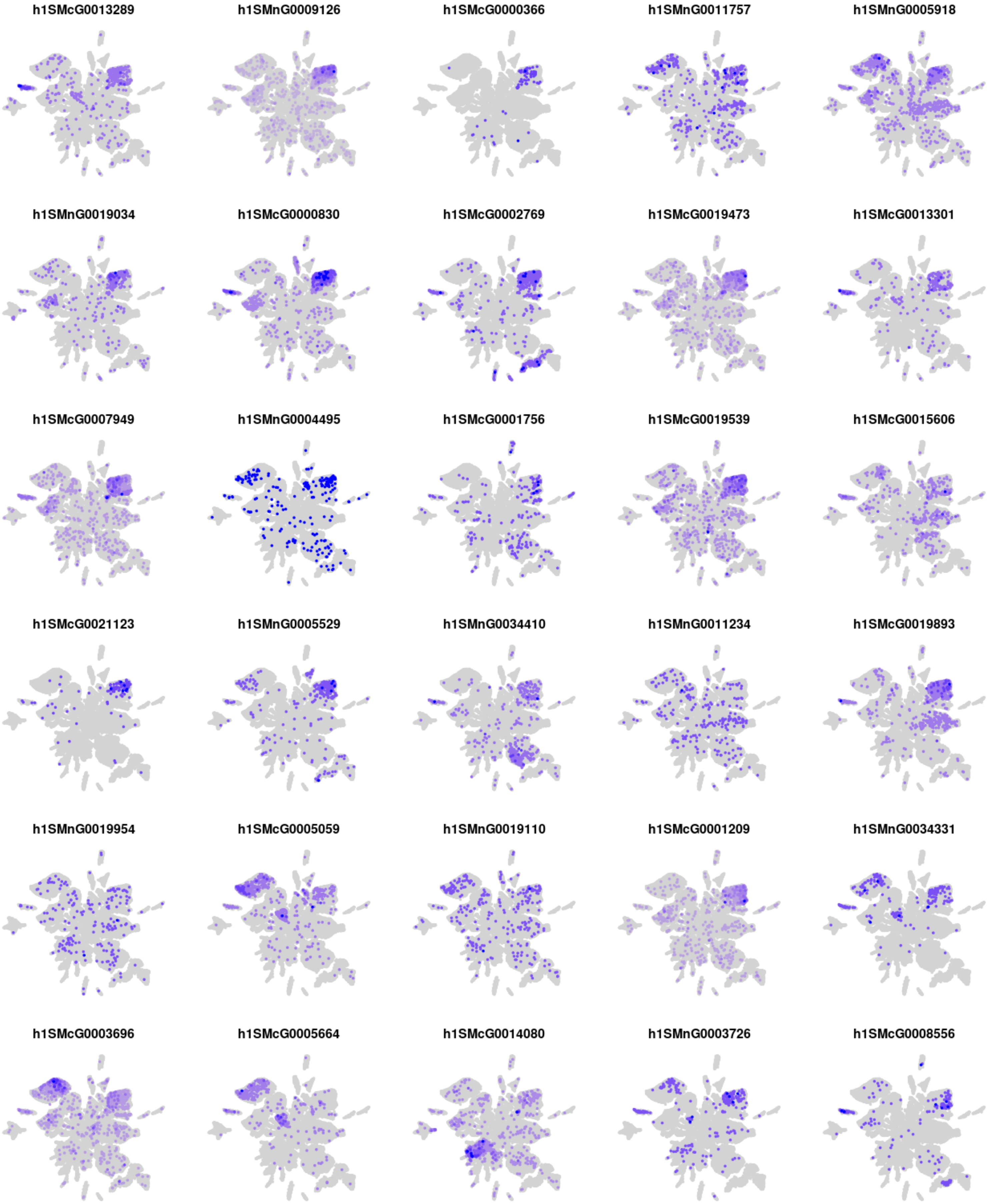
h1SMcG0013289
h1SMnG0009126
h1SMcG0000366
h1SMnG0011757
h1SMnG0005918
h1SMnG0019034
h1SMcG0000830
h1SMcG0002769
h1SMcG0019473
h1SMcG0013301
h1SMcG0007949
h1SMnG0004495
h1SMcG0001756
h1SMcG0019539
h1SMcG0015606
h1SMcG0021123
h1SMnG0005529
h1SMnG0034410
h1SMnG0011234
h1SMcG0019893
h1SMnG0019954
h1SMcG0005059
h1SMnG0019110
h1SMcG0001209
h1SMnG0034331
h1SMcG0003696
h1SMcG0005664
h1SMcG0014080
h1SMnG0003726
h1SMcG0008556

### Module m43

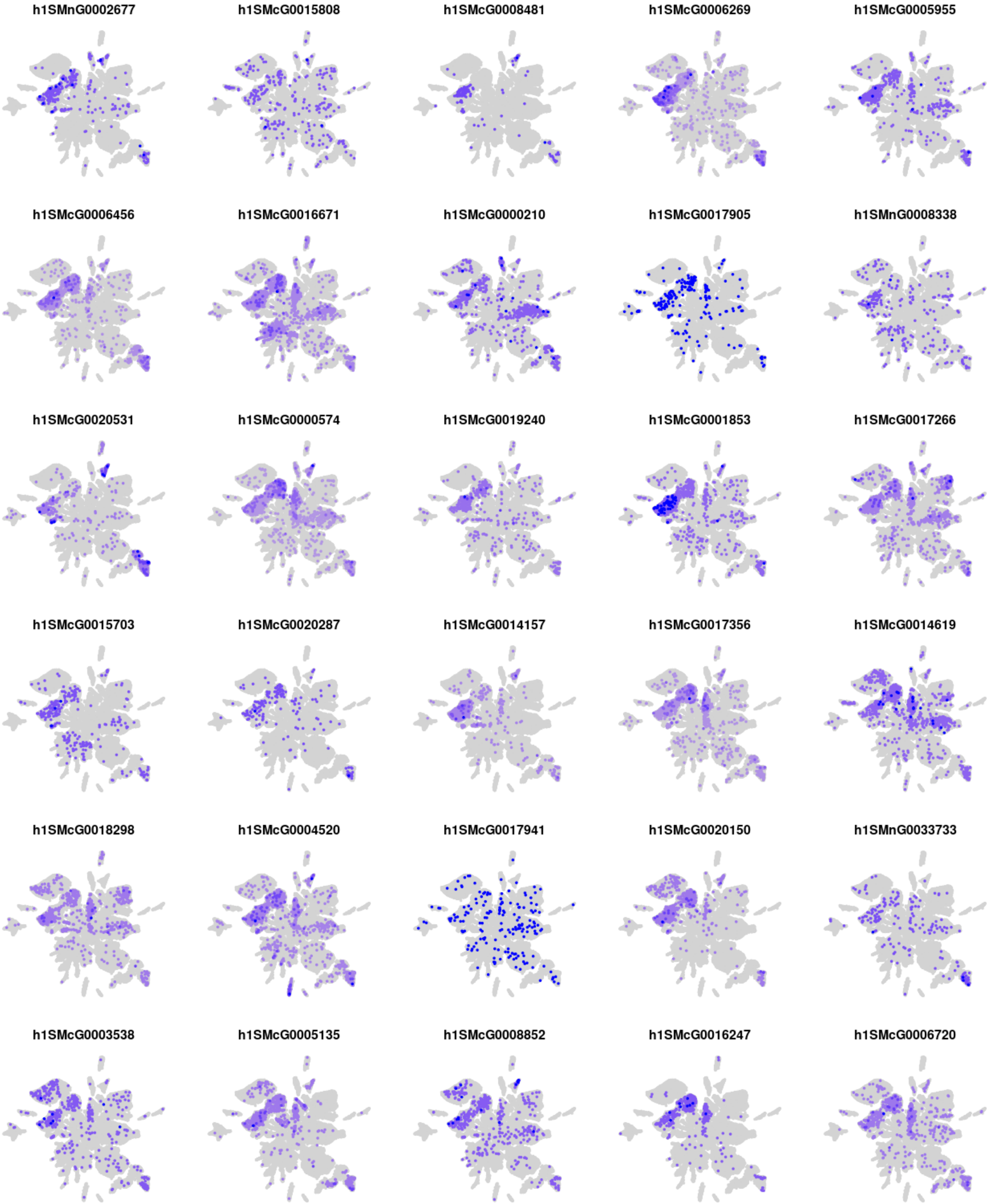
h1SMnG0002677
h1SMcG0015808
h1SMcG0008481
h1SMcG0006269
h1SMcG0005955
h1SMcG0006456
h1SMcG0016671
h1SMcG0000210
h1SMcG0017905
h1SMnG0008338
h1SMcG0020531
h1SMcG0000574
h1SMcG0019240
h1SMcG0001853
h1SMcG0017266
h1SMcG0015703
h1SMcG0020287
h1SMcG0014157
h1SMcG0017356
h1SMcG0014619
h1SMcG0018298
h1SMcG0004520
h1SMcG0017941
h1SMcG0020150
h1SMnG0033733
h1SMcG0003538
h1SMcG0005135
h1SMcG0008852
h1SMcG0016247
h1SMcG0006720

### Module m44

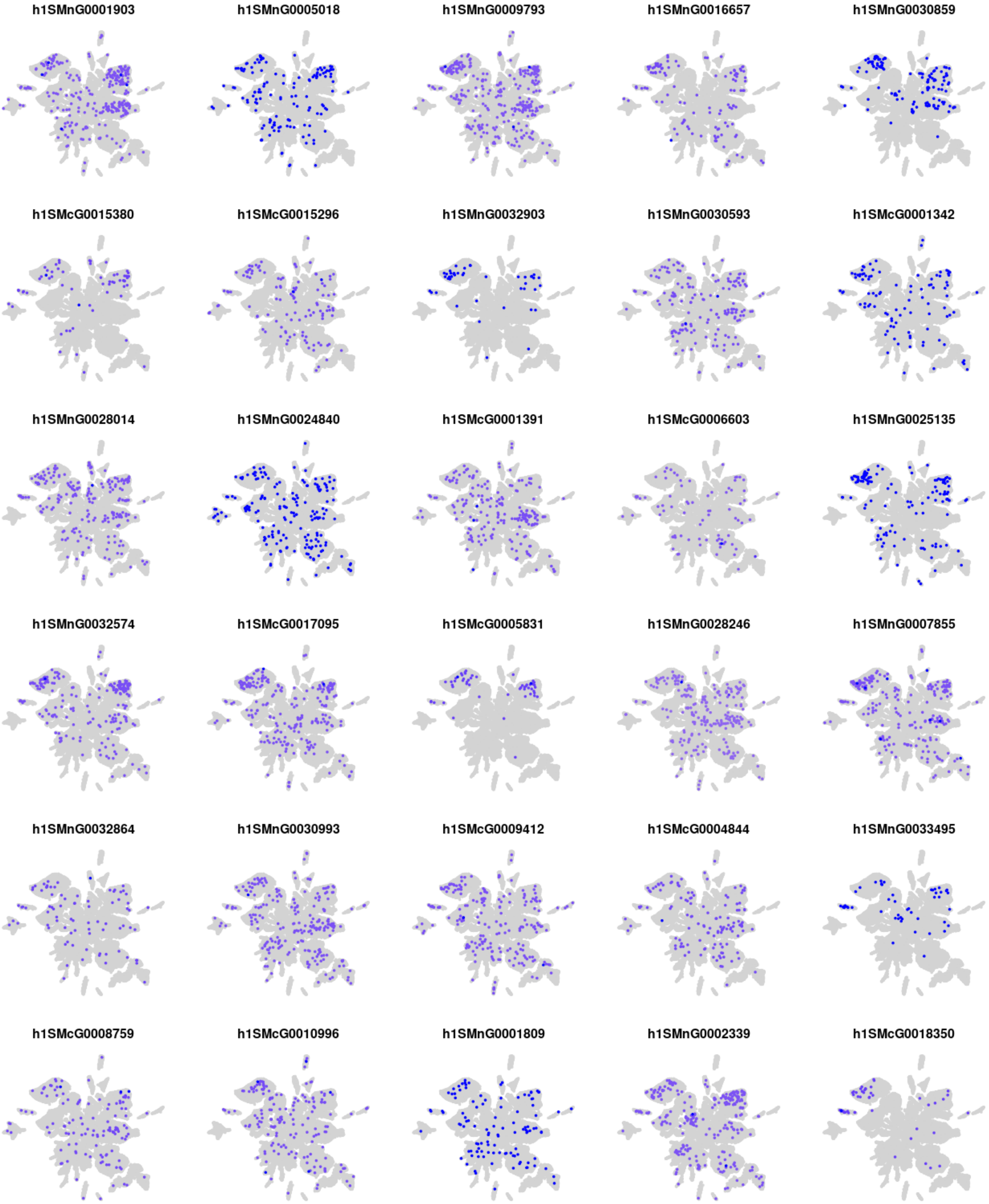
h1SMnG0001903
h1SMnG0005018
h1SMnG0009793
h1SMnG0016657
h1SMnG0030859
h1SMcG0015380
h1SMcG0015296
h1SMnG0032903
h1SMnG0030593
h1SMcG0001342
h1SMnG0028014
h1SMnG0024840
h1SMcG0001391
h1SMcG0006603
h1SMnG0025135
h1SMnG0032574
h1SMcG0017095
h1SMcG0005831
h1SMnG0028246
h1SMnG0007855
h1SMnG0032864
h1SMnG0030993
h1SMcG0009412
h1SMcG0004844
h1SMnG0033495
h1SMcG0008759
h1SMcG0010996
h1SMnG0001809
h1SMnG0002339
h1SMcG0018350

### Module m45

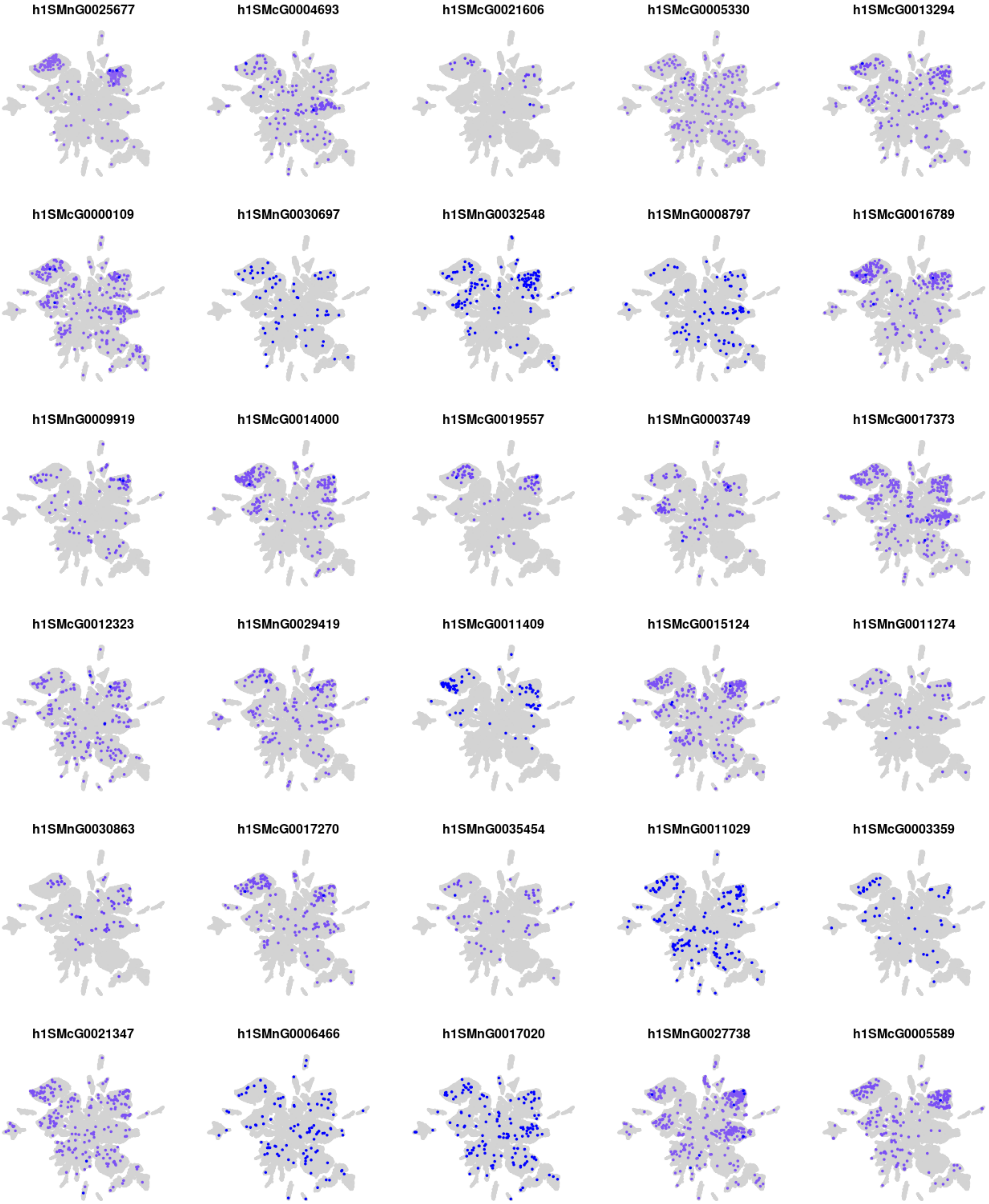
h1SMnG0025677
h1SMcG0004693
h1SMcG0021606
h1SMcG0005330
h1SMcG0013294
h1SMcG0000109
h1SMnG0030697
h1SMnG0032548
h1SMnG0008797
h1SMcG0016789
h1SMnG0009919
h1SMcG0014000
h1SMcG0019557
h1SMnG0003749
h1SMcG0017373
h1SMcG0012323
h1SMnG0029419
h1SMcG0011409
h1SMcG0015124
h1SMnG0011274
h1SMnG0030863
h1SMcG0017270
h1SMnG0035454
h1SMnG0011029
h1SMcG0003359
h1SMcG0021347
h1SMnG0006466
h1SMnG0017020
h1SMnG0027738
h1SMcG0005589

### Module m46

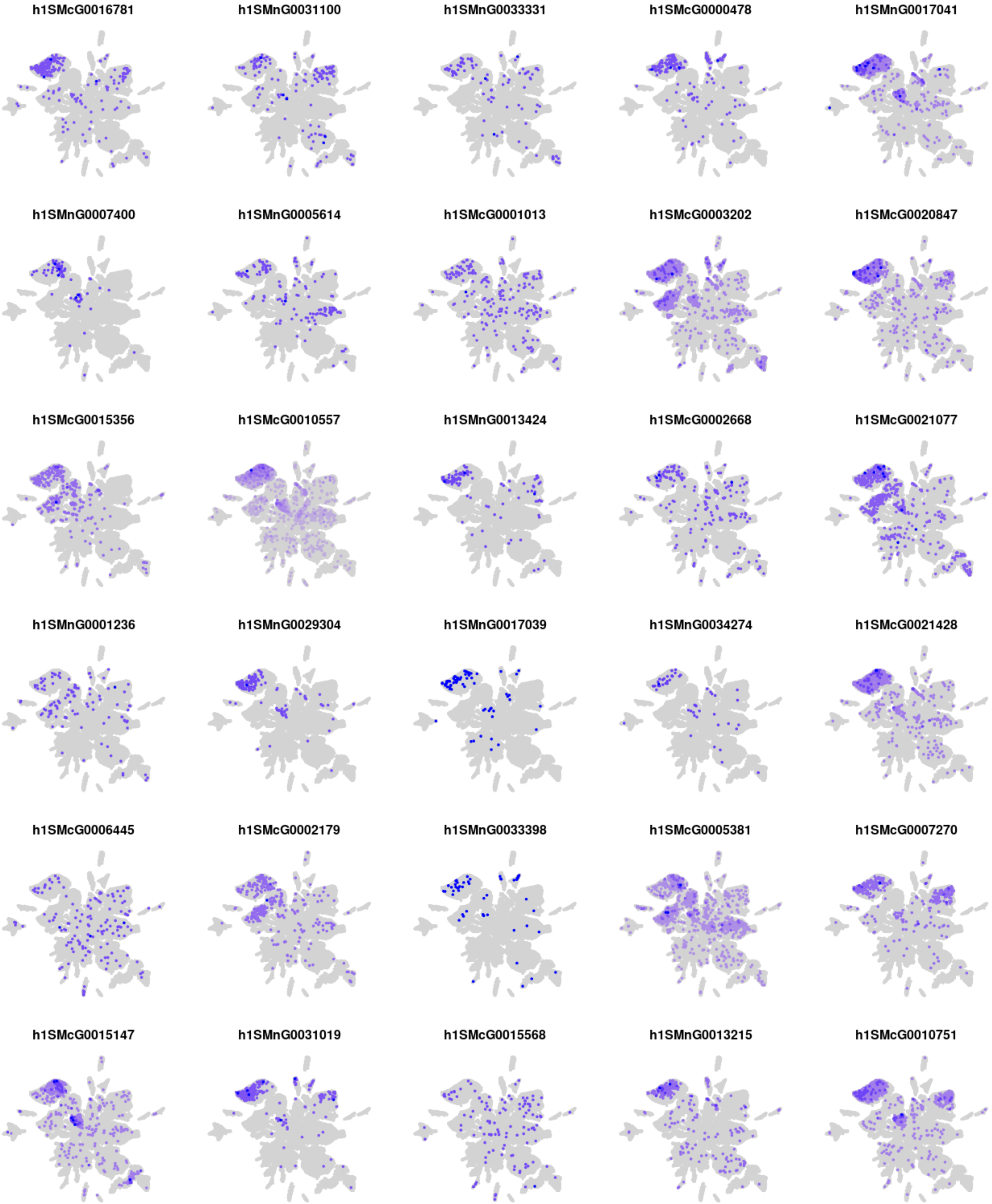
h1SMcG0016781
h1SMnG0031100
h1SMnG0033331
h1SMcG0000478
h1SMnG0017041
h1SMnG0007400
h1SMnG0005614
h1SMcG0001013
h1SMcG0003202
h1SMcG0020847
h1SMcG0015356
h1SMcG0010557
h1SMnG0013424
h1SMcG0002668
h1SMcG0021077
h1SMnG0001236
h1SMnG0029304
h1SMnG0017039
h1SMnG0034274
h1SMcG0021428
h1SMcG0006445
h1SMcG0002179
h1SMnG0033398
h1SMcG0005381
h1SMcG0007270
h1SMcG0015147
h1SMnG0031019
h1SMcG0015568
h1SMnG0013215
h1SMcG0010751

### Module m47

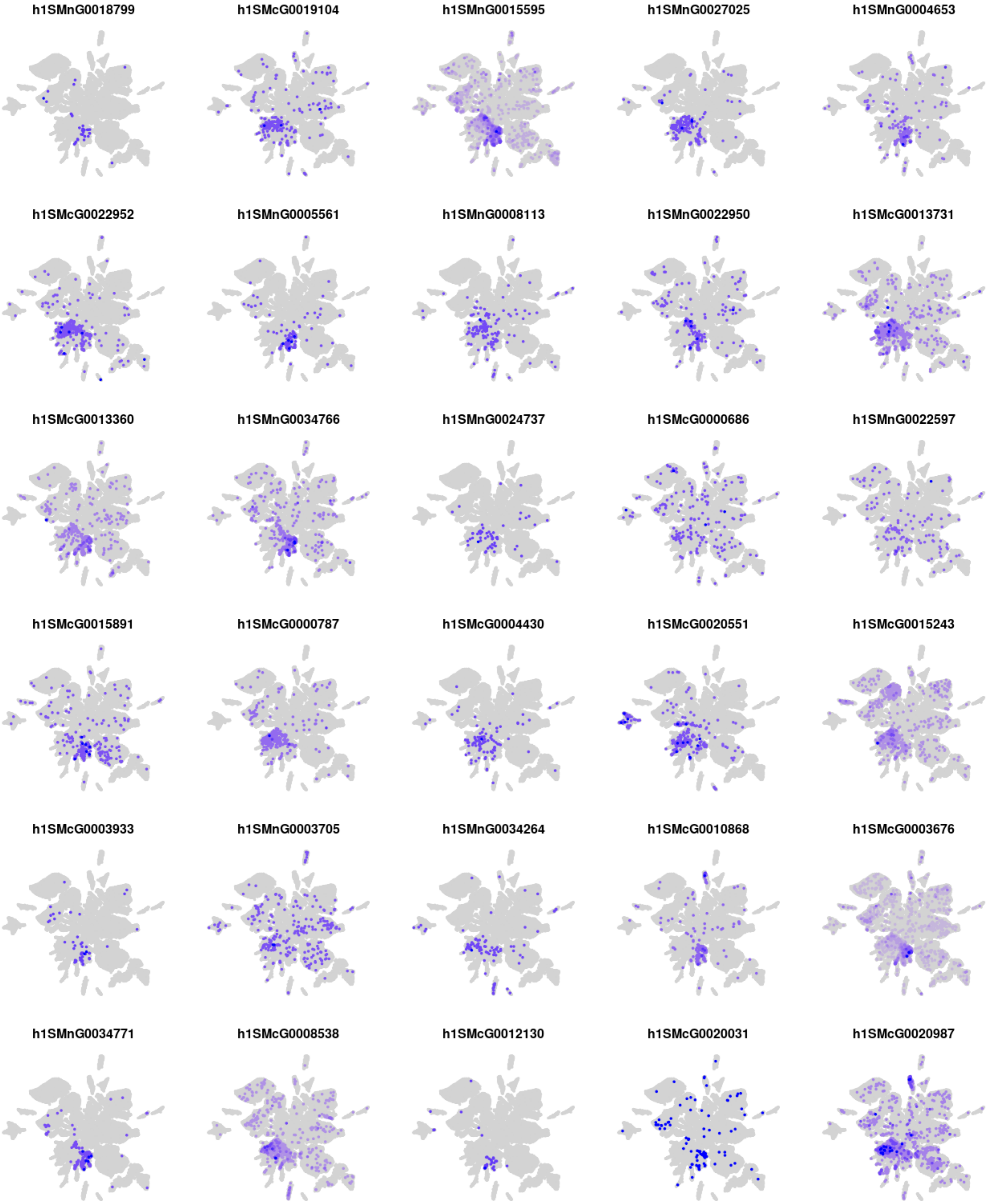
h1SMnG0018799
h1SMcG0019104
h1SMnG0015595
h1SMnG0027025
h1SMnG0004653
h1SMcG0022952
h1SMnG0005561
h1SMnG0008113
h1SMnG0022950
h1SMcG0013731
h1SMcG0013360
h1SMnG0034766
h1SMnG0024737
h1SMcG0000686
h1SMnG0022597
h1SMcG0015891
h1SMcG0000787
h1SMcG0004430
h1SMcG0020551
h1SMcG0015243
h1SMcG0003933
h1SMnG0003705
h1SMnG0034264
h1SMcG0010868
h1SMcG0003676
h1SMnG0034771
h1SMcG0008538
h1SMcG0012130
h1SMcG0020031
h1SMcG0020987

### Module m48

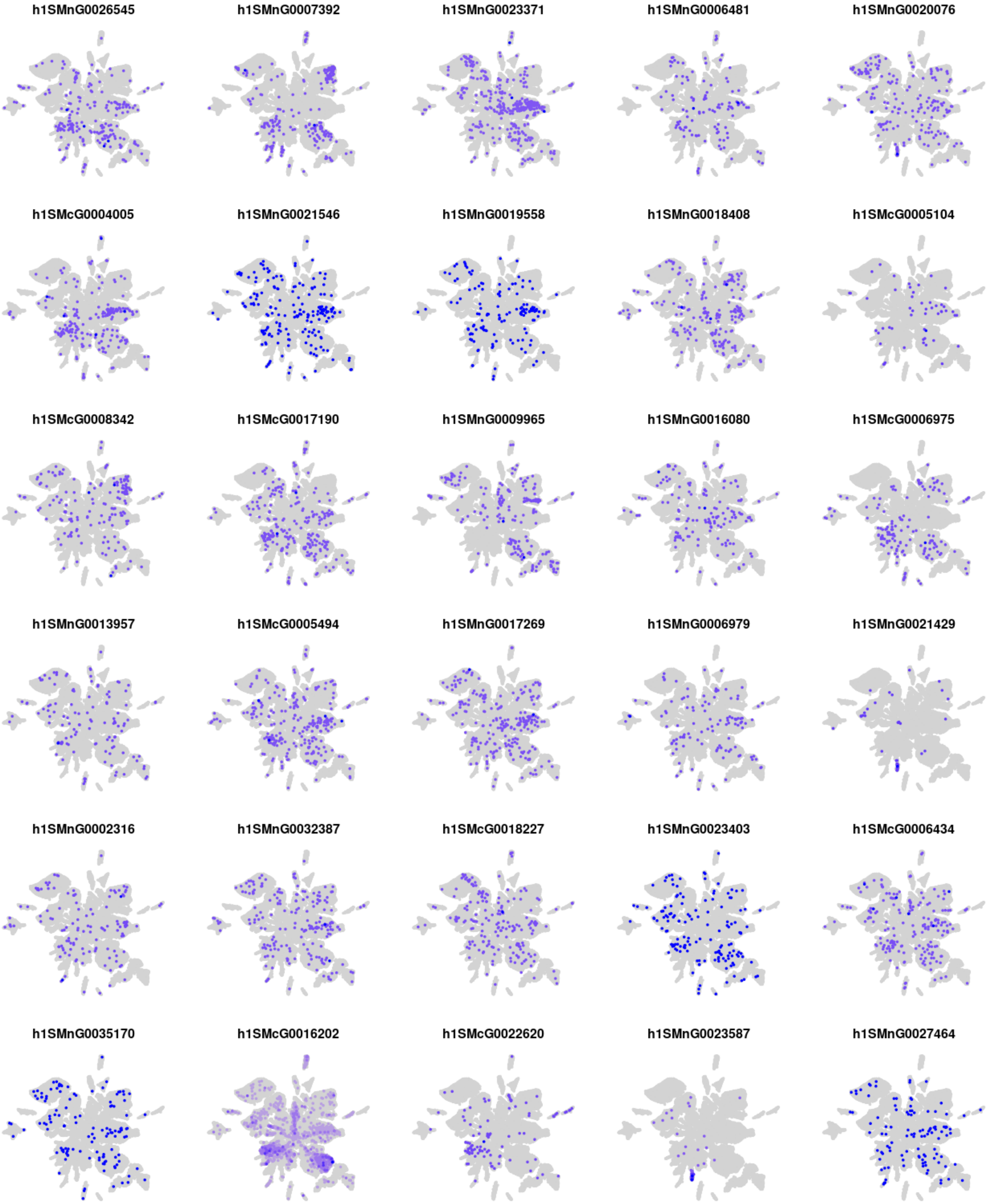
h1SMnG0026545
h1SMnG0007392
h1SMnG0023371
h1SMnG0006481
h1SMnG0020076
h1SMcG0004005
h1SMnG0021546
h1SMnG0019558
h1SMnG0018408
h1SMcG0005104
h1SMcG0008342
h1SMcG0017190
h1SMnG0009965
h1SMnG0016080
h1SMcG0006975
h1SMnG0013957
h1SMcG0005494
h1SMnG0017269
h1SMnG0006979
h1SMnG0021429
h1SMnG0002316
h1SMnG0032387
h1SMcG0018227
h1SMnG0023403
h1SMcG0006434
h1SMnG0035170
h1SMcG0016202
h1SMcG0022620
h1SMnG0023587
h1SMnG0027464

### Module m49

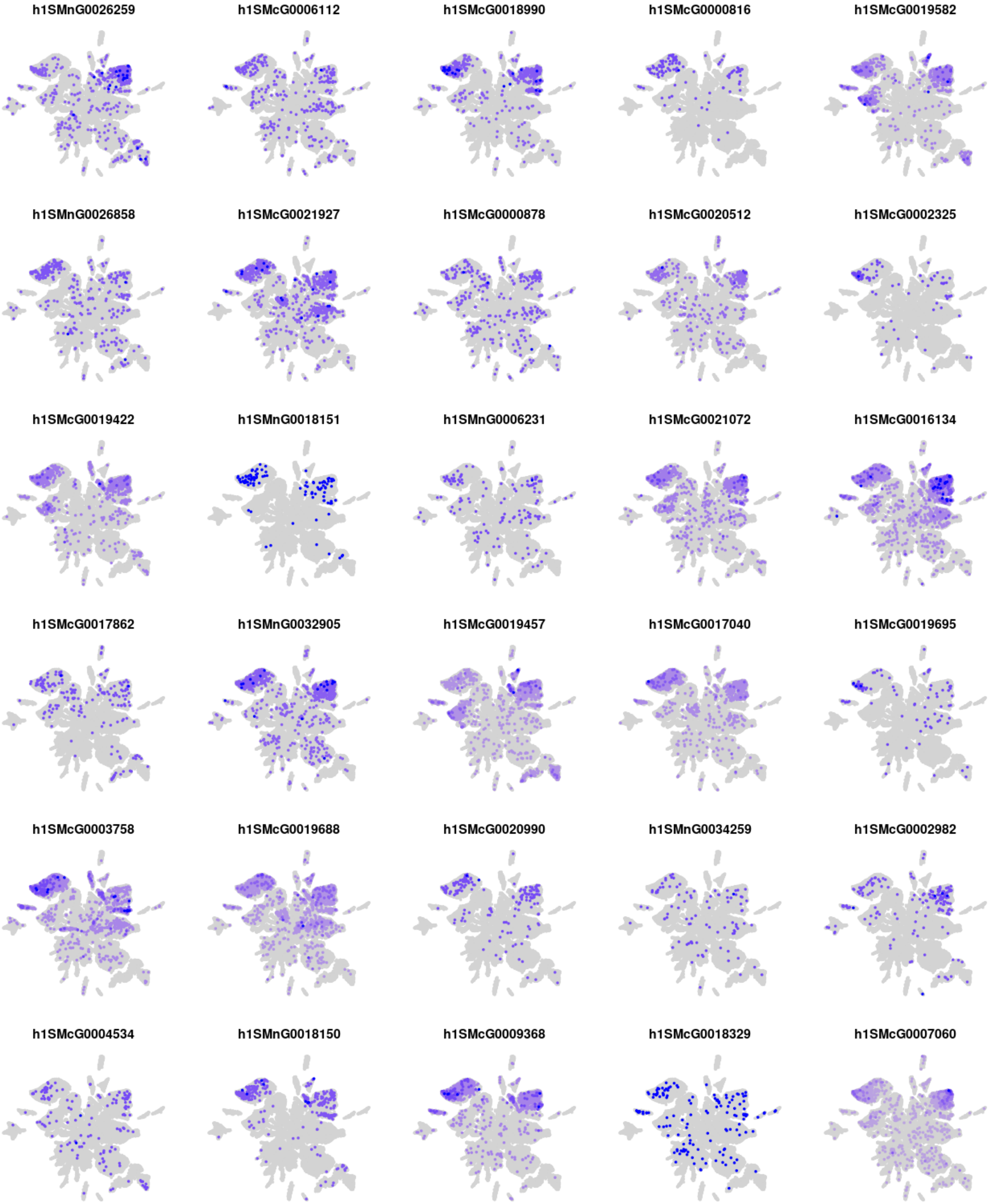
h1SMnG0026259
h1SMcG0006112
h1SMcG0018990
h1SMcG0000816
h1SMcG0019582
h1SMnG0026858
h1SMcG0021927
h1SMcG0000878
h1SMcG0020512
h1SMcG0002325
h1SMcG0019422
h1SMnG0018151
h1SMnG0006231
h1SMcG0021072
h1SMcG0016134
h1SMcG0017862
h1SMnG0032905
h1SMcG0019457
h1SMcG0017040
h1SMcG0019695
h1SMcG0003758
h1SMcG0019688
h1SMcG0020990
h1SMnG0034259
h1SMcG0002982
h1SMcG0004534
h1SMnG0018150
h1SMcG0009368
h1SMcG0018329
h1SMcG0007060

### Module m50

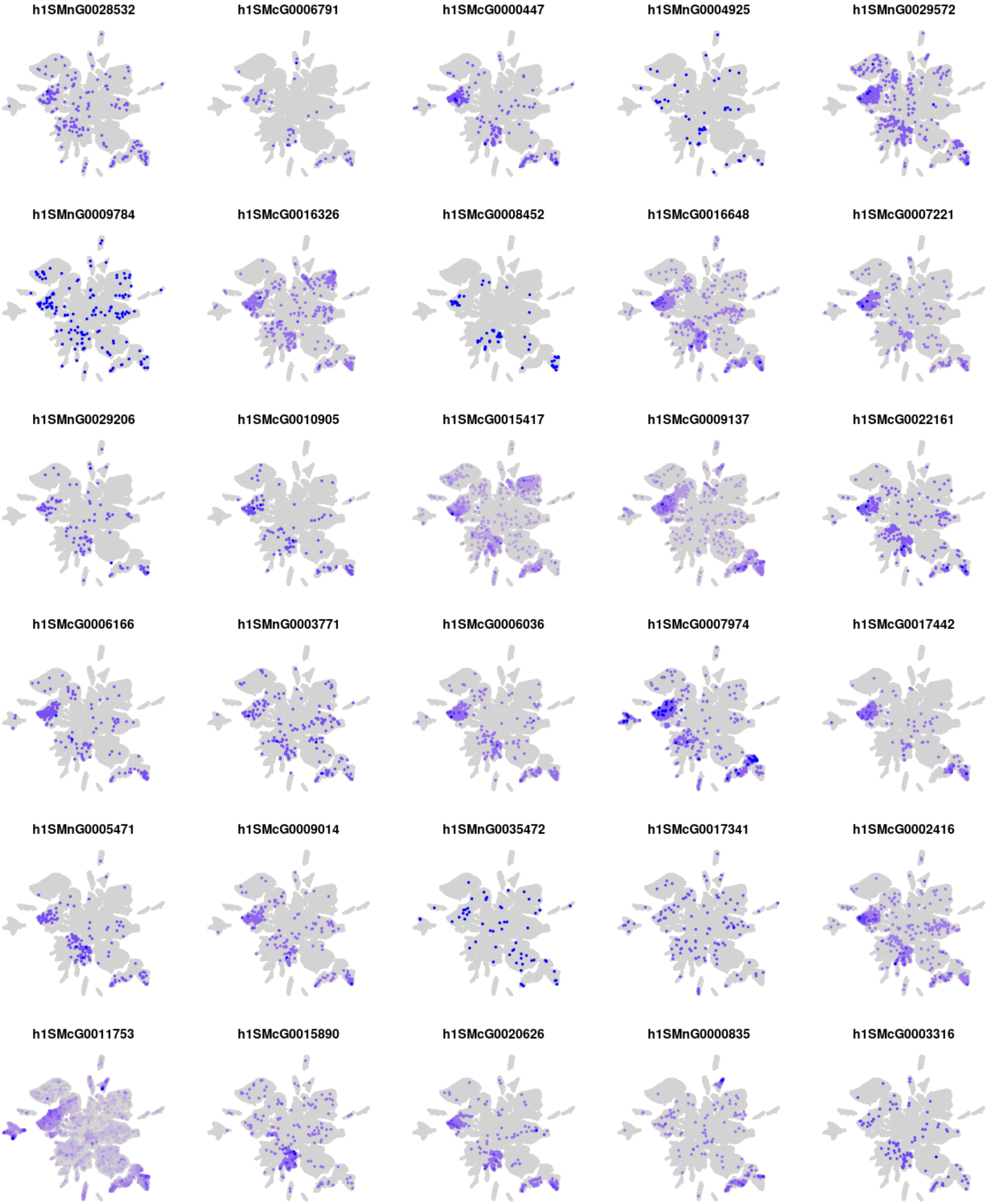
h1SMnG0028532
h1SMcG0006791
h1SMcG0000447
h1SMnG0004925
h1SMnG0029572
h1SMnG0009784
h1SMcG0016326
h1SMcG0008452
h1SMcG0016648
h1SMcG0007221
h1SMnG0029206
h1SMcG0010905
h1SMcG0015417
h1SMcG0009137
h1SMcG0022161
h1SMcG0006166
h1SMnG0003771
h1SMcG0006036
h1SMcG0007974
h1SMcG0017442
h1SMnG0005471
h1SMcG0009014
h1SMnG0035472
h1SMcG0017341
h1SMcG0002416
h1SMcG0011753
h1SMcG0015890
h1SMcG0020626
h1SMnG0000835
h1SMcG0003316

#### Module m51

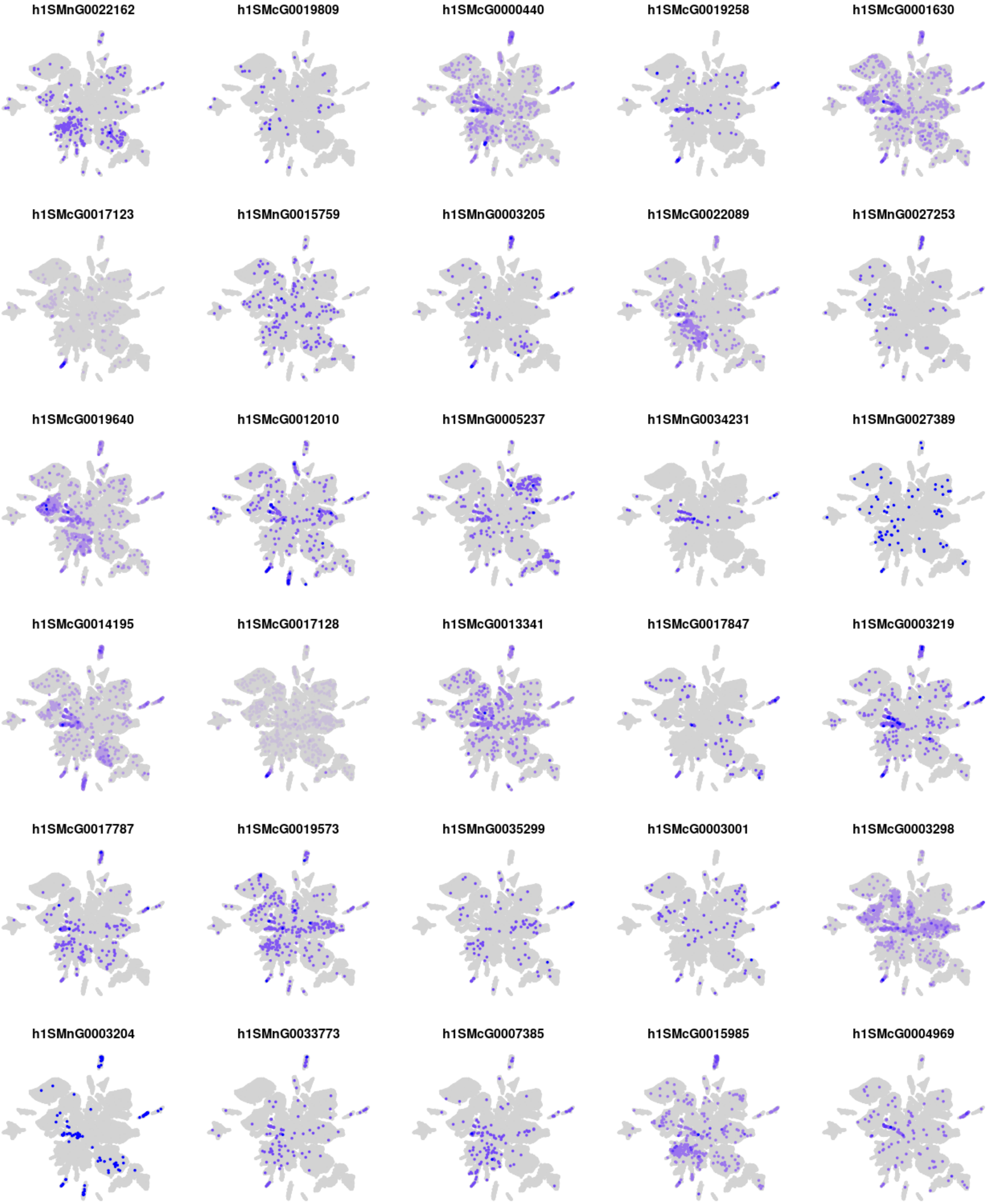
h1SMnG0022162
h1SMcG0019809
h1SMcG0000440
h1SMcG0019258
h1SMcG0001630
h1SMcG0017123
h1SMnG0015759
h1SMnG0003205
h1SMcG0022089
h1SMnG0027253
h1SMcG0019640
h1SMcG0012010
h1SMnG0005237
h1SMnG0034231
h1SMnG0027389
h1SMcG0014195
h1SMcG0017128
h1SMcG0013341
h1SMcG0017847
h1SMcG0003219
h1SMcG0017787
h1SMcG0019573
h1SMnG0035299
h1SMcG0003001
h1SMcG0003298
h1SMnG0003204
h1SMnG0033773
h1SMcG0007385
h1SMcG0015985
h1SMcG0004969

### Module m52

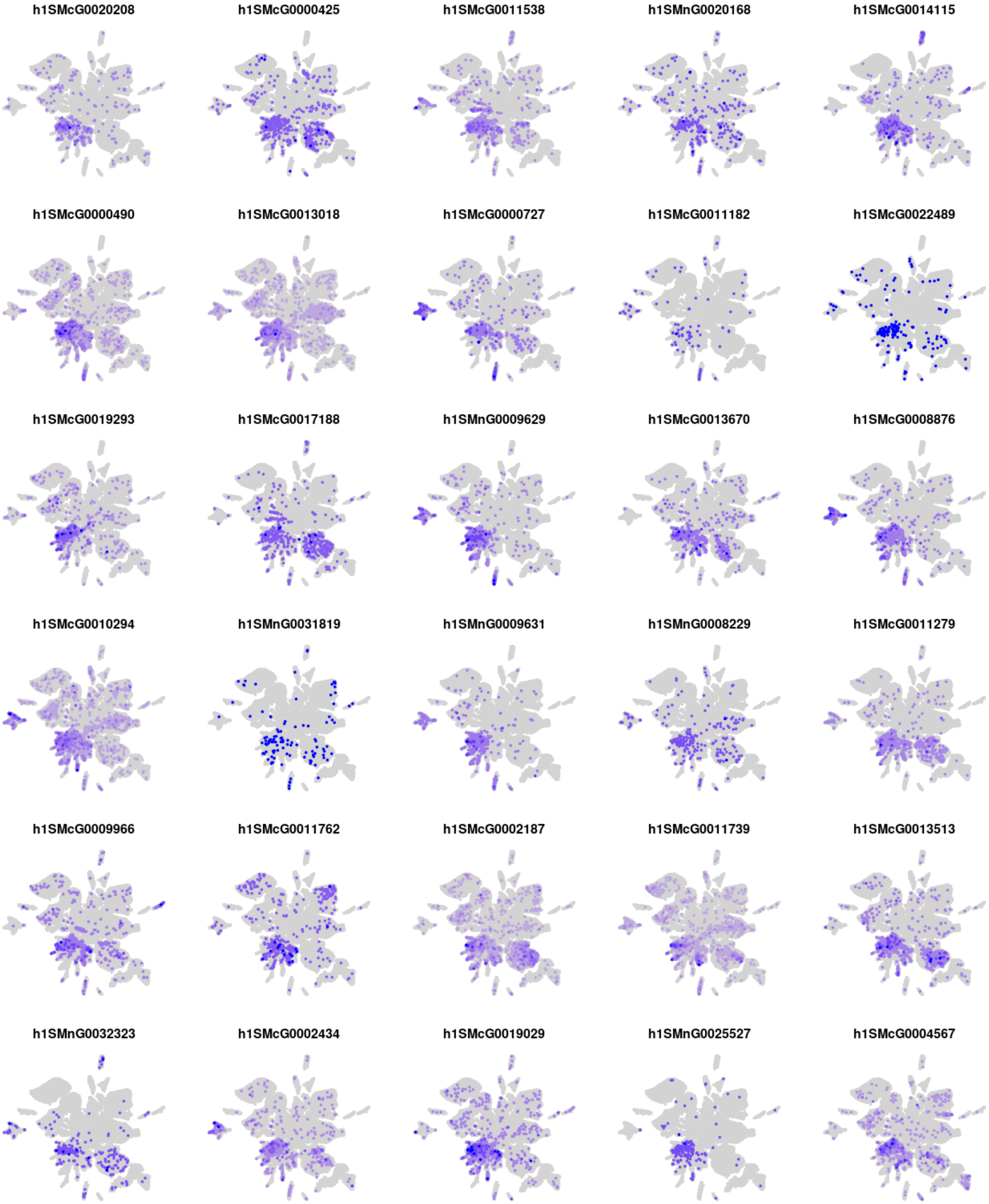
h1SMcG0020208
h1SMcG0000425
h1SMcG0011538
h1SMnG0020168
h1SMcG0014115
h1SMcG0000490
h1SMcG0013018
h1SMcG0000727
h1SMcG0011182
h1SMcG0022489
h1SMcG0019293
h1SMcG0017188
h1SMnG0009629
h1SMcG0013670
h1SMcG0008876
h1SMcG0010294
h1SMnG0031819
h1SMnG0009631
h1SMnG0008229
h1SMcG0011279
h1SMcG0009966
h1SMcG0011762
h1SMcG0002187
h1SMcG0011739
h1SMcG0013513
h1SMnG0032323
h1SMcG0002434
h1SMcG0019029
h1SMnG0025527
h1SMcG0004567

### Module m53

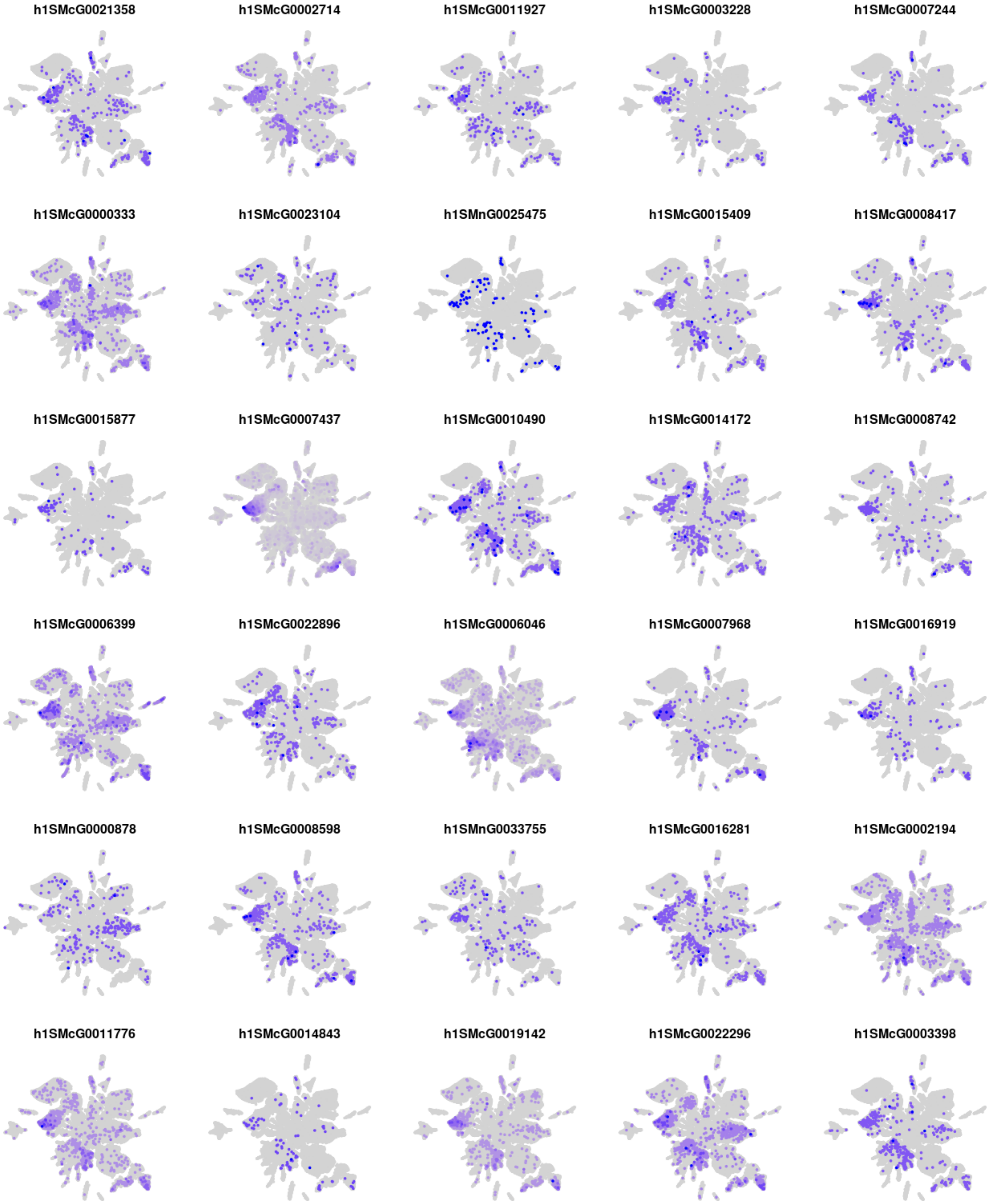
h1SMcG0021358
h1SMcG0002714
h1SMcG0011927
h1SMcG0003228
h1SMcG0007244
h1SMcG0000333
h1SMcG0023104
h1SMnG0025475
h1SMcG0015409
h1SMcG0008417
h1SMcG0015877
h1SMcG0007437
h1SMcG0010490
h1SMcG0014172
h1SMcG0008742
h1SMcG0006399
h1SMcG0022896
h1SMcG0006046
h1SMcG0007968
h1SMcG0016919
h1SMnG0000878
h1SMcG0008598
h1SMnG0033755
h1SMcG0016281
h1SMcG0002194
h1SMcG0011776
h1SMcG0014843
h1SMcG0019142
h1SMcG0022296
h1SMcG0003398
