## Supplementary material for "The Regulatory Logic of Planarian Stem Cell Differentiation": supp_file_08.pdf

Motif enrichment analysis (qvalue <0.1)

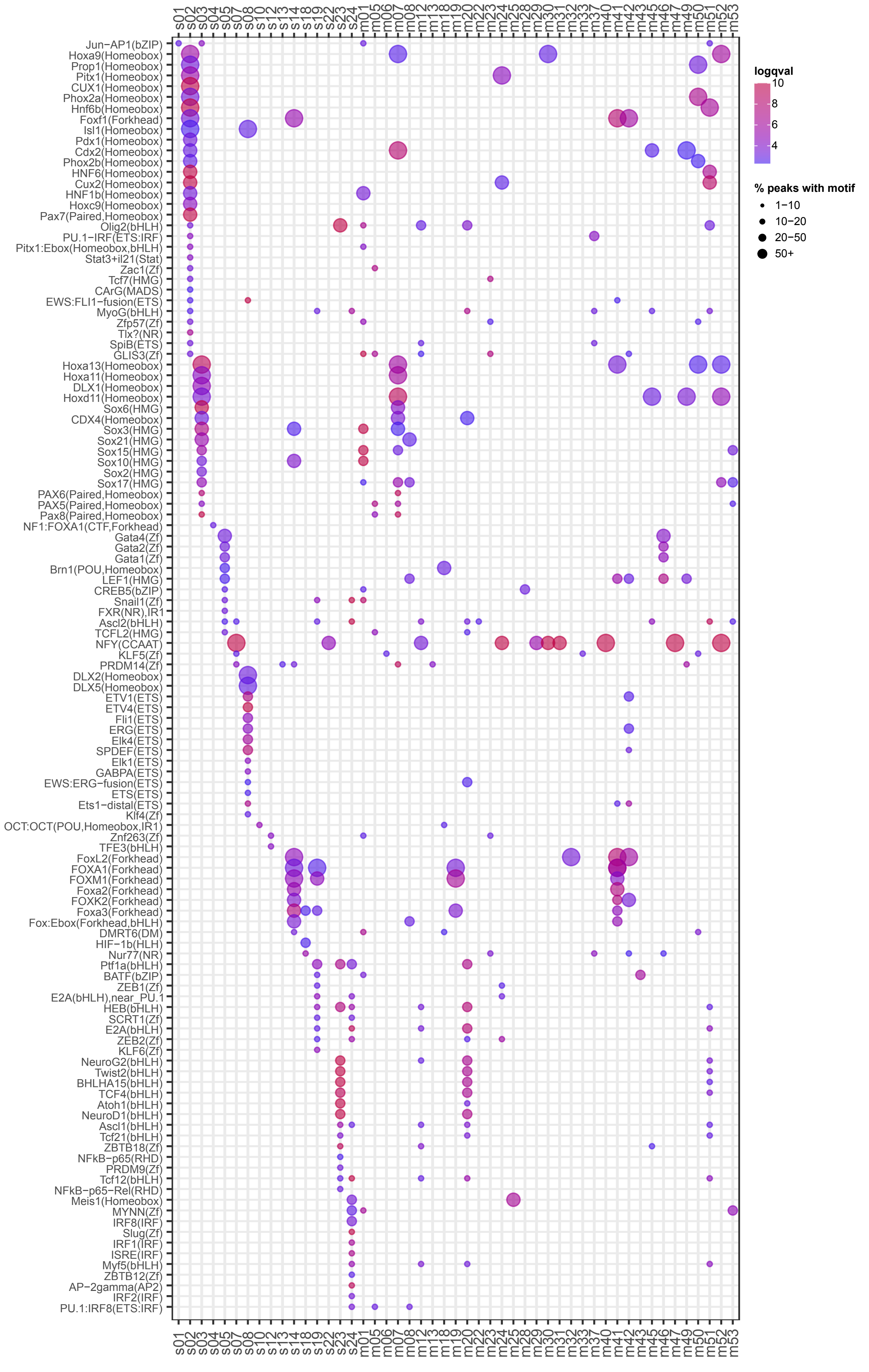


(continues in following page)

Motif enrichment analysis (qvalue <0.1) (continued from previous page)

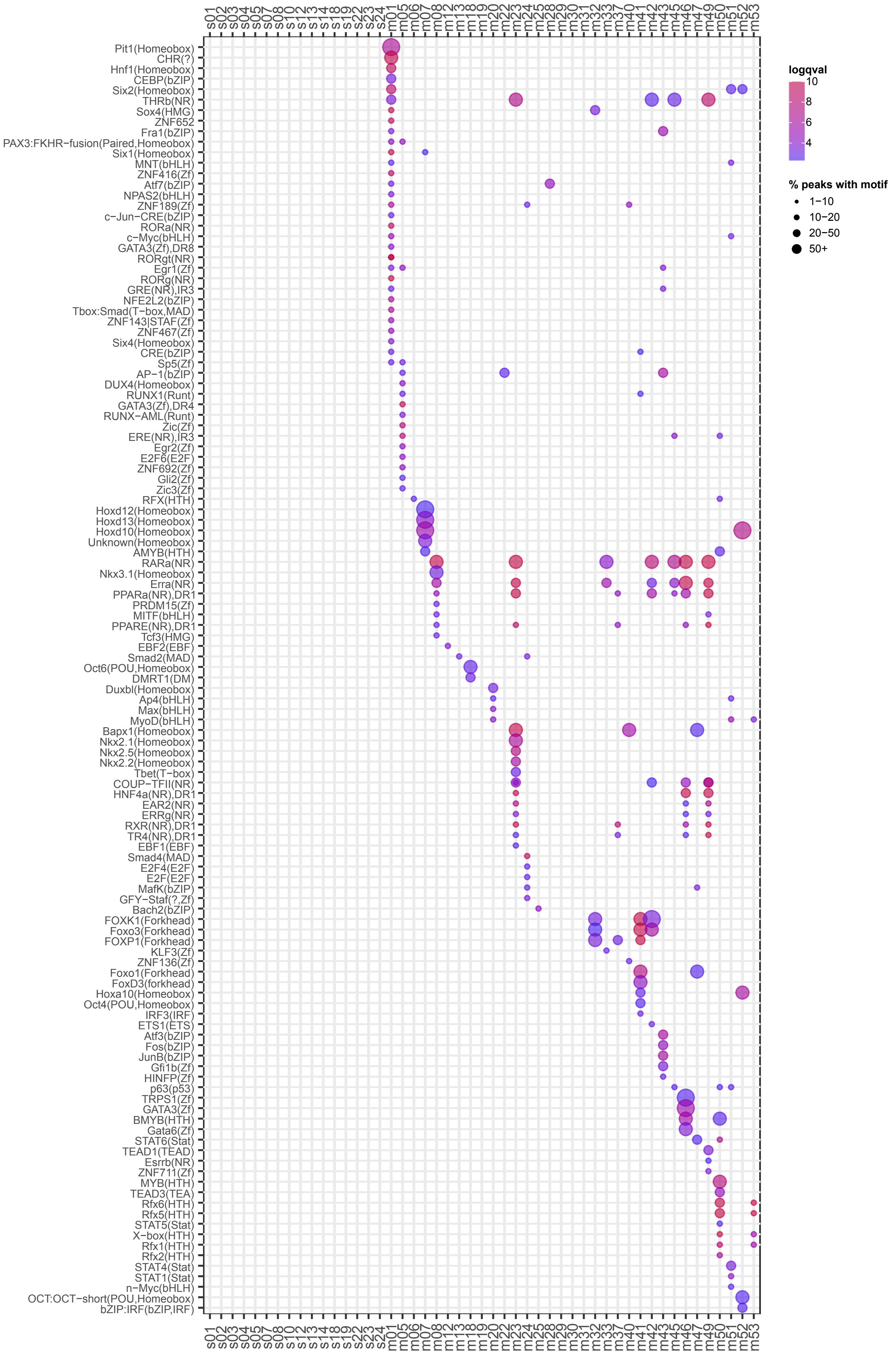
