## Supplementary figures and images for "The Regulatory Logic of Planarian Stem Cell Differentiation"

### supp_file_05.pdf

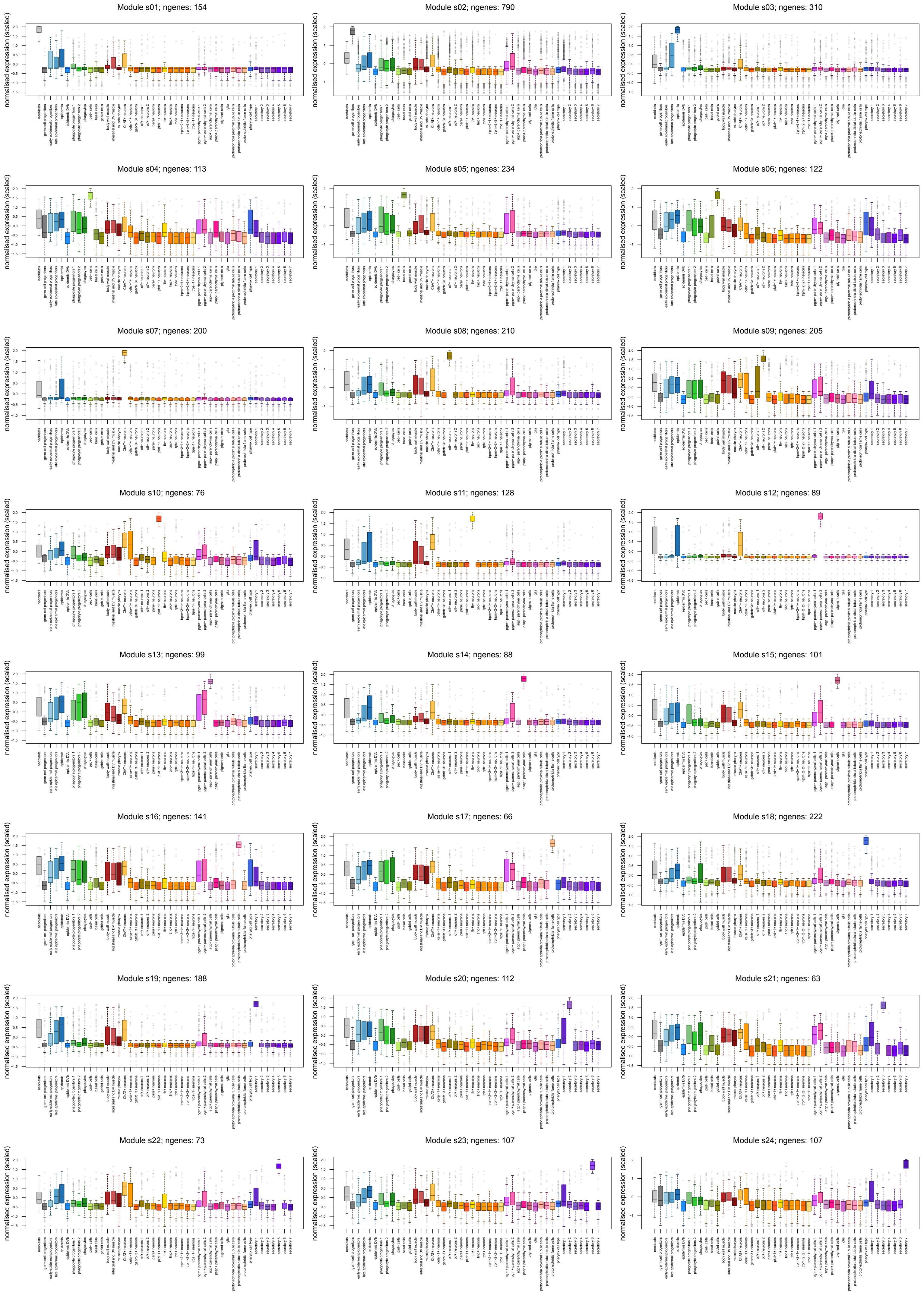

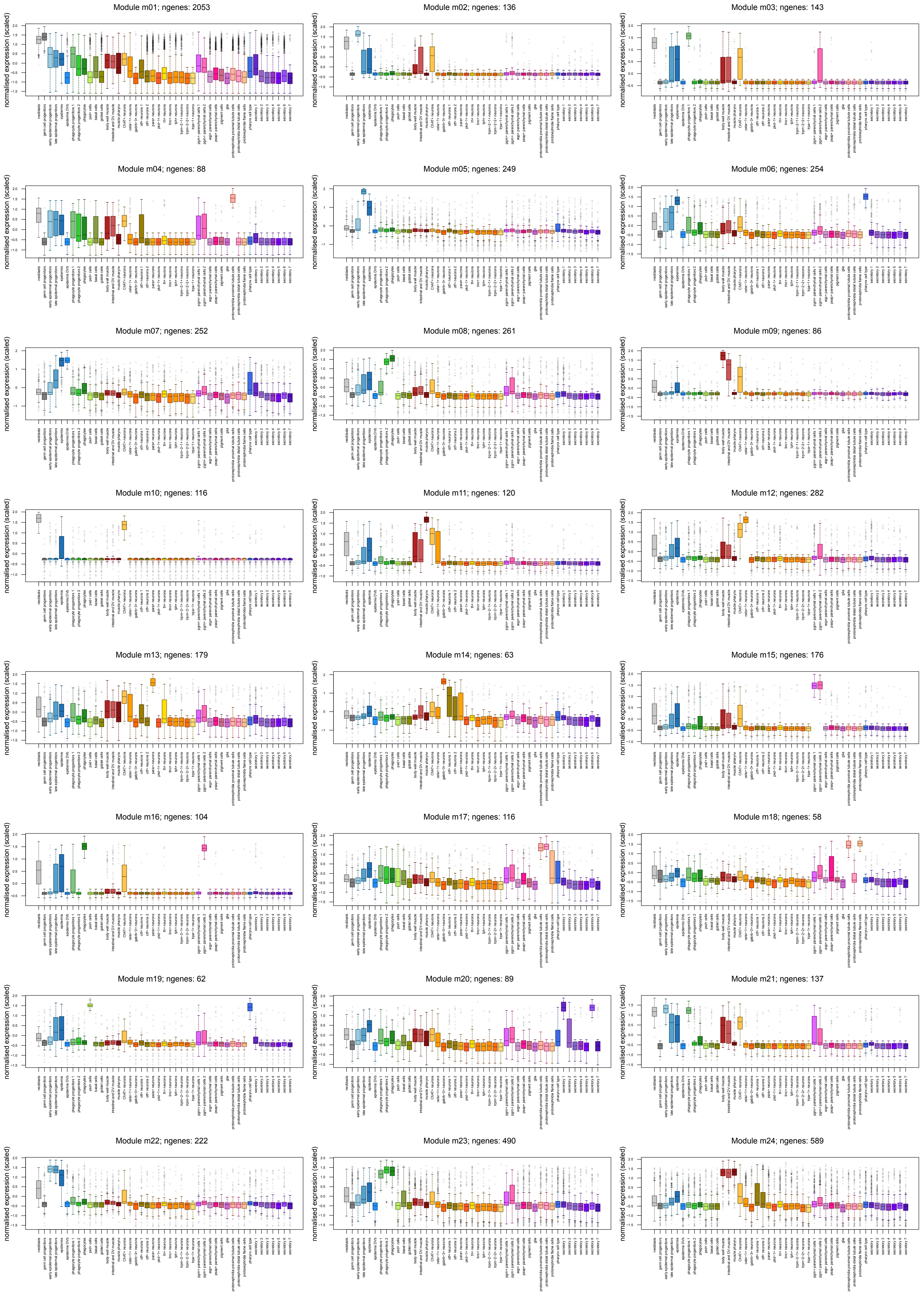

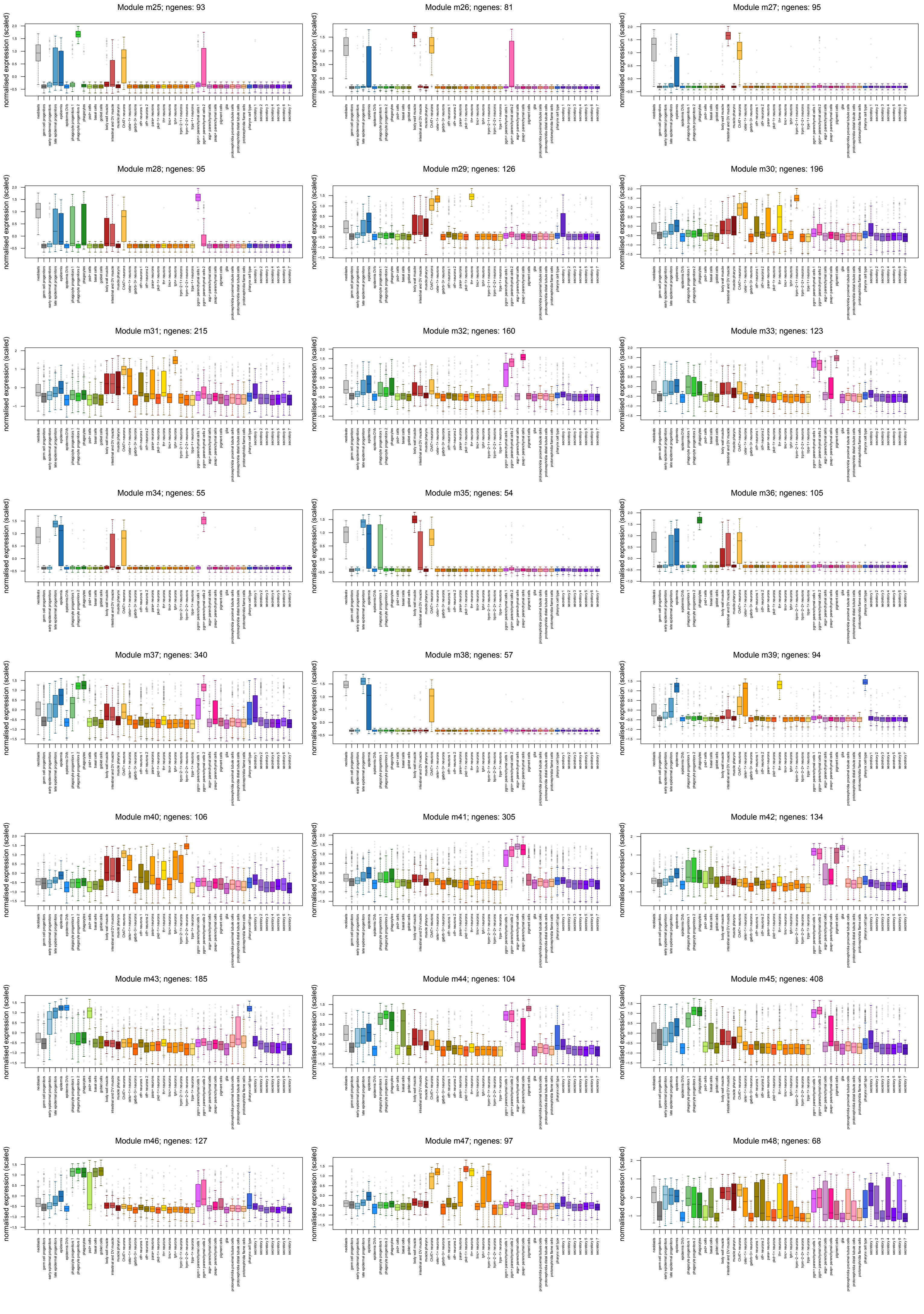

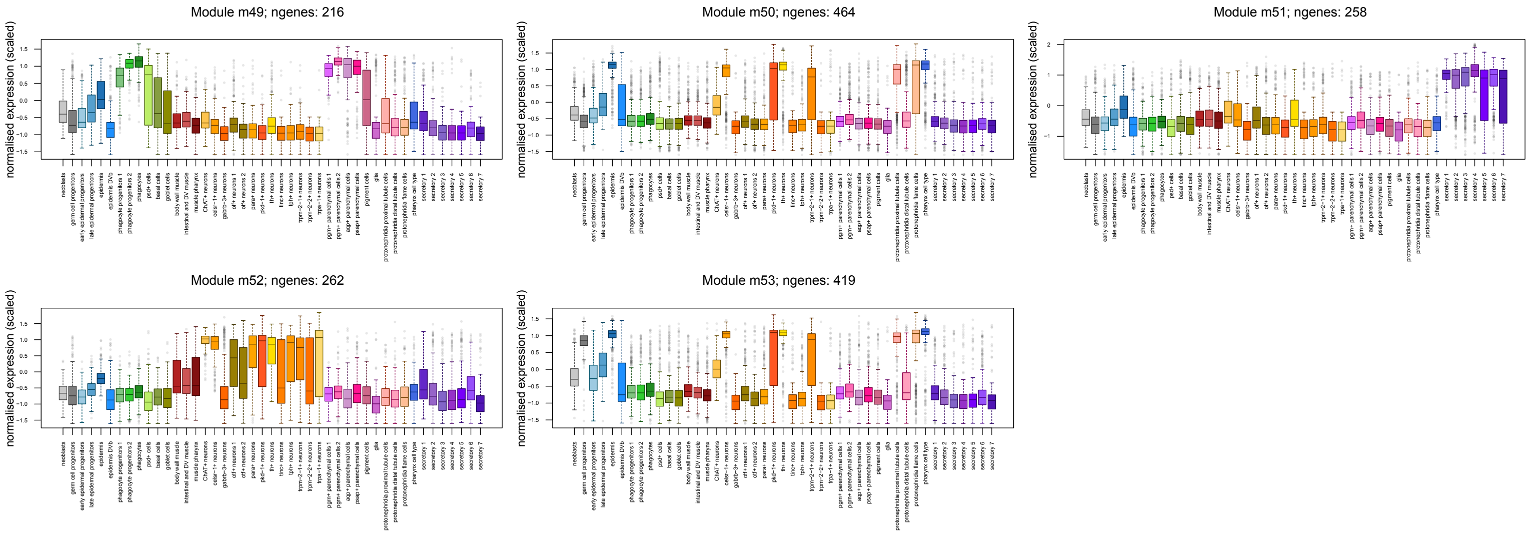

### supp_file_07.pdf

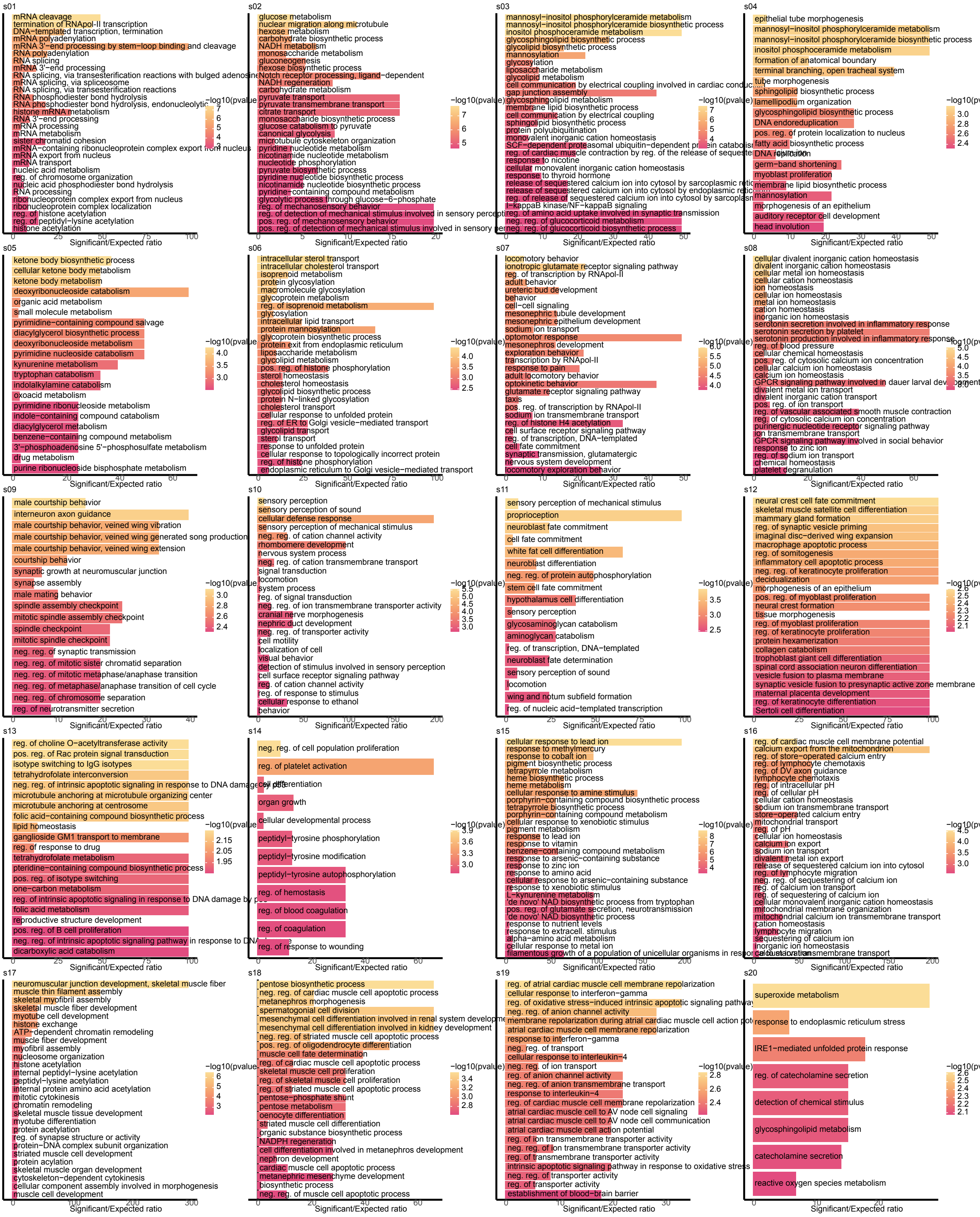

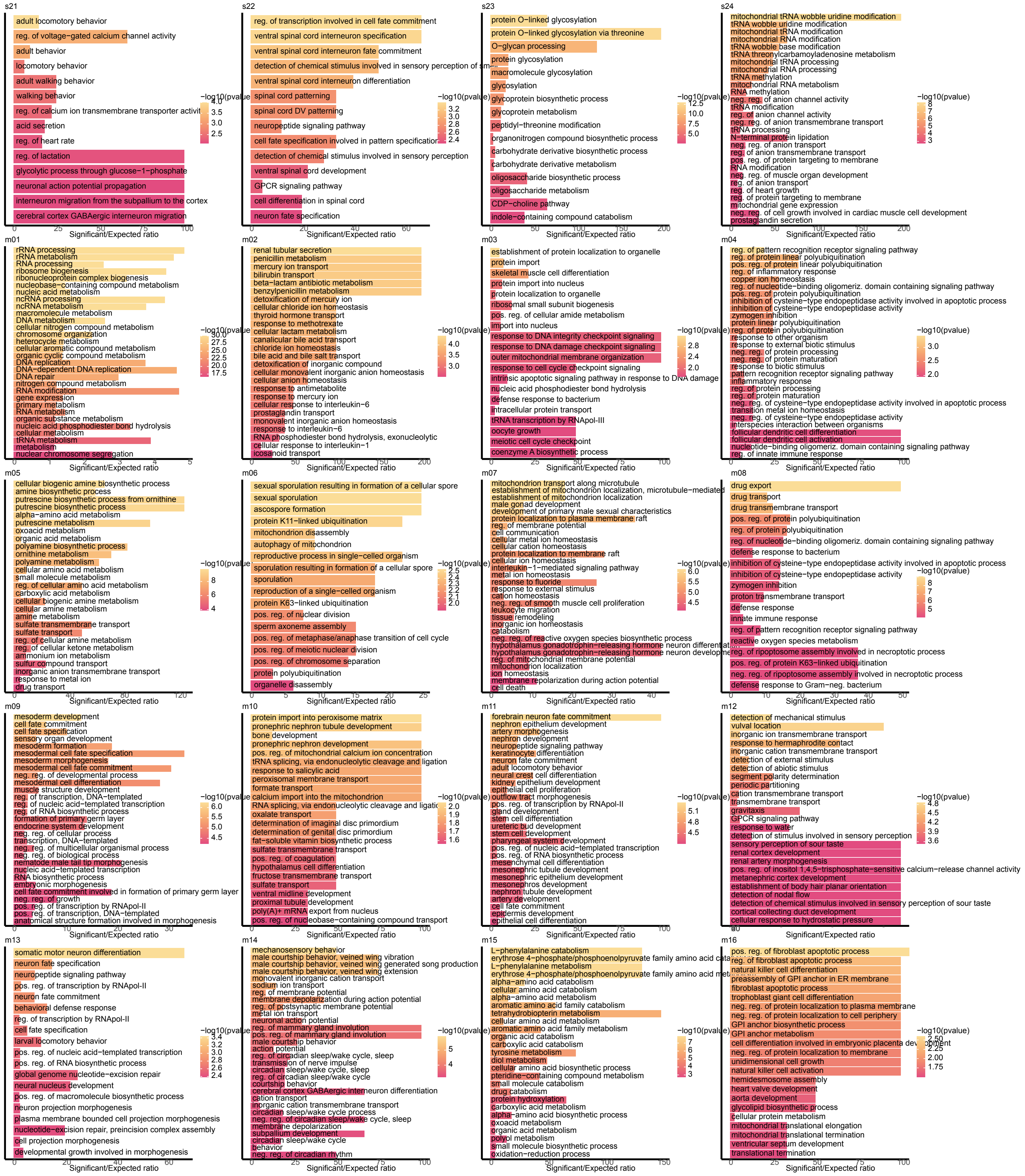

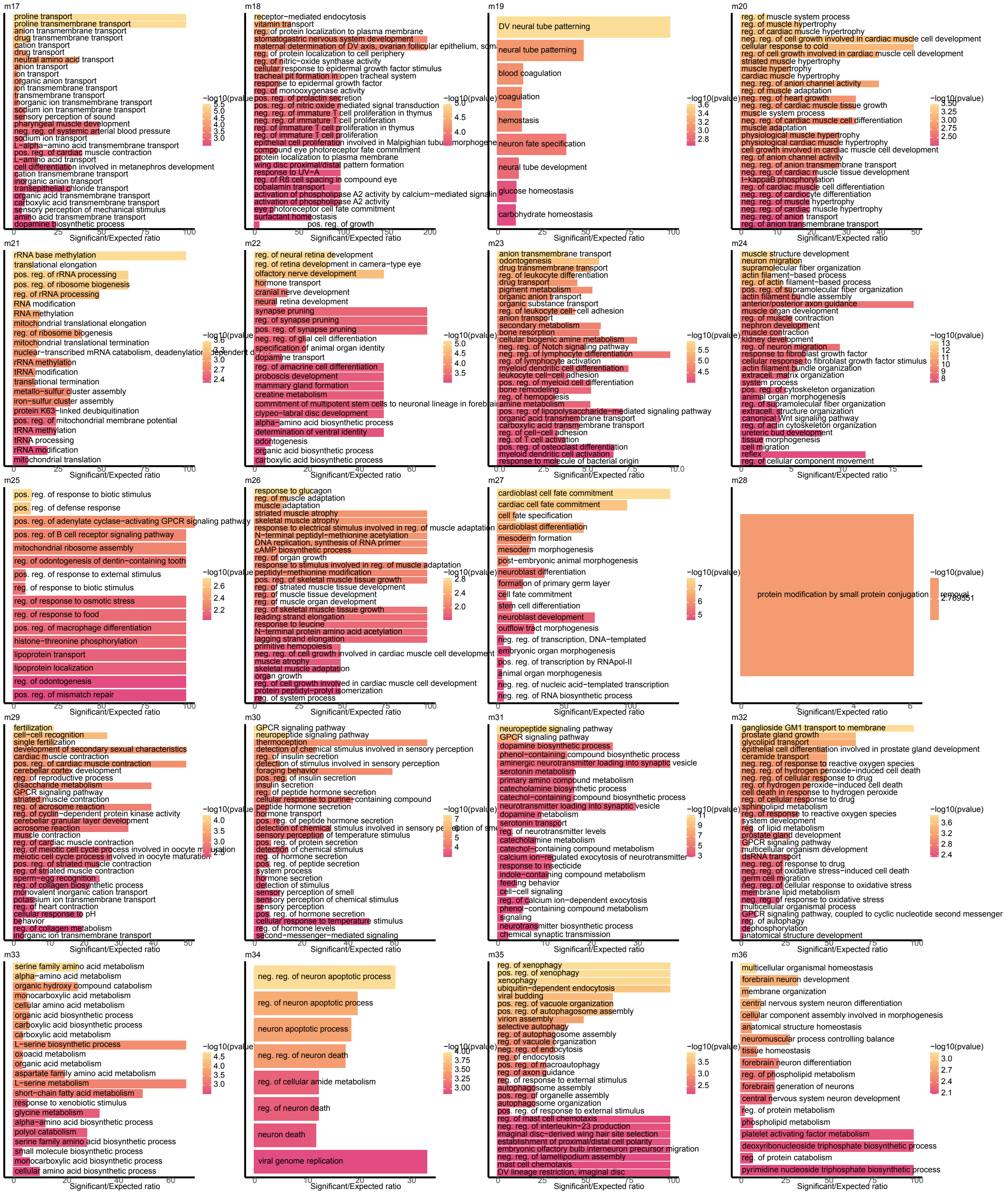

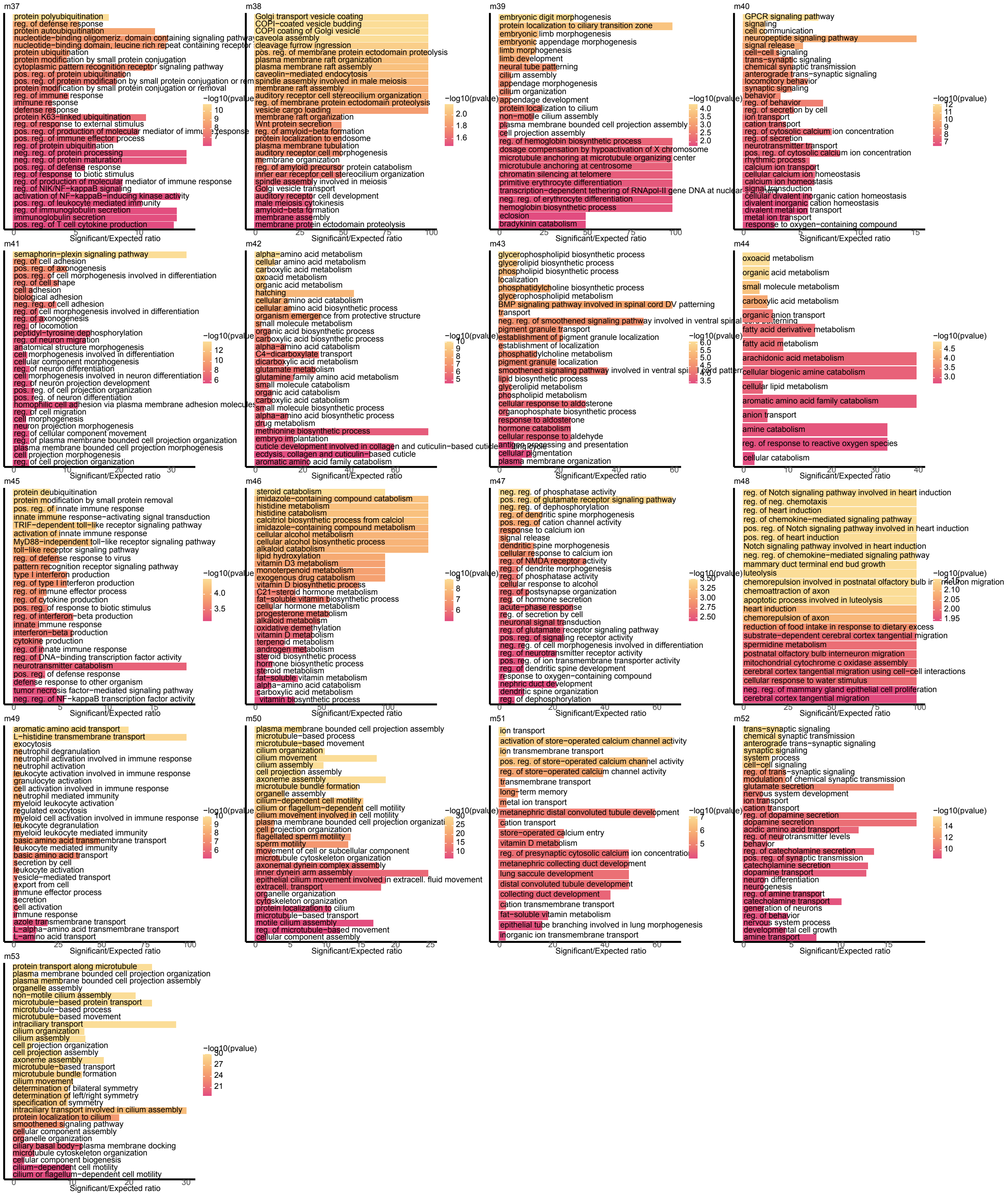

### supp_file_15.pdf

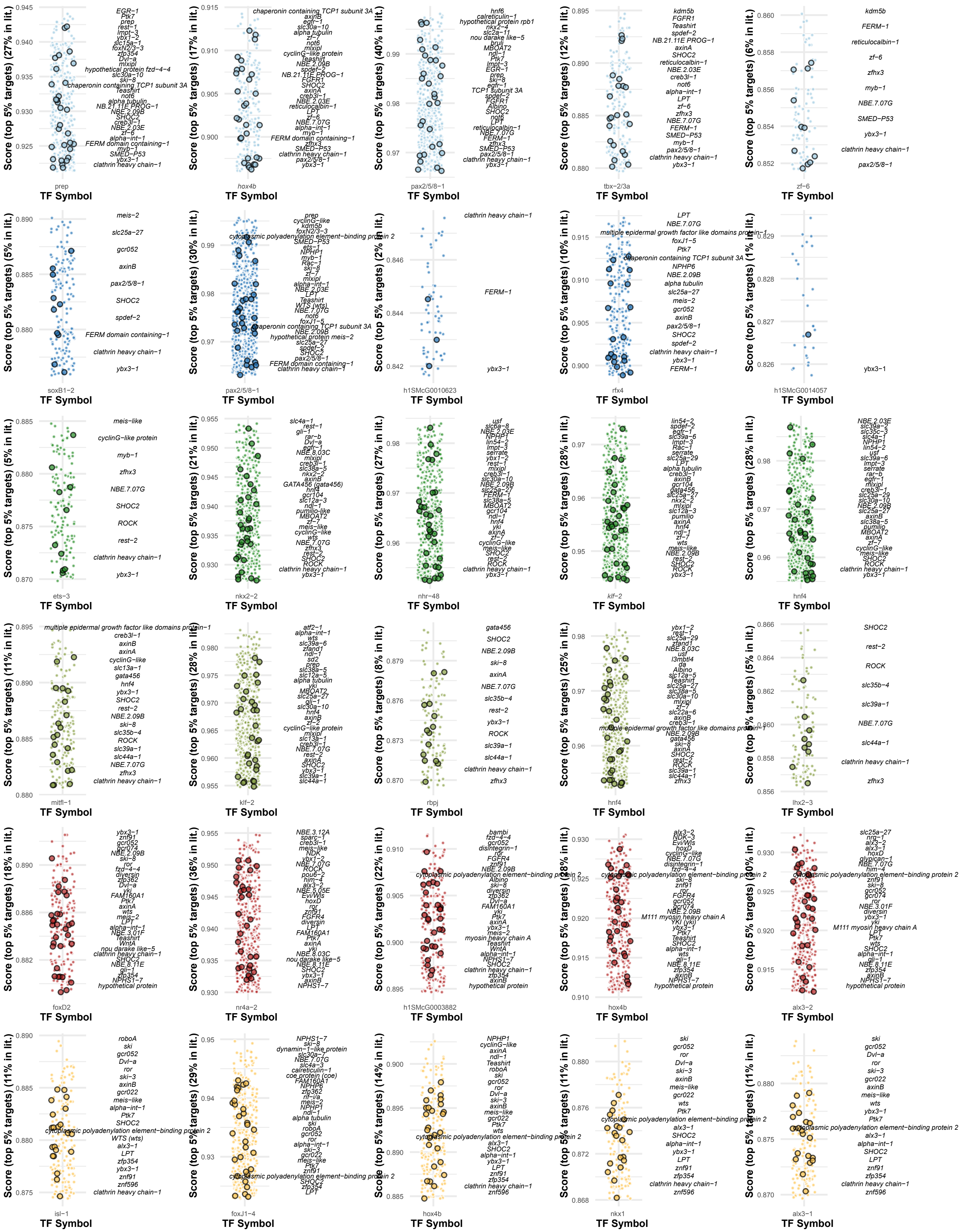

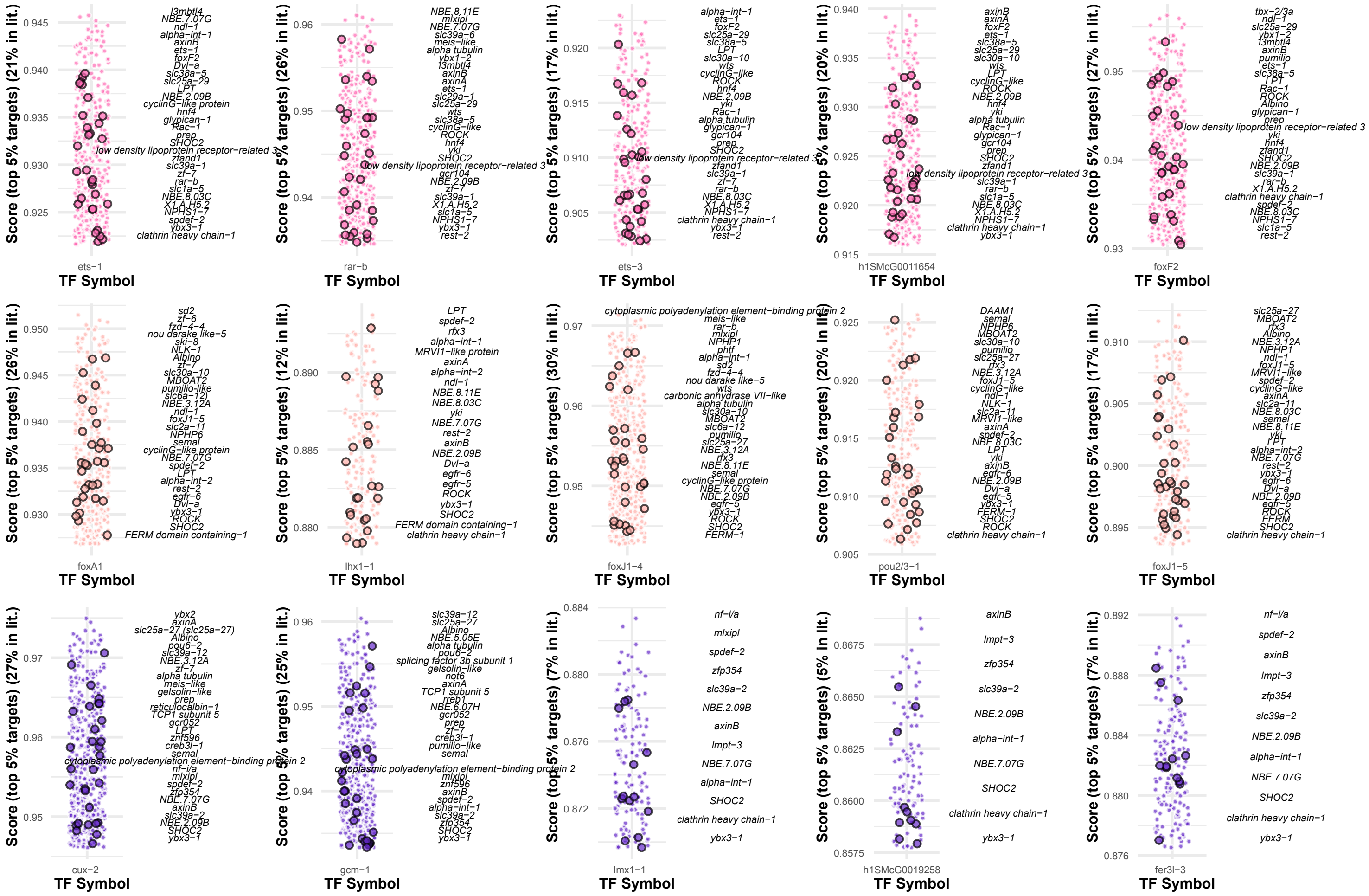

### supp_file_16.pdf

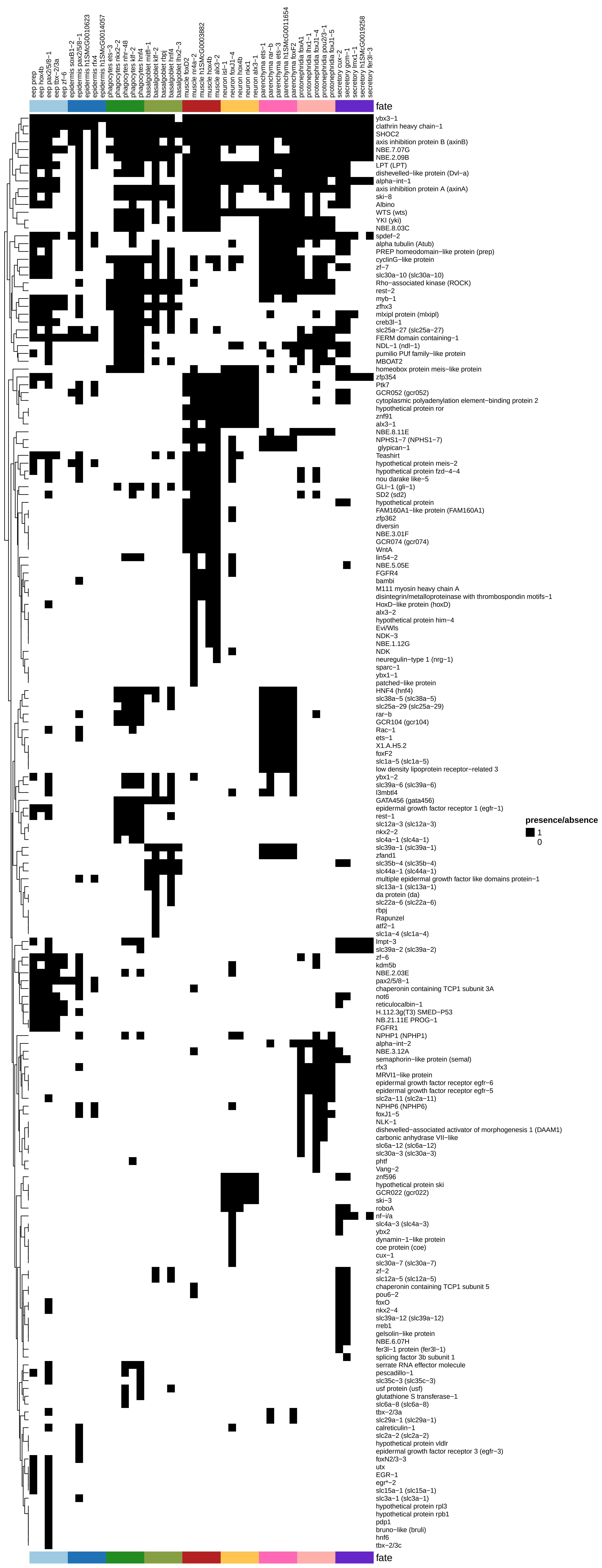
