## Supplementary Information for "The Regulatory Logic of Planarian Stem Cell Differentiation"

### Supplementary Notes

#### Supplementary Note 1: On pseudo-bulk data and Weighted Gene Correlation Network Analysis (WGCNA)

Single cell analyses have generally focused on the one-versus-all approach for finding markers of cell types. We sought to investigate gene expression with a focus on the gene space. In other words, after having found groups of cells that express similar sets of genes with an one-versus-all approach (standard procedure with tools like Scanpy or Seurat), we sought to find groups of genes that express in similar ways across one or more of the cell types detected in our dataset. For this, we focused on the use of well-established tools in the field of bulk-sequencing such as Weighted Gene Correlation Network analysis (WGCNA) [1].

Briefly, WGCNA detects groups of genes with similar expression profiles across samples (where a sample can be anything from a tissue, experimental condition, or in our case, cell clusters), using:

(i) weighted correlation to detect connections between genes based on expression;

(ii) a topology overlap algorithm of the connections detected in step (i) to further refine connections between genes sharing many neighbours;

and (iii) detecting gene modules based on dynamic cutting of a clustering of genes in a tree of similarity using the data from step (ii).

WGCNA has been implemented in several platforms and for different kinds of data. In this study, we used the latest version of the R package `WGCNA` [1]. We refer to Zhang and Horvath [2] for a fully detailed explanation and application of the algorithm. Of note is that the original algorithm of WGCNA uses absolute values of correlations in the adjacency step, meaning it can potentially group genes with high positive correlation together with genes to which they correlate strongly but negatively. In practice, data derived from single cell methods tends to attract genes with high positive correlations only. This is because there are more cases of genes very highly expressed in one or few cell types than cases of genes with an exact inverse pattern, where a gene is generally expressed in all but one or few cell types.

We opted for a pseudobulk approach in order to compensate for the sparsity of the single cell data, as previously done in Álvarez-Campos *et al.* [3]. By aggregating counts of cells for each cluster, one can retrieve a matrix of 'genes x clusters' like those used in bulk sequencing data. In preliminary analyses, we normalised the pseudobulk data based on the size of each "library" (each pseudobulk cluster of cells) in a similar fashion to quantile normalisation, as implemented in packages such as DESeq2 [4]; this had the benefit of equating the dynamic range of variability between clusters, allowing for comparisons between clusters of small and large numbers of cells. However, we found that this normalisation had the disadvantage of inflating counts of spurious or lowly expressed genes in relation to the respective normalisation applied to that same gene(s) on large clusters. For example, a housekeeping gene found in three cells of a large cluster (thousands of cells) might have its expression dampened compared to their expression in two cells of a small (dozens or few hundreds) of cells. This led, in our experience, to genes with artificially enlarged expression values in small clusters, making them indistinguishable from (or at least, at comparable orders of magnitude to those of) truly cell type-specific genes. To our understanding, library size normalisation on pseudobulk data does not fully account for the bias of detection due to the low abundance of several cell types. We sought to investigate alternative ways to account for small clusters.

From a methodological standpoint, it is safe to assume that not every gene that is being expressed in a cell is truly indicative of the function and the dynamics of a given cell type. Firstly, there are a number of genes essential to the function of any cell that will be expressed at basal levels or when required, and this can be retrieved by the sampling. By this we mean both basal housekeeping genes (genes with very few counts per cell, in many cells of any cell type) and genes with occasional expression in any cell type (genes with moderate or high counts in few cells of any given cell type). Secondly, different sources of background signal and experimental artefacts must be considered, such as the likelihood of finding doublets, errors in the barcoding, or artefacts of the k-NN graph and clustering [5, 6]. We reasoned that the likelihood of capturing expression of a gene that is truly part of the functional machinery of a given cell type relative to the rest of the organism, is directly proportional to the fraction of cells of a given cell type (or in a cell cluster) expressing that gene relative to the rest of the organism. Thus, for a gene to be considered reliably expressed on a cell type, one must consider the dynamics of that gene in the whole dataset. One possibility is to take into account the fraction of cells expressing that gene in that cell type, in relation to the fraction of cells expressing that gene in the rest of the dataset. The relation between these two fractions can potentially be used as a dynamic weight of gene expression on each cell type, in order to better model a representative transcriptomic snapshot of each cell type.

For this approach, we computed another pseudobulk matrix, this time counting not the number of counts of each gene in each cluster, but the number of cells expressing each gene in each cluster. Then, for every gene *i* and every cluster *j* and using this matrix of number of cells, we calculated:

(i) The fraction of cells from cluster *j* expressing gene *i* (we called this number $a_{ij}$). This number is calculated by dividing the number of cells from cluster *j* expressing gene *i* divided by the number of cells in cluster *j*, and can range between 0 (no cells) and 1 (every cell);

$$a_{ij} = \frac{n_{ij}}{N_j}$$

(ii) The fraction of cells not from cluster j expressing gene i (we called this number $b_{ij}$). This number is calculated by dividing the number of cells that do not belong to cluster *j* expressing gene *i* divided by the number of cells in the dataset that do not belong to cluster *j*, and can range between 0 (no cells) and 1 (every cell);

$$b_{ij} = \frac{n_{i\bar{j}}}{N_{\bar{j}}}$$

(iii) The fraction between $a_{ij}$ and $b_{ij}$, which we call $C_{ij}$ . This number can be as low as 0 (no cells in cluster *j* expressing the gene compared to the rest of the dataset) but the range of values has no upper boundary (meaning a large fraction of cells in cluster *j* is expressing that gene compared to the rest of the data, which can be disproportionate if few or no cells are expressing that gene outside cluster *j*).

$$C_{ij} = \frac{a_{ij}}{b_{ij}}$$

(iv) This number, whose values can range from zero to infinite, is mapped to a range [0,1] using the formula `*1 - exp(- $C_{ij}$)*`, in order to be used as a weight. We call this number $w_{ij}$.

$$w_{ij} = 1 - e^{-C_{ij}} = 1 - e^{\frac{-a_{ij}}{b_{ij}}}$$

The resulting count matrix from normalising by library size can be adjusted using this matrix of weights. In practice, this dampens the spurious counts of lowly-expressed genes in small clusters.

We performed some preliminary comparisons between different normalisations (data not shown). We ran WGCNA with each of them using broadly similar parameters and we observed that, compared to just library size normalisation or log(library size normalisation), using cell weights on top of log transformation of library size-normalisation performed better at detecting smaller modules for small cell clusters. With this method we did not observe large modules specific to small clusters that upon inspection corresponded to genes expressed broadly in the dataset, as it happened with normalisation by library size only. Therefore we decided to adjust the library size normalisation by log transforming and using these weights for adjustment. Following WGCNA standard practices as well as previous works [7], we filtered out genes with less than thirty counts across the dataset (<30) and genes with low coefficient of variation too. As examples, lowly expressed genes expressed in infrequent cell types such as *pitx* (h1SMcG0012776) [8, 9] and *estrella* (h1SMcG0019080) [10] had 89 and 178 counts respectively.

WGCNA posits the idea that biological networks such as those of gene-gene interactions follow a scale-free model of graph topology [11], whereby the degree (number of connections per gene) distribution of the graph follows a power law. In other words, a scale-free network is expected to have many genes with few neighbours and few genes with many neighbours, these few genes acting as "hubs". Despite correlation alone (such as Pearson or Spearman) can coalesce genes in groups of similar expression profiles, weighing these correlation values using a soft threshold (or soft power) parameter is able to highlight strong correlations relative to weaker ones. This soft thresholding method can effectively yield a scale-free graph from correlation data, hence its use.

In order to detect the best-suited soft power, we ran the `pickSoftThreshold()` function from the WGCNA R package on our weighted, normalised data, and chose soft power 8 as it showed a high scale free topology fit (based on Zhang and Horvath [2]) as well as a small median connectivity in order to detect smaller modules (Supplementary Figure 6B,C). After this, we ran the `adjacency()` and `TOMsimilarity()` functions to generate a topology overlap matrix (TOM), the opposite of which (1 - TOM matrix) can be used as a distance matrix for clustering the genes (Supplementary Figure 6D). We clustered the distance matrix derived from the TOM similarity matrix, and we finally ran the `cutreedynamic()` function of WGCNA. We set the parameter `deepSplit = 3` to retrieve small modules, and set a minimum module size to 50 genes. We explored these modules by checking their overall expression dynamics (Supplementary File 5), their gene ontology enrichments (Supplementary File 7), and individual expression patterns on our scRNA original dataset (Supplementary File 6).

We sorted these modules in a semi-automated way by leveraging the dynamics of expression patterns of the genes constituting each module. We reasoned that modules could be majoritarily distributed in two groups: those which are highly specific for a given cell type (which we called "s" modules), and those whose genes are expressed in two or more cell types (which we called "m" modules). One way to tell these apart can be by checking in which cell type, or cell types, are their genes most often, and most highly, expressed. There are several ways to define what "highly expressed" means; for example, it can be measured as outlier values on the distribution of means, medians, or upper quartiles of expression on each cell type. These mean, median and upper quartile values can be derived from all of the genes of a module, or from those with intra-modular connectivity (==correlation with the average expression profile of the module) above a threshold value. In this study, for every module *i*, we calculated the upper quartiles of the distributions of normalised counts from every gene of module *i* on every cell type. We defined outliers of this distribution of upper quartiles as those values higher than the sum of the mean and 1.5 times the standard deviation of that distribution (Supplementary Figure 6E). Modules with only one outlier value of upper quartiles were defined as "s"; and those with two or more outlier upper quartiles were defined as "m". After defining them as "s" or "m", we reordered these modules following a similar pattern to the one chosen for listing the different cell clusters (i.e. neoblast first, epidermis second, etc, and then, neoblast and epidermis first, neoblast and phagocytes second, etc) (Supplementary File 4). Alternatively, there exist other methods such as the Tau metric as originally described in Yanai *et al* [12] and used more recently by Mantica *et al*. [13].

To visualise these modules we generated a heatmap using the ComplexHeatmap R package [14]. To facilitate the visualisation of the modules, we created profiles of the relative amount of gene expression of each module. For this, we first calculate the average expression profile of every gene from module *i* on every cell type, and then we then divide every value by the sum of values. This results in the frequency of expression of each module *i* on every cell type. The resulting frequencies represent the average expression signal in each cluster, and these were visualised as stacked bar plots on the side of the heatmap (Figure 2A,B,C).

To further investigate these modules we generated a graph out of the TOM matrix using `igraph` [15], for different analyses and visualisation (Supplementary Figure 7A). We removed the spurious connections (connections between genes with values below the second-lowest value, below 0.01) and further explored these connections, observing the gene-gene connectivity value has an exponential decay-like pattern that can be bi or multimodal (Supplementary Figure 7B). For this dataset this value was around 0.3 and 0.4, which is slightly similar but not equal to our previous analyses [3]. We suggest exploring this for each dataset. Based on inspection of the distribution, we chose nine threshold values (Supplementary Figure 7C) over which we iterated to generate nine graphs and measure several metrics, such as the number of connected components, median degree, and median and number of genes per connected component (Supplementary Figure 7 D-G). This was instrumental to ascertain the appropriate threshold value, as e.g. the graph with edge values above threshold 0.25 showed a much higher number of connected components (Supplementary Figure 7D), but these were formed by much fewer genes (Supplementary Figure 7G). We decided to focus on the threshold value 0.35, and observed that the vast majority of all the connected components correspond to unique modules of those detected by the dynamic tree cutting of WGCNA (Supplementary Figure 7I).

Alternatively, since every gene in the graph belongs to a WGCNA module as computed earlier, one can isolate the different WGCNA modules from the graph which can be useful for intra-modular analyses, but this precludes further cross-module analyses as any connection between modules is lost. We followed this approach to study the relationship between TF connectivity and centrality. As suggested in our previous publication [3] we noticed TF intramodular connectivity is in agreement with TF centrality, both globally (when correlating the relative intramodular connectivity and the relative centrality of TFs from all modules) and module-wise (Supplementary Figure 7J). This can help identify potentially relevant TFs for regulating the expression of genes in a given module.

We then wondered what was the structure of this network at the module level. Are genes from one module more connected to other modules? If so, are they functionally similar? Are they similar at the transcriptional regulatory landscape (TF and motifs) too? To answer these questions:

(i) We first generated a cross connection graph as described previously [3], by counting how often a gene of a given module *i* is connected to genes from other modules. We normalised this number by module size, as large modules likely tend to have more cross-connections. We used this 'module x module' matrix to generate a module-wise graph of cross-connections (Supplementary Figure 7K).

(ii) Secondly, we used the motif enrichment analyses we did previously (see Methods) to calculate a motif x module matrix of percentage of enrichment in gene promoters, in turn used to create a module-wise graph of motif enrichment similarity (Supplementary Figure 7L).

(iiI) Thirdly, for further exploration we did module connectivity of the TFs against the eigengene of every module (average profile of a module as done in WGCNA). We retrieved the profile of TF connectivity for each module and correlated this to make another module-wise graph (Supplementary Figure 7M).

(iv) Next, we did a COG functional category enrichment analysis as described in Álvarez-Campos *et al.* [3] and we used the percentages of over- or under-representation to generate a functional category x module matrix, to create a module-wise graph for functional category enrichment (Supplementary Figure 7N).

(v) We finally aggregated these graphs together counting how often a pair of modules appears connected in each (Supplementary Figure 7O).

Using the aggregated module-wise graph from step (v), we detected communities of modules using the `cluster_label_prop()` function from igraph, which overall aligned with our previous observations and provided a broad view of the dynamics at the gene space level in *S. mediterranea* at the coexpression, functional, and potential regulatory level.

Of note, we also followed a similar rationale when exploring the sets of co-accessible regions of open chromatin (Figure 3). We ran WGCNA and chose a soft threshold power, and we later sorted the modules of co-accessible regions in a similar fashion. Because our module sorting approach is actually agnostic from the source of the data (as long as these are groups of features such as genes or OCRs), we also followed this approach to group and sort the clusters of co-influential TFs.

#### Supplementary Note 2: ANANSE

Despite gene expression analysis and chromatin analysis providing valuable insights, these analyses are run separately from each other. For an integrated analysis of the two data modalities, we used ANANSE [16]. ANANSE is a tool that leverages TF/target gene annotation, gene expression, chromatin accessibility, and motif enrichment analysis to assign a probability to TF-target gene interactions based on a score from an additive model. If available, ANANSE can also leverage histone modification data such as H3K4me3 or H3K27Ac ChIP-Seq. The resulting output is a list of TF-target gene interactions defining a graph, or network, of TF-target gene predicted interactions.

Because we can computationally dissect our single cell data to isolate signals coming from each cell type, it is feasible to construct such networks of TF/target gene interactions for every *S. mediterranea* broad cell type. In addition, ANANSE incorporates a feature (ANANSE influence) to compare pairs of networks using differential gene expression analysis, in order to e.g. identify the potential TFs driving the transition from one given cell state to another. This approach proves instrumental to study these networks from the perspective of pluripotent stem cells committing to different cell fates, such as is the case of *S. mediterranea* neoblasts. Therefore we sought to investigate if our data is capable of recapitulating known dynamics related to cell type differentiation in *S. mediterranea*.

To create these networks, it becomes necessary to consider which genes in the dataset are TFs, and which motifs correspond to which TFs. ANANSE needs to tell TFs apart from non-TF genes, which can be achieved by providing a database of TF-motif association. There are several ways to do this, and we implemented a two-step solution:

(i) Firstly, the motif database of choice is relevant. ANANSE relies on gimmemotifs, a suite for motif enrichment analysis [17]. gimmemotifs incorporates motif databases from a number of projects, notably HOMER [18] or JASPAR [19]. The latter provides a tool to infer the predicted motif of a TF based on sequence homology and DNA-binding protein domain prediction [20]. Therefore, choosing the JASPAR database has the advantage that this information can be provided with a tool for that database.

(ii) Secondly, since gimmemotifs was designed for human and mouse data, it also has a tool called `motif2factors` that relies on automated orthology inference to identify TFs based on sequence homology, using Orthofinder [21] to transfer TF (and motif) annotation from human and mouse to the species of interest.

As explained in Methods, we ran `motif2factors` to transfer motif annotation from the JASPAR database, using a set of several protostome metazoan species (Supplementary File 13, Supplementary Figure 10A) to provide additional phylogenetic signal for the automated transfer. Second to this, we observed that not every TF detected by our TF annotation workflow (see Methods) received a motif. For this we ran the JASPAR similarity prediction tool [20] on those TFs that did not get any transferred motif, using the JASPAR 2024 motif database which overlaps with the JASPAR 2020 database.

Finally, to ensure the vast majority of TFs from the literature had an annotated motif for ANANSE, we supplied this motif annotation with any remaining TF that had an associated motif as detected by Neiro et al., 2022 [22].

The resulting database of 401 (out of 665) TFs was adapted to a format compliant to gimmemotifs, and subsequently used for running ANANSE.

Next, we took our database of OCRs (the output peak file from cellranger) and re-centered the coordinates of the OCRs around the summit of the ATAC-seq signal, for optimal results with gimmemotifs.

In parallel to these steps, the input data for ANANSE must be prepared. Briefly, ANANSE runs on two steps:

(i) the "binding" step where a TF x region matrix of predicted binding activity is computed using the TF/motif annotation and the BAM file, effectively generating a profile of TF binding activity on every region of open chromatin in the dataset;

(ii) the "network" step where the binding information is leveraged with gene expression data to generate the TF-target gene predicted interactions.

To run these steps for each cell type, we computationally dissected the scRNA-seq and the scATAC-seq data to separate the signal from each cell type (Figure 4A). Since the lowest resolution of the data comes from the scATAC-seq, we used the eleven broad cell types as defined by the scATAC-seq data. These are neoblasts, early epidermal progenitors (also referred in the manuscript and figures as "early epid. prog.", or "eep"), late epidermal progenitors (also referred in the manuscript and figures as "late epid. prog.", or "lep"), epidermis, phagocytes, basal/goblet cells, muscle, neurons, parenchyma, protonephridia, and secretory.

(i) First, as explained in Methods, we split the scATAC-seq BAM file generated by `cellranger-atac` [23] using the software `sinto` (https://github.com/timoast/sinto) and the list of cell barcodes belonging to each of the eleven broad types (which we know because of our scATAC-seq analysis and the because of the alignment of the scATAC-seq and the scRNA-seq data) to generate independent BAM files for each of the eleven broad types. These BAM files were later used as input for ANANSE binding.

(ii) Second, we performed a pseudobulk computational dissection on the scRNA-seq dataset at the broad type level, to generate eleven tables of counts of RNA, one per broad cell type. These tables of counts were later used as input for ANANSE network.

We then proceeded to run ANANSE binding for each of the eleven broad types, as explained in Methods. The resulting binding profiles were used alongside the counts tables to generate eleven networks of interactions.

For a preliminary assessment of these networks, we ported these networks to R and igraph. Since ANANSE's workflow scores every putative TF-target gene interaction based on a rank of probability, one must prune the network from gene-gene interactions based on a threshold. For every network, we did a pre-processing by keeping TF-target gene interactions with score values above 0.8, and any gene without neighbours (degree == 0) was removed.

Next, as explained in Methods, we calculated centrality, out-centrality, in-degree, and out-degree for every gene on every network. Relative out-degree was calculated as in-degree divided by the sum of in-degree and out-degree. We calculated the number of active TFs as the number of genes with outdegree above 0.

Then, for every network, we extracted the centrality values for all the TF genes in the network. We did so by running the function `closeness()` from igraph [15] with `method = "out"`. Lastly, we collapsed these values together in a TF x cell type network matrix of centrality values (Supplementary Figure 10C). To assess the degree of similarity of these cell types based on the profile of TF centrality, we clustered the columns of this matrix using the following parameters: `method = "ward.D2"`. With this we observed, as explained in Results, that the epidermis networks group together which provides support to our analyses. Interestingly, we also observed that gut and parenchyma networks group together, which aligns with some of our previous analyses. Overall, we interpreted these observations as confirming the validity of the methodology.

With this, we proceeded to run ANANSE influence, the third step. ANANSE influence takes as input two networks, A and B, and a list of differentially expressed genes between conditions A and B. By default, ANANSE network focuses on up-regulation from a condition to the other, which leaves out genes downregulated in B relative to A. To generate the list of differentially expressed genes (DEGs), we ran our pseudobulk DGE analysis comparing each of the ten non-neoblast broad types against neoblast (Supplementary Figure 11A), as explained in Methods. In particular, we excluded late epidermal progenitors as this is an intermediate state. Thus, we retrieved nine lists of DEGs, one for each transition from neoblast to a non-pluripotent cell type. We called these transitions “fates” in the manuscript.

For each fate, we provided ANANSE influence with the neoblast network, the relevant non-neoblast network, and the list of DEGs in the comparison of the relevant non-neoblast broad type against neoblasts. ANANSE influence focuses on a top number of interactions based on the probability score, which we set to top 250,000, and computes an influence score based on the differential gene expression analysis. The resulting output comprises a list of top influential TFs (visualised in Supplementary Figure 11B-J, Supplementary File 14) as well as a “differential network” –containing the influential TFs and their differentially expressed target genes.

We ported these differential networks to R and generated graphs out of them. For the sake of visualisation only, we pruned these networks and kept the top two interactions per TF (Figure 4C-K). In this same figure, we coloured the TFs based on the relative outdegree in their differential network. We then crossed our differential network data with gene annotation from the planarian literature (using PlanMine [24] to label target genes, and visualised them using ggplot2 (Supplementary Files 15, 16, 17).

We observed that several TFs appeared as influential in more than one cell type (Figure 4C-L, Supplementary File 16). To assess the degree of co-influence of each TF, we collapsed the influence scores of each TF on each fate, generating a TF x fate matrix of influence scores. We subjected this matrix to hierarchical clustering and cut the clustering tree very granularly to detect small groups of co-influential TFs (Supplementary Figure 11K). For each of these clusters, we applied sorting following the same approach as previously described, we computed the average influence profile across fates and correlated every TF of the same cluster against that average influence profile (Supplementary Figure 11L,M). We selected the top five highly-correlating genes per cluster as the top co-influential per group (Figure 4M).

In addition to the probability score, ANANSE also reports the individual parameters estimated for each TF-target interaction, of which weighted binding is particularly interesting. In Figure 6, we show that differentially expressed genes of the *hnf4*i knock-down have higher weighted binding of the *hnf4* TF than non-DEGs (Figure 6E).

Overall, these analyses show that our data recapitulates known dynamics related to cell type differentiation in *S. mediterranea*. Supported by further validation (Figure 5, Figure 6, see Results), we showed that analyses have the potential of generating effective hypotheses about interactions driving cell fate specification in *S. mediterranea*.

#### Supplementary Figures

**Expression of markers from (Emili et al., 2023) in the scRNA-seq dataset from this study**

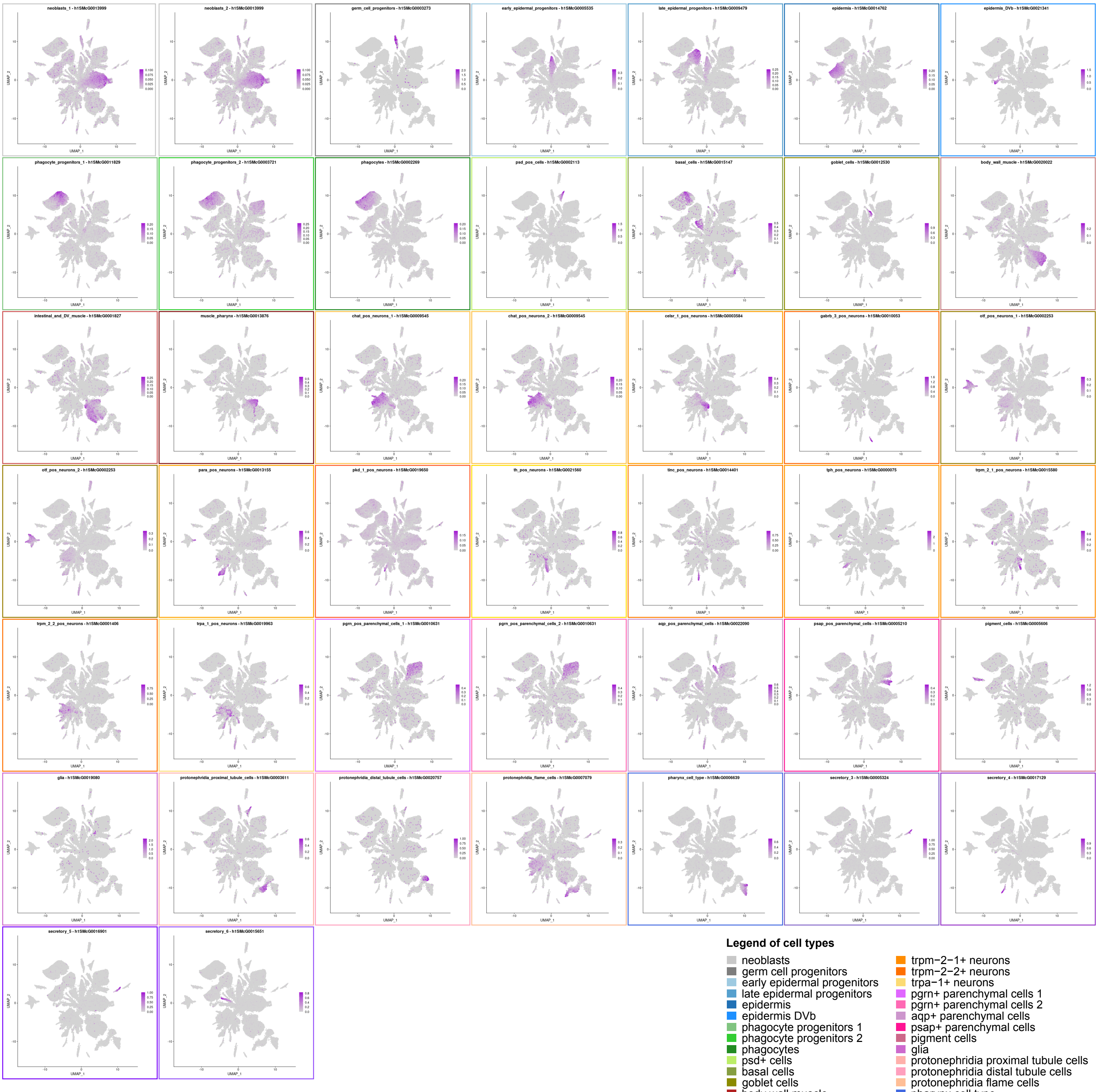

**Legend of cell types**

- neoblasts
- germ cell progenitors
- early epidermal progenitors
- late epidermal progenitors
- epidermis
- epidermis DVb
- phagocyte progenitors 1
- phagocyte progenitors 2
- phagocytes
- psd+ cells
- basal cells
- goblet cells
- body wall muscle
- intestinal and DV muscle
- muscle pharynx
- ChAT+ neurons
- celsr−1+ neurons
- gabrb−3+ neurons
- otf+ neurons 1
- otf+ neurons 2
- para+ neurons
- pkd−1+ neurons
- th+ neurons
- tinc+ neurons
- tph+ neurons
- trpm−2−1+ neurons
- trpm−2−2+ neurons
- trpa−1+ neurons
- pgrn+ parenchymal cells 1
- pgrn+ parenchymal cells 2
- aqp+ parenchymal cells
- psap+ parenchymal cells
- pigment cells
- glia
- protonephridia proximal tubule cells
- protonephridia distal tubule cells
- protonephridia flame cells
- pharynx cell type
- secretory 1
- secretory 2
- secretory 3
- secretory 4
- secretory 5
- secretory 6
- secretory 7

**Expression of marker genes of cell types that did not cluster together in our analysis**

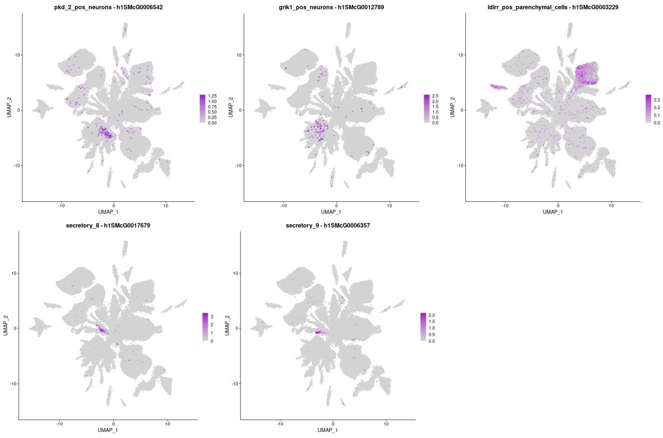

Perez-Posada et al 2024, Supplementary Figure 1

**Supplementary Figure 1.** Expression of cell markers from Emili et al., 2023 in our scRNA dataset.

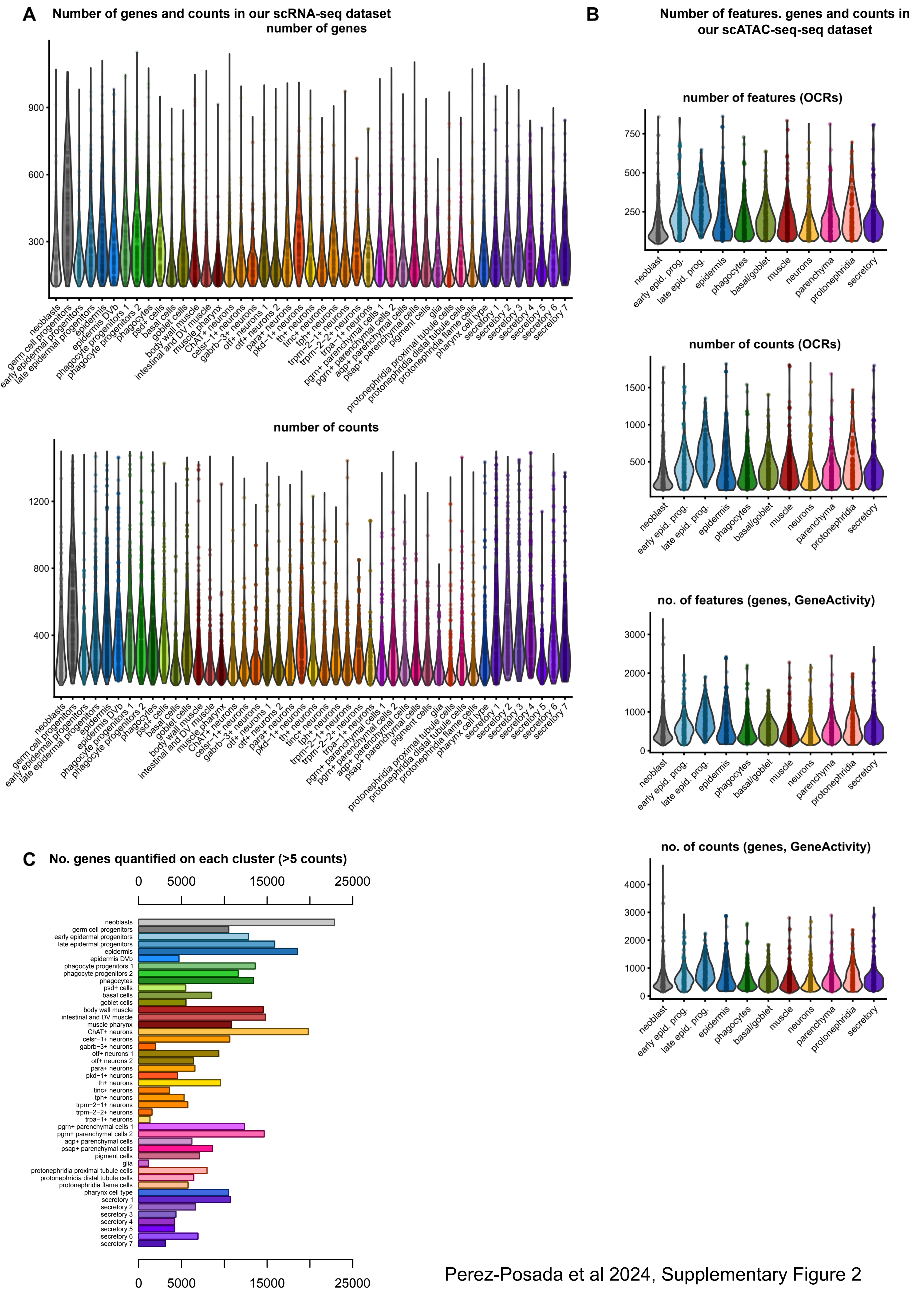


Perez-Posada et al 2024, Supplementary Figure 2

**Supplementary Figure 2.** A: Violin plots showing (top) number of genes and (bottom) number of counts per cell on each of the detected cell types in our dataset. B: From top to bottom: violin plots showing the number of OCRs and number of counts per cell, and the number of genes and Gene Activity counts per cell on the scATAC-seq dataset. C: Barplot showing number of genes quantified on each cluster (above 5 counts). In all panels: points, data points.

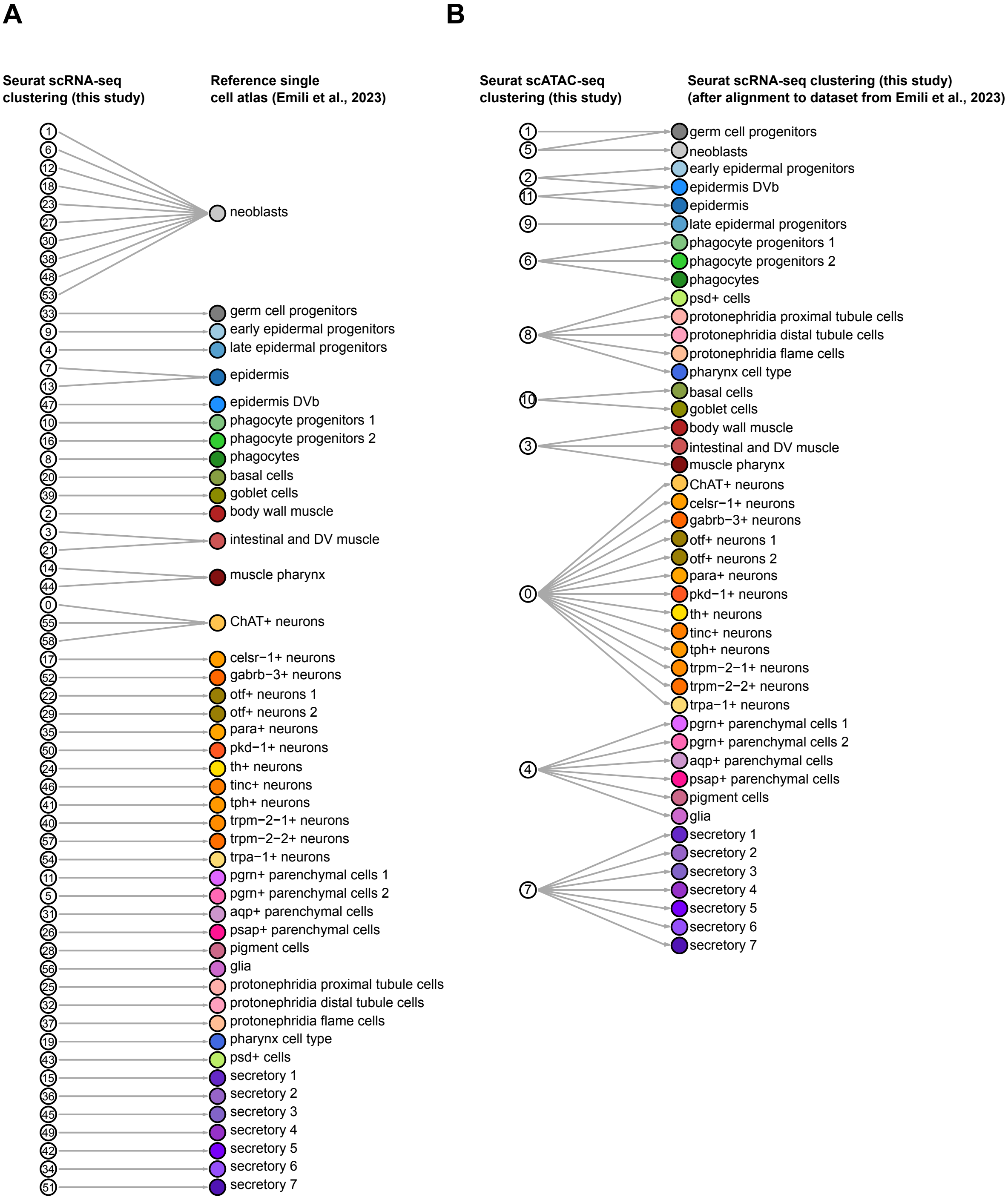


Perez-Posada et al 2024, Supplementary Figure 3

**Supplementary Figure 3.** A: Bipartite graph showing the transfer of labels from the reference dataset to our scRNA-seq dataset. B: Bipartite graph showing the transfer of labels from the scATAC-seq to our scRNA-seq dataset, after aligning to reference scRNA-seq dataset and transferring the cell type labels.

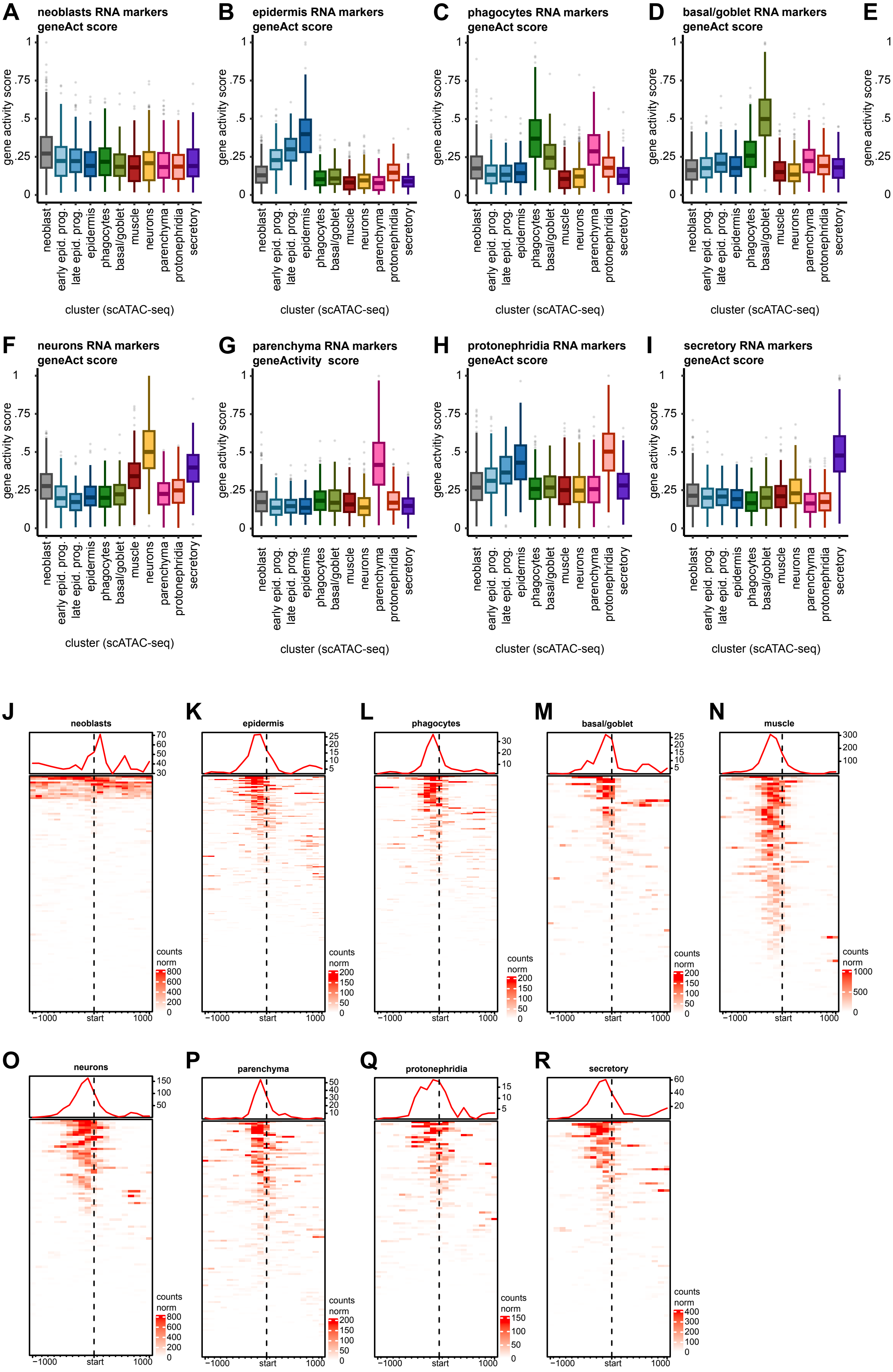


Perez-Posada et al 2024, Supplementary Figure 4

**Supplementary Figure 4.** A-I: Boxplots showing gene activity score of the scRNA-seq markers on each of the broad cell types of the scATAC-seq dataset. Center line, median; box limits, upper and lower quartiles; whiskers, 1.5x interquartile range; points, outliers. J-R: Chromatin accessibility profile of scRNA-seq markers on each of the broad cell types of the scATAC-seq dataset.

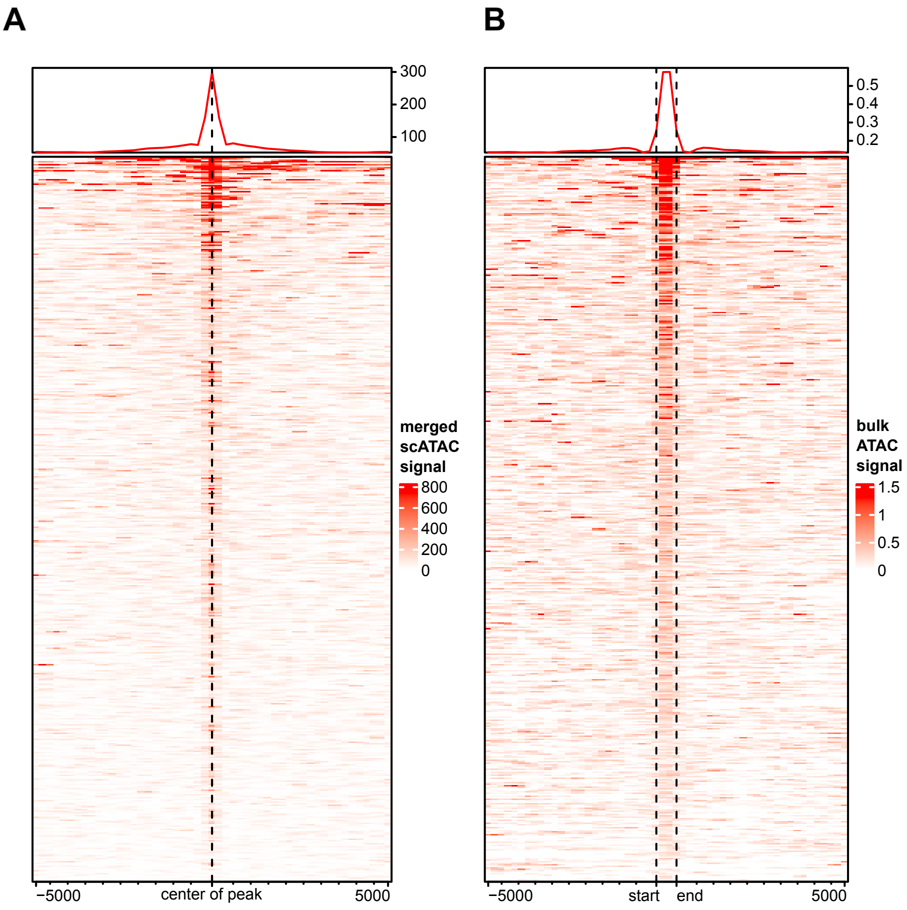


Perez-Posada et al 2024, Supplementary Figure 5

**Supplementary Figure 5.** A: Chromatin profile of the OCRs detected in bulk ATAC seq using the mapping data from the scATAC-seq. B: Chromatin profile of the OCRs detected in scATAC-seq using the mapping data from the bulk ATAC-seq.

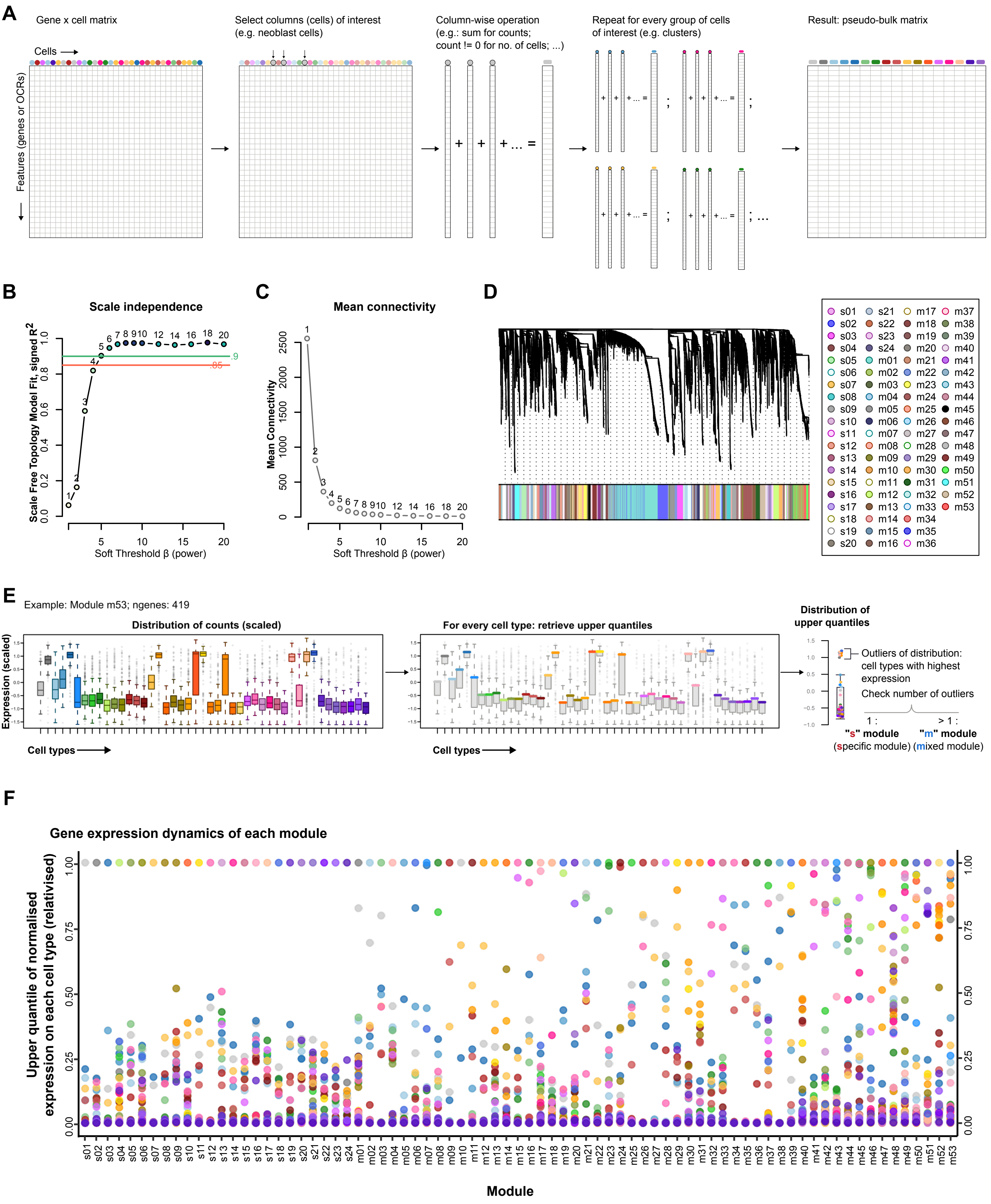


Perez-Posada et al 2024, Supplementary Figure 6

**Supplementary Figure 6.** A: Schematic of the pseudobulk computational dissection without replicates or conditions. B: Scale-free topology model fit of different networks obtained from raising the gene-wise correlations of the pseudobulk data raised to fifteen different soft power thresholds. C: Mean connectivity of different networks obtained from raising the gene-wise correlations of the pseudobulk data to fifteen different soft power thresholds. D: (Top) gene tree of topology overlap and (bottom) module association. E: Schematics of the rationale for sorting modules based on upper quartile values of expression across cell types. F: Distribution of relativised upper quartiles of normalised expression on each cell type per module.

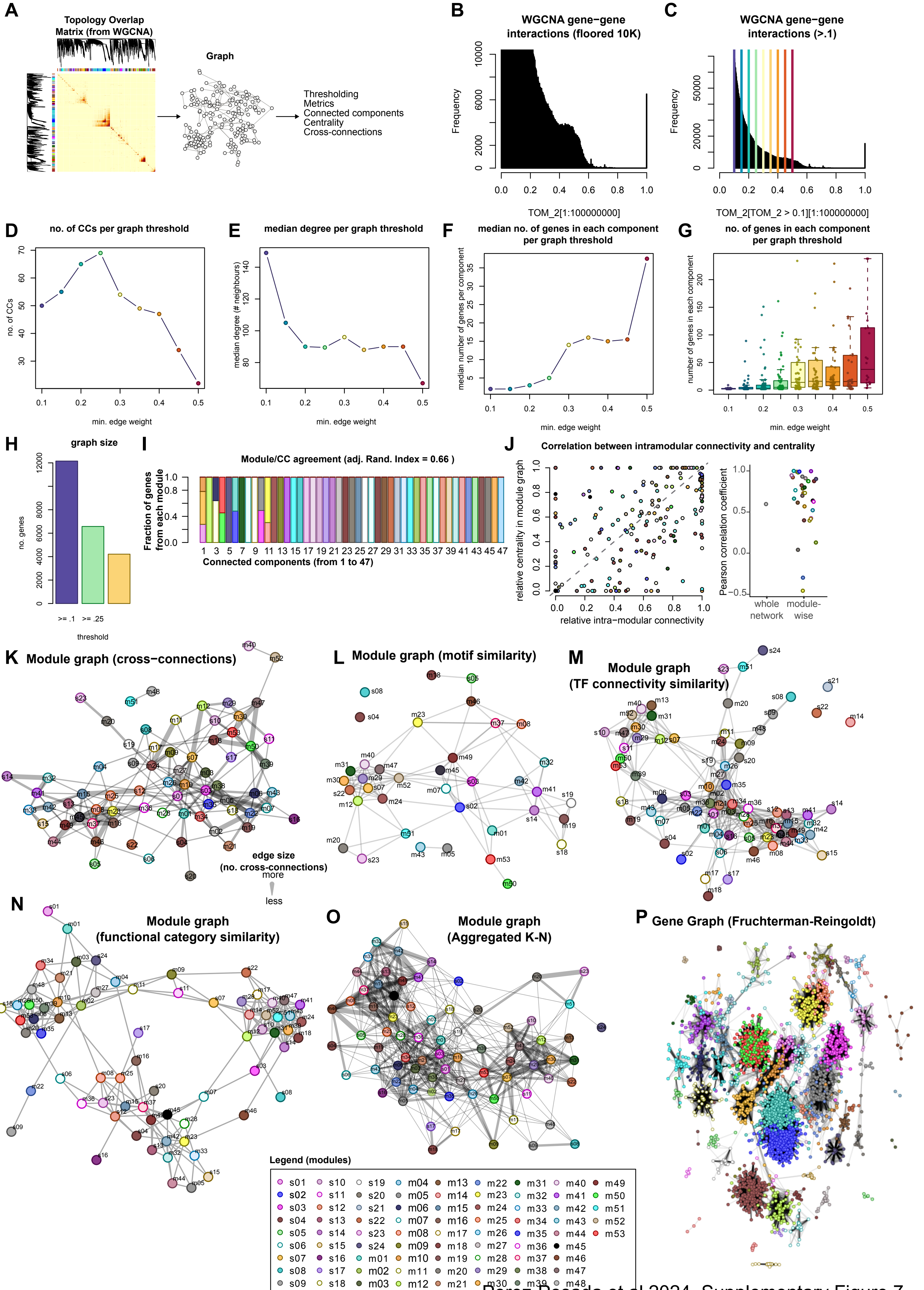


Perez-Posada et al 2024, Supplementary Figure 7

**Supplementary Figure 7.** A: Schematics of graph analysis. B: Distribution of gene-gene weighted correlation values of the whole co-expression WGCNA data. X axis, values of TOM matrix. Y axis, number of edges. Y axis is floored to 10,000. C: Distribution of gene-gene weighted correlation values above 0.1. Coloured lines indicate candidate thresholds for downstream graph analysis. D: Number of connected components per thresholds. E: Median degree (median number of neighbours per gene) per thresholds. F: Median number of genes in each connected component per threshold. G: Boxplot of number of genes in each connected component per threshold. Center line, median; box limits, upper and lower quartiles; whiskers, 1.5x interquartile range; points, all data points. H: Number of genes on each graph per graph threshold. I: Stacked barplot of fractions of genes from each WGCNA module per connected component. J: Correlation between intramodular TF intra-modular connectivity and TF centrality. (Left) scatter plot of relative TF intra-modular connectivity and relative TF centrality. Every dot is a TF. Colour code indicates module membership. (Right) jitter plot of Pearson correlation between TF connectivity and TF centrality all together without considering modules, or module-wise. Every dot is a module (or the whole network for the dot classified as "whole network"). K: Module-wise graph connecting modules (nodes) based on the number of genes between the two with high weighted correlation. Edge thickness indicates the number of cross connections. L: Module-wise graph connecting modules based on Spearman correlation of motif enrichment. M: Module-wise graph connecting modules based on similarity of functional category (COG) enrichment. N: Module-wise graph connecting modules based on similarity of TF connectivity. O: Aggregated module-wise graph, where edge thickness indicates number of times a pair of modules was connected in the aforementioned module-wise graphs. P: Fruchterman-Reingold projection of the WGCNA graph, values >0.35. Every dot is a gene. Colour indicates module membership.

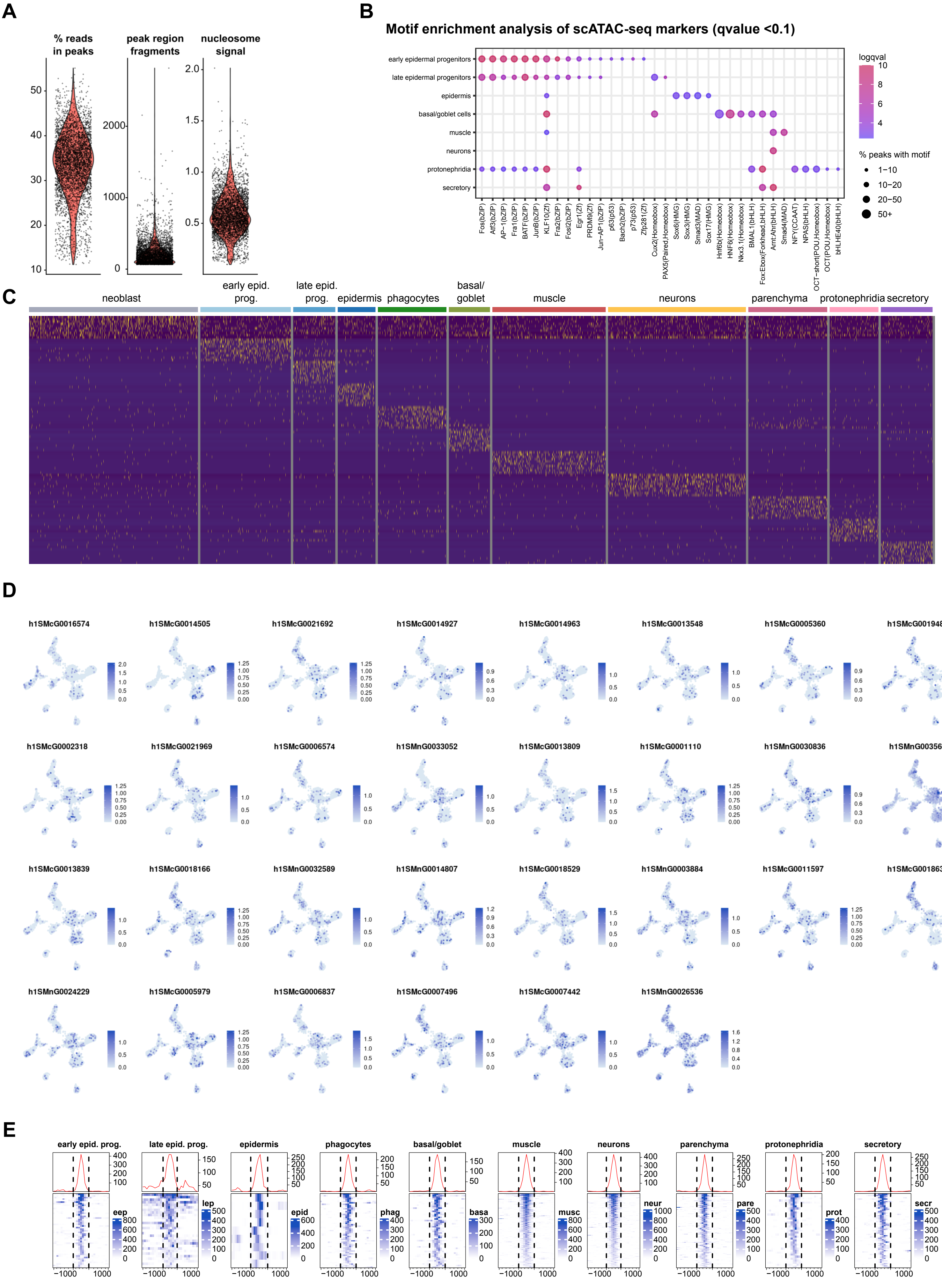


Perez-Posada et al 2024, Supplementary Figure 8

**Supplementary Figure 8.** A: Basic features of the scATAC-seq dataset. (Left) percentage of reads in OCRs, (middle) OCR region fragments, and (right) nucleosome signal per cell. B: Motif enrichment analysis of markers from the scATAC-seq data. C: Heatmap showing the scATAC-seq markers of all the broad cell types in the scATAC-seq data. D: Feature plots of the gene activity of the top fifty markers of neoblasts from the scRNA-seq dataset, in the scATAC-seq dataset. E: Chromatin profile of the cell type-specific (one-vs-all) differential OCRs.

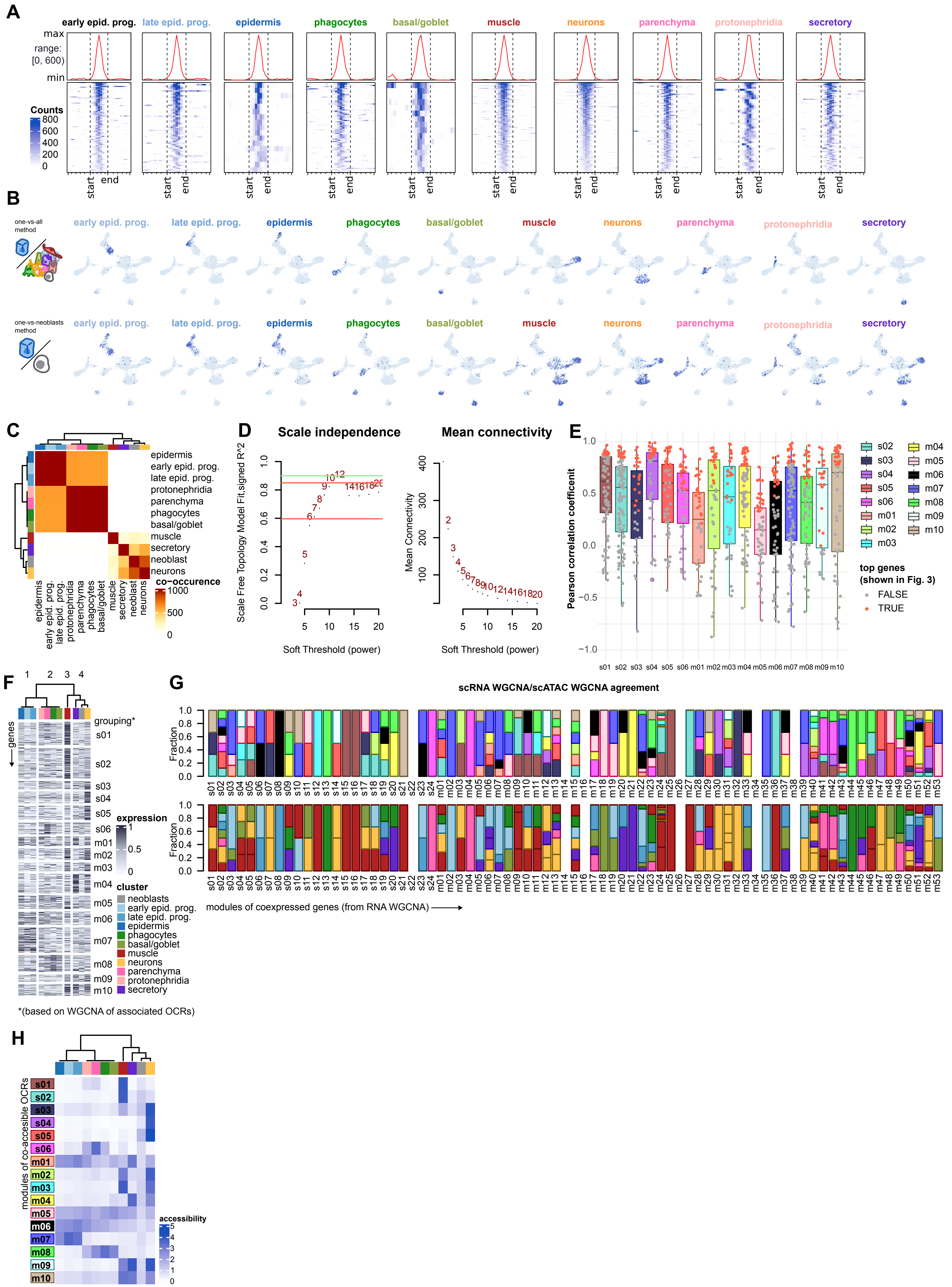


Perez-Posada et al 2024, Supplementary Figure 9

**Supplementary Figure 9.** A: Chromatin plot of the differentiated cell type (one-versus-neoblast) differentially accessible OCRs. B: (top) Example feature plots of cell type-specific (one-versus-all) differentially accessible OCRs; (bottom) Example feature plots of differentiated cell type (one-versus-neoblast) differentially accessible OCRs. C: Cell type similarity based on co-occurrency (Pearson correlation with bootstrapping) of OCR accessibility across all differentiated cell type (one-vs-neoblast) differential OCRs. D: (Left) scale-free topology model fit of different networks obtained from raising the gene-wise correlations of the pseudobulk scATAC-seq data (one-vs-neoblast OCRs ) to fifteen different soft power thresholds. (Right) Mean connectivity of different networks obtained from raising the gene-wise correlations of the pseudobulk scATAC-seq data (one-vs-neoblast OCRs ) to fifteen different soft power thresholds. E: Box plot showing correlation of genes to the co-accessibility profile of modules from their associated OCRs. Every box corresponds to a co-accessibility module. Every point is a gene. Genes labelled in grey do not appear in Figure 3G. Genes labelled in red do. Center line, median; box limits, upper and lower quartiles; whiskers, 1.5x interquartile range; points, data points. F: Heatmap showing gene expression profile of all the genes associated with the differentiated cell type (one-vs-neoblast) OCRs. Genes have been sorted and arranged based on the co-accessibility modules of their associated OCRs. Genes in rows, cell types in columns. Columns have been clustered as in B. G: (Top) stacked barplot of fractions of genes associated to OCRs from each module, for each of the co-expression WGCNA modules of the scRNA dataset. Colour indicates co-accessibility module membership. (Bottom) same as above, but colour indicates cell type of highest average accessibility of a given module. H: Heatmap of average OCR accessibility of each module of co-accessibility. Modules in rows, cell types in columns. Columns have been clustered as in B.

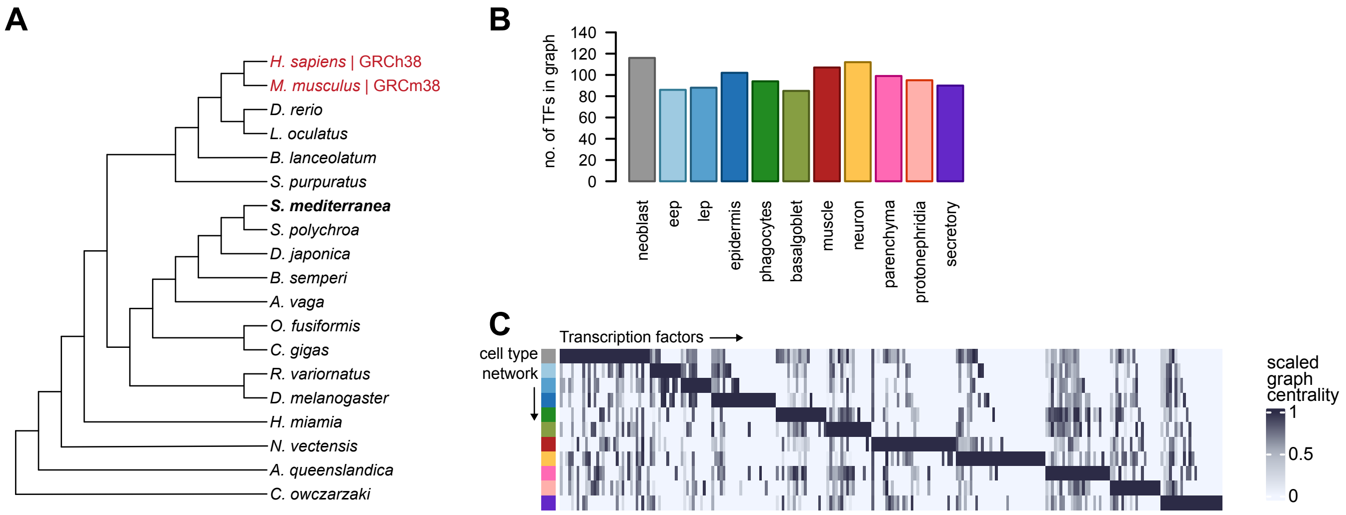


Perez-Posada et al 2024, Supplementary Figure 10

**Supplementary Figure 10.** A: Tree of species used to transfer TF and motif annotation to *S. mediterranea* (bold) from a set of reference species (red) using automated orthology. B: Number of active TFs in each of the constructed graphs. C: Heatmap of TF centrality in each of the graphs. TFs in columns, cell type graphs in rows. TFs have been sorted by highest value of centrality.

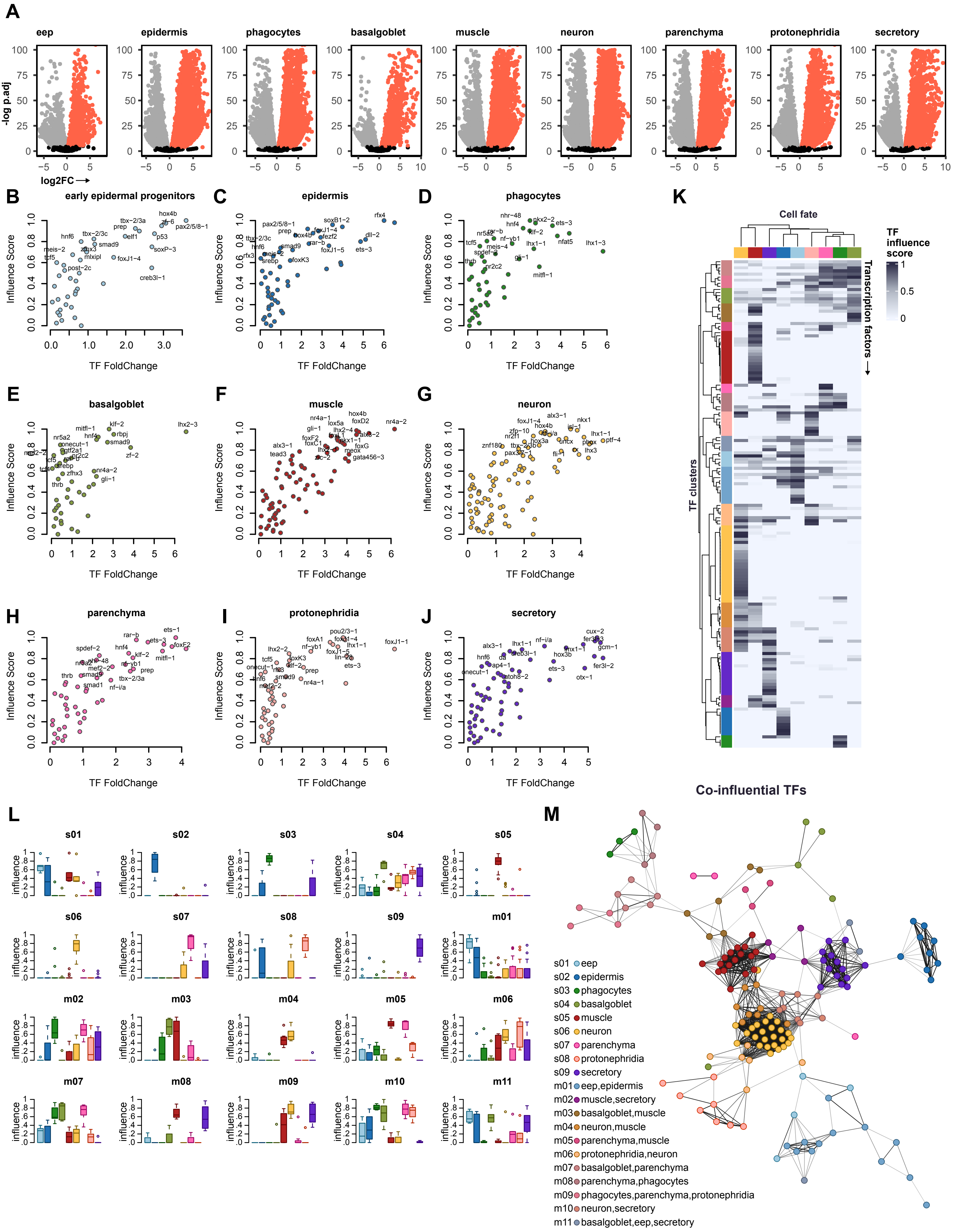


Perez-Posada et al 2024, Supplementary Figure 11

**Supplementary Figure 11.** A: Volcano plots showing log fold change (x-axis) and -log p. adjusted (y axis) of each of the differentiated cell type (one-vs-neoblasts) differential gene expression analysis, used as input for ANANSE influence. B-J: Scatter plot of TF fold change and influence score for the top factors of each transition from neoblast to a non-pluripotent cell type: (B) early epidermal progenitors, (C) epidermis, (D) phagocytes, (E) basal/goblet, (F) muscle, (G) neurons, (H) parenchyma, (I) protonephridia, (J) secretory. Labels shown for the 20 TFs with highest influence score. K: Heatmap of TF influence score across cell fates. TFs in columns, cell fates in rows. Top tree: hierarchical clustering of TFs based on their co-influence across cell fates. L: Boxplots of influence score profiles of each of the clusters of co-influential TFs. Center line, median; box limits, upper and lower quartiles; whiskers, 1.5x interquartile range; points, outliers. M: Graph of Pearson correlation between co-influential TFs.

***hnf4*i RNAi phenotype progression**

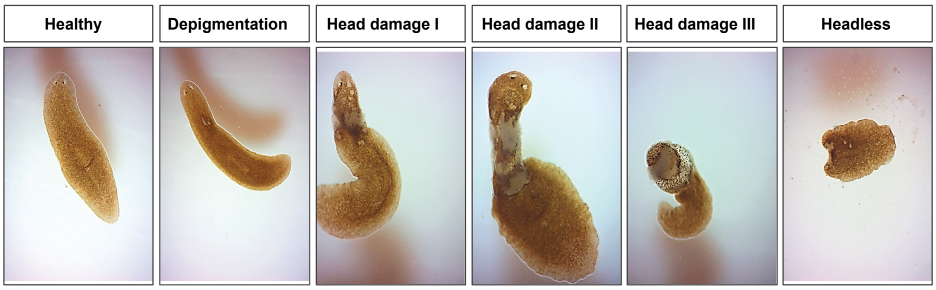


Perez-Posada et al 2024, Supplementary Figure 12

**Supplementary Figure 12.** Phenotype progression of the *hnf4*i RNAi knock-down in *Smichdtea mediterranea*.

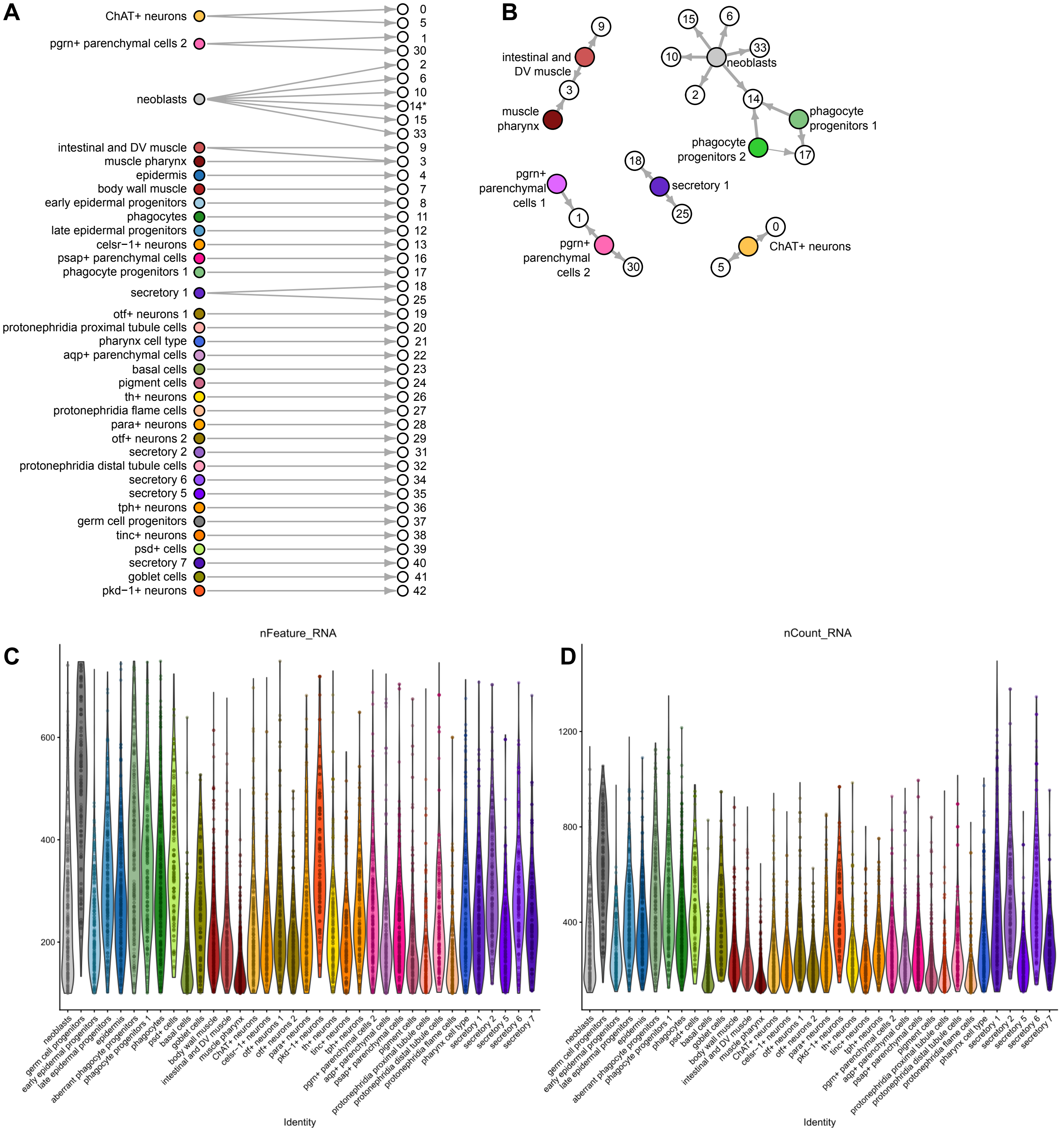


Perez-Posada et al 2024, Supplementary Figure 13

**Supplementary Figure 13.** A: Bipartite graph showing the transfer of labels from our whole scRNA-seq dataset to the *hnf4*i dataset. B: Graph showing which labels of our whole scRNA-seq dataset are received by more than one cluster from the *hnf4*i dataset. Of these, only cluster 14 contain cells that receive labels from more than one broad cell type (neoblasts and phagocyte progenitors). C: Violin plot of number of genes per cell on each cell type of the *hnf4*i dataset. D: Violin plot of number of counts per cell on each cell type of the *hnf4*i dataset. Points, data points.

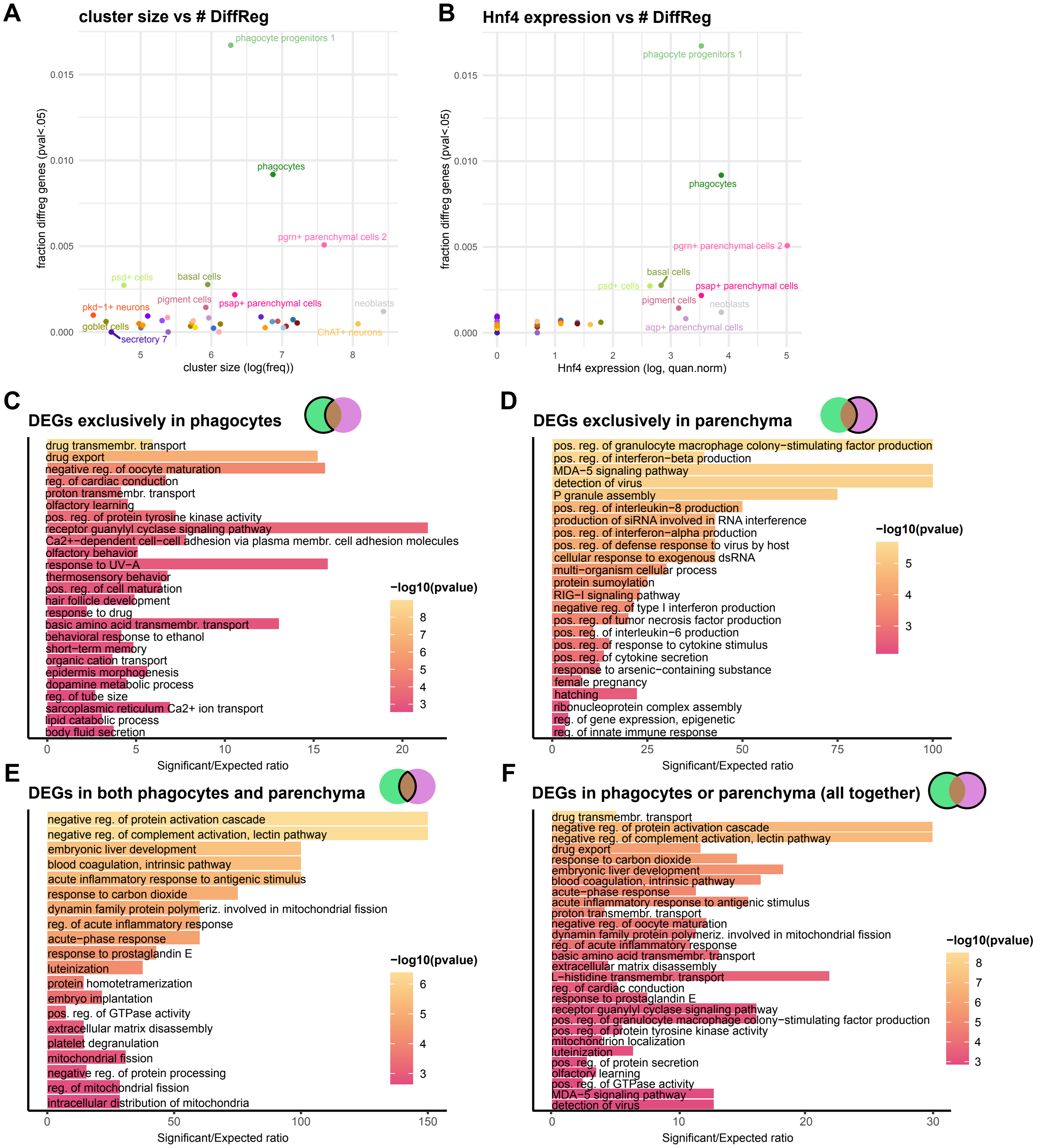


Perez-Posada et al 2024, Supplementary Figure 14

**Supplementary Figure 14.** A: Scatter plot showing the number of differentially expressed genes on each cell type (DEGs) in relation to cell type cluster size (number of cells). B: Scatter plot showing the number of DEGs on each cell type in relation to the level of expression of *hnf4* on each cell type. C-F: Bar plots of Gene Ontology Enrichment for the phagocytes-exclusive (C), parenchyma-exclusive (D), common (E) or all (F) DEGs.

#### Supplementary Files

**Supplementary File 1.** Details on sequencing libraries used in this study.

**Supplementary File 2.** Gene and functional annotation used in this study, including transcription factor annotation and WGCNA module membership.

**Supplementary File 3.** Cell type annotation of Seurat clusters of the main scRNA-seq dataset in this study.

**Supplementary File 4.** Table of modules detected by WGCNA in the main scRNA-seq dataset in this study.

**Supplementary File 5.** Boxplots of normalised gene counts on each cell type, for each WGCNA module of gene co-expression. Center line, median; box limits, upper and lower quartiles; whiskers, 1.5x interquartile range; points, outliers.

**Supplementary File 6.** Feature plots of thirty genes per WGCNA module of gene co-expression, randomly sampled for each module.

**Supplementary File 7.** Barplots of Gene Ontology enrichment of each WGCNA module of gene co-expression.

**Supplementary File 8.** Motif enrichment analysis on the promoters of the genes from each WGCNA module of gene co-expression.

**Supplementary File 9.** Lists of differential regions of open chromatin (OCRs) between every differentiated cell type and the rest, in a one-versus-all manner.

**Supplementary File 10.** Lists of differential regions of open chromatin (OCRs) between every differentiated cell type and neoblasts.

**Supplementary File 11.** List of OCR/module associations resulting from running WGCNA on one-versus-neoblasts OCRs.

**Supplementary File 12.** List of OCRs associated with their closest gene, with module membership of OCRs and correlation values between the expression of every gene and their associated OCRs.

**Supplementary File 13.** List of species used in orthology transferring of TFs using gimmemotifs motif2factors.

**Supplementary File 14.** List of outputs of ANANSE influence between the network of every differentiated cell type and the neoblast network.

**Supplementary File 15.** Top targets of five example TFs of the differential networks when comparing neoblasts to every differentiated cell type. From top row to bottom row, and following colour code as in the rest of the manuscript: early epidermal progenitors, epidermis, phagocytes, basal/goblet cells, muscle cells, neurons, parenchymal cells, protonephridia cells, and secretory cells.

**Supplementary File 16.** Heatmap showing presence/absence of putative predicted interaction between the top influential TFs of each fate and the targets with an identified gene symbol in the literature.

**Supplementary File 17.** Table of the putative predicted interactions between the top influential TFs of each fate and the targets with an identified gene symbol in the literature.

**Supplementary File 18.** List of influential TFs and their associated set of co-influence.

**Supplementary File 19.** Cell type annotation of Seurat clusters of the main scRNA-seq dataset in this study, including number of cells from each sample for each cluster.

**Supplementary File 20.** List of gene markers used to calculate the neoblast gene score and the phagocyte gene score.

**Supplementary File 21.** List of Differentially Expressed Genes (DEGs) for all broad cell types when comparing broad cell types of control and *hnf4*i samples.
